# Glutamine antagonism suppresses tumor growth in adrenocortical carcinoma through inhibition of de novo nucleotide biosynthesis

**DOI:** 10.1101/2025.09.28.674326

**Authors:** Vasileios Chortis, Kleiton Silva Borges, Cong-Hui Yao, Claudio Ribeiro, Luis Fernando Nagano, Mesut Berber, Alessandro Prete, Lukáš Najdekr, Michail E. Klontzas, Andris Jankevics, Pedro Vendramini, Jean Lucas Kremer, Liam Kelley, Sathuwarman Raveenthiraraj, Alcides Euzebio Tavares Xavier, Stylianos Tsagarakis, Magdalena Macech, Ivana D. Pupovac, Thomas G Papathomas, Betul Haykir, Catherine Winder, Marcus Quinkler, M. Conall Dennedy, Grethe Å. Ueland, Felix Beuschlein, Antoine Tabarin, Martin Fassnacht, Angela E. Taylor, Darko Kastelan, Urszula Ambroziak, Dimitra A. Vassiliadi, Margaret E. Wierman, Katja Kiseljak-Vassiliades, Irina Bancos, Diana L. Carlone, Warwick B. Dunn, Wiebke Arlt, Marcia C. Haigis, David T. Breault

## Abstract

Dysregulation of cellular metabolism is a hallmark of cancer, yet metabolic dependencies in adrenocortical carcinoma (ACC) remain poorly defined and effective systemic therapies are lacking. Here, we dissected ACC metabolism by integrating transcriptional profiling from human and mouse ACC, targeted tissue metabolomics from a genetically engineered mouse ACC model, and untargeted serum metabolomics from a large patient cohort, providing cross-species validation of metabolic rewiring in ACC. This study revealed global metabolic dysregulation, involving glutamine-dependent pathways such as non-essential amino-acid and hexosamine biosynthesis, nucleotide metabolism, and glutathione biosynthesis, suggesting glutamine catabolism is a critical metabolic vulnerability in ACC. Pharmacologic inhibition of glutamine metabolism using the glutamine antagonist 6-Diazo-5-Oxo-L-Norleucine (DON) and its pro-drug JHU-083 elicited robust anti-tumor responses *in vitro* and *in vivo*. Mechanistic studies revealed that the anti-tumor effect of glutamine antagonism was primarily driven by selective inhibition of glutamine-fueled *de novo* nucleotide biosynthesis, leading to nucleotide depletion and DNA damage. The resulting metabolic stress is associated with increased sensitivity to inhibition of the DNA damage response, particularly through pharmacological targeting of Ataxia telangiectasia and Rad3-related (ATR) signaling, resulting in synergistic anti-tumor effects. Collectively, our study identifies glutamine-dependent nucleotide biosynthesis as a therapeutically exploitable metabolic vulnerability in ACC and provides a rationale for combining glutamine antagonism with DNA damage response–targeted therapies.

**Highlights:**

- Mouse and human ACC share conserved, glutamine-dependent metabolic programs.
- Glutamine antagonism potently inhibits ACC tumor growth *in vivo*
- Nucleotide depletion and DNA damage underpin the anti-tumor effect of glutamine metabolism
- Glutamine antagonism synergizes with ATR inhibition in ACC cells

## Introduction

Adrenocortical Carcinoma (ACC) is a rare and aggressive malignancy, originating from the adrenal cortex. Most patients present with, or develop, metastatic disease; for such patients, there are no effective medical treatments and overall survival remains poor with a median survival of less than 15 months with cytotoxic chemotherapy^1,2^. Despite considerable progress in understanding the molecular biology of ACC^3,4^, no effective targeted treatments have yet emerged. Therefore, there is a critical unmet clinical need to develop effective medical treatments for patients with inoperable tumors.

Over the last decade, seminal studies have characterized the molecular landscape of ACC, identifying molecular clusters associated with distinct clinical outcomes^3–5^. This work demonstrated that constitutive activation of the Wnt/β-catenin signaling pathway and cell cycle dysregulation due to TP53/RB1 loss are the two most common driver events in ACC, and are most prevalent in patients with poor prognosis. Notably, concurrent alterations in both pathways are observed in up to 30% of ACC cases^6^. Building on these findings, we recently developed a genetically engineered mouse model of ACC (BPCre), which combines Wnt/β-catenin activation with p53 loss^7^. This model faithfully recapitulates major features of the most aggressive molecular subtype of ACC, including metastasis and active steroidogenesis as well as epigenetic dysregulation^7–9^.

Dysregulation of cellular metabolism is recognized as a hallmark of cancer^10,11^, which supports cancer cells’ specific metabolic needs, including the rapid generation of biomass and energy, and a robust antioxidant defence system. Despite ACC’s unique metabolic trait as an actively steroidogenic tumor, there is limited understanding of how ACC tumors rewire their metabolic network and what diagnostic or therapeutic opportunities this aspect of ACC biology may present.

Here, we present a comprehensive analysis of metabolic dysregulation in ACC by integrating transcriptional profiling from human and mouse ACC, targeted tissue metabolomics from a mouse model of ACC, and untargeted serum metabolomics from a large cohort of patients with adrenal tumors, achieving cross-species validation of metabolic rewiring in ACC. These findings reveal dysregulation in pathways supporting non-essential amino-acid and nucleotide biosynthesis, hexosamine biosynthesis, and antioxidant production, all of which are heavily dependent on glutamine (Gln) as a key metabolic substrate. Given Gln’s central role in fueling anabolic and redox-supporting processes, we used the Gln antagonist 6-diazo-5-oxo-L-norleucine (DON), and its recently developed tumor-selective pro-drug JHU-083^12–14^, to target this putative metabolic dependency in models of ACC. We demonstrate that Gln antagonism is highly effective against ACC using *in vitro* and *in vivo* pre-clinical models. Furthermore, we show that DON’s anti-tumor activity against ACC is specifically contingent on disruption of Gln’s role in *de novo* nucleotide biosynthesis, a process that is up-regulated in the most aggressive human ACCs. Moreover, we discovered that DON-driven nucleotide depletion leads to DNA damage, resulting in potent synergism with inhibition of the DNA damage response pathway. Consistent with these findings, untargeted serum metabolomics from patients with adrenal tumors identified key changes in nucleotide and amino-acid homeostasis in ACC. These pre-clinical findings reveal Gln antagonism as a promising new therapeutic target for this aggressive cancer.

## Results

### Cross-species analysis reveals conserved transcriptional-metabolic programs in ACC

To identify actionable metabolic vulnerabilities in ACC, we sought to identify common up-regulated metabolic programs utilized by mouse and human ACC. To achieve this, we transcriptionally profiled our recently developed BPCre genetically engineered mouse model of ACC^7^ to identify dysregulated metabolic pathways and compared these with pathways also dysregulated in human ACC. First, we performed bulk RNA-sequencing (RNA-seq) on BPCre tumors (*n* = 7 mice, 5 female and 2 male mice, aged 9-12 months) and control mouse adrenals without mutations (*n* = 6 mice, all female, aged 5-11 months). Then, we analyzed bulk RNA-seq data from human ACC tumors using the TCGA-ACC database (*n* = 79)^3^ and normal human adrenals from the GTEx database (*n* = 295)^15^ (**Fig. 1a**). We focused on the expression of 2,754 defined metabolic genes encoding metabolic enzymes and transporters, using an unbiased, standardized transcriptomic approach to identify uniquely dysregulated tumor pathways^16–18^. Comparison of mouse and human ACC tumors revealed a 68% overlap in up-regulated metabolic pathways and a 75% overlap in down-regulated metabolic pathways (**Fig. 1b, Suppl. Fig. 1a, Suppl. Tables 1-2**), underscoring the strong conservation of transcriptional–metabolic programs between species in ACC. Strikingly, all 15 of the most significantly up-regulated pathways in mouse tumors were also up-regulated in human tumors (**Fig. 1c, Suppl. Table 1**), revealing a conserved metabolic signature.

**Fig. 1:**
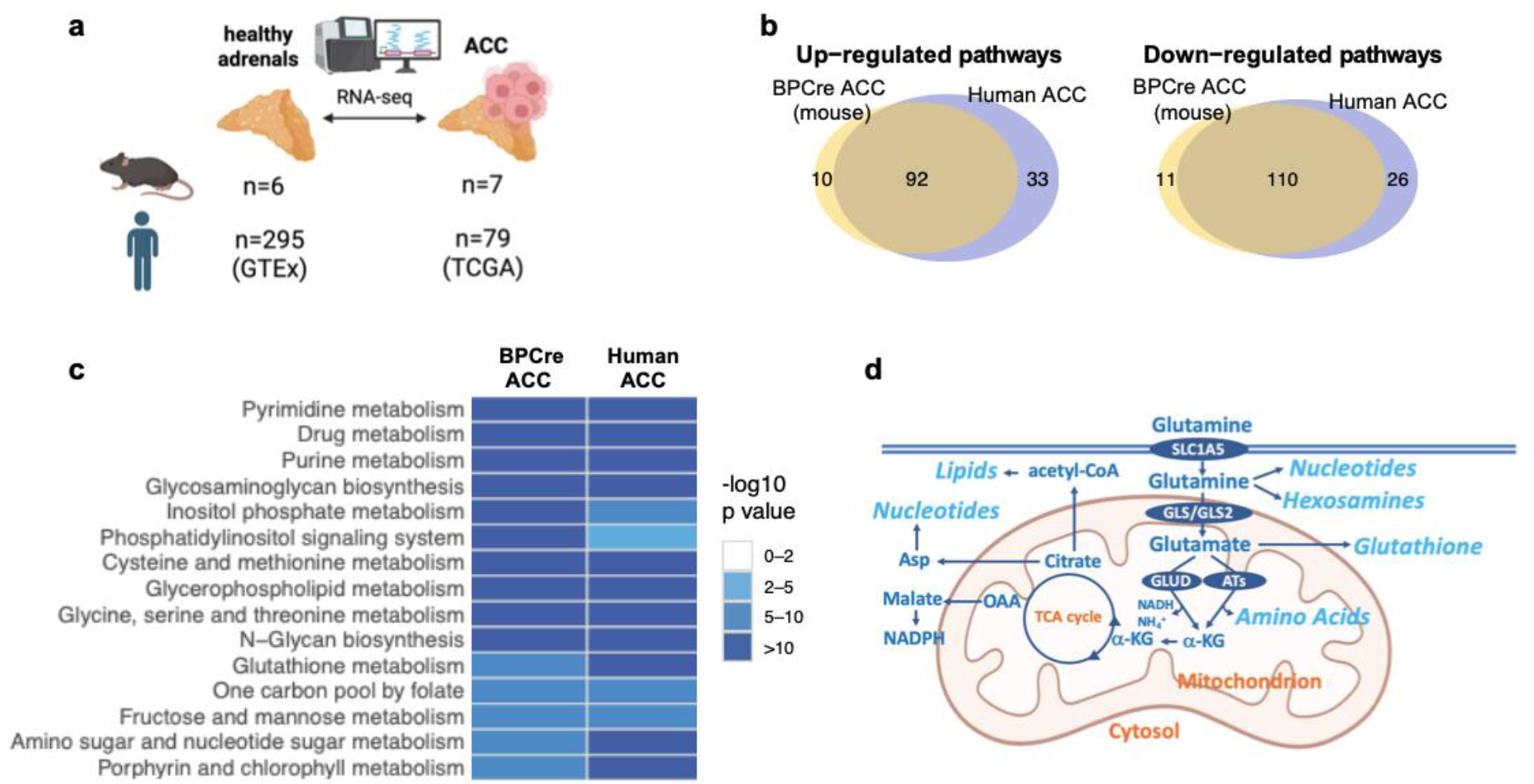
Cross-species transcriptomic analysis of metabolic reprogramming in ACC. **a** Schematic overview of cross-species transcriptomic analysis of metabolic pathways in ACC. **b** Venn diagrams showing overlap between significantly (FDR<0.05) up-regulated and down-regulated metabolic pathways in mouse (BPCre) and human ACC tumors. Numbers refer to dysregulated pathways. **c** Top 15 most significantly up-regulated pathways in mouse ACC; all pathways are also significantly up-regulated in human ACC (TCGA/GTEx samples). **d** Schematic overview of Gln metabolism. α-KG: α-ketoglutarate, Asp: aspartate, ATs: aminotransferases, NADH: reduced nicotinamide adenine dinucleotide, OAA: oxaloacetate.

Dysregulated purine and pyrimidine metabolism was a prominent feature in this analysis, with up-regulation of several genes involved in *de novo* biosynthesis (a Gln-dependent multi-step process synthesizing nucleotides from basic precursors) and salvage biosynthesis (mediating nucleotide recycling) (**Suppl. Fig. 1b,c**). Interestingly, amino sugar and nucleotide sugar metabolism (**Suppl. Fig. 2a**), N-glycan biosynthesis, and glycosaminoglycan biosynthesis were all among the most up-regulated pathways, consistent with an increased requirement for activated sugar donors to support protein glycosylation and glycosaminoglycan biosynthesis. Taken together, these prominent changes indicate an increased demand for Gln, an essential nitrogen donor to both *de novo* nucleotide biosynthesis and amino sugar (hexosamine) biosynthesis (**Fig. 1d**). Additional indications of an increased demand for Gln included i) an up-regulation of transamidases channeling Gln-derived nitrogen to aspartate and asparagine biosynthesis (*GOT2*, *ASNS*) in both species (**Suppl. Table 1-2**) and ii) extensive dysregulation of glutathione metabolism, a major Gln-dependent antioxidant system (**Suppl. Fig. 2b**). One-carbon metabolism, encompassing the folate cycle and serine/glycine metabolism, was also prominently up-regulated, further supporting nucleotide synthesis and redox homeostasis. Overall, this cross-species analysis revealed extensive rewiring of Gln and one-carbon metabolism to support nucleotide and protein biosynthesis, redox balance, and protein glycosylation.

### Gln-dependent metabolic pathways are dysregulated in mouse ACC

To quantify tumor-intrinsic metabolic alterations and to validate the functional consequences of the transcriptional alterations observed by RNA-seq, we performed targeted tissue metabolomics in BPCre tumors from both sexes compared with age-matched control adrenals (lacking mutations), as well as adrenals from BPCre mice prior to tumor formation (BPCre-Pre) (**Fig. 2a**)^7^. Targeted polar metabolomic analysis by liquid chromatography-mass spectrometry (LC-MS), using a library that provides a representative cross-section of core carbon and nitrogen-utilizing metabolic pathways, confirmed extensive metabolic rewiring involving key programs highlighted by the transcriptomic analysis (**Fig. 2b**). We observed a robust increase in the Gln-dependent amino-acids glutamate, aspartate, proline and ornithine, as well as the major antioxidant glutathione and its analogue ophthalmic acid, in ACC tumors (**Fig. 2c**, **Suppl. Table 3**). Elevated levels of nucleotide sugars, including UDP-GlcNAc/UDP-GalNAc and UDP-glucose/UDP-galactose, were consistent with increased availability of activated sugar donors for protein glycosylation and glycan biosynthesis, in keeping with the transcriptional upregulation of hexosamine and nucleotide sugar metabolic pathways. We also identified prominent changes in nucleotide metabolism, particularly related to alterations in the tissue-level abundance of purine salvage/degradation intermediates. Interestingly, there was a depletion in purine metabolites xanthine, xanthosine and uric acid (potentially reflecting increased recycling to meet biosynthetic demands), but an increase in pyrimidine metabolites thymidine, cytidine, and 3-ureidopropionic acid (suggestive of increased turnover/overflow catabolism). Despite transcriptional up-regulation of several enzymes mediating *de novo* purine/pyrimidine biosynthesis, steady-state tissue metabolomics did not show changes in *de novo* intermediates, with the notable exception of the strong depletion of xanthosine monophosphate (XMP), an obligate substrate of the Gln-dependent enzyme GMP synthetase (GMPS). This pattern could be consistent with increased metabolic flux coupled with tight channeling to down-stream processes (DNA/RNA synthesis, hexosamine biosynthesis). In addition to these changes, there was a robust increase in the Gln-dependent co-factors nicotinamide adenine dinucleotide (NAD) and reduced NAD (NADH). Collectively, these findings highlight Gln metabolism as a putative metabolic dependency in ACC. Interestingly, several of these metabolites exhibited progressive changes across the tumorigenic trajectory from normal adrenals to BPCre adrenals pre-malignant transformation to established BPCre ACC tumors, suggestive of involvement in early ACC tumorigenesis (**Fig. 2d, Suppl. Fig. 3a**).

**Fig. 2:**
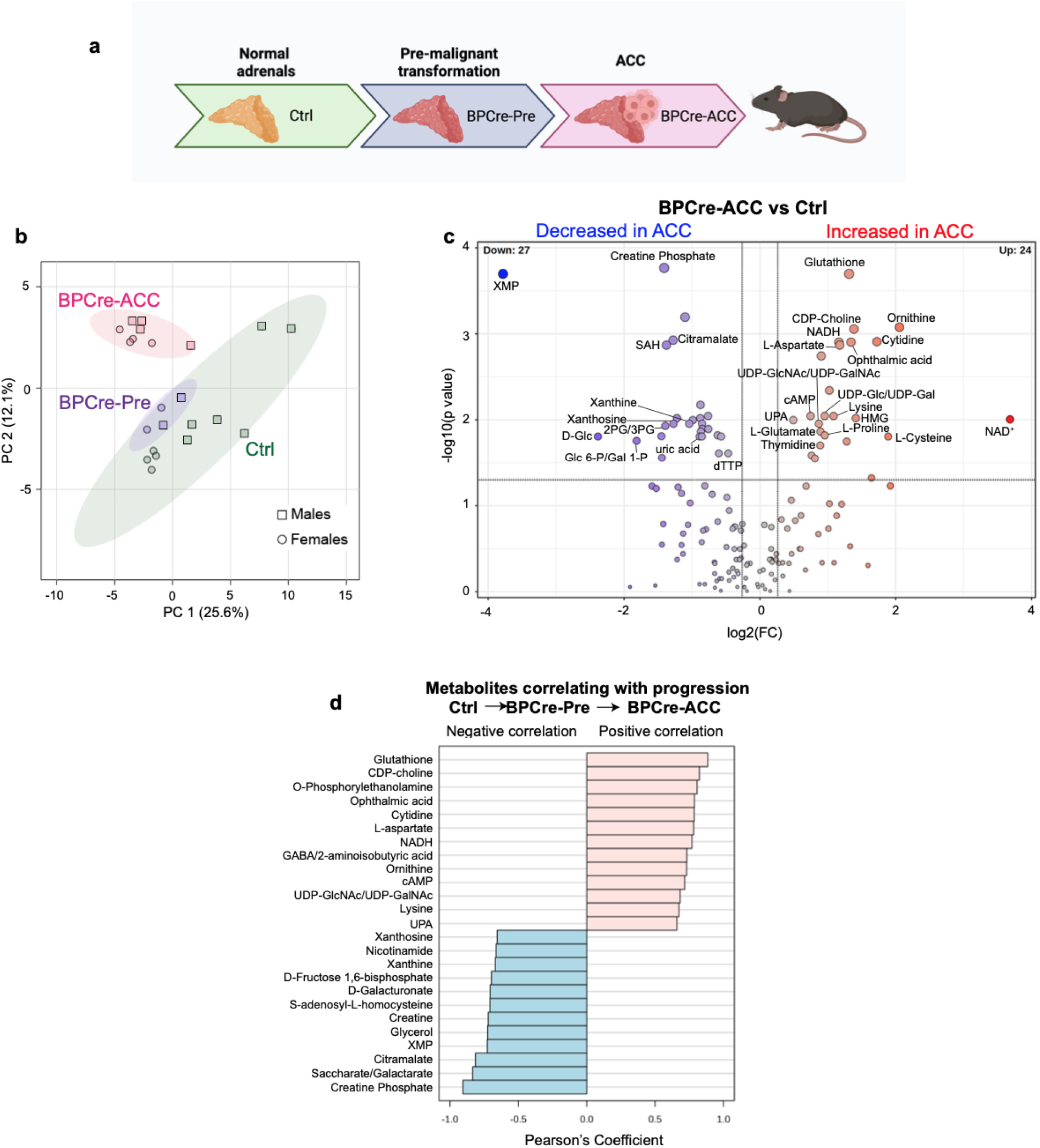
Targeted tissue metabolomics in BPCre tumors. **a** Schematic of malignant transformation in BPCre mice. **b** Principal Component Analysis (PCA) comparing the polar metabolome of control adrenals from C57BL/6J mice (7-11 months, n=10), adrenals from young BPCre mice pre-tumor formation (BPCre-Pre, 4 months, n=4), and ACC tumors from older BPCre mice (BPCre-ACC, 8-13 months, n=8). Circles represent female mice; squares represent male mice. For control and BPCre-Pre adrenals, each sample represents tissue from two adrenals taken from either the same mouse or from two mice of the same genotype and sex. **c** Polar metabolite changes comparing BPCre-ACC vs control adrenal tissue selected by Volcano plot with fold change threshold (x) 1.2 and t-tests threshold (y) 0.05 (FDR-corrected p values). Dashed lines indicate statistical thresholds. **d** Top 20 metabolites showing progressive changes from control adrenals → BPCre-Pre adrenals → BPCre-ACC. Metabolites that could not be unambiguously distinguished are reported using dual annotations. 2PG: 2-phosphoglycerate; 3PG: 3-phosphoglycerate. CDP, cytidine diphosphate; D-Glc: D-glucose; dTTP, deoxythymidine triphosphate; HMG: 3-hydroxy-3-methylglutarate; NAD, oxidized nicotinamide adenine dinucleotide; NADH, reduced nicotinamide adenine dinucleotide; SAH, S-adenosylhomocysteine; UDP, Uridine diphosphate; UDP-GalNAc: UDP-N-Acetylgalctosamine; UDP-GlcNAc: UDP-N-acetylglucosamine; UPA, β-Ureidopropionic acid; XMP, Xanthosine monophosphate.

As additional controls, we compared the metabolomes of adrenals from male and female mice, from young and older mice, and from single-mutant mice. No significant sex-specific differences were observed in tumor metabolomes (**Suppl. Fig. 3b**), although several nucleotide metabolites differed between normal male and female adrenal glands (**Suppl. Fig. 3c**). Additionally, there were no significant differences between control adrenals from younger mice (3-5 months) and older mice (7-10 months) (**Suppl. Fig. 3d**), indicating that metabolic differences between BPCre-Pre adrenals and ACC tumors or older control adrenals are not a function of increasing age. Analysis of adrenals from single-mutant (BCre or PCre) mice revealed only modest changes compared to the complex metabolic rewiring seen in BPCre adrenals prior to tumor formation and older mice with BPCre tumors (**Suppl. Fig. 4a–c**).

### ACC is highly sensitive to disruption of Gln metabolism *in vitro* and *in vivo*

To assess whether Gln metabolism is a druggable metabolic dependency in ACC, we employed the Gln antagonist 6-diazo-5-oxo-L-norleucine (DON)^12–14,19^. To test its efficacy, we treated a panel of ACC cell lines with DON, including a new murine ACC cell line (BCH-ACC3) derived from a BPCre tumor and three human ACC cell lines (NCI-H295R, CU-ACC1, CU-ACC2)^20^ harboring common ACC driver alterations (β-Catenin gain-of-function and/or p53 loss-of-function). The genetic profile and doubling time of each cell line are summarized in **Suppl. Fig. 5a**. All ACC cell lines displayed marked sensitivity to DON, with IC_50_ values ranging from 0.03 to 4.07 μM (**Fig. 3a**). Notably, low micromolar concentrations (2.5–5 μM) were sufficient to fully impede colony formation in clonogenic assays across all cell lines (**Fig. 3b**).

**Fig. 3:**
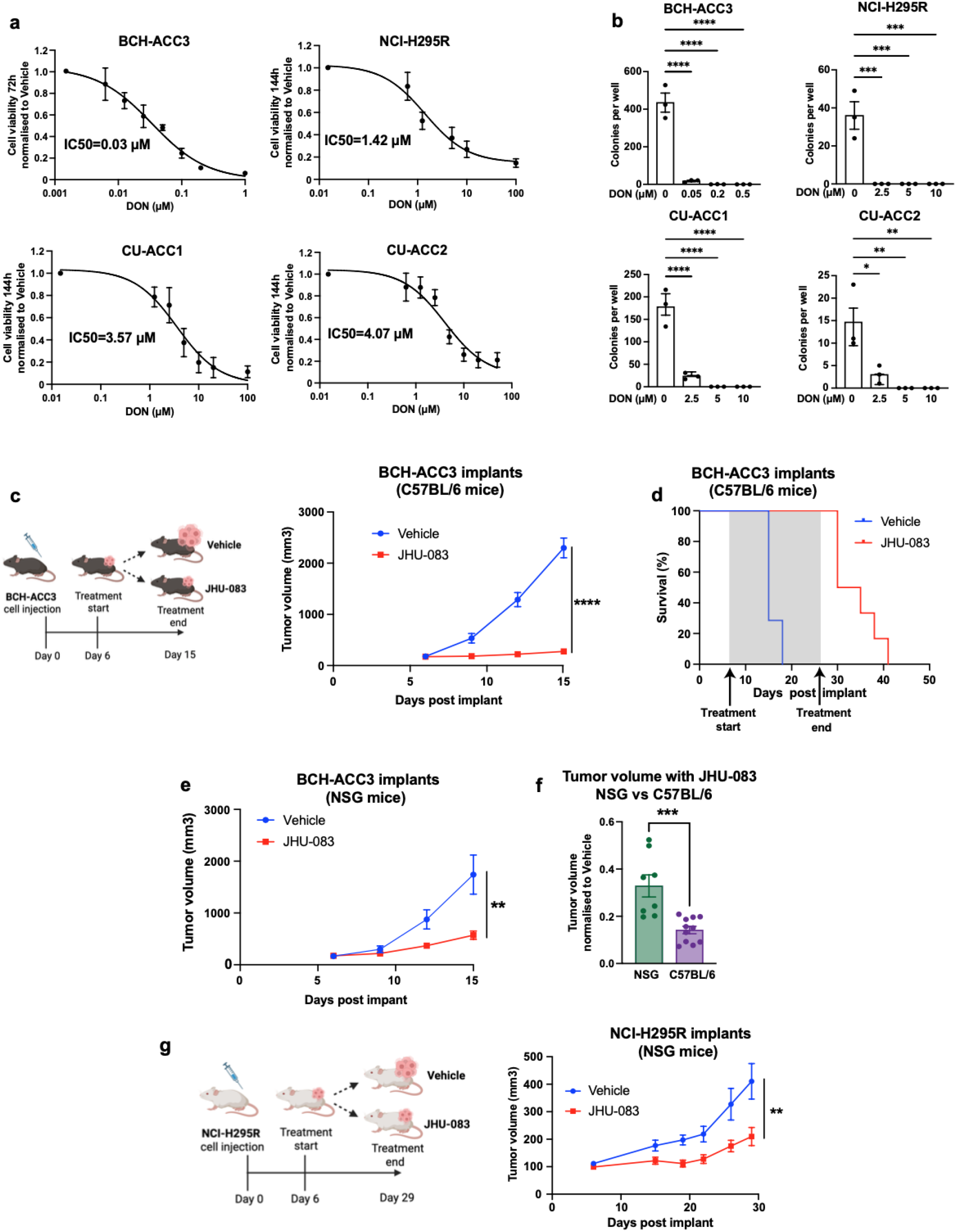
Gln antagonism is highly effective against ACC *in vitro* and *in vivo.* **a** Dose-response curves for four ACC cell lines treated with DON *in vitro*. Data are displayed as mean ± S.E.M of at least three independent experiments (pooled data) with 5 replicates per dose per experiment. In each experiment, cell viability was normalized to the viability of vehicle-treated cells. **b** Clonogenic assays in four ACC cell lines treated with DON *in vitro*. N=3 replicates per dose; *p<0.05, **p<0.01, ***p<0.001, ****p<0.0001. One-way ANOVA with post-hoc Tukey’s tests. Data representative of three independent experiments. **c** Schematic and tumor growth curve from subcutaneous (SQ) BCH-ACC3 implants in male C57BL/6J mice. Mice were treated with JHU-083 1 mg/kg (or vehicle) from day 6-10 post-implantation and 0.33 mg/kg from days 11-15. Tumor inhibition is shown until first sacrifice (day 15). N=9,13; ****p<0.0001 (t-test at the end of the treatment course). Data are representative of three independent experiments. **d** Survival curves post SQ implantation of BCH-ACC3 cells in male C57BL/6J mice. JHU-083 was administered at 1 mg/kg from day 6-10 post-implantation and 0.33 mg/kg from days 11-26. Results were analysed by log-rank (Mantel-Cox) test. Data are representative of two independent experiments. All deaths are due to tumor burden reaching humane endpoints. **e** Tumor growth curve from NCI-H295R cell implants in male immunodeficient NSG mice. 1 mg/kg (or vehicle) from day 6-10 post-implantation and 0.33 mg/kg from days 11-15. N=7,8; **p<0.01 (t-test at the end of the treatment course). **f** Tumor volume on day 15 post BCH-ACC3 implant in male C57BL/6J and NSG mice treated in parallel. Values have been normalized to vehicle-treated mice of the same strain. N=8-13; ***p<0.001 (t-test). Data are representative of three independent experiments. **g** Schematic and tumor growth curve from NCI-H295R cell implants in female immunodeficient NSG mice. JHU-083 was administered six days per week for three weeks (days 6-29 post implant). N=13-14 per group. **p<0.01 (t-test at the end of the treatment course).

To test the effectiveness of Gln antagonism *in vivo*, we established a syngeneic subcutaneous (SQ) implant model using BCH-ACC3 cells and immunocompetent male C57BL/6J mice. BCH-ACC3 cells were injected SQ into the flanks of C57BL/6J mice, where they rapidly formed tumors (**Fig. 3c, Suppl. Fig. 5b,c**). To assess the impact of Gln antagonism in this model, mice harboring established SQ tumors were treated with JHU-083, a DON pro-drug that is selectively activated within the tumor microenvironment^12–14^. Treatment with JHU-083 was well tolerated (**Suppl. Fig. 5d**) and led to >80% inhibition of tumor growth after nine days (**Fig. 3c**), demonstrating potent anti-tumor efficacy, while also markedly prolonging survival (**Fig. 3d**). To assess the contribution of immune-mediated mechanisms on tumor growth (as seen in other cancer models^13,14^), we treated immunodeficient NSG mice harboring SQ BCH-ACC3 tumors with JHU-083, which confirmed the strong anti-tumor effect of Gln antagonism, albeit with somewhat lower efficacy than in immunocompetent mice (**Fig. 3e,f**). These results are consistent with JHU-083 having an effect on ACC tumor growth that is in large part immune cell-independent. Importantly, JHU-083 also significantly inhibited tumor growth in a xenograft model of human ACC (using NCI-H295R implants in NSG mice) (**Fig. 3g**), underscoring its preclinical efficacy.

### Inhibition of Gln-dependent nucleotide biosynthesis drives DON’s anti-tumor effect in ACC

Broad-spectrum Gln antagonism by DON can target multiple Gln-dependent metabolic pathways essential for tumor cell proliferation and survival, including glutaminolysis, non-essential amino-acid biosynthesis, hexosamine biosynthesis, glutathione biosynthesis, and *de novo* purine and pyrimidine biosynthesis^12,21,22^. To identify the metabolic pathway(s) responsible for DON’s cytotoxicity in ACC, we performed rescue experiments with cultured ACC cells to functionally bypass specific Gln-dependent processes. Supplementation of ACC cell lines with a cell-permeable nucleoside (Nuc) cocktail was able to fully rescue cell viability in all four ACC cell lines (**Fig. 4a, Suppl. Fig. 6a**). In contrast, supplementation with (1) non-essential amino-acids (NEAA), (2) N-acetylglucosamine (GlcNAc) to support protein glycosylation pathways, (3) α-ketoglutarate (αKG) to restore TCA cycle function, or (4) antioxidants such as glutathione (GSH) or vitamin E (Trolox), failed to rescue the effect of DON on cell viability (**Fig. 4a,b**). These findings identify the blockade of Gln-fueled *de novo* nucleotide biosynthesis as the principal mechanism underpinning DON’s effect on ACC cell viability.

**Fig. 4:**
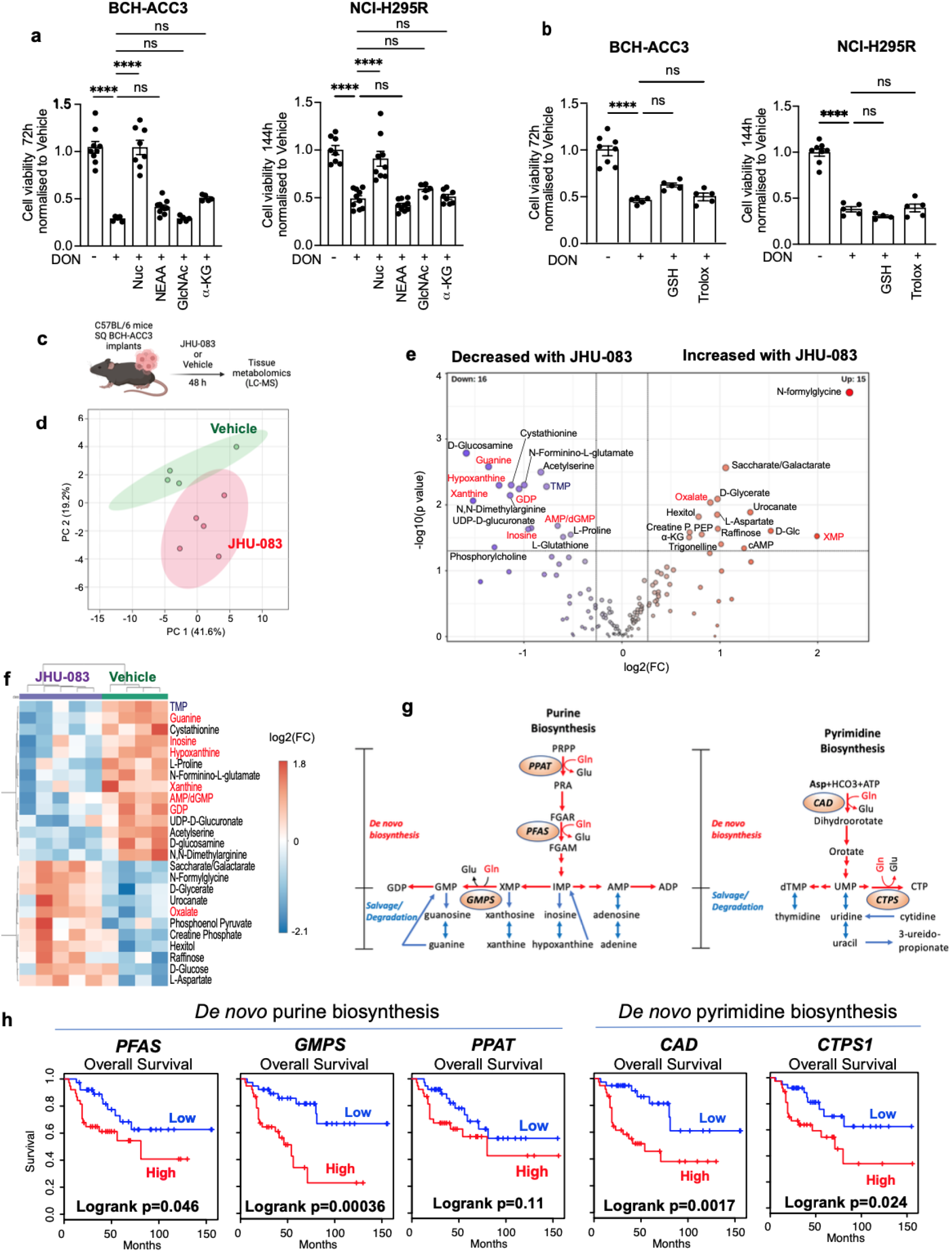
Gln-fueled *de novo* nucleotide biosynthesis as a metabolic vulnerability in ACC. **a** BCH-ACC3 and NCI-H295R cell viability normalized to Vehicle (DON-) after treatment with DON (DON +), alone or supplemented with the indicated metabolites. DON was administered at 50 nM for 72h for BCH-ACC3 and 5 µM for 144h for NCI-H295R cells. Nuc: Embryomax nucleoside cocktail 1x, NEAA: non-essential amino-acid cocktail (0.1 mM), NAG: n-acetyl-glucosamine (10 μM), α-KG: dimethyl-2-oxo-glutarate, cell-permeable form of α-ketoglutarate, 5.5 mM. Data representative of three independent experiments (5-8 technical replicates per experiment). ****p<0.0001, one-way ANOVA followed by Tukey’s test for multiple comparisons between DON monotherapy and each other group. **b** BCH-ACC3 (left) or NCI-H295R (right) viability normalized to Vehicle (DON-) after treatment with DON (DON +), alone or with co-treatment with antioxidants (reduced glutathione (GSH) 2 mM or Trolox 10 μΜ). DON was administered at 50 nM for 72h for BCH-ACC3 and 5 μM for 144h for NCI-H295R. Data representative of three independent experiments (4-8 technical replicates per experiment). ****p<0.0001, one-way ANOVA followed by Tukey’s test for multiple comparisons between DON monotherapy and each other group. **c** Schematic of JHU-083 treatment of BCH-ACC3 implants for metabolomic analysis. **d** Principal Component Analysis (PCA) comparing the polar metabolome of BCH-ACC3 implants in C57BL/6 mice after 48h treatment with JHU-083 (1 mg/kg once daily, n=5) or vehicle (n=4). **e** Polar metabolite changes comparing JHU-083 vs vehicle-treated tumors by Volcano plot with fold change threshold (x) >1.2 and t-tests threshold (y) p<0.05. Dashed lines indicate statistical thresholds. Purine metabolite names are highlighted in red font; pyrimidine metabolite names are highlighted in blue font. Metabolites that could not be unambiguously distinguished are reported using dual annotations. **f** Heat-map of the top 25 differential polar metabolites in JHU-083 tumors vs vehicle. Purine metabolite names are highlighted in red font. Pyrimidine metabolite names are highlighted in blue font. **g** Schematic summarizing the two pathways contributing to purine (left) and pyrimidine (right) nucleotide biosynthesis: *de novo* biosynthesis (Gln-dependent, red arrows) and salvage/degradation pathways (blue arrows). The five enzymes that use Gln as a substrate are indicated in circles. **h** Survival curves comparing overall survival in patients with high (top 50%) versus low (bottom 50%) levels of expression of indicated Gln-metabolizing enzymes, based on The Cancer Genome Atlas (TCGA). Significance and hazards ratio were assessed by Log-rank (Mantel-Cox) tests. ADP, adenosine diphosphate; AMP, adenosine monophosphate; CAD, carbamoyl-phosphate synthetase II/aspartate transcarbamylase/dihydroorotase; CTP, cytidine triphosphate; CTPS, CTP synthase; dTMP, deoxythymidine monophosphate; FGAM, formylglycinamidine ribonucleotide; FGAR, formylglycinamide ribonucleotide; GDP, guanosine diphosphate; Gln, glutamine; Glu, glutamate; GMP, guanosine monophosphate; GMPS, GMP synthase; IMP, inosine monophosphate; PFAS, phosphoribosylformylglycinamidine synthase; PPAT, phosphoribosyl pyrophosphate amidotransferase; PRA, 5-phosphoribosylamine; PRPP, 5-phosphoribosyl 1-pyrophosphate; UMP, uridine monophosphate.

To assess the metabolic impact of Gln antagonism on ACC *in vivo*, we treated mice harboring SQ BCH-ACC3 tumors with JHU-083 and applied LC-MS-based targeted metabolomics 48 hours after treatment (**Fig. 4c**). This analysis revealed a marked depletion of purine metabolites (including guanine, GDP, AMP/dGMP, inosine, xanthine and hypoxanthine) and, to a lesser extent, pyrimidine intermediates (e.g., thymidine monophosphate, TMP) (**Fig. 4d–g, Suppl. Fig. 6b**). Consistent with on-target inhibition of Gln-dependent purine biosynthesis, JHU-083 treatment led to a marked accumulation of xanthosine monophosphate (XMP), a substrate of the Gln-utilizing enzyme GMPS in the purine biosynthetic pathway. JHU-083 treatment also led to decreased glutathione levels, consistent with partial inhibition of Gln-dependent antioxidant synthesis, and a modest trend towards decreased hexosamine biosynthesis. However glutamate, α-ketoglutarate, and TCA cycle intermediates were not depleted by JHU-083, suggesting that mitochondrial metabolism is not broadly suppressed by JHU-083 (**Suppl. Fig. 6b**). Collectively, these findings validate *de novo* purine biosynthesis as the dominant Gln-dependent pathway disrupted by JHU-083 *in vivo* and confirm this metabolic axis as a critical vulnerability in ACC.

### High expression of Gln-metabolizing enzymes in *de novo* nucleotide biosynthesis correlates with poor prognosis

Building on the findings that ACC is highly sensitive to Gln antagonism, we examined the gene expression of all known Gln-utilizing enzymes in human ACC using the TCGA dataset. Remarkably, high expression of four out of five key Gln-dependent enzymes directly involved in *de novo* purine and pyrimidine biosynthesis (*PFAS*, *GMPS*, *CAD*, and *CTPS1*) were significantly associated with worse survival (**Fig. 4g,h**). Interestingly, among 32 other cancer types in TCGA, only five (mesothelioma, low-grade glioma, hepatocellular carcinoma, sarcoma, and cutaneous melanoma) showed a comparable correlation between high expression of two or more Gln-metabolizing enzymes involved in nucleotide biosynthesis and poor prognosis (**Suppl. Fig. 7a**), highlighting the distinctive dependence of ACC on this metabolic axis, particularly in aggressive tumors with poor prognosis. A significant association with overall survival was also observed for *ASNS*, encoding asparagine synthetase, and a trend was noted for *NADSYN1* (coenzyme synthesis), but no differences were observed for enzymes involved in alternative Gln-related pathways such as *GLS* and *GLS2* (glutaminolysis) or *GFPT1* (hexosamine synthesis) (**Suppl. Fig. 7b**).

### Inhibition of the DNA damage response synergizes with Gln antagonism to enhance anti-tumor effects in ACC

Nucleotide depletion is a well-established driver of replication stress, leading to replication fork stalling, accumulation of DNA damage, and ultimately cell death^23^. Given DON disrupts *de novo* nucleotide biosynthesis, we hypothesized that its anti-tumor activity in ACC might involve induction of replication stress. To test this, we assessed γH2AX levels (a marker of DNA damage and replication stress^24^) in SQ BCH-ACC3 tumors treated with JHU-083 or vehicle. This revealed a marked increase in γH2AX staining in treated tumors compared with controls (**Fig. 5a**). Consistent with this, we confirmed that DNA damage was also induced by JHU-083 in human NCI-H295R xenografts implanted in NSG mice (**Fig. 5b**).

**Fig. 5:**
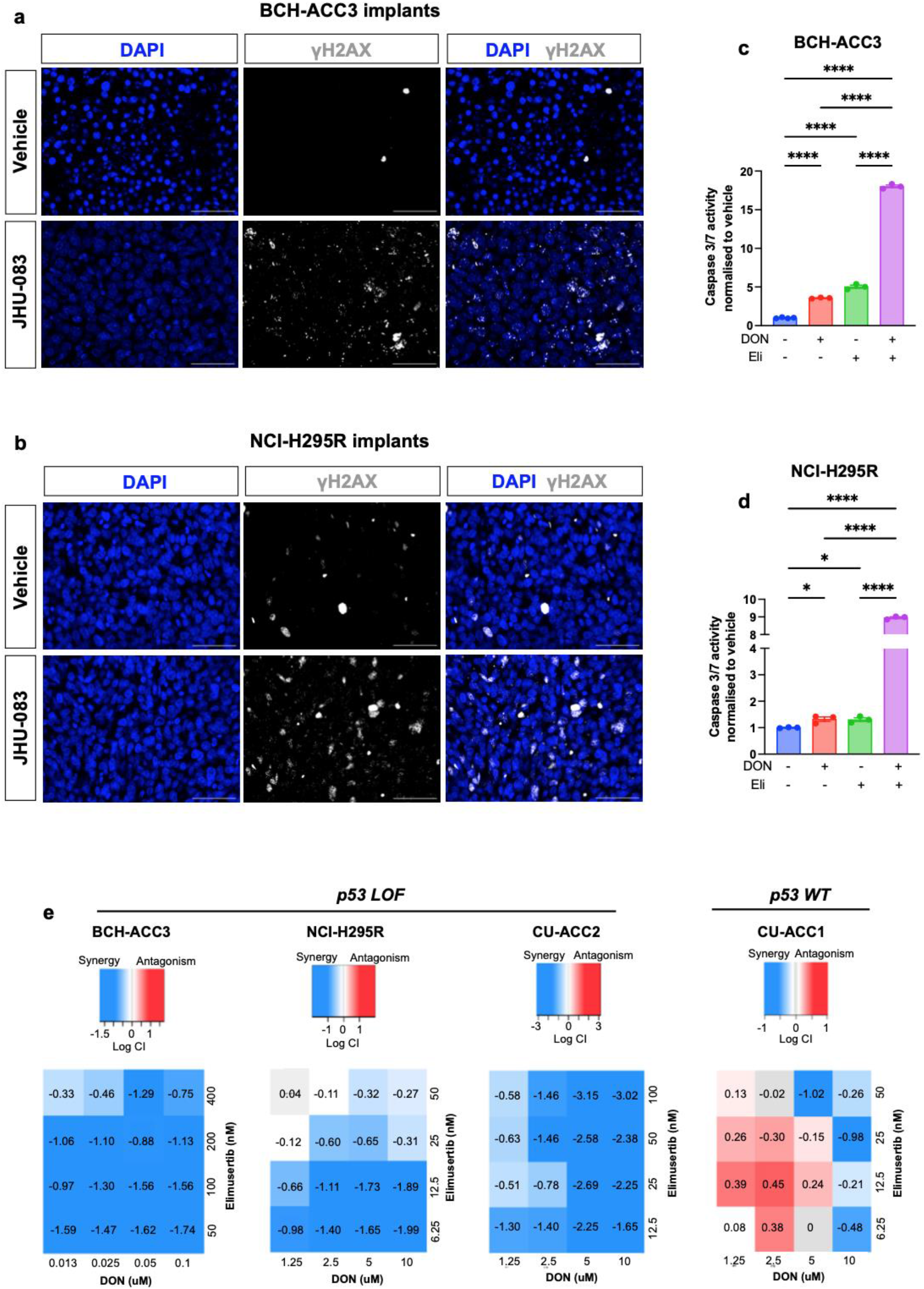
ATR inhibition exhibits synergistic cytotoxicity with Gln antagonism. **a** Representative phosphorylated histone H2A.X (γ-H2AX) staining (grey) of SQ BCH-ACC3 implants in C57BL/6J mice treated with JHU-083 or vehicle for nine days (days 6-15 post-implant). Blue, DAPI. Scale bars: 50 μM. **b** Representative phosphorylated histone H2A.X (γ-H2AX) staining (grey) of SQ NCI-H295R implants in NSG mice treated with JHU-083 or vehicle (days 6-29 post-implant). Blue, DAPI. Scale bars: 50 μM. **c,d** Assessment of DON and Elimusertib (Eli)-induced apoptosis (luminescence-based quantification of activated caspase 3/7) in BCH-ACC3 (**c**) and NCI-H295R (**d**) cells, treated for 48 hours (BCH-ACC3) and 96 hours (NCI-H295R). Data are representative of three independent experiments (3-4 technical replicates per experiment). BCH-ACC3 cells: DON 100 nM, Elimusertib 100 nM; NCI-H295R cells: DON 10 uM, Elimusertib 12.5 nM. For both cell lines, results were analysed by two-way ANOVA, showing very high statistical significance (p<0.0001) for the effect of DON and Elimusertib on apoptosis, as well as for their interaction (consistent with synergy). Post-hoc pairwise comparisons (Tukey’s) are shown on the graphs. *p<0.05, ****p<0.0001. **e** Synergistic effects of DON and Elimusertib on ACC cell viability. ACC cell lines were treated with varying doses of DON and Elimusertib for 72 h (BCH-ACC3) or 144 h (NCI-H295R, CU-ACC1 and CU-ACC2), followed by assessment of cell viability. BCH-ACC3, NCI-H295R and CU-ACC2: data are representative of three independent experiments (4-8 technical replicates per experiment). CU-ACC1: Data represent combination (median viabilities) of three independent experiments. Drug interactions were analysed using SiCoDEA (Single and Combined Drug Effect Analysis). Drug synergy or antagonism was assessed using combination indices calculated according to the Loewe additivity model. Blue color indicates synergy; red color indicates antagonism. Numbers represent log (Combination Index, CI).

Cells counteract the effect of replication stress through activation of the DNA Damage Response (DDR), including the ataxia telangiectasia and Rad3-related (ATR) and ataxia telangiectasia-mutated (ATM) signaling pathways, which promote cell survival by halting cell cycle progression, up-regulating nucleotide synthesis, and activating DNA repair programs^24,25^. Based on the observation that treatment with the DON pro-drug JHU-083 leads to DNA damage in ACC, we hypothesized that inhibiting ATR signaling could block this protective response and drive synthetic lethality in Gln-deprived cancer cells. To test this, we treated ACC cells with the combination of DON and the ATR inhibitor elimusertib, which led to a dramatic increase in apoptotic cell death in both murine (BCH-ACC3) and human (NCI-H295R) ACC cell lines compared to the modest increase seen with either agent alone (**Fig. 5c,d**). We further evaluated the effects of combination treatment on cell viability across a matrix of dose combinations in all four ACC cell lines. Remarkably, we observed robust synergy in all p53-deficient cell lines (BCH-ACC3, NCI-H295R, CU-ACC2), and in CU-ACC1 (p53 wild-type) at higher elimusertib concentrations (**Fig. 5e**). Together, these findings demonstrate that DON-induced replication stress renders ACC cells highly dependent on ATR-mediated DDR for survival. Pharmacological co-inhibition of Gln metabolism and ATR signaling produces synergistic anti-tumor effects, particularly in p53-deficient ACC cells, and highlights a potentially promising combination treatment approach.

### Untargeted serum metabolomics reveals dysregulation of nucleotide and amino-acid metabolism in patients with ACC

To validate the findings showing dysregulation of Gln-dependent metabolic pathways in ACC, we performed untargeted serum metabolomics on pre-operative samples from 54 patients with ACC and 291 age- and body mass index-matched patients with benign adrenocortical adenomas (ACA), prospectively collected across 10 centers in Europe and the U.S. (**Fig. 6a,b**). Differential abundance analysis using Benjamini–Hochberg correction revealed 437 metabolites that differed significantly between ACC and ACA (FDR < 0.05), of which 144 were elevated and 293 were reduced in the serum of patients with ACC (**Fig. 6c**, **Suppl. Table 4**). Polar metabolite pathway analysis revealed eight significantly dysregulated pathways (FDR<0.05), including pyrimidine metabolism and several amino-acid metabolic pathways directly linked to Gln utilization (‘Arginine and Proline Metabolism’, ‘Arginine Biosynthesis’, ‘Alanine, Aspartate and Glutamate metabolism’) (**Fig. 6d and Suppl. Table 5**). Importantly, there was a significant depletion of circulating Gln levels, suggestive of increased uptake of Gln by ACC tumors (**Fig. 6e**). Focusing on nucleotide metabolites, we found an increase in several purine and pyrimidine intermediates in the salvage/degradation pathways, including hypoxanthine, xanthosine, xanthine (**Fig. 6f**), cytidine, uridine, and deoxyuridine (**Fig. 6g**), reflecting enhanced nucleotide turnover consistent with the up-regulated nucleotide biosynthesis programs identified in tumor transcriptional profiles. As expected, there was a marked increase in several steroid conjugates (**Suppl. Fig. 8 and Suppl. Table 4**), in keeping with previous studies using targeted steroid profiling in patients with adrenal tumors^26–28^. Together, these serum metabolomic data complement the tumor-specific analyses and reinforce the presence of key changes in nucleotide and amino-acid metabolism.

**Fig. 6:**
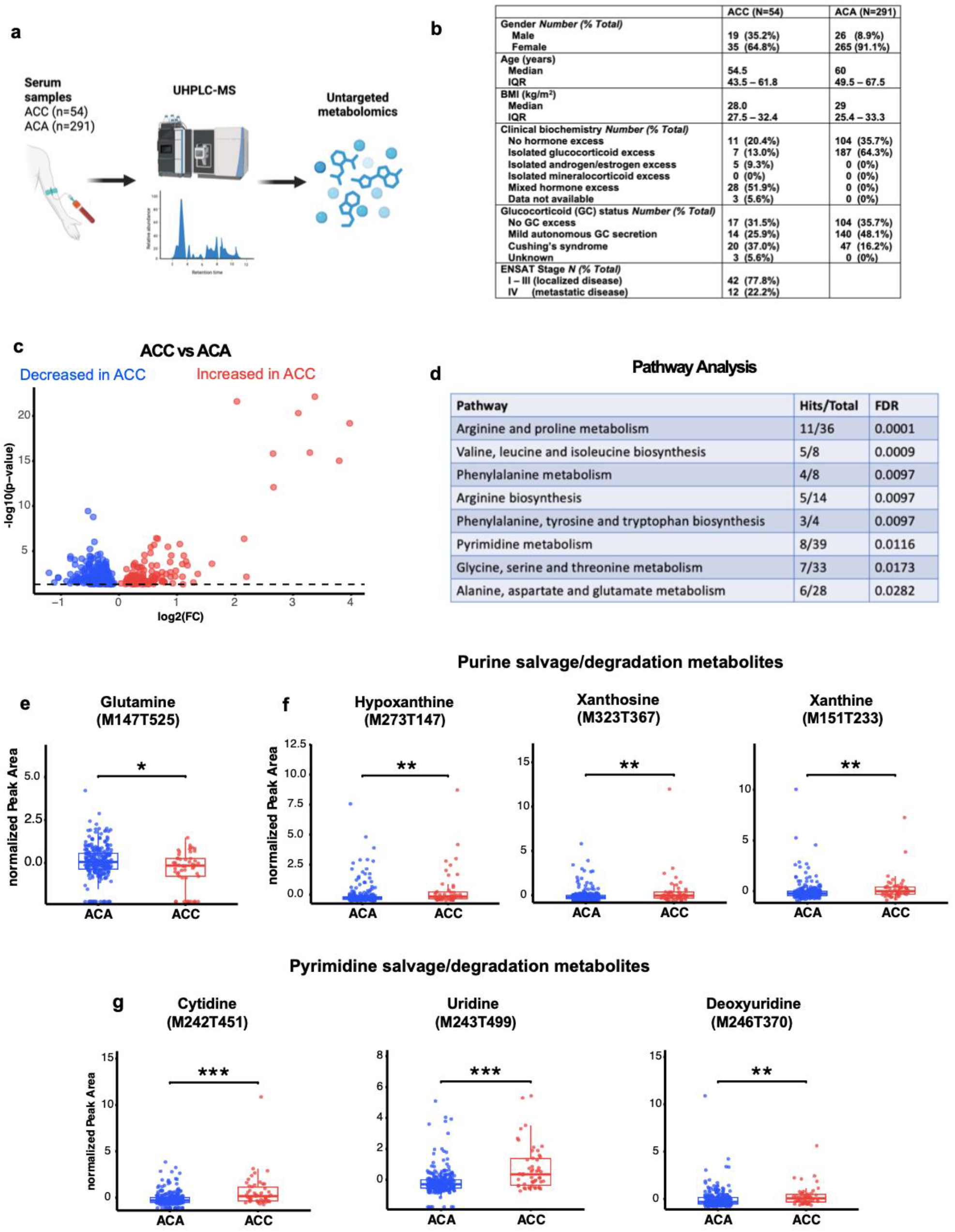
Untargeted global serum metabolomics in patients with ACC vs benign adrenocortical adenomas (ACA). **a** Schematic overview of clinical study. UHPLC-MS: Ultra-high performance liquid chromatography-mass spectrometry. **b** Clinical characteristics of study participants. **c** Volcano plot with significantly altered metabolites in the serum of patients with ACC, n=54 versus ACA, n=291 (red dots: increased in ACC, blue dots: decreased in ACC). **d** Dysregulated pathways (FDR<0.05) derived from analysis of polar metabolites. **e-g** Box and whiskers plots comparing relative serum abundance of Gln (**e**) and selected purine (**f**) and pyrimidine (**g**) metabolites from the salvage/degradation pathways in patients with ACC vs. ACA. The box edges represent the 1st and 3rd quartile and the upper and lower whiskers represent Q3 + 1.5x Interquartile range (IQR) and Q1 - 1.5x IQR, respectively. The median is depicted as the horizontal line within the box. *p<0.05, ** p<0.01, ***p<0.001 (t-test). For all metabolites, alternative putative annotations can be found in Suppl. Table 4 using peak identifiers indicated on the graph titles.

## Discussion

By integrating transcriptional profiling, targeted tissue metabolomics in genetically engineered mouse models, and untargeted serum metabolomics from a large patient cohort, our study provides cross-species validation of metabolic rewiring in ACC. With this approach, we identified Gln metabolism as a central, actionable vulnerability, based on transcriptional up-regulation of common pathways in mouse and human ACC, validation of changes in key metabolites and metabolic pathways in mouse ACC tissues, demonstration of ACC tumor sensitivity to Gln inhibition *in vitro* and *in vivo*, and validation of key changes in circulating metabolites from patients with ACC. Additionally, we showed that addition of ATR inhibition to Gln antagonism results in potent synergy in multiple ACC cell lines *in vitro*, introducing a new treatment paradigm.

Consistent with this work, a previous pan-cancer analysis of the TCGA database used a gene signature consisting of Gln transporters and glutaminase isozymes and showed an association with poor prognosis that was stronger for ACC than for nearly any other cancer type^29^. We selected DON to target Gln metabolism due to its effects on multiple Gln-metabolizing pathways (rather than focused targeting of glutaminolysis), and additionally considering the highly promising new therapeutic opportunities provided by the recent discovery of DON pro-drugs such as JHU-083. While clinical development of DON was previously hindered by the significant systemic toxicity, these new pro-drugs, which are preferentially activated in the tumor microenvironment through moiety cleavage, provide a new avenue to clinical translation for this metabolic treatment^12,19^. Indeed, a drug from this new class (DRP-104) is in Phase I-II trials for glutamine-addicted cancers, including NFE2L2/KEAP1-mutant non-small celllung cancer and fibrolamellar cancer (NCT07249372, NCT06027086). Moving forward, it will be critical to identify cancer types that are most likely to respond to Gln antagonism^30,31^, as well as identify rational combination treatment approaches.

These data reveal that ACC cells are highly sensitive to Gln antagonism, *in vitro,* compared with other cancer cell lines^13,14,22^, and *in vivo*, using both WT and NSG SQ implant models with murine and human cell lines. DON theoretically has the capacity to inhibit all Gln-metabolizing enzymes, including glutaminase as well as transamidases that participate in amino-acid, nucleotide, co-factor and hexosamine biosynthesis^12^. Individual enzyme inhibition kinetics are incompletely described and the relative importance of different Gln-utilizing pathways in the anti-tumor effects of DON is not well understood. The *in vitro* rescue experiments suggest that, in ACC, DON primarily affects cell proliferation and viability through inhibition of *de novo* purine and pyrimidine biosynthesis. Consistent with this, depletion of purine metabolites, in particular, was a prominent change in the metabolome of ACC tumors treated with JHU-083, with a lesser effect on pyrimidine metabolites. This asymmetric nucleotide response is suggestive of preferential functional inhibition of purine *de novo* glutamine-amidotransferases (PPAT, PFAS, GMPS) and/or with greater salvage buffering on the pyrimidine arm. These findings are consistent with recent studies assessing DON as a treatment in KEAP1-mutant lung cancer^31^ and prostate cancer^32^, which also showed that cancer cell-intrinsic sensitivity to Gln antagonism was contingent on inhibition of *de novo* nucleotide biosynthesis. Conversely, nucleoside substitution had no ability to rescue the effect of DON on pancreatic cancer cell viability, while substitution with the amino-acid asparagine did^22^. Previous work in a colon cancer cell line using shRNA-mediated knockdown of individual Gln-metabolizing enzymes suggested that targeting of three Gln-metabolizing enzymes involved in pyrimidine biosynthesis (*Cad* and *Ctps1*) and hexosamine biosynthesis (*Gfpt1*) had the strongest effect on cell viability in comparison to other Gln-metabolizing enzymes^13^. Therefore, an important message of this study is that, although DON may be effective across a number of cancer types, the mechanisms underpinning cancer cell-intrinsic sensitivity to the drug are cancer type-dependent. Interestingly, among the ACC cell lines tested in our study, BCH-ACC3 exhibited the highest sensitivity; this may be explained by the substantially higher proliferation rate of this cell line, which is likely to increase the demand for rapid nucleotide biosynthesis.

Metabolic dysregulation in the nucleotide salvage/degradation pathways was a prominent finding in both the tissue-level analysis in BPCre ACC tumors and in the serum-level metabolomic analysis in the cohort of patients with ACC. Pyrimidine salvage/degradation metabolites were increased at both a tissue level (BPCre tumors) and at a serum level in patients with ACC. Conversely, purine metabolites were relatively depleted in BPCre tumors but increased in the serum of patients. This discrepancy may reflect accelerated release of these metabolites into the bloodstream from viable or necrotic ACC cells, or contributions from other tissues. Alternatively, it is also possible that there may be differences in purine degradation metabolite handling between mouse and human ACC. The observed dysregulation of nucleotide salvage/degradation pathways may plausibly be related to the marked sensitivity of ACC to pharmacological inhibition of the alternative nucleotide biosynthetic pathway (*de novo* biosynthesis), which requires nitrogen contribution by Gln. Dedicated future studies employing metabolic tracing (e.g. by ^13^C and ^15^N-labeled Gln) will be required to probe the dynamics of nucleotide salvage/degradation in ACC.

A key translational question is how to build rational drug combinations that will enhance the efficacy of Gln antagonism. In this work, we demonstrate that Gln antagonism by JHU-083 induces DNA damage in ACC tumors *in vivo*. Interestingly, the same effect was very recently shown in an independent study using the DON pro-drug DRP-104 in other cancer models including prostate, lung, and glioma^33^, supporting the concept that disruption of nucleotide homeostasis by glutamine antagonism can induce DNA damage across tumor types. Building on this observation, we hypothesized that concurrent inhibition of the DNA damage response through the ATR pathway inhibitor elimusertib could yield synergy with DON. Indeed, we confirmed that this new combination can elicit synergistic cytotoxicity across all p53-deficient ACC cell lines tested and, at higher doses, in a p53 WT line. Given that p53 deficiency sensitizes cells to replication stress^34^, this likely underlies the enhanced response observed in p53-mutant cell lines. These findings introduce a new combination treatment paradigm that may be of broader relevance for other p53-deficient cancers, or cancers in which the anti-tumor effect of glutamine antagonism is primarily driven by nucleotide depletion, such as prostate cancer or KEAP1-mutant lung cancer^31–33^.

The clinical study reported here represents, to our knowledge, the largest global metabolomic analysis to date in patients with adrenal tumors and provides a unique metabolomic dataset that will be valuable for future studies exploring new targeted therapeutic approaches or biomarker development. The observed reduction in circulating Gln in patients with ACC is likely to reflect increased tumor uptake and utilization, although we cannot exclude the possibility that this decrease reflects changes in systemic metabolism. It is important to note that the control group for this clinical study was patients with benign adrenocortical adenomas rather than healthy adrenals, in order to allow exploration of diagnostic biomarkers in future work to distinguish benign from malignant adrenal tumors. A limitation of serum metabolomic analyses is that observed differences may, at least in part, reflect changes in systemic metabolism rather than tumor metabolism (due to a lack of specificity). Additionally, mass-to-charge ratios prior to fragmentation (MS1) were used for putative metabolite identification in most cases, rendering distinction between isomeric compounds difficult, with all putative metabolites being reported. Finally, while the two patient cohorts were well-matched for age and BMI, the cohort of patients with ACAs had a stronger female preponderance; however, no statistically significant differences were observed between male and female patients with ACC.

A previous study in human ACC using spatial metabolomics by MALDI mass spectrometry imaging revealed that high intra-tumoral metabolic heterogeneity is associated with poor prognosis^35^. We were not able to assess intra-tumoral heterogeneity in this work. Another limitation is that we did not use metabolic tracing to directly quantify changes in the intracellular metabolic fate of Gln in primary ACC tumors, or in SQ tumors treated with JHU-083. These *in vivo* targeted metabolomic data, however, clearly demonstrate that depletion of the pool of purine metabolites is the prominent metabolic change in JHU-083-treated tumors *in vivo*. Interestingly, there was no apparent depletion of alpha-ketoglutarate or TCA cycle metabolites. This is in keeping with recent metabolic tracing studies in xenograft models of renal cell carcinoma, which showed unimpeded [U-^13^C] Gln labeling of TCA cycle metabolites^36^.

In summary, this study provides a systematic exploration of metabolic rewiring in ACC, leading to the discovery that this rare cancer is sensitive to targeting of Gln’s contribution to *de novo* nucleotide biosynthesis through treatment with Gln antagonists. With the recent development of DON pro-drugs with a broad therapeutic window, these robust pre-clinical findings are of high translational potential for this aggressive and treatment-resistant cancer. We also show that DNA damage induced by Gln antagonism is an important mechanism, serving as the basis for potent synergistic anti-tumor effects through combination with ATR pathway inhibition.

## Methods

### Transcriptomic analysis

To characterize the molecular similarities between BPCre tumors and human ACC, we performed bulk RNA-seq on BPCre tumors (*n* = 7, 5 from female and 2 from male mice, aged 9-12 months) and control mouse adrenals without mutations (*n* = 6, 5 from female ASCre/+ mice and 1 from a C57BL/6J female mouse, aged 5-11 months). We also analyzed publicly available bulk RNA-seq data from human ACC tumors (TCGA-ACC, n=79)^3^ and normal human adrenals (GTEx, n=295)^15^. The GTEx and TCGA datasets were normalized and batch-corrected using the Variance Stabilizing Transformation (vst) function in the DESeq2 package^37^. Differential expression analysis was then performed with DESeq2 as well, restricted to a curated list of 2,754 metabolic genes and their mouse orthologs^16^. Genes with a Benjamini-Hochberg false discovery rate (FDR) < 0.05 were considered significantly differentially expressed.

For pathway interpretation, lists of significant genes were tested for KEGG over-representation using Enrichr^38^ (human and mouse separately), and cross-species conservation was assessed at the pathway level. For prespecified pathways (Purine metabolism, Pyrimidine metabolism, Glutathione metabolism, Amino sugar and nucleotide sugar metabolism), we generated heatmaps using variance-stabilized expression values of the significant genes in each pathway. A curated set of genes involved in de novo and salvage nucleotide biosynthesis or nucleotide metabolism was retrieved from the Reactome database via the Molecular Signatures Database (MSigDB, v2025.1), with cross-referencing to the KEGG database. All analyses and plots were produced in R, utilizing the DESeq2, ComplexHeatmap^39^, and ggplot2 packages^40^.

### Mice

All animal procedures were approved by the Institutional Animal Care and Use Committees of Boston Children’s Hospital. Mouse strains used in this work were: C57BL/6J (Jackson Laboratory, #000664), ASCre/+ (Cyp11b2tm1.1(cre)Brlt^41^), Ctnnb1fl(ex3) (Ctnnb1tm1Mmt^42^), Trp53flox/flox^43^, and NSG (NOD.Cg-Prkdcscid Il2rgtm1Wjl/SzJ) mice (Jackson laboratory, strain #005557). Cross-breeding of transgenic mice led to the generation of the following adrenal-specific transgenic mice: (1) PCre (ASCre/+:: Trp53flox/flox), P53 loss-of-function (LOF) model; (2) BCre (ASCre/+::Ctnnbflox(ex3)/+), β-catenin gain-of-function (GOF) model and (3) BPCre (ASCre/+::Trp53flox/flox::Ctnnbflox(ex3)/+), p53-LOF/βcat-GOF bigenic model. Mouse sex is indicated in each figure legend or Methods. Littermate animals were studied whenever possible.

### Drugs

6-diazo-5-oxo-L-norleucine (DON) was purchased from Sigma (D2141). JHU-083 (HY-122218) and elimusertib (HY-101566) were purchased from Medchem Express.

### Cell Culture

The human ACC cell line NCI-H295R was purchased from the ATCC. Human ACC cell lines CU-ACC1 and CU-ACC2 were provided by Dr. Kiseljak-Vassiliades and Professor Margaret E. Wierman (University of Colorado)^21^. The murine cell line BCH-ACC3 was derived from a 538 mg primary ACC tumor with documented lung metastases in a male BPCre^9^ transgenic mouse. Before using BCH-ACC3 cells in tumor formation experiments, they were first assessed for GFP (as the BPCre mouse model includes the ROSA^mT/mG^ reporter allele) and Cre expression (both known to be immunogenic^44,45^) and found that both genes had been lost during serial SQ passaging in C57BL/6J mice during cell line derivation. Genotyping was performed by PCR using the following primers, as previously described^41,42^: ASCre F 5’-GAGCTGGGGCCCATTTTCAGG-3’, R1 5’-GCTCCAGGTGCATCCGACGG-3’, R2: 5’-AACTTGCACCATGCCGCCCA-3’; GFP F: 5’-CACATGAAGCAGCACGACTT-3’, R: 5’ AGTTCACCTTGATGCCGTTC-3’; β-Cat F: 5′-GGTAGTGGTCCCTGCCCTTGACAC-3′, R: 5′-CTAAGCTTGGCTGGACGTAAACTC-3′. NCI-H295R and BCH-ACC3 cells were cultured in Advanced DMEM/F12 (Thermo Fisher, 12634010) supplemented with 2% Nu serum (Corning, 355100), 1% ITS (Corning, 354350), 1x Glutamax (Thermo Fisher, 35050061) and 1% Penicillin-Streptomycin (ThermoFisher, 15140122). CU-ACC1 and CU-ACC2 cells were cultured in 3:1 v/v mix of F12 Nutrient Mixture (Ham) (Thermo Fisher, 11765-054) and DMEM-high glucose, pyruvate (Thermo Fisher, 11995-065), 10% fetal bovine serum (Thermo Fisher, A5256701), 0.4 μg/mL hydrocortisone (Sigma, H-0888), 5 μg/mL insulin (Sigma, 16634), 8.4 ng/mL cholera toxin (Sigma, C9903), 10 ng/mL epidermal growth factor (Invitrogen, PHG0311), 24 μg/mL adenine (Sigma, A2786), and 1% Penicillin-Streptomycin (ThermoFisher, 15140122), as previously described^20^. Cells were tested routinely for Mycoplasma infection by PCR.

### *In vitro* assessment of cell viability and apoptosis

To assess the *in vitro* efficacy of DON against murine and human ACC cell lines, cells were plated in 96-well plates (Falcon) at different seeding densities depending on the growth kinetics of each cell line: 3,500 cells per well (BCH-ACC3), 6,000 cells per well (CU-ACC1 and CU-ACC2) and 8,000 cells/well (NCI-H295R). The next day, cell culture medium was removed and replaced by 200 µl treatment medium (DON, elimusertib, or combination) at varying concentrations as specified in individual figure legends. Treatment was applied for a total of 72 hours (BCH-ACC3) or 144 hours (NCI-H295R, CU-ACC1 and CU-ACC2) to allow two doubling times. For rescue experiments, cells were co-treated with individual metabolites as specified in figure legends, including 1x non-essential amino-acid cocktail (Thermo Fisher, 11140050), 1x nucleoside cocktail Embryomax (Millipore, ES-008-D), dimethyl-2-oxoglutarate 5.5 mM (Sigma, 349631), GlcNAc 10 μM (Sigma, A8625), 2 mM reduced glutathione (Sigma, G1404), or Trolox 10 μM (Abcam, ab120747). At the end of the treatment course, relative viable cell numbers were measured using the CyQuant Direct Cell Proliferation Assay (Thermo Fisher, C35011) as per the manufacturer’s instructions. Baseline cancer cell line doubling times (in regular growth medium) were calculated by culturing each cell line for 72 hours and using the formula Doubling Time (hours) = (72 * ln(2)) / ln(final cell number / initial cell number). Drug interactions were analysed using SiCoDEA (Single and Combined Drug Effect Analysis). Drug synergy or antagonism was assessed using combination indices calculated according to the Loewe additivity model^46,47^. Cellular apoptosis was assessed using the Caspase-Glo 3/7 Assay kit (Promega, G8090), a luminescence-based assay measuring Caspase 3 and 7 activity in cell lysates following the manufacturer’s instructions, as described^48^. To account for differences in cell numbers present at the time of apoptosis measurements, caspase assay results were normalized to relative viable cell numbers, as assessed by the CyQuant Direct Cell Proliferation Assay run in wells treated in parallel.

### *In vitro* clonogenic assays

For 2D clonogenic assays, cells were plated onto 6-well plates at a seeding density of 1,000 cells per well (BCH-ACC3, NCI-H295R, CU-ACC2) and 5,000 cells per well (CU-ACC1, as seeding 1,000 cells per well did not lead to colony development in this latter cell line). The next day, DON was added directly to wells at the desired concentrations, as specified in figure legends (medium was not changed to avoid washing out cells). Medium containing the drug was replenished every 3-4 days. The treatment course was ended once confluent colony formation was observed in vehicle-treated wells or at pre-specified endpoints (whichever was earlier). Pre-specified endpoints were 10 days for BCH-ACC3 cells and 28 days for NCI-H295R, CU-ACC1 and CU-ACC2. At the end of the course, plates were washed with Phosphate-Buffered Saline (PBS) and fixed in 4% paraformaldehyde in PBS for 30 min. After washing with PBS again, wells were stained with crystal violet (0.1% crystal violet in 5% ethanol in water) for 30 min, then washed with water and left to dry inverted overnight. The next day, cells were imaged at 2x magnification using an Invitrogen EVOS FL 2 Auto microscope. Colonies comprising at least 50 cells were counted in Fiji using the ‘Analyse Particles’ tool.

### JHU-083 treatment *in vivo*

For experiments involving BCH-ACC3 cells, 3x10^6^ cells in 100 μl PBS/Matrigel (Corning) (1:1) were injected subcutaneously in the right flank of 2-month-old male C57BL/6J or NSG mice (Jackson). Tumors were measured by electronic caliper on day 6 and volume was calculated using the formula volume=length x (width)^2^/2. Mice were randomized to treatment groups and treatment was commenced on the same day. JHU-083 was diluted in 100% ethanol and stored at -20°C. Working solution of JHU-083 in PBS with 2.5% ethanol was prepared every day and administered by oral gavage at a dose of 1 mg/kg/day (days 6-10) and 0.33 mg/kg thereafter, as previously described^13^. Tumor growth and mouse weight was monitored by electronic caliper every 72 hours. For tumor analysis by histology, mice were sacrificed and tumors were collected on day 15. JHU-083 or vehicle was administered 60-90 min before tumor collection.

For experiments involving NCI-H295R cells, 10x10^6^ cells in 200 μl PBS/Matrigel (1:1) were injected subcutaneously in the right flank of 2-month-old female NSG mice (Jackson). Tumors were measured by electronic caliper on day 6 and volume was calculated using the formula volume=length x (width)^2^/2. Mice were randomized to treatment groups and treatment was commenced on the same day. JHU-083 was diluted in 100% ethanol and stored at -20°C. Working solution of JHU-083 in PBS with 2.5% ethanol was prepared every day and administered by oral gavage at a dose of 1 mg/kg/day 6 days per week (days 6-29). Treatment dose was reduced to 0.5 mg/kg/day in animals with weight loss>7.5% from baseline and suspended in animals with weight loss >12.5% from baseline until weight improved (no mice lost >20% of their body weight or reached endpoints for euthanasia under this protocol). Tumor growth and mouse weight was monitored by electronic caliper every 72 hours. For both implant models, mice were sacrificed when tumor volume exceeded 2,000 mm^3^, tumors became ulcerated or necrotic, or when mice lost >20% weight from baseline.

### Targeted tissue metabolomic analysis

For metabolomic analysis comparing primary ACC tumors in BPCre mice with adrenals from other transgenic mice and from wild-type mice, mice were euthanized and tumors were immediately dissected and a snap-frozen on dry ice. Because of the small weight of mouse adrenals, two pooled adrenals were used per sample, either deriving from the same mouse or from two separate mice of the same sex, age and genotype. For ACC tumors up to 10 mg of tissue was used per sample. For metabolomic analysis comparing SQ BCH-ACC3 implants treated with JHU-083 or vehicle, mice were euthanized after 48 h of treatment and tumors were immediately dissected and snap-frozen on dry ice. Up to 10 mg of tissue was used per sample. Samples were extracted with methanol: water: chloroform (1:1:1). The water-methanol phase containing polar metabolites was dried down and reconstituted with water/acetonitrile (1:1) with the volume normalized to tissue wet weight. The polar extract was separated on an iHILIC column (5 μm, 150 mm x 2.1 mm I.D., Nest Group)^49^. The column was coupled to an Agilent 6546 LC/Q-TOF with an ESI source operated in negative and/or positive mode. The identity of each metabolite was confirmed by matching retention time and MS/MS fragmentation data to standard compounds and/or a database. Metabolite quantification was performed using MassHunter Profinder 10.0 software (Agilent). Data were analysed using Metaboanalyst 6.0. Features with >40% missing data for all groups were removed, and other missing variables were replaced by zero. Peak intensities were normalized to internal standards (D8-phenylalanine for polar metabolites) and to sum, followed by log transformation and scaling before statistical analysis.

### Paraffin Section Immunofluorescence

After fixation in 4% paraformaldehyde, harvested ACC implants were embedded in paraffin blocks. Paraffin sections were cut at 5 µm thickness. Sections were rinsed in xylene, an ethanol gradient and then phosphate-buffered saline (PBS). Antigen retrieval was performed in 10 mM Sodium Citrate pH 6.0. Sections were blocked in 5% Normal Goat Serum (Sigma, G9023) in PBS for 1 hour at room temperature. Primary Phospho-Histone H2A.X (Ser139) Antibody (Cell Signaling, #2577) was diluted 1:200 in 5% normal goat serum in PBS and incubated on sections at 4°C overnight. Slides were washed three times for 5 min in 0.1% Tween-20 in PBS. Secondary antibody (Alexa Fluor 594-conjugated goat anti-rabbit IgG antibody) was diluted 1:200 in PBS and incubated on sections at room temperature for 1-2 hours. For nuclear staining, DAPI (4’,6-diamidino-2-phenylindole) was added to the secondary antibody mixture at a final concentration of 1:1000. After three 5-min washes with 0.1% Tween-20 in PBS, slides were mounted with ProLong Gold Antifade Mountant (Thermo Fisher, P36930). Images were acquired using a Nikon upright Eclipse 90i microscope and adjusted for brightness and contrast in ImageJ.

### TCGA transcriptome analysis

GEPIA (online analysis tool) was used to access and process publicly available transcriptomic data from ACC and 32 other human cancers, collected in the context of The Cancer Genome Atlas (TCGA)^50^. Survival comparisons were performed by logrank (Mantel-cox) tests.

### Clinical study design

All patients were prospectively recruited through the cross-sectional Evaluation of Urine Steroid Metabolomics in the Differential Diagnosis of Adrenocortical Tumors (EURINE-ACT) study, whose primary aim was to validate urine steroid profiling as a new diagnostic test for ACC^26^. The EURINE-ACT study was registered and approved by the European Network for the Study of Adrenal Tumors (ENSAT), in addition to local institute review board approvals. All patient recruitment centers had ethical approval for pseudonymized phenotype recording in the online ENSAT database and all participants of the EURINE-ACT study provided written informed consent prior to study participation.

Inclusion criteria included adult participants (age ≥18 years) with a newly identified adrenal mass of more than 1 cm diameter. Exclusion criteria were biochemical evidence of pheochromocytoma, pregnancy, lactation, and current or recent (<6 months) intake of drugs known to alter steroid synthesis or metabolism. Patients with an adrenal mass discovered during imaging for cancer staging or monitoring were also not eligible. Additional details on patient recruitment for EURINE-ACT are available here^26^. Centers that contributed serum samples for this study came from the following countries: Croatia (Zagreb), France (Bordeaux), Greece (Athens), Germany (Berlin, Munich, Wurzburg, Dresden), Ireland (Galway), Norway (Bergen), Poland (Warsaw), United Kingdom (Birmingham), and the United States (Mayo Clinic). 54 patients with ACC were able to provide serum samples for metabolomic analysis. As a control group, we selected 291 patients with benign adrenocortical adenomas (ACAs) from the same study (EURINE-ACT), who were age- and BMI-matched for the cohort of ACCs. This included patients with non-functioning adrenocortical adenomas, as defined by clinical biochemistry, and patients with glucocorticoid excess, as defined by clinical biochemistry (1 mg dexamethasone suppression test showing cortisol suppression to <1.8 mg/dl).

### Untargeted serum metabolome analysis

For each serum sample, stored at -80°C, 50μL aliquots were used for untargeted metabolome analysis. Two assays were applied to increase the coverage of metabolic features detected; polar (water-soluble) metabolites were analysed by hydrophilic interaction chromatography (HILIC) UHPLC-MS, and non-polar (lipid) metabolites by a C_18_ reversed-phase lipidomics UHPLC-MS. All samples were analyzed separately in positive and negative ion modes. Raw data files were deconvoluted using the XCMS software. Scatterplot smoothing signal correction was used for signal correction. The mean and relative standard deviation of metabolic feature concentrations were calculated against pooled quality control samples. Following data normalization by probabilistic quotient normalization, metabolic features were annotated using the in-house BEAMS software and the mass spectral library mzCloud. All non-lipid metabolites were grouped into classes based on KEGG metabolic pathway involvement by applying pathway enrichment analysis in MetaboAnalyst. For lipid-based results, lipids were classified according to LipidMaps ontologies manually. Data were visualized using R v4.2.2 (r-project.org) as implemented in R studio v. 2024.04.0 for MacOS. A more detailed description of the method can be found in the **Supplementary Data**.

### Quantification and Statistical Analysis

All statistical analyses were performed and graphs prepared using Prism 10 (GraphPad), Metaboanalyst version 6.0^51^ or R v4.2.2 (r-project.org) as implemented in R studio v. 2024.04.0 for MacOS (for metabolomic data). The statistical test used to assess significance for each experiment is indicated in figure legends. To correct for multiple comparisons in the untargeted metabolomic analysis of human serum samples and targeted metabolomic analysis of murine tissue samples, FDR-corrected p values using the Benjamini-Hochberg method were used. To detect metabolites that showed step-wise progression from control adrenals to BPCre-Pre adrenals to ACC tumors, we used Pearson’s correlation coefficients through the Pattern Finder function on Metaboanalyst. For survival analyses, significance and hazards ratio were assessed by Log-rank (Mantel-Cox) tests. In bar graphs, data are presented as Mean ± Standard Error of the Mean (SEM), unless otherwise stated. p<0.05 was considered statistically significant, unless otherwise stated. Data collection and analysis were not performed blind to the experimental conditions. Biorender was used for preparation of schematics.

### Sex as a biological variable

RNA-seq analysis and targeted transcriptomics on BPCre tumors was performed on mice from both sexes. The murine ACC cell line was derived from a tumor that developed in a male mouse. Human cell lines were derived from female patients. Male C57BL/6J and NSG mice were used as hosts in the BCH-ACC3 implant model. Female NSG mice were used as hosts for the NCI-H295R xenograft model. Both male and female patients were included in the clinical study. The ACC cohort had a female preponderance (65%), in line with disease demographics. We were not able to fully match the ACA cohort for sex (91% female preponderance).

## Acknowledgement

We thank members of the Breault and Haigis Labs and Dr. Esin Isik for helpful discussions. This work was supported by R01 DK123694 to D.T.B., a Michail Papamichail, MD, PhD, Postdoctoral Fellowship at Harvard Medical School to V.C., an Academy of Medical Sciences U.K. Clinical Lecturer Starter Grant SGL020\1018 to V.C., the Wellcome Trust (Investigator Award 209492/Z/17/Z to W.A. and Clinical Research Training Fellowship WT101671 to V.C.), the European Commission under the 7th Framework Program (FP7/2007-2013, grant agreement 259735, ENSAT-CANCER, to W.A., M.F., and F.B.), the UK Medical Research Council (MRC) through the construction of the Phenome Center Birmingham (MR/M009157/1) and core programme grant MC_UP_1605/15 to W.A., and a São Paulo Research Foundation (FAPESP) scholarship 2024/14165-6 to J.L.K. Biorender was used to generate schematics for this work (https://BioRender.com/y9blqb5, https://BioRender.com/w6r6cqp, https://biorender.com/dl0p0rq, https://biorender.com/k7qp0fh, https://biorender.com/t50p462 ).

## Author Contributions

V.C., K.S.B., W.A., W.B.D., M.C.H. and D.T.B. conceptualized the study and developed the methodology; V.C., K.S.B., C.Y., C.R., M.B., P.V., A.E.T.X., J.L.K., L.N., A.J. and C.W. performed the experiments; V.C., K.S.B., L.F.N., M.E.K., L.K., S.R. and T.G.P. performed formal data analysis; V.C. and M.E.K. visualized data for figures; V.C., A.P., M.M., I.D.P., M.Q., M.C.D., G.A.U., F.B., A.T., M.F., D.K., U.A., D.A.V., M.E.W., K.K.V. and I.B. provided resources (serum samples, cell lines); V.C., A.P., A.E.T. and I.B. provided project administration for the clinical study; all authors interpreted the data; V.C., K.S.B. and D.T.B. wrote the original paper with editorial inputs from all authors; W.B.D., W.A., M.C.H. and D.T.B. provided the supervision; W.A., M.C.H. and D.T.B. acquired the funds.

## Abbreviations

ACC: Adrenocortical carcinoma
ACA: Adrenocortical adenoma
ATR: Ataxia telangiectasia and Rad3-related
DDR: DNA damage response
DON: 6-diazo-5-oxo-L-norleucine
FDR: False discovery rate
GOF: Gain-of-function
LC-MS: Liquid chromatography–mass spectrometry
LOF: Loss-of-function
TCA: Tricarboxylic acid cycle TCGA The cancer genome atlas

## References

1. Fassnacht M, Terzolo M, Allolio B, et al. Combination chemotherapy in advanced adrenocortical carcinoma. N Engl J Med 2012;366(23):2189–97. DOI: 10.1056/NEJMoa1200966.

2. Fassnacht M, Dekkers OM, Else T, et al. European Society of Endocrinology Clinical Practice Guidelines on the management of adrenocortical carcinoma in adults, in collaboration with the European Network for the Study of Adrenal Tumors. Eur J Endocrinol 2018;179(4):G1–G46. DOI: 10.1530/EJE-18-0608.

3. Zheng S, Cherniack AD, Dewal N, et al. Comprehensive Pan-Genomic Characterization of Adrenocortical Carcinoma. Cancer Cell 2016;29(5):723–736. DOI: 10.1016/j.ccell.2016.04.002.

4. Assie G, Letouze E, Fassnacht M, et al. Integrated genomic characterization of adrenocortical carcinoma. Nat Genet 2014;46(6):607–12. DOI: 10.1038/ng.2953.

5. Mohan DR, Lerario AM, Finco I, Hammer GD. New strategies for applying targeted therapies to adrenocortical carcinoma. Curr Opin Endocr Metab Res 2019;8:72–79. DOI: 10.1016/j.coemr.2019.07.006.

6. Pozdeyev N, Fishbein L, Gay LM, et al. Targeted genomic analysis of 364 adrenocortical carcinomas. Endocr Relat Cancer 2021;28(10):671–681. DOI: 10.1530/ERC-21-0040.

7. Borges KS, Pignatti E, Leng S, et al. Wnt/beta-catenin activation cooperates with loss of p53 to cause adrenocortical carcinoma in mice. Oncogene 2020;39(30):5282–5291. DOI: 10.1038/s41388-020-1358-5.

8. Mariniello K, Pittaway JFH, Altieri B, et al. Dlk1 is a novel adrenocortical stem/progenitor cell marker that predicts malignancy in adrenocortical carcinoma. Cancer Commun (Lond) 2025;45(6):663–668. DOI: 10.1002/cac2.70012.

9. Mohan DR, Borges KS, Finco I, et al. beta-Catenin-Driven Differentiation Is a Tissue-Specific Epigenetic Vulnerability in Adrenal Cancer. Cancer Res 2023;83(13):2123–2141. DOI: 10.1158/0008-5472.CAN-22-2712.

10. Hanahan D, Weinberg RA. Hallmarks of cancer: the next generation. Cell 2011;144(5):646–74. DOI: 10.1016/j.cell.2011.02.013.

11. Hanahan D. Hallmarks of Cancer: New Dimensions. Cancer Discov 2022;12(1):31–46. DOI: 10.1158/2159-8290.CD-21-1059.

12. Lemberg KM, Vornov JJ, Rais R, Slusher BS. We’re Not "DON" Yet: Optimal Dosing and Prodrug Delivery of 6-Diazo-5-oxo-L-norleucine. Mol Cancer Ther 2018;17(9):1824–1832. DOI: 10.1158/1535-7163.MCT-17-1148.

13. Leone RD, Zhao L, Englert JM, et al. Glutamine blockade induces divergent metabolic programs to overcome tumor immune evasion. Science 2019;366(6468):1013–1021. DOI: 10.1126/science.aav2588.

14. Oh MH, Sun IH, Zhao L, et al. Targeting glutamine metabolism enhances tumor-specific immunity by modulating suppressive myeloid cells. J Clin Invest 2020;130(7):3865–3884. DOI: 10.1172/JCI131859.

15. Consortium GT. The Genotype-Tissue Expression (GTEx) project. Nat Genet 2013;45(6):580–5. DOI: 10.1038/ng.2653.

16. Possemato R, Marks KM, Shaul YD, et al. Functional genomics reveal that the serine synthesis pathway is essential in breast cancer. Nature 2011;476(7360):346–50. DOI: 10.1038/nature10350.

17. Panwalkar P, Tamrazi B, Dang D, et al. Targeting integrated epigenetic and metabolic pathways in lethal childhood PFA ependymomas. Sci Transl Med 2021;13(614):eabc0497. DOI: 10.1126/scitranslmed.abc0497.

18. Dang D, Deogharkar A, McKolay J, et al. Isocitrate dehydrogenase 1 primes group-3 medulloblastomas for cuproptosis. Cancer Cell 2025;43(6):1159–1174 e8. DOI: 10.1016/j.ccell.2025.04.013.

19. DeBerardinis RJ. Tumor Microenvironment, Metabolism, and Immunotherapy. N Engl J Med 2020;382(9):869-871. DOI: 10.1056/NEJMcibr1914890.

20. Kiseljak-Vassiliades K, Zhang Y, Bagby SM, et al. Development of new preclinical models to advance adrenocortical carcinoma research. Endocr Relat Cancer 2018;25(4):437–451. DOI: 10.1530/ERC-17-0447.

21. Encarnacion-Rosado J, Sohn ASW, Biancur DE, et al. Targeting pancreatic cancer metabolic dependencies through glutamine antagonism. Nat Cancer 2024;5(1):85–99. DOI: 10.1038/s43018-023-00647-3.

22. Recouvreux MV, Grenier SF, Zhang Y, et al. Glutamine mimicry suppresses tumor progression through asparagine metabolism in pancreatic ductal adenocarcinoma. Nat Cancer 2024;5(1):100–113. DOI: 10.1038/s43018-023-00649-1.

23. Mullen NJ, Singh PK. Nucleotide metabolism: a pan-cancer metabolic dependency. Nat Rev Cancer 2023;23(5):275–294. DOI: 10.1038/s41568-023-00557-7.

24. Saxena S, Zou L. Hallmarks of DNA replication stress. Mol Cell 2022;82(12):2298–2314. DOI: 10.1016/j.molcel.2022.05.004.

25. Qiu Z, Oleinick NL, Zhang J. ATR/CHK1 inhibitors and cancer therapy. Radiother Oncol 2018;126(3):450–464. DOI: 10.1016/j.radonc.2017.09.043.

26. Bancos I, Taylor AE, Chortis V, et al. Urine steroid metabolomics for the differential diagnosis of adrenal incidentalomas in the EURINE-ACT study: a prospective test validation study. Lancet Diabetes Endocrinol 2020;8(9):773–781. DOI: 10.1016/S2213-8587(20)30218-7.

27. Arlt W, Biehl M, Taylor AE, et al. Urine steroid metabolomics as a biomarker tool for detecting malignancy in adrenal tumors. J Clin Endocrinol Metab 2011;96(12):3775–84. DOI: 10.1210/jc.2011-1565.

28. Yu K, Athimulam S, Saini J, et al. Serum Steroid Profiling in the Diagnosis of Adrenocortical Carcinoma: A Prospective Cohort Study. J Clin Endocrinol Metab 2025;110(4):1177–1186. DOI: 10.1210/clinem/dgae604.

29. Zou J, Du K, Li S, et al. Glutamine Metabolism Regulators Associated with Cancer Development and the Tumor Microenvironment: A Pan-Cancer Multi-Omics Analysis. Genes (Basel) 2021;12(9). DOI: 10.3390/genes12091305.

30. Yokoyama Y, Estok TM, Wild R. Sirpiglenastat (DRP-104) Induces Antitumor Efficacy through Direct, Broad Antagonism of Glutamine Metabolism and Stimulation of the Innate and Adaptive Immune Systems. Mol Cancer Ther 2022;21(10):1561–1572. DOI: 10.1158/1535-7163.MCT-22-0282.

31. Pillai R, LeBoeuf SE, Hao Y, et al. Glutamine antagonist DRP-104 suppresses tumor growth and enhances response to checkpoint blockade in KEAP1 mutant lung cancer. Sci Adv 2024;10(13):eadm9859. DOI: 10.1126/sciadv.adm9859.

32. Moon D, Hauck JS, Jiang X, et al. Targeting glutamine dependence with DRP-104 inhibits proliferation and tumor growth of castration-resistant prostate cancer. Prostate 2024;84(4):349–357. DOI: 10.1002/pros.24654.

33. Yu J, Jin C, Su C, et al. Resilience and vulnerabilities of tumor cells under purine shortage stress. Clin Cancer Res 2025. DOI: 10.1158/1078-0432.CCR-25-1667.

34. Dembitz V, Tomic B, Kodvanj I, Simon JA, Bedalov A, Visnjic D. The ribonucleoside AICAr induces differentiation of myeloid leukemia by activating the ATR/Chk1 via pyrimidine depletion. J Biol Chem 2019;294(42):15257–15270. DOI: 10.1074/jbc.RA119.009396.

35. Wang Q, Sun N, Meixner R, et al. Metabolic heterogeneity in adrenocortical carcinoma impacts patient outcomes. JCI Insight 2023;8(16). DOI: 10.1172/jci.insight.167007.

36. Kaushik AK, Tarangelo A, Boroughs LK, et al. In vivo characterization of glutamine metabolism identifies therapeutic targets in clear cell renal cell carcinoma. Sci Adv 2022;8(50):eabp8293. DOI: 10.1126/sciadv.abp8293.

37. Love MI, Huber W, Anders S. Moderated estimation of fold change and dispersion for RNA-seq data with DESeq2. Genome Biol 2014;15(12):550. DOI: 10.1186/s13059-014-0550-8.

38. Kuleshov MV, Jones MR, Rouillard AD, et al. Enrichr: a comprehensive gene set enrichment analysis web server 2016 update. Nucleic Acids Res 2016;44(W1):W90–7. DOI: 10.1093/nar/gkw377.

39. Gu Z, Eils R, Schlesner M. Complex heatmaps reveal patterns and correlations in multidimensional genomic data. Bioinformatics 2016;32(18):2847–9. DOI: 10.1093/bioinformatics/btw313.

40. Wickham H. ggplot2 – Elegant Graphics for Data Analysis. Second ed. New York: Springer-Verlag, 2016.

41. Freedman BD, Kempna PB, Carlone DL, et al. Adrenocortical zonation results from lineage conversion of differentiated zona glomerulosa cells. Dev Cell 2013;26(6):666–673. DOI: 10.1016/j.devcel.2013.07.016.

42. Harada N, Tamai Y, Ishikawa T, et al. Intestinal polyposis in mice with a dominant stable mutation of the beta-catenin gene. EMBO J 1999;18(21):5931–42. DOI: 10.1093/emboj/18.21.5931.

43. Marino S, Vooijs M, van Der Gulden H, Jonkers J, Berns A. Induction of medulloblastomas in p53-null mutant mice by somatic inactivation of Rb in the external granular layer cells of the cerebellum. Genes Dev 2000;14(8):994–1004. (https://www.ncbi.nlm.nih.gov/pubmed/10783170).

44. Schultheiss C, Binder M. Overcoming unintended immunogenicity in immunocompetent mouse models of metastasis: the case of GFP. Signal Transduct Target Ther 2022;7(1):68. DOI: 10.1038/s41392-022-00929-9.

45. Dubrot J, Lane-Reticker SK, Kessler EA, et al. In vivo screens using a selective CRISPR antigen removal lentiviral vector system reveal immune dependencies in renal cell carcinoma. Immunity 2021;54(3):571–585 e6. DOI: 10.1016/j.immuni.2021.01.001.

46. Spinozzi G, Tini V, Ferrari A, et al. SiCoDEA: A Simple, Fast and Complete App for Analyzing the Effect of Individual Drugs and Their Combinations. Biomolecules 2022;12(7). DOI: 10.3390/biom12070904.

47. Foucquier J, Guedj M. Analysis of drug combinations: current methodological landscape. Pharmacol Res Perspect 2015;3(3):e00149. DOI: 10.1002/prp2.149.

48. Chortis V, Taylor AE, Doig CL, et al. Nicotinamide Nucleotide Transhydrogenase as a Novel Treatment Target in Adrenocortical Carcinoma. Endocrinology 2018;159(8):2836–2849. DOI: 10.1210/en.2018-00014.

49. Yao CH, Park JS, Kurmi K, et al. Uncoupled glycerol-3-phosphate shuttle in kidney cancer reveals that cytosolic GPD is essential to support lipid synthesis. Mol Cell 2023;83(8):1340–1349 e7. DOI: 10.1016/j.molcel.2023.03.023.

50. Tang Z, Li C, Kang B, Gao G, Li C, Zhang Z. GEPIA: a web server for cancer and normal gene expression profiling and interactive analyses. Nucleic Acids Res 2017;45(W1):W98–W102. DOI: 10.1093/nar/gkx247.

51. Pang Z, Lu Y, Zhou G, et al. MetaboAnalyst 6.0: towards a unified platform for metabolomics data processing, analysis and interpretation. Nucleic Acids Res 2024;52(W1):W398–W406. DOI: 10.1093/nar/gkae253.

