## Supplementary material for "Glutamine antagonism suppresses tumor growth in adrenocortical carcinoma through inhibition of de novo nucleotide biosynthesis": Suppl. Table 1A

**Suppl. Table 1A mouse Up-regulated pathways BPCre ACC tumors versus Control adrenals**

| Term | Overlap | P-value | Adjusted P-v | Odds Ratio | Combined Score | Genes |
|---|---|---|---|---|---|---|
| Pyrimidine metabolism | 22/58 | 1.13E-19 | 2.76E-17 | 21.91528 | 956.1387097 | DTYMK;DUT;RRM1;RRM2;DPYS;NME4;TYMS;NT5C2;DCK;NT5C3;UCKL1;DHODH;DCTD;NT5E;UCK2;UCK1;NME7;TK2;UPP2;NT5M;TK1;UMPS |
| Drug metabolism | 28/114 | 7.15E-19 | 8.68E-17 | 11.776309 | 492.0340815 | MAOA;DPYS;GSTP1;GMPS;GSTT2;GSTT1;MPO;ADH5;TK2;UPP2;TK1;GUSB;GSTM4;GSTM3;GSTM2;DUT;RRM1;GSTM1;RRM2;NME4;UCKL1;UCK2;UCK1;NME7;IMPDH1;UMPS;ITPA;GSTM5 |
| Purine metabolism | 30/136 | 1.06E-18 | 8.68E-17 | 10.264611 | 424.8089396 | GMPS;AK4;NUDT5;NT5C2;AK7;PFAS;NT5C3;PAPSS1;DGUOK;NT5E;PDE11A;ATIC;PDE4A;NT5M;NUDT16;PDE8A;ADSL;RRM1;RRM2;NME4;DCK;APRT;PKM;NME7;IMPDH1;PDE5A;GART;PDE9A;ITPA;ADA |
| Glycosaminoglycan biosynthesis | 20/53 | 5.96E-18 | 3.65E-16 | 21.657459 | 858.9564626 | B4GALT2;CSGALNACT1;B4GALT3;B4GALT1;CHPF;CHPF2;XYLT2;HS6ST2;HS3ST1;NDST2;EXT2;FUT8;EXTL1;B3GNT7;CHST11;CHST12;B3GNT2;CHST14;B4GALT7;ST3GAL2 |
| Inositol phosphate metabolism | 20/73 | 7.50E-15 | 3.68E-13 | 13.470934 | 438.1261581 | ISYNA1;TPI1;IPMK;INPPL1;SYNJ2;ITPKB;INPP4A;ITPKC;PLCB3;PLCB4;IMPA2;ITPKA;INPP5E;PIP5K1A;PIP4K2B;PIK3C3;PIP4K2C;PLCG1;PI4KB;PLCD3 |
| Phosphatidylinositol signaling system | 22/98 | 3.13E-14 | 1.28E-12 | 10.359519 | 322.144491 | IPMK;INPPL1;PIK3R2;SYNJ2;DGKZ;INPP4A;ITPKB;ITPKC;PLCB3;PLCB4;PPIP5K2;IMPA2;INPP5E;ITPKA;PIP5K1A;PIP4K2B;PIK3C3;PIP4K2C;PLCG1;PI4KB;DGKK;PLCD3 |
| Cysteine and methionine metabolism | 16/50 | 2.56E-13 | 7.93E-12 | 16.692548 | 483.9913522 | MPST;AHCY;MDH2;GSS;GOT2;MAT2B;MTAP;LDHA;TST;CBS;ADI1;CTH;KYAT3;BCAT1;GCLM;ENOPH1 |
| Glycerophospholipid metabolism | 21/97 | 2.59E-13 | 7.93E-12 | 9.8703875 | 286.0657725 | PLA2G2F;LYPLA2;PLA2G2D;CHKB;PLA2G2E;PNPLA7;LPCAT1;MBOAT1;MBOAT2;PLA2G5;AGPAT1;DGKZ;PLD3;PLD2;AGPAT4;PHOSPHO1;ETNK2;LPCAT3;PNPLA6;DGKK;LPIN3 |
| Glycine, serine and threonine metabolism | 14/40 | 1.98E-12 | 5.39E-11 | 19.038391 | 513.0577594 | GAMT;ALAS2;MAOA;SHMT2;BPGM;GCAT;GCSH;GRHPR;CBS;PSAT1;CTH;SARDH;PHGDH;PSPH |
| N-Glycan biosynthesis | 14/50 | 6.17E-11 | 1.51E-09 | 13.742866 | 323.0739159 | B4GALT2;B4GALT3;ALG8;ST6GAL1;B4GALT1;SRD5A3;MOGS;DPM2;FUT8;DOLPP1;MGAT4B;MAN1C1;MGAT3;MAN1B1 |
| Glutathione metabolism | 15/64 | 2.05E-10 | 4.57E-09 | 10.830478 | 241.5981749 | GSTM4;GGT5;GSTM3;GSTM2;RRM1;GSTM1;RRM2;GSTP1;GSS;IDH2;GPX7;GSTT2;GSTT1;GCLM;GSTM5 |
| One carbon pool by folate | 19/9 | 7.48E-10 | 1.53E-08 | 31.560108 | 663.1799588 | DHFR;ATIC;MTHFD1L;SHMT2;MTHFD2;MTHFR;TYMS;ALDH1L2;GART |
| Fructose and mannose metabolism | 11/35 | 1.80E-09 | 3.40E-08 | 16.118886 | 324.5255502 | PFKFB1;PFKL;PFKFB3;TPI1;AKR1B10;GMPPA;PMM2;SORD;ALDOA;PFKM;PFKP |
| Amino sugar and nucleotide sugar metabolism | 12/49 | 7.86E-09 | 1.38E-07 | 11.419042 | 213.0926277 | GPI1;UGDH;CYB5R1;GNPDA1;UXS1;AMDHD2;GMPPA;UAP1L1;GFPT1;PMM2;NPL;GALK1 |
| Porphyrin and chlorophyll metabolism | 10/41 | 1.47E-07 | 2.41E-06 | 11.320072 | 178.0672101 | ALAD;ALAS2;MMAB;CPOX;HMOX1;BLVRB;GUSB;HMOX2;COX10;BLVRA |
| Selenocompound metabolism | 17/7 | 2.05E-07 | 3.14E-06 | 24.458453 | 376.6213092 | TXNRD3;TXNRD2;TXNRD1;CTH;KYAT3;SEPHS1;PAPSS1 |
| Metabolism of xenobiotics by cytochrome P450 | 12/66 | 2.71E-07 | 3.91E-06 | 7.8173019 | 118.196535 | GSTM4;AKR7A5;GSTM3;GSTM2;GSTM1;GSTP1;CYP1A1;GSTT2;GSTT1;ADH5;CBR3;GSTM5 |
| Mineral absorption | 10/44 | 3.02E-07 | 4.12E-06 | 10.319647 | 154.9155708 | SLC11A2;SLC40A1;HMOX1;TRPM7;ATP1A1;ATP1B2;SLC26A9;ATP1B1;SLC8A1;HMOX2 |
| Glycolysis / Gluconeogenesis | 12/67 | 3.22E-07 | 4.15E-06 | 7.6747731 | 114.7232885 | GPI1;LDHA;PFKL;TPI1;PKM;BPGM;ALDOA;GAPDH;PFKM;ADH5;PFKP;PCK2 |
| Glycosphingolipid biosynthesis | 10/45 | 3.79E-07 | 4.64E-06 | 10.024283 | 148.2160889 | B4GALT2;B4GALT3;B4GALT1;B3GALT4;ST8SIA5;NAGA;B3GNT2;ST6GALNAC4;ST3GAL2;ST6GALNAC6 |
| Calcium signaling pathway | 20/189 | 4.12E-07 | 4.81E-06 | 4.1993309 | 61.73944289 | SPHK2;CACNA1A;ATP2A1;GRIN2C;SLC8A1;CACNA1H;CACNA1G;GRIN2D;TPCN1;ITPKB;ITPKC;PLCB3;PLCB4;P2RX4;P2RX3;ORAI3;ITPKA;VDAC3;PLCG1;PLCD3 |
| Mucin type O-glycan biosynthesis | 28/8 | 7.09E-07 | 7.90E-06 | 13.994234 | 198.146779 | GALNT12;GALNT7;GALNT2;GALNT18;GALNT1;C1GALT1;ST3GAL2;B4GALT5 |
| Oxidative phosphorylation | 16/134 | 1.19E-06 | 1.23E-05 | 4.7888947 | 65.32392505 | NDUFA9;NDUFA8;NDUFA7;SDHC;TCIRG1;COX6A2;ATP4A;PPA2;NDUFS7;NDUFAB1;NDUFS3;NDUFS2;CYC1;ATP6V0C;ATP6V1C1;COX10 |
| Arachidonic acid metabolism | 13/89 | 1.21E-06 | 1.23E-05 | 6.0213636 | 82.06353861 | PLA2G2F;GGT5;PLA2G2D;PTGIS;PLA2G2E;PTGES2;PTGES3;ALOX15;GPX7;ALOX12;PLA2G5;LTA4H;CBR3 |
| Ferroptosis | 9/40 | 1.31E-06 | 1.28E-05 | 10.169675 | 137.7804696 | GSS;LPCAT3;ALOX15;SLC11A2;SLC40A1;VDAC3;HMOX1;SLC7A11;GCLM |
| Other types of O-glycan biosynthesis | 22/7 | 1.59E-06 | 1.50E-05 | 16.301439 | 217.6203819 | B4GALT2;B4GALT3;ST6GAL1;POFUT2;B4GALT1;POMT1;GXYLT2 |
| Citrate cycle (TCA cycle) | 8/32 | 2.17E-06 | 1.83E-05 | 11.659459 | 152.0320994 | MDH2;IDH2;IDH3B;SUCLG2;OGDHL;SDHC;FH1;PCK2 |
| Galactose metabolism | 8/32 | 2.17E-06 | 1.83E-05 | 11.659459 | 152.0320994 | B4GALT2;PFKL;B4GALT1;AKR1B10;LCT;PFKM;PFKP;GALK1 |
| Pentose phosphate pathway | 8/32 | 2.17E-06 | 1.83E-05 | 11.659459 | 152.0320994 | GPI1;RPIA;PFKL;H6PD;ALDOA;TKT;PFKM;PFKP |
| Mannose type O-glycan biosynthesis | 23/7 | 2.23E-06 | 1.83E-05 | 15.281812 | 198.8370631 | B3GALNT2;B4GALT2;B4GALT3;B4GALT1;POMT1;LARGE1;POMGNT1 |
| Glycosylphosphatidylinositol (GPI)-anchor biosynthesis | 25/7 | 4.17E-06 | 3.30E-05 | 13.582434 | 168.2452005 | PIGC;DPM2;PIGU;PIGT;PIGO;PIGM;PIGX |
| Sphingolipid metabolism | 9/48 | 6.56E-06 | 5.02E-05 | 8.0802555 | 96.43646388 | ACER2;UGCG;SMPD2;SGPL1;SPTLC3;SPHK2;SGMS2;GBA2;CERS1 |
| Fatty acid degradation | 9/50 | 9.32E-06 | 6.92E-05 | 7.6853042 | 89.02252739 | GCDH;ACADVL;ACAA2;ACADL;ECI1;ECI2;CPT1C;ACADS;ADH5 |
| Alzheimer disease | 17/175 | 9.61E-06 | 6.93E-05 | 3.799126 | 43.88844529 | NDUFA9;NDUFA8;NDUFA7;PSEN2;SDHC;ATP2A1;GRIN2C;COX6A2;GRIN2D;PLCB3;PLCB4;NDUFS7;NDUFAB1;NDUFS3;NDUFS2;CYC1;GAPDH |
| Central carbon metabolism in cancer | 10/64 | 1.11E-05 | 7.80E-05 | 6.4908579 | 74.02962567 | SLC7A5;LDHA;PFKL;PKM;SLC2A1;SLC2A2;PIK3R2;SLC16A3;PFKM;PFKP |
| Peroxisome | 11/84 | 2.31E-05 | 1.57E-04 | 5.2859837 | 56.43540129 | PECR;ACOT8;AMACR;ECI2;ECH1;IDH2;MLYCD;FAR2;DECR2;DHRS4;SOD1 |
| Proximal tubule bicarbonate reclamation | 22/6 | 2.46E-05 | 1.63E-04 | 13.07518 | 138.7604148 | CAR2;SLC25A10;ATP1A1;ATP1B2;ATP1B1;PCK2 |
| Sulfur relay system | 8/4 | 3.97E-05 | 2.50E-04 | 34.763864 | 352.2661904 | MPST;NFS1;MOCS3;TST |
| Pancreatic secretion | 12/105 | 3.98E-05 | 2.50E-04 | 4.5299456 | 45.90095901 | PLA2G2F;CAR2;PLCB3;PLA2G2D;PLCB4;PLA2G2E;KCNMA1;ATP2A1;ATP1B2;PLA2G5;ATP1A1;ATP1B1 |
| Ether lipid metabolism | 8/47 | 4.48E-05 | 2.74E-04 | 7.169508 | 71.78798751 | PLA2G2F;PLA2G2D;PLA2G2E;LPCAT1;ENPP2;PLA2G5;PLD3;PLD2 |
| Cardiac muscle contraction | 10/78 | 6.50E-05 | 3.89E-04 | 5.1507818 | 49.65551154 | CACNG6;CACNB2;CACNB3;CACNG8;ATP1B2;ATP1A1;CYC1;ATP1B1;COX6A2;SLC8A1 |
| Chemical carcinogenesis | 11/94 | 6.66E-05 | 3.89E-04 | 4.6467173 | 44.68549301 | GSTM4;GSTM3;GSTM2;GSTM1;GSTP1;CYP1A1;KYAT3;GSTT2;GSTT1;ADH5;GSTM5 |
| Arginine and proline metabolism | 8/50 | 7.11E-05 | 4.05E-04 | 6.6563707 | 63.58219455 | GAMT;CKM;MAOA;SMOX;GOT2;PYCR1;PYCR2;CNDP2 |
| Collecting duct acid secretion | 27/6 | 8.66E-05 | 4.82E-04 | 9.9594768 | 93.16278146 | CAR2;ATP4A;SLC4A1;TCIRG1;ATP6V0C;ATP6V1C1 |
| Hepatocellular carcinoma | 15/171 | 1.03E-04 | 5.60E-04 | 3.3831064 | 31.0615783 | GSTM4;GSTM3;NQO1;GSTM2;GSTM1;TXNRD3;TXNRD2;GSTP1;TXNRD1;GSTT2;GSTT1;PIK3R2;HMOX1;PLCG1;GSTM5 |
| Glucagon signaling pathway | 11/102 | 1.40E-04 | 7.48E-04 | 4.2364628 | 37.58063491 | PFKFB1;GYS1;LDHA;PLCB3;PFKL;PKM;PLCB4;SLC2A1;SLC2A2;CPT1C;PCK2 |
| Insulin secretion | 10/86 | 1.50E-04 | 7.81E-04 | 4.6066908 | 40.56912637 | PLCB3;PLCB4;ABCC8;KCNMA1;SLC2A1;SLC2A2;ATP1B2;ATP1A1;ATP1B1;TRPM4 |
| Valine, leucine and isoleucine degradation | 8/56 | 1.62E-04 | 8.28E-04 | 5.8225225 | 50.81291464 | BCKDHA;MCCC2;ACAA2;HIBADH;HMGCS2;BCAT1;ACADS;ACSF3 |
| Ubiquinone and other terpenoid-quinone biosynthesis | 11/4 | 1.75E-04 | 8.76E-04 | 19.861998 | 171.8071267 | VKORC1;NQO1;COQ6;COQ5 |
| Glyoxylate and dicarboxylate metabolism | 6/31 | 1.96E-04 | 9.59E-04 | 8.364237 | 71.41843166 | GCSH;GRHPR;SHMT2;MDH2;AFMID;HYI |
| Sphingolipid signaling pathway | 12/124 | 2.00E-04 | 9.62E-04 | 3.7577781 | 32.00202955 | ACER2;SMPD2;SGPL1;ABCC1;PLCB3;PLCB4;SPTLC3;SPHK2;PIK3R2;SGMS2;CERS1;PLD2 |
| Glycerolipid metabolism | 8/61 | 2.97E-04 | 0.0014006 | 5.2718681 | 42.81230471 | AKR1B10;MBOAT1;MBOAT2;DGKK;AGPAT1;DGKZ;LPIN3;AGPAT4 |
| Thyroid hormone signaling pathway | 11/115 | 4.04E-04 | 0.0018683 | 3.704414 | 28.94509445 | PLCB3;PLCB4;SLC2A1;PIK3R2;SLC16A10;ATP1B2;PLCG1;ATP1A1;ATP1B1;PLCD3;PFKP |
| Alanine, aspartate and glutamate metabolism | 6/37 | 5.36E-04 | 0.0024309 | 6.7432675 | 50.7886951 | ADSL;GFPT1;GOT2;ASNS;NIT2;ASS1 |
| Pyruvate metabolism | 6/38 | 6.21E-04 | 0.0027676 | 6.5322038 | 48.23182221 | GRHPR;LDHA;PKM;MDH2;FH1;PCK2 |
| Thiamine metabolism | 15/4 | 6.62E-04 | 0.0028971 | 12.636852 | 92.50131479 | THTPA;NFS1;AK4;AK7 |
| Fluid shear stress and atherosclerosis | 12/143 | 7.38E-04 | 0.0031731 | 3.2095981 | 23.1452536 | GSTM4;GSTM3;NQO1;GSTM2;GSTM1;GSTP1;GSTT2;HMOX1;GSTT1;PIK3R2;ASS1;GSTM5 |
| Parkinson disease | 12/144 | 7.85E-04 | 0.003317 | 3.185118 | 22.77203359 | NDUFA9;NDUFA8;NDUFA7;NDUFS7;NDUFAB1;VDAC3;NDUFS3;NDUFS2;SDHC;CYC1;COX6A2;SLC6A3 |
| Fat digestion and absorption | 6/40 | 8.24E-04 | 0.0033636 | 6.1473228 | 43.65617134 | PLA2G2F;PLA2G2D;PLA2G2E;GOT2;PLA2G5;AGPAT1 |
| Nicotine addiction | 6/40 | 8.24E-04 | 0.0033636 | 6.1473228 | 43.65617134 | GABRB3;CHRNA4;GABRA5;CACNA1A;GRIN2C;GRIN2D |
| Primary bile acid biosynthesis | 16/4 | 8.63E-04 | 0.0034676 | 11.583184 | 81.71547647 | CYP39A1;HSD3B7;ACOT8;AMACR |
| Adrenergic signaling in cardiomyocytes | 12/148 | 9.99E-04 | 0.0039482 | 3.0907975 | 21.35317119 | CACNG6;CACNB2;CACNB3;PLCB3;CACNG8;PLCB4;ATP1B2;SCN5A;ATP1A1;ATP1B1;SLC8A1;SCN1B |
| Huntington disease | 14/192 | 0.0011179 | 0.004299 | 2.7591126 | 18.75166656 | NDUFA9;NDUFA8;NDUFA7;SDHC;COX6A2;SOD1;PLCB3;PLCB4;NDUFS7;NDUFAB1;VDAC3;NDUFS3;NDUFS2;CYC1 |
| Retrograde endocannabinoid signaling | 12/150 | 0.001123 | 0.004299 | 3.0456877 | 20.68552425 | GABRB3;NDUFA9;NDUFA8;PLCB3;NDUFA7;PLCB4;NDUFS7;GABRA5;NDUFAB1;NDUFS3;CACNA1A;NDUFS2 |
| Carbohydrate digestion and absorption | 6/43 | 0.0012184 | 0.0045925 | 5.6480179 | 37.89929433 | SLC2A2;PIK3R2;ATP1A1;ATP1B2;ATP1B1;LCT |
| Sulfur metabolism | 9/3 | 0.0016419 | 0.0060038 | 17.349107 | 111.2411751 | MPST;TST;PAPSS1 |
| Vitamin B6 metabolism | 9/3 | 0.0016419 | 0.0060038 | 17.349107 | 111.2411751 | PDXK;PSAT1;PDXP |
| ABC transporters | 6/48 | 0.0021787 | 0.0076256 | 4.9743524 | 30.48782865 | ABCC3;ABCC1;ABCC2;ABCB6;ABCC8;ABCG1 |
| Tryptophan metabolism | 6/48 | 0.0021787 | 0.0076256 | 4.9743524 | 30.48782865 | GCDH;MAOA;AFMID;CYP1A1;KYAT3;DHTKD1 |
| Type II diabetes mellitus | 6/48 | 0.0021787 | 0.0076256 | 4.9743524 | 30.48782865 | PKM;ABCC8;CACNA1A;SLC2A2;PIK3R2;CACNA1G |
| Thermogenesis | 15/231 | 0.0023876 | 0.0082389 | 2.4357512 | 14.70576188 | NDUFA9;NDUFA8;NDUFA7;KDM1A;COX18;SDHC;CPT1C;COX6A2;NDUFS7;NDUFAB1;NDUFS3;NDUFS2;NDUFAF1;CYC1;COX10 |
| Pentose and glucuronate interconversions | 5/34 | 0.002459 | 0.0083675 | 5.9967866 | 36.02865416 | UGDH;AKR1B10;SORD;GUSB;XYLB |
| HIF-1 signaling pathway | 9/104 | 0.0026753 | 0.0089786 | 3.3075812 | 19.59315205 | LDHA;PFKL;PFKFB3;SLC2A1;HMOX1;PIK3R2;PLCG1;ALDOA;GAPDH |
| AMPK signaling pathway | 10/126 | 0.0030044 | 0.0099469 | 3.0119411 | 17.49242098 | PFKFB1;GYS1;PFKL;PFKFB3;PIK3R2;MLYCD;CPT1C;PFKM;PFKP;PCK2 |
| Taurine and hypotaurine metabolism | 11/3 | 0.0030923 | 0.0101014 | 13.010491 | 75.18572059 | GGT5;ADO;CSAD |
| Nicotinate and nicotinamide metabolism | 5/36 | 0.0031815 | 0.0102561 | 5.609319 | 32.25585683 | NT5E;NNT;NT5M;NT5C2;NT5C3 |
| Phospholipase D signaling pathway | 11/149 | 0.0033412 | 0.010631 | 2.7868226 | 15.88887547 | PLCB3;PLCB4;SPHK2;PIP5K1A;PIK3R2;PLCG1;DGKK;AGPAT1;DGKZ;PLD2;AGPAT4 |
| Non-alcoholic fatty liver disease (NAFLD) | 11/151 | 0.0036996 | 0.0116204 | 2.7467262 | 15.38040018 | NDUFA9;NDUFA8;NDUFA7;NDUFS7;NDUFAB1;NDUFS3;NDUFS2;SDHC;PIK3R2;CYC1;COX6A2 |
| Aldosterone-regulated sodium reabsorption | 5/38 | 0.0040454 | 0.0125389 | 5.2688172 | 29.03206768 | SCNN1A;PIK3R2;ATP1A1;ATP1B2;ATP1B1 |
| Bile secretion | 7/72 | 0.0040943 | 0.0125389 | 3.7521859 | 20.63008904 | ABCC3;CAR2;ABCC2;SLC2A1;ATP1B2;ATP1A1;ATP1B1 |
| Endocrine and other factor-regulated calcium reabsorption | 6/55 | 0.0043634 | 0.0131981 | 4.2621918 | 23.16284916 | PLCB3;PLCB4;ATP1B2;ATP1A1;ATP1B1;SLC8A1 |
| Gastric acid secretion | 7/74 | 0.0047677 | 0.014245 | 3.6398046 | 19.45798858 | CAR2;ATP4A;PLCB3;PLCB4;ATP1B2;ATP1A1;ATP1B1 |
| alpha-Linolenic acid metabolism | 25/4 | 0.004911 | 0.0144964 | 6.6158957 | 35.17189753 | PLA2G2F;PLA2G2D;PLA2G2E;PLA2G5 |
| Tyrosine metabolism | 5/40 | 0.0050651 | 0.0147731 | 4.9672299 | 26.2537342 | HGD;MAOA;GOT2;COMT;ADH5 |
| Folate biosynthesis | 26/4 | 0.0056768 | 0.0163624 | 6.3148479 | 32.65643704 | DHFR;AKR1B10;PAH;FPGS |
| Synaptic vesicle cycle | 7/77 | 0.0059299 | 0.0168933 | 3.4832734 | 17.86135384 | SLC6A7;SLC6A9;CACNA1A;TCIRG1;ATP6V0C;SLC6A3;ATP6V1C1 |
| Salivary secretion | 7/78 | 0.0063607 | 0.0179122 | 3.4340359 | 17.36805674 | PLCB3;PLCB4;KCNMA1;AQP5;ATP1B2;ATP1A1;ATP1B1 |
| Aldosterone synthesis and secretion | 8/102 | 0.0081464 | 0.0226805 | 2.9661491 | 14.26769076 | PLCB3;PLCB4;PRKD2;ATP1B2;ATP1A1;ATP1B1;CACNA1H;CACNA1G |
| Fatty acid elongation | 29/4 | 0.0084399 | 0.0230329 | 5.5562075 | 26.52971755 | ACAA2;TECR;ACOT1;HSD17B12 |
| Lysosome | 9/124 | 0.0084611 | 0.0230329 | 2.7295244 | 13.02605663 | MANBA;NPC1;NPC2;SLC11A2;NAGA;TCIRG1;NAGPA;GUSB;ATP6V0C |
| cGMP-PKG signaling pathway | 11/172 | 0.0096285 | 0.025923 | 2.3858583 | 11.07759837 | PLCB3;PLCB4;KCNJ8;KCNMA1;VDAC3;ATP2A1;PDE5A;ATP1B2;ATP1A1;ATP1B1;SLC8A1 |
| Propanoate metabolism | 4/31 | 0.0106986 | 0.0284909 | 5.1441065 | 23.34209969 | BCKDHA;LDHA;SUCLG2;MLYCD |
| Nitrogen metabolism | 17/3 | 0.0112416 | 0.0296149 | 7.4322704 | 33.35703803 | CAR2;CAR1;CAR9 |
| beta-Alanine metabolism | 4/32 | 0.0119608 | 0.0311744 | 4.9601329 | 21.95415838 | SMOX;DPYS;MLYCD;CNDP2 |
| Insulin resistance | 8/110 | 0.0125479 | 0.0323604 | 2.7323794 | 11.96290772 | GYS1;SLC27A1;GFPT1;SLC2A1;SLC2A2;PIK3R2;SLC27A3;PCK2 |
| Linoleic acid metabolism | 5/50 | 0.0129757 | 0.0330307 | 3.8614098 | 16.77658961 | PLA2G2F;PLA2G2D;PLA2G2E;ALOX15;PLA2G5 |
| Other glycan degradation | 18/3 | 0.0132123 | 0.0330307 | 6.9364286 | 30.01121472 | MANBA;MAN2B2;GBA2 |
| Pantothenate and CoA biosynthesis | 18/3 | 0.0132123 | 0.0330307 | 6.9364286 | 30.01121472 | PANK2;DPYS;BCAT1 |
| Arginine biosynthesis | 19/3 | 0.0153672 | 0.0380299 | 6.502567 | 27.15161061 | ACY1;GOT2;ASS1 |
| Thyroid hormone synthesis | 6/73 | 0.0168128 | 0.0411913 | 3.1142314 | 12.72355595 | PLCB3;PLCB4;GPX7;ATP1B2;ATP1A1;ATP1B1 |
| Phenylalanine, tyrosine and tryptophan biosynthesis | 8/2 | 0.0197906 | 0.0475363 | 11.545455 | 45.28759323 | PAH;GOT2 |
| Riboflavin metabolism | 8/2 | 0.0197906 | 0.0475363 | 11.545455 | 45.28759323 | FLAD1;BLVRB |
| Choline metabolism in cancer | 7/99 | 0.0217615 | 0.0517628 | 2.6473061 | 10.13286355 | CHKB;PIP5K1A;PIK3R2;PLCG1;DGKK;DGKZ;PLD2 |
| Phenylalanine metabolism | 23/3 | 0.0258559 | 0.0603304 | 5.2009821 | 19.01072008 | MAOA;PAH;GOT2 |
| Terpenoid backbone biosynthesis | 23/3 | 0.0258559 | 0.0603304 | 5.2009821 | 19.01072008 | PDSS2;PDSS1;HMGCS2 |
| Vitamin digestion and absorption | 24/3 | 0.028947 | 0.0669059 | 4.9530612 | 17.54516606 | ABCC1;AWAT2;SLC19A1 |
| Serotonergic synapse | 8/132 | 0.033291 | 0.0762271 | 2.245045 | 7.63869201 | GABRB3;PLCB3;PLCB4;MAOA;ALOX15;CACNA1A;ALOX12;HTR3A |
| Fc gamma R-mediated phagocytosis | 6/87 | 0.0360652 | 0.0818147 | 2.5741073 | 8.552280213 | SPHK2;INPPL1;PIP5K1A;PIK3R2;PLCG1;PLD2 |
| Butanoate metabolism | 27/3 | 0.0393336 | 0.0884105 | 4.3332589 | 14.02101945 | ACSM4;HMGCS2;ACADS |
| Protein digestion and absorption | 6/90 | 0.0415 | 0.0924319 | 2.4817902 | 7.897208355 | KCNE3;SLC16A10;ATP1B2;ATP1A1;ATP1B1;SLC8A1 |
| Glutamatergic synapse | 7/114 | 0.0422686 | 0.0932956 | 2.2744235 | 7.195616351 | PLCB3;PLCB4;GRIK3;CACNA1A;GRIN2C;PLD2;GRIN2D |
| Cortisol synthesis and secretion | 5/69 | 0.0448105 | 0.0980229 | 2.7123936 | 8.422831272 | PLCB3;PLCB4;PDE8A;CACNA1H;CACNA1G |
| Morphine addiction | 6/92 | 0.045395 | 0.0981227 | 2.4238236 | 7.495320056 | GABRB3;PDE11A;GABRA5;PDE4A;CACNA1A;PDE8A |
| Cocaine addiction | 4/48 | 0.0456571 | 0.0981227 | 3.1538462 | 9.734650188 | MAOA;GRIN2C;SLC6A3;GRIN2D |
| Cholesterol metabolism | 4/49 | 0.0486535 | 0.1036532 | 3.0836017 | 9.3218234 | STARD3;NPC1;NPC2;VDAC3 |
| Arrhythmogenic right ventricular cardiomyopathy (ARVC) | 5/72 | 0.0521674 | 0.1101812 | 2.5905419 | 7.650640397 | CACNG6;CACNB2;CACNB3;CACNG8;SLC8A1 |
| Biosynthesis of unsaturated fatty acids | 3/32 | 0.0602306 | 0.1261238 | 3.5852217 | 10.07295016 | TECR;ACOT1;HSD17B12 |
| Circadian entrainment | 6/99 | 0.0607723 | 0.1261798 | 2.2405745 | 6.27500006 | PLCB3;PLCB4;GRIN2C;CACNA1H;CACNA1G;GRIN2D |
| Starch and sucrose metabolism | 3/33 | 0.0649222 | 0.1336633 | 3.4655357 | 9.476735893 | GPI1;GYS1;PGM2L1 |
| cAMP signaling pathway | 10/211 | 0.0758414 | 0.1548429 | 1.7305876 | 4.463377212 | CNGA2;PDE4A;PIK3R2;ATP1B2;ATP1A1;GRIN2C;ATP1B1;HCN2;GRIN2D;PLD2 |
| PPAR signaling pathway | 5/85 | 0.0916504 | 0.1855731 | 2.1681228 | 5.181323944 | SLC27A1;ACADL;HMGCS2;CPT1C;PCK2 |
| Hypertrophic cardiomyopathy (HCM) | 5/86 | 0.0951868 | 0.1911537 | 2.1412452 | 5.03602539 | CACNG6;CACNB2;CACNB3;CACNG8;SLC8A1 |
| Vascular smooth muscle contraction | 7/140 | 0.1008976 | 0.200975 | 1.8273381 | 4.191271921 | PLA2G2F;PLCB3;PLA2G2D;PLCB4;PLA2G2E;KCNMA1;PLA2G5 |
| Taste transduction | 5/88 | 0.1024646 | 0.2024501 | 2.089433 | 4.760226386 | P2RX3;GABRA5;SCNN1A;CACNA1A;HTR3A |
| Steroid hormone biosynthesis | 5/89 | 0.1062044 | 0.2081607 | 2.0644521 | 4.62930538 | SRD5A1;SRD5A3;CYP1A1;HSD17B12;COMT |
| Dilated cardiomyopathy (DCM) | 5/90 | 0.1100106 | 0.2139095 | 2.040059 | 4.502774744 | CACNG6;CACNB2;CACNB3;CACNG8;SLC8A1 |
| Glycosaminoglycan degradation | 21/2 | 0.1169588 | 0.2256292 | 3.6434938 | 7.818694573 | HPSE;GUSB |
| Long-term potentiation | 4/67 | 0.1198718 | 0.2294422 | 2.2005282 | 4.668051004 | PLCB3;PLCB4;GRIN2C;GRIN2D |
| Amphetamine addiction | 4/68 | 0.1247073 | 0.2368471 | 2.1660331 | 4.509217745 | MAOA;GRIN2C;SLC6A3;GRIN2D |
| Histidine metabolism | 24/2 | 0.1456571 | 0.2745077 | 3.1461676 | 6.061091238 | MAOA;CNDP2 |
| AGE-RAGE signaling pathway in diabetic complications | 5/101 | 0.1559811 | 0.2917203 | 1.8052755 | 3.354239222 | PLCB3;PLCB4;PIK3R2;PLCG1;PLCD3 |
| Cushing syndrome | 7/159 | 0.1622881 | 0.3012165 | 1.5973471 | 2.904587605 | PLCB3;PDE11A;PLCB4;PDE8A;CACNA1H;FH1;CACNA1G |
| Pathways in cancer | 19/535 | 0.1788938 | 0.3290349 | 1.2807049 | 2.204046065 | GSTM4;GSTM3;NQO1;GSTM2;GSTM1;TXNRD3;TXNRD2;GSTP1;TXNRD1;SLC2A1;GSTT2;GSTT1;PIK3R2;FH1;PLCB3;PLCB4;HMOX1;PLCG1;GSTM5 |
| Amyotrophic lateral sclerosis (ALS) | 3/52 | 0.1800053 | 0.3290349 | 2.1196793 | 3.634759971 | GRIN2C;GRIN2D;SOD1 |
| Dopaminergic synapse | 6/135 | 0.181305 | 0.3290349 | 1.6122918 | 2.753108754 | PLCB3;PLCB4;MAOA;CACNA1A;COMT;SLC6A3 |
| RNA degradation | 4/83 | 0.2058273 | 0.3707919 | 1.7534023 | 2.771633985 | PFKL;NUDT16;PFKM;PFKP |
| Cholinergic synapse | 5/113 | 0.2134719 | 0.3817563 | 1.6036937 | 2.47650446 | PLCB3;PLCB4;CHRNA4;CACNA1A;PIK3R2 |
| VEGF signaling pathway | 3/58 | 0.2235763 | 0.396929 | 1.8878571 | 2.828014576 | SPHK2;PIK3R2;PLCG1 |
| Long-term depression | 3/61 | 0.2460846 | 0.4337463 | 1.7899323 | 2.509627858 | PLCB3;PLCB4;CACNA1A |
| Retinol metabolism | 4/91 | 0.254066 | 0.4446155 | 1.5915119 | 2.180627753 | AWAT2;CYP1A1;ADH5;DHRS4 |
| Oxytocin signaling pathway | 6/154 | 0.2671225 | 0.4641491 | 1.4039255 | 1.853248691 | CACNG6;CACNB2;CACNB3;PLCB3;CACNG8;PLCB4 |
| Synthesis and degradation of ketone bodies | 11/1 | 0.2696058 | 0.465165 | 3.4567616 | 4.531103427 | HMGCS2 |
| Inflammatory mediator regulation of TRP channels | 5/127 | 0.2873372 | 0.4922911 | 1.4186351 | 1.769177978 | PLCB3;PLCB4;PIK3R2;ALOX12;PLCG1 |
| Renal cell carcinoma | 3/68 | 0.2997078 | 0.50808 | 1.5965934 | 1.923810857 | SLC2A1;PIK3R2;FH1 |
| African trypanosomiasis | 2/39 | 0.3007004 | 0.50808 | 1.8692489 | 2.246165719 | PLCB3;PLCB4 |
| Adipocytokine signaling pathway | 3/71 | 0.322919 | 0.5418846 | 1.5259191 | 1.724828566 | SLC2A1;CPT1C;PCK2 |
| Fatty acid biosynthesis | 14/1 | 0.3296024 | 0.5493374 | 2.6586367 | 2.950736018 | MCAT |
| Apelin signaling pathway | 5/138 | 0.3482127 | 0.5764331 | 1.3005632 | 1.372018623 | PLCB3;PLCB4;SPHK2;PIK3C3;SLC8A1 |
| Parathyroid hormone synthesis, secretion and action | 4/107 | 0.3554801 | 0.5845142 | 1.3431752 | 1.389227385 | PLCB3;PLCB4;PDE4A;PLD2 |
| Renin secretion | 3/76 | 0.3615757 | 0.5905736 | 1.4210372 | 1.445598296 | PLCB3;PLCB4;KCNMA1 |
| Rheumatoid arthritis | 3/84 | 0.422565 | 0.6856187 | 1.2801587 | 1.102744192 | TCIRG1;ATP6V0C;ATP6V1C1 |
| GABAergic synapse | 3/90 | 0.4669829 | 0.747783 | 1.1915025 | 0.907284723 | GABRB3;GABRA5;CACNA1A |
| GnRH signaling pathway | 3/90 | 0.4669829 | 0.747783 | 1.1915025 | 0.907284723 | PLCB3;PLCB4;PLD2 |
| Ras signaling pathway | 7/233 | 0.4844805 | 0.7707644 | 1.0701996 | 0.775550251 | PLA2G2F;PLA2G2D;PLA2G2E;PIK3R2;PLA2G5;PLCG1;PLD2 |
| Lysine degradation | 2/59 | 0.4977743 | 0.7868045 | 1.2121212 | 0.845586166 | GCDH;DHTKD1 |
| Ascorbate and aldarate metabolism | 27/1 | 0.5376625 | 0.8390274 | 1.3284287 | 0.824322293 | UGDH |
| Maturity onset diabetes of the young | 27/1 | 0.5376625 | 0.8390274 | 1.3284287 | 0.824322293 | SLC2A2 |
| Protein export | 28/1 | 0.5506948 | 0.8493149 | 1.2791617 | 0.763115242 | OXA1L |
| Insulin signaling pathway | 4/139 | 0.5525476 | 0.8493149 | 1.0230968 | 0.606917153 | GYS1;INPPL1;PIK3R2;PCK2 |
| Chagas disease (American trypanosomiasis) | 3/103 | 0.5574106 | 0.8493149 | 1.0359107 | 0.605441276 | PLCB3;PLCB4;PIK3R2 |
| Non-small cell lung cancer | 2/66 | 0.5581212 | 0.8493149 | 1.0791555 | 0.629340977 | PIK3R2;PLCG1 |

| | | | | | | |
|---|---|---|---|---|---|---|
| Fc epsilon RI signaling pathway | 2/68 | 0.5743888 | 0.8618903 | 1.0463458 | 0.580145163 | PIK3R2;PLCG1 |
| Phagosome | 5/180 | 0.5746469 | 0.8618903 | 0.9862775 | 0.546397323 | PIK3C3;TCIRG1;MPO;ATP6V0C;ATP6V1C1 |
| Amoebiasis | 3/106 | 0.5769388 | 0.8618903 | 1.0055825 | 0.553089569 | PLCB3;PLCB4;PIK3R2 |
| Acute myeloid leukemia | 2/69 | 0.5823574 | 0.8647126 | 1.0306755 | 0.557256221 | PIK3R2;MPO |
| p53 signaling pathway | 2/71 | 0.5979634 | 0.8825363 | 1.0006975 | 0.514584408 | RRM2;CHEK2 |
| B cell receptor signaling pathway | 2/72 | 0.6056006 | 0.8831675 | 0.9863509 | 0.494689102 | INPPL1;PIK3R2 |
| Prolactin signaling pathway | 2/72 | 0.6056006 | 0.8831675 | 0.9863509 | 0.494689102 | SLC2A2;PIK3R2 |
| Prion diseases | 1/34 | 0.6215384 | 0.9010468 | 1.0462633 | 0.497558437 | SOD1 |
| Glioma | 2/75 | 0.6278501 | 0.9048428 | 0.9456694 | 0.440165433 | PIK3R2;PLCG1 |
| Primary immunodeficiency | 1/36 | 0.6425816 | 0.9206579 | 0.9863752 | 0.436235717 | ADA |
| Neuroactive ligand-receptor interaction | 9/348 | 0.6486205 | 0.9239072 | 0.9152104 | 0.396201369 | GABRB3;P2RX4;P2RX3;GLRB;GABRA5;CHRNA4;GRIK3;GRIN2C;GRIN2D |
| Neurotrophin signaling pathway | 3/121 | 0.6661203 | 0.9433496 | 0.8770732 | 0.356341633 | PSEN2;PIK3R2;PLCG1 |
| Protein processing in endoplasmic reticulum | 4/163 | 0.6771767 | 0.9534959 | 0.867587 | 0.338205427 | EDEM2;MAN1C1;MOGS;MAN1B1 |
| Platelet activation | 3/125 | 0.687449 | 0.9587384 | 0.8481411 | 0.31785586 | PLCB3;PLCB4;PIK3R2 |
| ErbB signaling pathway | 2/84 | 0.6887264 | 0.9587384 | 0.8414852 | 0.313799263 | PIK3R2;PLCG1 |
| Gap junction | 2/86 | 0.7010886 | 0.9691726 | 0.8213649 | 0.291683904 | PLCB3;PLCB4 |
| Rap1 signaling pathway | 5/209 | 0.7041336 | 0.9691726 | 0.8447976 | 0.296344175 | PLCB3;PLCB4;PRKD2;PIK3R2;PLCG1 |
| Relaxin signaling pathway | 3/131 | 0.7175028 | 0.982057 | 0.8081334 | 0.268282806 | PLCB3;PLCB4;PIK3R2 |
| MAPK signaling pathway | 7/294 | 0.7247729 | 0.9864964 | 0.8400597 | 0.270412634 | CACNG6;CACNB2;CACNB3;CACNG8;CACNA1A;CACNA1H;CACNA1G |
| Estrogen signaling pathway | 3/134 | 0.7316682 | 0.9877073 | 0.7895038 | 0.246663208 | PLCB3;PLCB4;PIK3R2 |
| Necroptosis | 4/176 | 0.7337254 | 0.9877073 | 0.8014727 | 0.248152335 | ALOX15;VDAC3;TRPM7;PGAM5 |
| Tuberculosis | 4/178 | 0.7417282 | 0.993024 | 0.7921782 | 0.236681016 | SPHK2;PIK3C3;TCIRG1;ATP6V0C |
| Notch signaling pathway | 1/49 | 0.7536097 | 0.9999952 | 0.71875 | 0.20332049 | PSEN2 |
| Prostate cancer | 2/97 | 0.7619536 | 0.9999952 | 0.7258467 | 0.19733566 | GSTP1;PIK3R2 |
| Cellular senescence | 4/185 | 0.7683002 | 0.9999952 | 0.7612647 | 0.200650101 | CHEK2;VDAC3;PIK3R2;TRPM7 |
| Melanogenesis | 2/100 | 0.7765753 | 0.9999952 | 0.7035178 | 0.177892683 | PLCB3;PLCB4 |
| T cell receptor signaling pathway | 2/101 | 0.7812718 | 0.9999952 | 0.6963755 | 0.171887845 | PIK3R2;PLCG1 |
| Longevity regulating pathway | 2/102 | 0.7858816 | 0.9999952 | 0.6893761 | 0.166104553 | PIK3R2;SOD1 |
| Regulation of lipolysis in adipocytes | 1/56 | 0.7983541 | 0.9999952 | 0.6270463 | 0.141212701 | PIK3R2 |
| Ovarian steroidogenesis | 1/57 | 0.8040464 | 0.9999952 | 0.6158172 | 0.134308723 | CYP1A1 |
| Endometrial cancer | 1/58 | 0.8095782 | 0.9999952 | 0.6049822 | 0.127797614 | PIK3R2 |
| mTOR signaling pathway | 3/154 | 0.8121931 | 0.9999952 | 0.6842242 | 0.142330401 | SLC7A5;PIK3R2;ATP6V1C1 |
| Alcoholism | 4/199 | 0.814973 | 0.9999952 | 0.7060961 | 0.144467429 | MAOA;GRIN2C;SLC6A3;GRIN2D |
| TNF signaling pathway | 2/110 | 0.8197752 | 0.9999952 | 0.6380471 | 0.12679597 | PIK3R2;PGAM5 |
| Proteoglycans in cancer | 4/203 | 0.8268025 | 0.9999952 | 0.6917593 | 0.131565326 | PIK3R2;PLCG1;HPSE;NUDT16L1 |
| NOD-like receptor signaling pathway | 4/205 | 0.8324785 | 0.9999952 | 0.684805 | 0.125557572 | PLCB3;PLCB4;VDAC3;TRPM7 |
| Mitophagy | 1/63 | 0.8349847 | 0.9999952 | 0.5560498 | 0.100279071 | PGAM5 |
| Leukocyte transendothelial migration | 2/115 | 0.83844 | 0.9999952 | 0.6096572 | 0.107429105 | PIK3R2;PLCG1 |
| Natural killer cell mediated cytotoxicity | 2/118 | 0.8487822 | 0.9999952 | 0.593798 | 0.097354785 | PIK3R2;PLCG1 |
| Regulation of actin cytoskeleton | 4/217 | 0.8633613 | 0.9999952 | 0.6458213 | 0.094885357 | PIP5K1A;PIP4K2B;PIK3R2;PIP4K2C |
| Melanoma | 1/72 | 0.8724918 | 0.9999952 | 0.4853391 | 0.066201237 | PIK3R2 |
| Bacterial invasion of epithelial cells | 1/74 | 0.8795947 | 0.9999952 | 0.4719934 | 0.060553915 | PIK3R2 |
| Pancreatic cancer | 1/75 | 0.8829967 | 0.9999952 | 0.465591 | 0.057935275 | PIK3R2 |
| Autophagy | 2/130 | 0.8843873 | 0.9999952 | 0.5377952 | 0.066073629 | PIK3R2;PIK3C3 |
| Axon guidance | 3/180 | 0.8854938 | 0.9999952 | 0.5829298 | 0.070889987 | DPYSL2;PIK3R2;PLCG1 |
| Chronic myeloid leukemia | 1/76 | 0.8863027 | 0.9999952 | 0.4593594 | 0.055443196 | PIK3R2 |
| FoxO signaling pathway | 2/132 | 0.8895083 | 0.9999952 | 0.5294666 | 0.061993375 | PIK3R2;PCK2 |
| Transcriptional misregulation in cancer | 3/183 | 0.8920576 | 0.9999952 | 0.573125 | 0.065464958 | FUT8;SLC45A3;MPO |
| Human T-cell leukemia virus 1 infection | 4/245 | 0.9170314 | 0.9999952 | 0.5699567 | 0.049366005 | CHEK2;VDAC3;SLC2A1;PIK3R2 |
| Th1 and Th2 cell differentiation | 1/87 | 0.9170566 | 0.9999952 | 0.4003766 | 0.034667038 | PLCG1 |
| Chemokine signaling pathway | 3/197 | 0.9184338 | 0.9999952 | 0.5313789 | 0.045212586 | PLCB3;PLCB4;PIK3R2 |
| Colorectal cancer | 1/88 | 0.9194016 | 0.9999952 | 0.3957541 | 0.033256105 | PIK3R2 |
| Progesterone-mediated oocyte maturation | 1/90 | 0.9238949 | 0.9999952 | 0.3868207 | 0.030619547 | PIK3R2 |
| Small cell lung cancer | 1/92 | 0.9281381 | 0.9999952 | 0.3782801 | 0.028210123 | PIK3R2 |
| mRNA surveillance pathway | 1/96 | 0.9359291 | 0.9999952 | 0.3622776 | 0.023988394 | NUDT21 |
| Toll-like receptor signaling pathway | 1/99 | 0.9412138 | 0.9999952 | 0.351133 | 0.021273387 | PIK3R2 |
| Wnt signaling pathway | 2/160 | 0.9422385 | 0.9999952 | 0.4350053 | 0.025881447 | PLCB3;PLCB4 |
| Kaposi sarcoma-associated herpesvirus infection | 3/216 | 0.94486 | 0.9999952 | 0.483501 | 0.027423449 | PIK3R2;PIK3C3;PLCG1 |
| NF-kappa B signaling pathway | 1/102 | 0.9460632 | 0.9999952 | 0.3406504 | 0.018887665 | PLCG1 |
| Th17 cell differentiation | 1/102 | 0.9460632 | 0.9999952 | 0.3406504 | 0.018887665 | PLCG1 |
| Hepatitis B | 2/163 | 0.9461924 | 0.9999952 | 0.4268332 | 0.023607854 | VDAC3;PIK3R2 |
| MicroRNAs in cancer | 4/281 | 0.9580748 | 0.9999952 | 0.494953 | 0.021198528 | ABCC1;SLC45A3;HMOX1;PLCG1 |
| Viral carcinogenesis | 3/229 | 0.9580994 | 0.9999952 | 0.4553808 | 0.01949203 | PKM;VDAC3;PIK3R2 |
| C-type lectin receptor signaling pathway | 1/112 | 0.9595229 | 0.9999952 | 0.3098009 | 0.012800704 | PIK3R2 |
| Ribosome biogenesis in eukaryotes | 1/115 | 0.962864 | 0.9999952 | 0.3016014 | 0.011413534 | NAT10 |
| Cell cycle | 1/123 | 0.9704882 | 0.9999952 | 0.2817076 | 0.00843885 | CHEK2 |
| Osteoclast differentiation | 1/128 | 0.9744376 | 0.9999952 | 0.2705467 | 0.007005756 | PIK3R2 |
| Human papillomavirus infection | 5/360 | 0.9756013 | 0.9999952 | 0.4816498 | 0.011897357 | PKM;PIK3R2;TCIRG1;ATP6V0C;ATP6V1C1 |
| Human cytomegalovirus infection | 3/255 | 0.9761466 | 0.9999952 | 0.4078444 | 0.009846393 | PLCB3;PLCB4;PIK3R2 |
| Signaling pathways regulating pluripotency of stem cells | 1/137 | 0.9802642 | 0.9999952 | 0.2525251 | 0.005033622 | PIK3R2 |
| Apoptosis | 1/141 | 0.9824083 | 0.9999952 | 0.2452593 | 0.004352924 | PIK3R2 |
| Measles | 1/144 | 0.9838623 | 0.9999952 | 0.2400766 | 0.003905878 | PIK3R2 |
| Breast cancer | 1/147 | 0.9851963 | 0.9999952 | 0.235107 | 0.003506461 | PIK3R2 |
| Gastric cancer | 1/150 | 0.9864202 | 0.9999952 | 0.2303375 | 0.003149358 | PIK3R2 |
| Hepatitis C | 1/160 | 0.9898157 | 0.9999952 | 0.2157389 | 0.002208419 | PIK3R2 |
| JAK-STAT signaling pathway | 1/164 | 0.9909232 | 0.9999952 | 0.2104011 | 0.001918488 | PIK3R2 |
| Human immunodeficiency virus 1 infection | 2/238 | 0.9914333 | 0.9999952 | 0.2900541 | 0.002495518 | PIK3R2;PLCG1 |
| Influenza A | 1/168 | 0.9919104 | 0.9999952 | 0.2053189 | 0.001667695 | PIK3R2 |
| Endocytosis | 2/269 | 0.9961 | 0.9999952 | 0.2559635 | 0.001000198 | PIP5K1A;PLD2 |
| Focal adhesion | 1/199 | 0.9966878 | 0.9999952 | 0.1728944 | 5.74E-04 | PIK3R2 |
| PI3K-Akt signaling pathway | 3/357 | 0.9977219 | 0.9999952 | 0.2887863 | 6.59E-04 | GYS1;PIK3R2;PCK2 |
| Epstein-Barr virus infection | 1/229 | 0.9986051 | 0.9999952 | 0.149911 | 2.09E-04 | PIK3R2 |
| Herpes simplex virus 1 infection | 1/433 | 0.9999927 | 0.9999952 | 0.0782795 | 5.69E-07 | PIK3R2 |
| Olfactory transduction | 2/1133 | 0.9999952 | 0.9999952 | 0.0577029 | 2.79E-07 | CNGA2;SLC8A1 |
