## Supplementary material for "Glutamine antagonism suppresses tumor growth in adrenocortical carcinoma through inhibition of de novo nucleotide biosynthesis": Suppl. Table 1B

**Table 1B human up Up-regulated pathways Human ACC (TCGA) versus normal adrenals**

| Term | Overlap | P-value | Adjusted P-v | Odds Ratio | Combined Score | Genes |
|---|---|---|---|---|---|---|
| Oxidative phosphorylation | 81/133 | 1.54E-66 | 3.92E-64 | 27.7756471 | 4209.1504 | [illegible] |
| Parkinson disease | 69/142 | 1.02E-47 | 1.30E-45 | 16.64606951 | 1801.11848 | [illegible] |
| Non-alcoholic fatty liver disease (NAFLD) | 65/149 | 2.65E-41 | 2.25E-39 | 13.56876084 | 1267.748689 | [illegible] |
| Alzheimer disease | 69/171 | 4.52E-41 | 2.88E-39 | 11.89497767 | 1105.014462 | [illegible] |
| Thermogenesis | 76/231 | 8.55E-38 | 4.36E-36 | 8.654193548 | 738.6603179 | [illegible] |
| Huntington disease | 69/193 | 4.81E-37 | 2.04E-35 | 9.773105179 | 817.2718948 | [illegible] |
| Drug metabolism | 49/108 | 2.09E-32 | 7.62E-31 | 14.36772021 | 1048.05689 | [illegible] |
| Glycolysis / Gluconeogenesis | 36/68 | 3.66E-27 | 1.17E-25 | 19.26 | 1172.379072 | [illegible] |
| Purine metabolism | 47/129 | 5.10E-26 | 1.44E-24 | 9.885484557 | 575.713964 | [illegible] |
| Steroid hormone biosynthesis | 32/60 | 2.01E-24 | 5.13E-23 | 19.4989648 | 1063.915419 | [illegible] |
| Retrograde endocannabinoid signaling | 48/148 | 5.65E-24 | 1.31E-22 | 8.278235294 | 443.139699 | [illegible] |
| Chemical carcinogenesis | 36/82 | 1.55E-23 | 3.30E-22 | 13.3883004 | 703.1409081 | [illegible] |
| Glycosaminoglycan biosynthesis | 29/53 | 1.11E-22 | 2.18E-21 | 20.56458898 | 1039.576882 | [illegible] |
| Citrate cycle (TCA cycle) | 22/30 | 1.23E-21 | 2.23E-20 | 46.5475763 | 2241.310633 | [illegible] |
| Metabolism of xenobiotics by cytochrome P450 | 32/74 | 8.03E-21 | 1.37E-19 | 12.98964803 | 601.0431339 | [illegible] |
| Retinol metabolism | 29/67 | 5.20E-19 | 8.29E-18 | 12.97850996 | 546.3907234 | [illegible] |
| N-Glycan biosynthesis | 25/50 | 1.80E-18 | 2.70E-17 | 16.95679568 | 692.8573576 | [illegible] |
| Lysosome | 38/123 | 1.95E-18 | 2.76E-17 | 7.646008786 | 311.7884516 | [illegible] |
| Cardiac muscle contraction | 29/78 | 7.59E-17 | 1.02E-15 | 10.05908597 | 373.3686321 | [illegible] |
| Amino sugar and nucleotide sugar metabolism | 23/48 | 1.42E-16 | 1.81E-15 | 15.57221923 | 568.2060576 | [illegible] |
| Pyrimidine metabolism | 24/57 | 1.24E-15 | 1.51E-14 | 12.31589274 | 422.7187046 | [illegible] |
| Collecting duct acid secretion | 17/27 | 2.92E-15 | 3.38E-14 | 28.6432529 | 958.6539153 | [illegible] |
| Glycosphingolipid biosynthesis | 21/45 | 5.91E-15 | 6.55E-14 | 14.78475336 | 484.3815954 | [illegible] |
| Glycerophospholipid metabolism | 30/97 | 7.37E-15 | 7.83E-14 | 7.609916062 | 247.6381306 | [illegible] |
| Glutathione metabolism | 23/56 | 9.40E-15 | 9.58E-14 | 11.79212611 | 380.8684314 | [illegible] |
| Pentose phosphate pathway | 17/30 | 3.52E-14 | 3.45E-13 | 22.02976559 | 682.4490541 | [illegible] |
| Synaptic vesicle cycle | 26/78 | 6.25E-14 | 5.90E-13 | 8.473873874 | 257.6427437 | [illegible] |
| Glycine, serine and threonine metabolism | 19/40 | 7.79E-14 | 7.09E-13 | 15.26269344 | 460.6831556 | [illegible] |
| Butanoate metabolism | 16/28 | 1.69E-13 | 1.48E-12 | 22.44285714 | 660.0691065 | [illegible] |
| Cysteine and methionine metabolism | 20/47 | 2.33E-13 | 1.96E-12 | 12.50298686 | 363.6888985 | [illegible] |
| Porphyrin and chlorophyll metabolism | 19/42 | 2.38E-13 | 1.96E-12 | 13.93402359 | 405.0309831 | [illegible] |
| Peroxisome | 26/83 | 3.29E-13 | 2.62E-12 | 7.728496918 | 222.1344173 | [illegible] |
| Pyruvate metabolism | 18/39 | 6.63E-13 | 5.12E-12 | 14.44646052 | 405.1111958 | [illegible] |
| Alanine, aspartate and glutamate metabolism | 17/35 | 1.02E-12 | 7.62E-12 | 15.90616622 | 439.2467837 | [illegible] |
| Sphingolipid metabolism | 19/47 | 2.83E-12 | 2.06E-11 | 11.44276762 | 304.2868294 | [illegible] |
| Arginine and proline metabolism | 19/49 | 6.85E-12 | 4.85E-11 | 10.67878245 | 274.5122575 | [illegible] |
| Fatty acid degradation | 18/44 | 8.34E-12 | 5.75E-11 | 11.66519884 | 297.5767407 | [illegible] |
| Valine, leucine and isoleucine degradation | 18/48 | 4.77E-11 | 3.20E-10 | 10.10769231 | 240.2269015 | [illegible] |
| Vibrio cholerae infection | 18/50 | 1.06E-10 | 6.91E-10 | 9.474955277 | 217.6502772 | [illegible] |
| Pentose and glucuronate interconversions | 15/34 | 1.26E-10 | 8.04E-10 | 13.27174985 | 302.5178188 | [illegible] |
| Bile secretion | 21/72 | 2.77E-10 | 1.72E-09 | 6.947560011 | 152.887596 | [illegible] |
| Terpenoid backbone biosynthesis | 22/12 | 4.05E-10 | 2.46E-09 | 20.12882562 | 435.3186539 | [illegible] |
| Other glycan degradation | 18/11 | 4.17E-10 | 2.47E-09 | 26.33993651 | 568.9128898 | [illegible] |
| Tryptophan metabolism | 16/42 | 4.33E-10 | 2.51E-09 | 10.35054945 | 223.1543236 | [illegible] |
| Biosynthesis of unsaturated fatty acids | 13/27 | 5.69E-10 | 3.16E-09 | 15.58643938 | 331.7866369 | [illegible] |
| Fatty acid elongation | 13/27 | 5.69E-10 | 3.16E-09 | 15.58643938 | 331.7866369 | [illegible] |
| Glycosaminoglycan degradation | 19/11 | 9.39E-10 | 5.09E-09 | 23.04622222 | 479.0548176 | [illegible] |
| Central carbon metabolism in cancer | 19/65 | 1.88E-09 | 9.98E-09 | 6.95850687 | 139.8187332 | [illegible] |
| Glycosylphosphatidylinositol (GPI)-anchor biosynthesis | 25/12 | 2.78E-09 | 1.45E-08 | 15.48124829 | 305.0007894 | [illegible] |
| Arginine biosynthesis | 21/11 | 3.94E-09 | 2.01E-08 | 18.43502222 | 356.7516655 | [illegible] |
| Galactose metabolism | 13/31 | 4.73E-09 | 2.32E-08 | 12.12021371 | 232.3427806 | [illegible] |
| Mucin type O-glycan biosynthesis | 13/31 | 4.73E-09 | 2.32E-08 | 12.12021371 | 232.3427806 | [illegible] |
| Ascorbate and aldarate metabolism | 27/12 | 8.35E-09 | 4.02E-08 | 13.41565836 | 249.5397204 | [illegible] |
| Fructose and mannose metabolism | 13/33 | 1.18E-08 | 5.58E-08 | 10.90703473 | 199.1016172 | [illegible] |
| Ubiquinone and other terpenoid-quinone biosynthesis | 11/8 | 1.50E-08 | 6.94E-08 | 44.58865248 | 803.3774502 | [illegible] |
| Steroid biosynthesis | 19/10 | 1.94E-08 | 8.83E-08 | 18.60568384 | 330.4082409 | [illegible] |
| Ether lipid metabolism | 15/47 | 2.54E-08 | 1.14E-07 | 7.874665477 | 137.7205129 | [illegible] |
| Glyoxylate and dicarboxylate metabolism | 30/12 | 3.54E-08 | 1.56E-07 | 11.17793594 | 191.7598805 | [illegible] |
| Tyrosine metabolism | 13/36 | 4.06E-08 | 1.75E-07 | 9.482868094 | 161.3993469 | [illegible] |
| Propanoate metabolism | 12/32 | 8.32E-08 | 3.54E-07 | 10.05907473 | 163.9794934 | [illegible] |
| Nitrogen metabolism | 17/9 | 9.58E-08 | 4.00E-07 | 18.82253771 | 304.1954197 | [illegible] |
| Inositol phosphate metabolism | 18/74 | 1.16E-07 | 4.77E-07 | 5.407360082 | 86.35712387 | [illegible] |
| Epithelial cell signaling in Helicobacter pylori infection | 17/68 | 1.69E-07 | 6.85E-07 | 5.604110813 | 87.38256161 | [illegible] |
| Glycerolipid metabolism | 16/61 | 1.90E-07 | 7.56E-07 | 5.974285714 | 92.4702188 | [illegible] |
| Proximal tubule bicarbonate reclamation | 23/10 | 1.95E-07 | 7.64E-07 | 12.87812543 | 198.983418 | [illegible] |
| Fat digestion and absorption | 13/41 | 2.37E-07 | 9.16E-07 | 7.787431624 | 118.7953473 | [illegible] |
| Arachidonic acid metabolism | 16/63 | 3.08E-07 | 1.17E-06 | 5.719452888 | 85.75776493 | [illegible] |
| Starch and sucrose metabolism | 12/36 | 3.73E-07 | 1.40E-06 | 8.380782918 | 124.0421837 | [illegible] |
| Pancreatic secretion | 20/98 | 5.10E-07 | 1.89E-06 | 4.316239316 | 62.53622489 | [illegible] |
| One carbon pool by folate | 20/9 | 5.66E-07 | 2.06E-06 | 13.68694039 | 196.8707677 | [illegible] |
| Folate biosynthesis | 26/10 | 7.73E-07 | 2.78E-06 | 10.46181172 | 147.2254063 | [illegible] |
| ABC transporters | 13/45 | 7.94E-07 | 2.81E-06 | 6.812555654 | 95.68812078 | [illegible] |
| Calcium signaling pathway | 29/188 | 8.90E-07 | 3.11E-06 | 3.081846227 | 42.93645492 | [illegible] |
| Ferroptosis | 12/40 | 1.35E-06 | 4.65E-06 | 7.18200305 | 97.07430315 | [illegible] |
| Aldosterone synthesis and secretion | 19/98 | 2.21E-06 | 7.52E-06 | 4.044683431 | 52.67110826 | [illegible] |
| Ovarian steroidogenesis | 13/49 | 2.31E-06 | 7.75E-06 | 6.054318789 | 78.57362171 | [illegible] |
| PPAR signaling pathway | 16/74 | 3.13E-06 | 1.04E-05 | 4.632019704 | 58.70806482 | [illegible] |
| Mineral absorption | 13/51 | 3.77E-06 | 1.23E-05 | 5.735061161 | 71.62423012 | [illegible] |
| Carbohydrate digestion and absorption | 12/44 | 4.12E-06 | 1.33E-05 | 6.282918149 | 77.90242133 | G6PC3;AMY1A;ATP1A3;PIK3R2;PIK3CB;PIK3R1;SLC5A1;ATP1A1;SLC37A4;ATP1B1;HK2;HK1 |
| Synthesis and degradation of ketone bodies | 10/6 | 5.71E-06 | 1.82E-05 | 25.03539823 | 302.2539547 | BDH2;BDH1;HMGCS1;OXCT1;OXCT2;ACAT2 |
| Nicotine addiction | 11/40 | 9.51E-06 | 2.99E-05 | 6.350498084 | 73.43170935 | GRIN3A;CHRNA4;GABRA5;CHRNA7;CHRNA6;GABRA3;SLC17A8;GRIN2C;GABRD;GABRG1;GRIN2D |
| Rheumatoid arthritis | 17/91 | 1.25E-05 | 3.90E-05 | 3.857570707 | 43.53581872 | ATP6V1A;ATP6V1G1;ATP6V0B;ATP6V0E1;ATP6AP1;ATP6V0A2;ATP6V1H;ATP6V0A4;ATP6V1E1;ATP6V0D1;ATP6V0E2;ATP6V0C;ATP6V0D2;ATP6V1D;ATP6V1C1;ATP6V1B1;ATP6V1F |
| Primary bile acid biosynthesis | 17/7 | 2.20E-05 | 6.76E-05 | 11.68981399 | 125.3665194 | CYP39A1;CYP27A1;HSD3B7;ACOT8;AMACR;AKR1D1;SLC27A5 |
| Histidine metabolism | 23/8 | 2.41E-05 | 7.31E-05 | 8.912056738 | 94.77599208 | ALDH3B2;MAOB;UROC1;ALDH1B1;MAOA;HDC;HNMT;ALDH7A1 |
| Sulfur relay system | 8/5 | 2.84E-05 | 8.52E-05 | 27.79398762 | 290.9685556 | MPST;NFS1;MOCS2;MOCS3;TST |
| GABAergic synapse | 16/89 | 3.68E-05 | 1.09E-04 | 3.677299413 | 37.54394225 | SLC38A1;KCNJ6;SLC12A5;GLS2;GABRA5;GAD1;GABRA3;SLC6A13;ADCY3;ABAT;SLC6A11;SLC6A1;ADCY8;ADCY7;GABRG1;GABRD |
| beta-Alanine metabolism | 9/31 | 3.85E-05 | 1.13E-04 | 6.839477293 | 69.51536228 | ALDH3B2;ECHS1;ALDH1B1;SMOX;GAD1;EHHADH;SMS;ABAT;ALDH7A1 |
| Phosphatidylinositol signaling system | 17/99 | 3.93E-05 | 1.14E-04 | 3.479740186 | 35.30299031 | CDS1;IPPK;PI4K2B;ITPK1;PIK3R2;PIK3R1;PIK3CB;INPP5A;IMPA1;ITPKA;PLCE1;PIP5K1B;PIP4K2C;PIP5K1C;CDIPT;PLCD4;PI4K2A |
| Salivary secretion | 16/90 | 4.24E-05 | 1.22E-04 | 3.627413127 | 36.51827671 | AMY1A;ATP1A3;ADCY3;ATP2B3;LPO;ATP2B2;SLC4A2;ATP1A1;ATP2B1;ATP1B1;ADCY8;LYZ;ADCY7;SLC9A1;KCNMA1;KCNN4 |
| Morphine addiction | 16/91 | 4.88E-05 | 1.38E-04 | 3.578857143 | 35.52845945 | KCNJ6;GABRA5;PDE4D;PDE3B;GABRA3;PDE2A;ADCY3;ADCY8;ADCY7;GABRG1;PDE11A;PDE4B;PDE7B;PDE8B;PDE7A;GABRD |
| Caffeine metabolism | 5/4 | 4.94E-05 | 1.39E-04 | 66.65371025 | 660.8594903 | CYP2A6;NAT1;NAT2;XDH |
| cGMP-PKG signaling pathway | 22/166 | 1.84E-04 | 5.11E-04 | 2.567324955 | 22.07705066 | TRPC6;PDE2A;PDE3B;ATP2A3;ATP1A3;ADCY3;ATP2B3;ATP2B2;ATP1A1;ATP2B1;ATP1B1;ADCY8;ADCY7;KCNU1;KCNMA1;VDAC3;KCNMB3;VDAC2;VDAC1;SLC25A5;SLC25A4;SLC25A6 |
| Nicotinate and nicotinamide metabolism | 30/8 | 2.01E-04 | 5.52E-04 | 6.074145712 | 51.6949962 | NT5C;NNT;QPRT;NT5M;ENPP3;NMNAT2;NT5C2;NUDT12 |
| Phenylalanine metabolism | 17/6 | 2.38E-04 | 6.47E-04 | 9.100402253 | 75.91182849 | ALDH3B2;MAOB;GOT1;MAOA;PAH;GOT2 |
| Insulin secretion | 14/86 | 3.29E-04 | 8.82E-04 | 3.256684492 | 26.12013309 | RYR2;SLC2A1;ATP1A3;ADCY3;ATP1A1;ATP1B1;ADCY8;ADCY7;GCK;KCNU1;KCNMA1;KCNMB3;KCNN1;KCNN4 |
| cAMP signaling pathway | 25/212 | 4.38E-04 | 0.0011628 | 2.247457366 | 17.38154886 | RYR2;PDE3B;ADCY3;ATP1A3;PIK3R2;PIK3R1;PIK3CB;ATP1A1;ADCY8;ADCY7;SLC9A1;LIPE;PDE4B;PLCE1;ABCC4;PDE4D;ATP2B3;ATP2B2;ATP2B1;GRIN2C;ATP1B1;GRIN2D;GRIN3A;ORAI1;HCN2 |
| HIF-1 signaling pathway | 15/100 | 5.07E-04 | 0.0013331 | 2.956236553 | 22.42834642 | PDHA1;SLC2A1;PIK3R2;ENO1;PIK3R1;PIK3CB;PDHB;ENO2;HK2;HK1;LDHA;PFKL;PGK1;ALDOA;GAPDH |
| Cholinergic synapse | 16/112 | 5.83E-04 | 0.001518 | 2.792857143 | 20.7974821 | KCNJ4;ACHE;KCNJ6;CHRNA4;KCNJ12;CHRNA7;CHRNA6;KCNJ14;ADCY3;PIK3R2;PIK3R1;PIK3CB;ADCY8;ADCY7;KCNJ2;SLC18A3 |
| Riboflavin metabolism | 8/4 | 6.03E-04 | 0.0015522 | 16.66077739 | 123.5267945 | FLAD1;RFK;BLVRB;ENPP3 |
| Aldosterone-regulated sodium reabsorption | 8/37 | 9.32E-04 | 0.0023771 | 4.606260699 | 32.1422677 | HSD11B2;SCNN1G;ATP1A3;PIK3R2;PIK3CB;PIK3R1;ATP1A1;ATP1B1 |
| Linoleic acid metabolism | 29/7 | 9.69E-04 | 0.0024467 | 5.310169901 | 36.84807746 | CYP2J2;PLA2G12A;PLA2G2D;PLA2G4D;PLA2G1B;PLA2G4C;CYP3A4 |
| Cortisol synthesis and secretion | 11/65 | 0.0010015 | 0.0024889 | 3.405925926 | 23.52205477 | CYP21A2;CYP11A1;HSD3B1;ORAI1;ADCY3;PDE8B;ADCY8;ADCY7;CACNA1H;CACNA1G;CYP17A1 |
| Gastric acid secretion | 12/75 | 0.0010053 | 0.0024889 | 3.186070158 | 21.99166971 | KCNE2;KCNJ15;ATP1A3;ADCY3;KCNJ16;SLC4A2;ATP1A1;ATP1B1;ADCY8;ADCY7;KCNJ2;SLC9A1 |
| AMPK signaling pathway | 16/120 | 0.0012456 | 0.003054 | 2.576923077 | 17.23490482 | PFKFB4;SCD5;PIK3R2;PIK3R1;SLC2A4;PIK3CB;HMGCR;CPT1C;LIPE;GYS1;PFKL;G6PC3;SCD;PFKM;PFKP;PCK2 |
| Choline metabolism in cancer | 14/99 | 0.0013883 | 0.0033715 | 2.756694977 | 18.13824633 | SLC44A5;PCYT1B;SLC22A5;CHKA;SLC44A1;PLA2G4D;SLC44A2;PLA2G4C;PIK3R2;PIK3R1;PIK3CB;PIP5K1B;CHPT1;PIP5K1C |
| Neomycin, kanamycin and gentamicin biosynthesis | 5/3 | 0.0016761 | 0.0039944 | 24.9717564 | 159.6020317 | HK2;GCK;HK1 |
| Phenylalanine, tyrosine and tryptophan biosynthesis | 5/3 | 0.0016761 | 0.0039944 | 24.9717564 | 159.6020317 | GOT1;PAH;GOT2 |
| Cholesterol metabolism | 9/50 | 0.0018131 | 0.0042809 | 3.666262687 | 23.1440802 | CYP27A1;LIPC;NPC2;LIPG;VDAC3;VDAC2;LPL;VDAC1;LIPA |
| Glucagon signaling pathway | 14/103 | 0.002038 | 0.0047678 | 2.632237778 | 16.3087875 | PDHA1;PGAM1;PDE3B;SLC2A1;PDHB;CPT1C;GCK;LDHB;GYS1;LDHA;PFKL;G6PC3;PGAM4;PCK2 |
| alpha-Linolenic acid metabolism | 25/6 | 0.0023079 | 0.0053501 | 5.266418258 | 31.97469062 | PLA2G12A;PLA2G2D;PLA2G4D;PLA2G1B;PLA2G4C;ACAA1 |
| Taurine and hypotaurine metabolism | 11/4 | 0.0024756 | 0.0056873 | 9.518929833 | 57.12552407 | GGT6;GAD1;ADO;CDO1 |
| Phagosome | 18/152 | 0.0025039 | 0.0057009 | 2.250420527 | 13.47978131 | ATP6V1A;ATP6V1G1;ATP6V0B;ATP6V0E1;ATP6AP1;TAP1;ATP6V0A2;ATP6V1H;ATP6V0A4;ATP6V1E1;ATP6V0D1;ATP6V0E2;ATP6V0D2;ATP6V1D;ATP6V0C;ATP6V1C1;ATP6V1F;ATP6V1B1 |
| Phosphonate and phosphinate metabolism | 6/3 | 0.0032105 | 0.007245 | 16.64695499 | 95.57539403 | PCYT1B;PCYT2;CHPT1 |
| Regulation of lipolysis in adipocytes | 9/55 | 0.0035712 | 0.0079881 | 3.266887852 | 18.40847728 | LIPE;PDE3B;ADCY3;PIK3R2;PIK3CB;PIK3R1;ADCY8;ADCY7;MGLL |
| Type II diabetes mellitus | 8/46 | 0.0040098 | 0.0088913 | 3.513624487 | 19.39174293 | PIK3R2;PIK3CB;PIK3R1;SLC2A4;HK2;GCK;HK1;CACNA1G |
| Fatty acid biosynthesis | 13/4 | 0.004896 | 0.0107628 | 7.402826855 | 39.37808515 | ACSL6;ACSL4;ACSL3;MCAT |
| Glutamatergic synapse | 14/114 | 0.005209 | 0.0113529 | 2.341319073 | 12.30918909 | SLC38A1;PLA2G4D;GLS2;SLC1A1;PLA2G4C;SLC1A3;ADCY3;SLC1A6;GRIN2C;ADCY8;ADCY7;GRIN2D;GRIN3A;SLC17A8 |
| Lysine degradation | 9/59 | 0.0057662 | 0.0124609 | 3.004897959 | 15.49247829 | GCDH;ECHS1;TMLHE;ALDH1B1;EHHADH;HADH;ALDH7A1;DLD;ACAT2 |
| Thiamine metabolism | 15/4 | 0.0085351 | 0.0182894 | 6.056215869 | 28.84922257 | THTPA;NFS1;AK1;AK5 |
| Thyroid hormone synthesis | 10/74 | 0.0088207 | 0.018744 | 2.608792185 | 12.34129178 | GPX8;GPX7;ATP1A3;ADCY3;ATP1A1;ATP1B1;ADCY8;SLC26A4;ADCY7;SLC5A5 |
| Sulfur metabolism | 9/3 | 0.0118561 | 0.024986 | 8.322153575 | 36.90803748 | MPST;TST;PAPSS1 |
| Protein digestion and absorption | 11/90 | 0.0128887 | 0.0269394 | 2.325007032 | 10.11705173 | ACE2;MME;SLC7A8;KCNE3;SLC1A1;SLC3A1;ATP1A3;SLC3A2;ATP1A1;ATP1B1;KCNN4 |
| Insulin resistance | 12/108 | 0.0195453 | 0.0405207 | 2.087188612 | 8.213132144 | GYS1;SLC27A1;G6PC3;GFPT1;SLC2A1;PIK3R2;PIK3R1;SLC2A4;PIK3CB;SLC27A5;SLC27A4;PCK2 |
| Pantothenate and CoA biosynthesis | 19/4 | 0.0202331 | 0.0416084 | 4.440282686 | 17.31903942 | PANK1;ENPP3;PPCDC;AASDHPPT |
| Phospholipase D signaling pathway | 15/148 | 0.0212607 | 0.0433718 | 1.884494913 | 7.25699196 | AGPAT5;PLA2G4D;SPHK2;PLA2G4C;ADCY3;PIK3R2;PIK3R1;PIK3CB;ADCY8;AGPAT1;ADCY7;AGPAT2;AGPAT4;PIP5K1B;PIP5K1C |
| Inflammatory mediator regulation of TRP channels | 11/100 | 0.0264033 | 0.0534353 | 2.06267166 | 7.496295806 | CYP2J2;PLA2G4D;PLA2G4C;TRPV2;ADCY3;PIK3R2;PIK3R1;TRPM8;PIK3CB;ADCY8;ADCY7 |
| Fluid shear stress and atherosclerosis | 14/139 | 0.0267045 | 0.0536193 | 1.870559715 | 6.77689474 | GSTM4;NQO1;GSTO2;GSTO1;MGST3;MGST2;GSTT2;PIK3R2;PIK3R1;PIK3CB;ASS1;GSTA3;GSTA2;GSTA1 |
| D-Glutamine and D-glutamate metabolism | 5/2 | 0.0287325 | 0.0572406 | 11.08818342 | 39.36000469 | GLUD2;GLS2 |
| Thyroid hormone signaling pathway | 12/116 | 0.032064 | 0.0633823 | 1.925814399 | 6.624842114 | SLCO1C1;SLC2A1;PLCE1;ATP1A3;PIK3R2;PIK3R1;PIK3CB;ATP1A1;ATP1B1;PLCD4;PFKP;SLC9A1 |
| Sphingolipid signaling pathway | 12/119 | 0.0379831 | 0.0745054 | 1.871520271 | 6.121018999 | SGPL1;ABCC1;ASAH1;SPTLC1;SPHK2;SMPD1;PIK3R2;DEGS1;PIK3R1;PIK3CB;SGMS2;SGPP1 |
| Mannose type O-glycan biosynthesis | 23/4 | 0.0386548 | 0.0752441 | 3.504742421 | 11.401222 | B3GALNT2;MGAT5B;CHST10;FUT4 |
| Adipocytokine signaling pathway | 8/69 | 0.0412056 | 0.0795553 | 2.186141146 | 6.971999204 | G6PC3;SLC2A1;ACSL6;ACSL4;SLC2A4;ACSL3;CPT1C;PCK2 |
| Vitamin B6 metabolism | 6/2 | 0.0414936 | 0.0795553 | 8.315696649 | 26.46235297 | PDXK;PNPO |
| Vitamin digestion and absorption | 24/4 | 0.0443697 | 0.0844349 | 3.329328622 | 10.37151805 | SLC46A1;ABCC1;FOLH1;LRAT |
| Circadian entrainment | 10/97 | 0.0485846 | 0.0917709 | 1.916763643 | 5.79715298 | RYR1;RYR2;KCNJ6;ADCY3;GRIN2C;ADCY8;ADCY7;CACNA1H;CACNA1G;GRIN2D |
| Cocaine addiction | 6/49 | 0.0578487 | 0.1084663 | 2.324058448 | 6.62339014 | GRIN3A;MAOB;MAOA;GRIN2C;SLC6A3;GRIN2D |
| Selenocompound metabolism | 17/3 | 0.0685127 | 0.1275236 | 3.565124196 | 9.557159031 | SEPHS1;SEPHS2;PAPSS1 |
| Adrenergic signaling in cardiomyocytes | 13/145 | 0.0692107 | 0.1278893 | 1.642758844 | 4.387151762 | RYR2;ATP1A3;ADCY3;ATP2B3;ATP2B2;ATP1A1;ATP2B1;ATP1B1;ADCY8;ADCY7;SLC9A1;CACNB3;SCN5A |
| Insulin signaling pathway | 12/137 | 0.0898391 | 0.1648128 | 1.600483986 | 3.856741422 | LIPE;GYS1;G6PC3;PDE3B;PIK3R2;PIK3R1;SLC2A4;PIK3CB;HK2;GCK;PCK2;HK1 |
| mTOR signaling pathway | 13/152 | 0.0921173 | 0.1677851 | 1.559447011 | 3.718801218 | ATP6V1A;ATP6V1G1;PIK3R2;SLC3A2;PIK3R1;PIK3CB;SLC7A5;ATP6V1H;ATP6V1E1;ATP6V1D;ATP6V1C1;ATP6V1F;ATP6V1B1 |
| Oxytocin signaling pathway | 13/153 | 0.0957312 | 0.1731309 | 1.548225417 | 3.632463205 | RYR1;KCNJ4;RYR2;KCNJ6;PLA2G4D;KCNJ12;PLA2G4C;KCNJ14;ADCY3;ADCY8;ADCY7;CACNB3;KCNJ2 |
| Cushing syndrome | 13/155 | 0.103216 | 0.1853526 | 1.526256381 | 3.466023889 | FH;HSD3B1;ADCY3;ADCY8;ADCY7;CACNA1H;CACNA1G;CYP17A1;PDE11A;CYP11A1;CYP21A2;ORAI1;PDE8B |
| VEGF signaling pathway | 6/59 | 0.1170222 | 0.2086759 | 1.884555018 | 4.043109226 | PLA2G4D;SPHK2;PLA2G4C;PIK3R2;PIK3CB;PIK3R1 |
| Vascular smooth muscle contraction | 11/132 | 0.1305016 | 0.2310966 | 1.514585859 | 3.084256704 | PLA2G12A;KCNU1;PLA2G2D;PLA2G4D;PLA2G1B;PLA2G4C;KCNMA1;KCNMB3;ADCY3;ADCY8;ADCY7 |
| Renin-angiotensin system | 23/3 | 0.13942 | 0.245187 | 2.494792586 | 4.915400176 | ACE2;MME;ATP6AP2 |
| Hepatocellular carcinoma | 13/168 | 0.1600558 | 0.2795495 | 1.397276879 | 2.560136483 | GSTM4;NQO1;GSTO2;GSTO1;MGST3;MGST2;GSTT2;PIK3R2;PIK3R1;PIK3CB;GSTA3;GSTA2;GSTA1 |
| Amphetamine addiction | 6/68 | 0.1883208 | 0.3266789 | 1.610219812 | 2.688436863 | GRIN3A;MAOB;MAOA;GRIN2C;SLC6A3;GRIN2D |
| Serotonergic synapse | 9/113 | 0.1927921 | 0.3321756 | 1.440516006 | 2.371294914 | CYP2J2;KCNJ6;MAOB;KCND2;PLA2G4D;MAOA;PLA2G4C;ALOX12B;CYP2C18 |
| Fc gamma R-mediated phagocytosis | 7/91 | 0.2588159 | 0.44294 | 1.386182462 | 1.873617288 | PLA2G4D;SPHK2;PIK3R2;PIP5K1B;PIK3CB;PIK3R1;PIP5K1C |
| Platelet activation | 9/124 | 0.2723724 | 0.4630331 | 1.301963659 | 1.693314426 | PLA2G4D;PLA2G4C;ORAI1;ADCY3;PIK3R2;PIK3R1;PIK3CB;ADCY8;ADCY7 |
| Endocrine and other factor-regulated calcium reabsorption | 4/48 | 0.289781 | 0.4893653 | 1.511403791 | 1.872069698 | ATP1A3;ATP1A1;ATP2B1;ATP1B1 |
| Human papillomavirus infection | 21/330 | 0.326665 | 0.5480235 | 1.130959119 | 1.265339769 | ATP6V1A;ATP6V1G1;ATP6V0B;ATP6V0E1;ATP6AP1;PIK3R2;PIK3R1;PIK3CB;OASL;ATP6V0A2;ATP6V1H;ATP6V0A4;ATP6V1E1;ATP6V0D1;ATP6V0E2;ATP6V0D2;ATP6V0C;ATP6V1D;ATP6V1C1;ATP6V1F;ATP6V1B1 |
| Amyotrophic lateral sclerosis (ALS) | 4/51 | 0.3288619 | 0.5481032 | 1.414705661 | 1.573318669 | CYCS;GRIN2C;GRIN2D;SOD1 |
| Progesterone-mediated oocyte maturation | 7/99 | 0.3314211 | 0.5487817 | 1.265105711 | 1.397139032 | PDE3B;ADCY3;PIK3R2;PIK3R1;PIK3CB;ADCY8;ADCY7 |
| Fc epsilon RI signaling pathway | 5/68 | 0.3433812 | 0.5649174 | 1.319312871 | 1.410232223 | PLA2G4D;PLA2G4C;PIK3R2;PIK3CB;PIK3R1 |
| Renal cell carcinoma | 5/69 | 0.3548079 | 0.5799744 | 1.298629531 | 1.345612443 | FH;SLC2A1;PIK3R2;PIK3CB;PIK3R1 |
| Longevity regulating pathway | 7/102 | 0.3594446 | 0.5838113 | 1.224959209 | 1.253372353 | ADCY3;PIK3R2;PIK3R1;PIK3CB;ADCY8;ADCY7;SOD1 |
| Human T-cell leukemia virus 1 infection | 14/219 | 0.3633327 | 0.5863913 | 1.135715838 | 1.149840092 | FDPS;SLC2A1;ADCY3;PIK3R2;PIK3R1;PIK3CB;ADCY8;ADCY7;VDAC3;VDAC2;VDAC1;SLC25A5;SLC25A4;SLC25A6 |
| Prolactin signaling pathway | 5/70 | 0.3662494 | 0.5873812 | 1.278582602 | 1.284260345 | PIK3R2;PIK3CB;PIK3R1;GCK;CYP17A1 |
| Apelin signaling pathway | 9/137 | 0.3759175 | 0.5991185 | 1.168921917 | 1.143656303 | RYR1;LIPE;RYR2;SPHK2;PDE3B;ADCY3;ADCY8;ADCY7;SLC9A1 |
| Necroptosis | 10/162 | 0.4399612 | 0.6968329 | 1.093297186 | 0.897672167 | GLUD2;PLA2G4D;SMPD1;PLA2G4C;VDAC3;VDAC2;VDAC1;SLC25A5;SLC25A4;SLC25A6 |
| RNA degradation | 5/79 | 0.4683266 | 0.7371808 | 1.12254164 | 0.851548083 | PFKL;ENO1;ENO2;PFKM;PFKP |
| Phototransduction | 28/2 | 0.4776796 | 0.7472902 | 1.277845611 | 0.944091613 | PDE6G;PDE6A |
| AGE-RAGE signaling pathway in diabetic complications | 6/100 | 0.5052699 | 0.7856331 | 1.060252307 | 0.723794463 | PLCE1;NOX4;PIK3R2;PIK3R1;PIK3CB;PLCD4 |
| Influenza A | 10/171 | 0.5087102 | 0.7861884 | 1.031684741 | 0.697291848 | FDPS;OAS1;OAS2;OAS3;CYCS;VDAC1;PIK3R2;PIK3R1;PIK3CB;SLC25A6 |
| Renin secretion | 4/69 | 0.5562584 | 0.854356 | 1.02196249 | 0.599403865 | PDE3B;KCNMA1;ORAI1;KCNJ2 |
| Cellular senescence | 9/160 | 0.5606055 | 0.854356 | 0.989657827 | 0.572752488 | VDAC3;VDAC2;VDAC1;PIK3R2;PIK3R1;PIK3CB;SLC25A5;SLC25A4;SLC25A6 |
| Parathyroid hormone synthesis, secretion and action | 6/106 | 0.5628698 | 0.854356 | 0.996318584 | 0.572591129 | CYP24A1;PDE4D;PDE4B;ADCY3;ADCY8;ADCY7 |
| Tuberculosis | 10/179 | 0.5676498 | 0.8565129 | 0.982427192 | 0.556300068 | ATP6V0B;ATP6AP1;SPHK2;ATP6V0A2;ATP6V1H;CYCS;ATP6V0A4;ATP6V0D1;ATP6V0D2;ATP6V0C |
| Dilated cardiomyopathy (DCM) | 5/91 | 0.5948529 | 0.8922793 | 0.965291058 | 0.501411904 | RYR2;CACNB3;ADCY3;ADCY8;ADCY7 |
| GnRH signaling pathway | 5/93 | 0.6142688 | 0.9160149 | 0.94325215 | 0.4596681 | PLA2G4D;PLA2G4C;ADCY3;ADCY8;ADCY7 |
| African trypanosomiasis | 2/37 | 0.6293215 | 0.9276127 | 0.948803225 | 0.439403086 | IDO2;IDO1 |
| Primary immunodeficiency | 2/37 | 0.6293215 | 0.9276127 | 0.948803225 | 0.439403086 | TAP1;ORAI1 |
| Endometrial cancer | 3/58 | 0.64744 | 0.9488344 | 0.905512316 | 0.393652661 | PIK3R2;PIK3CB;PIK3R1 |
| Estrogen signaling pathway | 7/137 | 0.666736 | 0.9709039 | 0.893493221 | 0.362187403 | KCNJ6;ADCY3;PIK3R2;PIK3R1;PIK3CB;ADCY8;ADCY7 |
| Long-term depression | 3/60 | 0.6701141 | 0.9709039 | 0.873647048 | 0.349727319 | RYR1;PLA2G4D;PLA2G4C |
| Measles | 7/138 | 0.6741411 | 0.9712203 | 0.886625332 | 0.349610355 | OAS1;OAS2;OAS3;CYCS;PIK3R2;PIK3R1;PIK3CB |
| Melanogenesis | 5/101 | 0.6862048 | 0.9830462 | 0.864279399 | 0.325469648 | TYRP1;ADCY3;TYR;ADCY8;ADCY7 |
| Taste transduction | 4/83 | 0.7009748 | 0.9985954 | 0.840229011 | 0.298519377 | SCNN1G;GABRA5;GABRA3;ADCY8 |
| Epstein-Barr virus infection | 10/201 | 0.7112601 | 0.9999955 | 0.868245097 | 0.295825955 | ENTPD1;OAS1;OAS2;OAS3;ENTPD8;TAP1;CYCS;PIK3R2;PIK3R1;PIK3CB |
| Vasopressin-regulated water reabsorption | 2/44 | 0.721832 | 0.9999955 | 0.790375409 | 0.25763307 | ADCY3;AQP2 |
| Other types of O-glycan biosynthesis | 22/1 | 0.7239533 | 0.9999955 | 0.790560101 | 0.255373382 | POFUT1 |
| Colorectal cancer | 4/86 | 0.7271016 | 0.9999955 | 0.809359648 | 0.257934115 | CYCS;PIK3R2;PIK3CB;PIK3R1 |
| Acute myeloid leukemia | 3/66 | 0.7316403 | 0.9999955 | 0.790190392 | 0.246907888 | PIK3R2;PIK3CB;PIK3R1 |
| Non-small cell lung cancer | 3/66 | 0.7316403 | 0.9999955 | 0.790190392 | 0.246907888 | PIK3R2;PIK3CB;PIK3R1 |
| Neuroactive ligand-receptor interaction | 17/338 | 0.7321053 | 0.9999955 | 0.877594314 | 0.273661046 | CHRNA1;CHRNA5;GABRA5;CHRNA4;GRID1;CHRNA7;GABRA3;CHRNA6;GRIN2C;GABRG1;GRIN2D;GLRA2;P2RX5;GRIN3A;P2RX4;CHRNG;GABRD |
| Protein export | 23/1 | 0.7396499 | 0.9999955 | 0.754585503 | 0.227566604 | OXA1L |
| Long-term potentiation | 3/67 | 0.7409635 | 0.9999955 | 0.777802295 | 0.233188195 | ADCY8;GRIN2C;GRIN2D |
| Relaxin signaling pathway | 6/130 | 0.7544149 | 0.9999955 | 0.802455039 | 0.226142087 | ADCY3;PIK3R2;PIK3R1;PIK3CB;ADCY8;ADCY7 |
| FoxO signaling pathway | 6/132 | 0.7671743 | 0.9999955 | 0.789633375 | 0.209285463 | G6PC3;PIK3R2;PIK3R1;SLC2A4;PIK3CB;PCK2 |
| B cell receptor signaling pathway | 3/71 | 0.7756959 | 0.9999955 | 0.731893463 | 0.185897109 | PIK3R2;PIK3CB;PIK3R1 |
| Small cell lung cancer | 4/93 | 0.7813469 | 0.9999955 | 0.745424227 | 0.183922992 | CYCS;PIK3R2;PIK3CB;PIK3R1 |
| Maturity onset diabetes of the young | 26/1 | 0.7815886 | 0.9999955 | 0.663929515 | 0.163609998 | GCK |
| Melanoma | 3/72 | 0.7837566 | 0.9999955 | 0.721247937 | 0.175736899 | PIK3R2;PIK3CB;PIK3R1 |
| p53 signaling pathway | 3/72 | 0.7837566 | 0.9999955 | 0.721247937 | 0.175736899 | RRM2;RRM2B;CYCS |
| Hepatitis C | 7/155 | 0.7837666 | 0.9999955 | 0.784071051 | 0.191034241 | OAS1;OAS2;OAS3;CYCS;PIK3R2;PIK3R1;PIK3CB |
| Bacterial invasion of epithelial cells | 3/74 | 0.7991592 | 0.9999955 | 0.700856507 | 0.157128606 | PIK3R2;PIK3CB;PIK3R1 |
| Amoebiasis | 4/96 | 0.801818 | 0.9999955 | 0.72100169 | 0.159250254 | ARG2;PIK3R2;PIK3CB;PIK3R1 |
| Glioma | 3/75 | 0.8065095 | 0.9999955 | 0.691085613 | 0.148610753 | PIK3R2;PIK3CB;PIK3R1 |
| Pancreatic cancer | 3/75 | 0.8065095 | 0.9999955 | 0.691085613 | 0.148610753 | PIK3R2;PIK3CB;PIK3R1 |
| Chronic myeloid leukemia | 3/76 | 0.8136318 | 0.9999955 | 0.681582415 | 0.140574545 | PIK3R2;PIK3CB;PIK3R1 |
| ErbB signaling pathway | 3/85 | 0.8682439 | 0.9999955 | 0.606483973 | 0.085685645 | PIK3R2;PIK3CB;PIK3R1 |
| Prion diseases | 1/35 | 0.8710688 | 0.9999955 | 0.487950246 | 0.067353883 | SOD1 |
| Dopaminergic synapse | 5/131 | 0.8718326 | 0.9999955 | 0.657446002 | 0.090173843 | KCNJ6;MAOB;MAOA;COMT;SLC6A3 |
| Gap junction | 3/88 | 0.8830261 | 0.9999955 | 0.584985203 | 0.072772466 | ADCY3;ADCY8;ADCY7 |
| NOD-like receptor signaling pathway | 7/178 | 0.8848969 | 0.9999955 | 0.67777726 | 0.082881381 | OAS1;OAS2;OAS3;VDAC3;TRPV2;VDAC2;VDAC1 |
| Hepatitis B | 6/163 | 0.9066418 | 0.9999955 | 0.632670086 | 0.062006636 | ATP6AP1;VDAC3;CYCS;PIK3R2;PIK3R1;PIK3CB |
| Protein processing in endoplasmic reticulum | 6/165 | 0.9124275 | 0.9999955 | 0.624645183 | 0.057246665 | GANAB;DAD1;EDEM2;MAN1C1;MAN1B1;DDOST |
| Ras signaling pathway | 9/232 | 0.9156749 | 0.9999955 | 0.667548673 | 0.058806987 | PLA2G12A;PLA2G2D;PLA2G4D;PLA2G1B;PLA2G4C;PLCE1;PIK3R2;PIK3R1;PIK3CB |
| Prostate cancer | 3/97 | 0.9188824 | 0.9999955 | 0.528722465 | 0.044728389 | PIK3R2;PIK3R1;PIK3CB |
| Type I diabetes mellitus | 1/43 | 0.9193158 | 0.9999955 | 0.394839522 | 0.033216107 | GAD1 |
| Arrhythmogenic right ventricular cardiomyopathy (ARVC) | 2/72 | 0.9211739 | 0.9999955 | 0.473519778 | 0.038879023 | RYR2;CACNB3 |
| T cell receptor signaling pathway | 3/101 | 0.9313359 | 0.9999955 | 0.507033882 | 0.036067976 | PIK3R2;PIK3R1;PIK3CB |
| Chagas disease (American trypanosomiasis) | 3/103 | 0.9368797 | 0.9999955 | 0.496840247 | 0.032394158 | PIK3R2;PIK3R1;PIK3CB |
| C-type lectin receptor signaling pathway | 3/104 | 0.9394938 | 0.9999955 | 0.491894821 | 0.030701139 | PIK3R2;PIK3R1;PIK3CB |
| Toll-like receptor signaling pathway | 3/104 | 0.9394938 | 0.9999955 | 0.491894821 | 0.030701139 | PIK3R2;PIK3R1;PIK3CB |
| Human cytomegalovirus infection | 8/225 | 0.9455397 | 0.9999955 | 0.609438834 | 0.034128202 | ADCY3;TAP1;CYCS;PIK3R2;PIK3R1;PIK3CB;ADCY8;ADCY7 |
| Alcoholism | 6/180 | 0.9467002 | 0.9999955 | 0.570338724 | 0.031239031 | GRIN3A;MAOB;MAOA;GRIN2C;SLC6A3;GRIN2D |
| ECM-receptor interaction | 2/82 | 0.951205 | 0.9999955 | 0.414109347 | 0.020716122 | SV2B;SV2A |
| Rap1 signaling pathway | 7/206 | 0.9515611 | 0.9999955 | 0.581539228 | 0.028874225 | PLCE1;ADCY3;PIK3R2;PIK3R1;PIK3CB;ADCY8;ADCY7 |
| TNF signaling pathway | 3/110 | 0.9531835 | 0.9999955 | 0.464163457 | 0.02225563 | PIK3R2;PIK3R1;PIK3CB |
| Leukocyte transendothelial migration | 3/112 | 0.9570629 | 0.9999955 | 0.455598112 | 0.019994468 | PIK3R2;PIK3R1;PIK3CB |
| Hypertrophic cardiomyopathy (HCM) | 2/85 | 0.9578355 | 0.9999955 | 0.399077793 | 0.017191969 | RYR2;CACNB3 |
| Legionellosis | 1/55 | 0.9600716 | 0.9999955 | 0.306901615 | 0.012505463 | CYCS |
| Chemokine signaling pathway | 6/190 | 0.9622962 | 0.9999955 | 0.539053482 | 0.020717412 | ADCY3;PIK3R2;PIK3R1;PIK3CB;ADCY8;ADCY7 |
| Regulation of actin cytoskeleton | 7/214 | 0.9628076 | 0.9999955 | 0.558824662 | 0.021180398 | PIK3R2;PIP5K1B;PIP4K2C;PIK3R1;PIP5K1C;PIK3CB;SLC9A1 |
| Apoptosis | 4/143 | 0.965274 | 0.9999955 | 0.476014948 | 0.016823912 | CYCS;PIK3R2;PIK3R1;PIK3CB |
| Neurotrophin signaling pathway | 3/119 | 0.9683939 | 0.9999955 | 0.427945339 | 0.013744035 | PIK3R2;PIK3R1;PIK3CB |
| Viral myocarditis | 1/59 | 0.9684203 | 0.9999955 | 0.285675224 | 0.00916705 | CYCS |
| Proteoglycans in cancer | 6/201 | 0.9745596 | 0.9999955 | 0.508345813 | 0.013099869 | PLCE1;PIK3R2;PIK3R1;PIK3CB;NUDT16L1;SLC9A1 |
| Viral carcinogenesis | 6/201 | 0.9745596 | 0.9999955 | 0.508345813 | 0.013099869 | VDAC3;PIK3R2;PIK3R1;PIK3CB;ATP6V0D1;ATP6V0D2 |
| Oocyte meiosis | 3/125 | 0.9757958 | 0.9999955 | 0.406768625 | 0.009966615 | ADCY3;ADCY8;ADCY7 |
| Osteoclast differentiation | 3/127 | 0.9778734 | 0.9999955 | 0.400165134 | 0.008953709 | PIK3R2;PIK3R1;PIK3CB |
| Autophagy | 3/128 | 0.9788474 | 0.9999955 | 0.39694263 | 0.008486426 | PIK3R2;PIK3R1;PIK3CB |
| Axon guidance | 5/181 | 0.9788501 | 0.9999955 | 0.469415642 | 0.010034578 | TRPC6;TRPC3;PIK3R2;PIK3R1;PIK3CB |
| Natural killer cell mediated cytotoxicity | 3/131 | 0.9815301 | 0.9999955 | 0.387577229 | 0.007225431 | PIK3R2;PIK3R1;PIK3CB |
| Signaling pathways regulating pluripotency of stem cells | 3/139 | 0.9871859 | 0.9999955 | 0.364622813 | 0.004702511 | PIK3R2;PIK3R1;PIK3CB |
| Antigen processing and presentation | 1/77 | 0.9890164 | 0.9999955 | 0.217806631 | 0.002405531 | TAP1 |
| Pathways in cancer | 19/530 | 0.9902201 | 0.9999955 | 0.610923164 | 0.006004205 | GSTM4;NQO1;FH;GSTO2;GSTO1;MGST3;SLC2A1;ADCY3;GSTT2;MGST2;PIK3R2;PIK3CB;PIK3R1;ADCY8;ADCY7;GSTA3;GSTA2;GSTA1;CYCS |
| Breast cancer | 3/147 | 0.9911567 | 0.9999955 | 0.344218888 | 0.003057571 | PIK3R2;PIK3R1;PIK3CB |
| Gastric cancer | 3/149 | 0.991946 | 0.9999955 | 0.339467289 | 0.002745144 | PIK3R2;PIK3R1;PIK3CB |
| Human immunodeficiency virus 1 infection | 5/212 | 0.9939427 | 0.9999955 | 0.398454619 | 0.002420882 | TAP1;CYCS;PIK3R2;PIK3R1;PIK3CB |
| Kaposi sarcoma-associated herpesvirus infection | 4/186 | 0.9943909 | 0.9999955 | 0.362715023 | 0.002040242 | CYCS;PIK3R2;PIK3R1;PIK3CB |
| mRNA surveillance pathway | 1/91 | 0.9951717 | 0.9999955 | 0.183788546 | 8.90E-04 | NUDT21 |
| JAK-STAT signaling pathway | 3/162 | 0.9956435 | 0.9999955 | 0.311495612 | 0.001359984 | PIK3R2;PIK3R1;PIK3CB |
| Hematopoietic cell lineage | 1/97 | 0.9966054 | 0.9999955 | 0.172246696 | 5.86E-04 | MME |
| Focal adhesion | 4/199 | 0.9968609 | 0.9999955 | 0.338298451 | 0.001063633 | PIK3R2;PIK3R1;PIP5K1C;PIK3CB |
| Toxoplasmosis | 1/113 | 0.9986732 | 0.9999955 | 0.14751416 | 1.96E-04 | CYCS |
| Transcriptional misregulation in cancer | 2/186 | 0.9997759 | 0.9999955 | 0.179050686 | 4.01E-05 | FUT8;HPGD |
| MAPK signaling pathway | 5/295 | 0.9998421 | 0.9999955 | 0.283148876 | 4.47E-05 | CACNB3;PLA2G4D;PLA2G4C;CACNA1H;CACNA1G |
| Endocytosis | 3/244 | 0.999923 | 0.9999955 | 0.204608629 | 1.58E-05 | PIP5K1B;PIP5K1C;FOLR2 |
| PI3K-Akt signaling pathway | 6/354 | 0.9999537 | 0.9999955 | 0.282514495 | 1.31E-05 | GYS1;G6PC3;PIK3R2;PIK3R1;PIK3CB;PCK2 |
| MicroRNAs in cancer | 4/299 | 0.9999695 | 0.9999955 | 0.22242319 | 6.79E-06 | ABCC1;CYP24A1;GLS2;PIK3R2 |
| Herpes simplex virus 1 infection | 8/492 | 0.9999941 | 0.9999955 | 0.269327706 | 1.60E-06 | OAS1;OAS2;OAS3;TAP1;CYCS;PIK3R2;PIK3CB;PIK3R1 |
| Olfactory transduction | 2/444 | 0.9999955 | 0.9999955 | 0.073507466 | 3.31E-07 | PDE2A;ADCY3 |
