## Supplementary material for "Glutamine antagonism suppresses tumor growth in adrenocortical carcinoma through inhibition of de novo nucleotide biosynthesis": Suppl. Table 2A

**Suppl. Table 2A mouse down  Down-regulated pathways BPCre ACC tumors versus Control adrenal**

| Term | Overlap | P-value | Adjusted P-va | Odds Ratio | Combined Sc | Genes |
|---|---|---|---|---|---|---|
| Retrograde endocannabinoid signaling | 30/150 | 6.25E-18 | 1.49E-15 | 9.4803922 | 375.55991 | GRIA1;GRIA2;GABRB1;NDUFB8;DAGLA;NDUFB6;CACNA1B;ADCY4;NDUFA4L2;ITPR2;ABHD6;ADCY2;ADCY8;PTGS2;ADCY6;ADCY5;NDUFV3;GRIA3;GRIA4;KCNJ3;GABRQ;KCNJ5;KCNJ6;GABRA4;GABRA3;NDUFA1;GABRG3;GABRG2;NDUFS8;DAGLB |
| Morphine addiction | 21/92 | 3.76E-14 | 4.19E-12 | 11.049662 | 341.57812 | GABRQ;KCNJ5;GABRB1;KCNJ6;PDE1C;PDE1B;PDE1A;GABRA4;GABRA3;ADCY4;CACNA1B;ADCY2;ADCY8;GABRG3;GABRG2;ADCY6;ADCY5;PDE10A;PDE3A;PDE7B;KCNJ3 |
| Nicotine addiction | 15/40 | 5.26E-14 | 4.19E-12 | 22.211429 | 679.12366 | GRIA1;CHRNB2;GABRQ;GRIA2;GABRB1;CHRNA7;GABRA4;GABRA3;CACNA1B;GABRG3;GABRG2;GRIN1;GRIN3A;GRIA3;GRIA4 |
| ABC transporters | 15/48 | 1.17E-12 | 7.00E-11 | 16.819913 | 462.07793 | ABCA1;ABCD2;ABCD3;ABCA5;ABCA8B;ABCB5;ABCA4;TAP2;ABCA9;TAP1;ABCC9;ABCB9;ABCA13;ABCB1B;ABCG4 |
| Purine metabolism | 23/136 | 1.95E-12 | 9.32E-11 | 7.6167988 | 205.37054 | GUCY1A1;GUCY1B1;ENTPD2;ENTPD3;PDE1C;PDE1B;ENTPD5;PDE1A;NPR2;ADCY4;AK3;NME3;ADCY2;AK5;ADCY8;ADCY6;ADCY5;PDE10A;NT5C1A;PDE3A;ENPP4;PDE6A;PDE7B |
| Glutamatergic synapse | 21/114 | 3.25E-12 | 1.30E-10 | 8.4261918 | 222.88859 | GRIA1;GRIA2;SLC38A1;PLA2G4D;TRPC1;ADCY4;SLC1A3;ITPR2;GRIK4;ADCY2;GRIK2;PLD1;ADCY8;ADCY6;ADCY5;GRIN1;GRIN3A;GLUL;GRIA3;GRIA4;KCNJ3 |
| Glycerolipid metabolism | 16/61 | 4.23E-12 | 1.44E-10 | 13.173876 | 345.01741 | DGKG;DGKE;DGAT2;DGKB;MOGAT1;DGKA;LCLAT1;AGPAT2;ALDH3A2;GPAM;LIPG;PNPLA3;PLPP3;LPIN1;PLPP1;DGKH |
| Glycosphingolipid biosynthesis | 14/45 | 7.16E-12 | 2.13E-10 | 16.681344 | 428.08099 | B3GALNT1;ST8SIA1;SLC33A1;B3GALT5;GGTA1;GBGT1;FUT9;B3GALT2;B3GNT5;B3GALT1;GCNT2;B4GALNT1;ST6GALNAC3;ST6GALNAC5 |
| Peroxisome | 18/84 | 8.01E-12 | 2.13E-10 | 10.132706 | 258.88935 | GSTK1;ABCD2;ABCD3;EPHX2;ACSL6;ACSL4;ACSL3;SOD2;DDO;SLC25A17;ACOX1;EHHADH;FAR1;HAO1;SLC27A2;HMGCLL1;HAO2;NUDT12 |
| Cholinergic synapse | 20/113 | 2.31E-11 | 5.51E-10 | 8.009512 | 196.17927 | CHRNB2;ACHE;CHRNB4;CHRNA3;KCNJ6;CHRNA7;CACNA1B;ADCY4;ITPR2;PIK3R3;ADCY2;PIK3R1;PIK3CB;ADCY8;ADCY6;ADCY5;KCNQ2;KCNQ3;KCNQ4;KCNJ3 |
| cAMP signaling pathway | 26/211 | 1.17E-10 | 2.43E-09 | 5.2702703 | 120.53728 | GRIA1;HCN4;GRIA2;ADCY4;ATP1A4;PIK3R3;ADCY2;ATP1A2;PIK3R1;PIK3CB;ADCY8;PLD1;ADCY6;ADCY5;PLCE1;CNGA3;GRIA3;GRIA4;ATP2B4;ATP2B2;ATP2B1;GRIN1;GRIN3A;PDE10A;ACOX1;PDE3A |
| Phosphatidylinositol signaling system | 18/98 | 1.22E-10 | 2.43E-09 | 8.3534483 | 190.6727 | DGKG;IPPK;DGKE;DGKB;DGKA;PTEN;ITPR2;PIK3R3;PIK3C2G;PIK3R1;PIK3CB;PPIP5K1;INPP5F;SYNJ1;PLCE1;PI4K2A;IP6K3;DGKH |
| Salivary secretion | 16/78 | 2.35E-10 | 4.04E-09 | 9.553312 | 211.82356 | SLC12A2;GUCY1A1;GUCY1B1;ATP1A4;ADCY4;ATP2B4;ITPR2;ATP1A2;ADCY2;ATP2B2;ATP2B1;ADCY8;ADCY6;ADCY5;BST1;NOS1 |
| Aldosterone synthesis and secretion | 18/102 | 2.43E-10 | 4.04E-09 | 7.954023 | 176.07058 | KCNJ5;DAGLA;HSD3B1;ATP1A4;ADCY4;ATP2B4;ITPR2;ATP1A2;ADCY2;ATP2B2;ATP2B1;ADCY8;ADCY6;ADCY5;STAR;CYP11A1;CYP11B1;DAGLB |
| GABAergic synapse | 17/90 | 2.53E-10 | 4.04E-09 | 8.6324681 | 190.74719 | GABRQ;GABRB1;SLC38A1;KCNJ6;GABRA4;GAD1;GABRA3;ADCY4;CACNA1B;ADCY2;ADCY8;GABRG3;GPHN;GABRG2;ADCY6;ADCY5;GLUL |
| Sphingolipid metabolism | 13/48 | 2.85E-10 | 4.26E-09 | 13.690702 | 300.8883 | CERS4;ASAH2;CERS6;SGMS1;SGPP2;SGPP1;SMPD3;SPTLC1;SMPD1;DEGS1;PLPP3;B4GALT6;PLPP1 |
| Valine, leucine and isoleucine degradation | 13/56 | 2.29E-09 | 3.22E-08 | 11.139005 | 221.59011 | ECHS1;HMGCS1;AGXT2;ACADSB;AACS;ALDH3A2;MCEE;ALDH6A1;PCCA;EHHADH;DBT;PCCB;HMGCLL1 |
| Ovarian steroidogenesis | 13/57 | 2.90E-09 | 3.85E-08 | 10.885285 | 213.99375 | PLA2G4D;HSD3B1;ADCY4;ADCY2;HSD17B7;ADCY8;PTGS2;ADCY6;ADCY5;CYP17A1;STAR;CYP11A1;CYP1B1 |
| Propanoate metabolism | 10/31 | 5.05E-09 | 6.35E-08 | 17.465409 | 333.67267 | ACSS3;MCEE;ALDH6A1;ECHS1;PCCA;EHHADH;DBT;PCCB;ECHDC1;ACACA |
| Glycerophospholipid metabolism | 16/97 | 6.74E-09 | 8.06E-08 | 7.3052493 | 137.44623 | DGKG;ACHE;DGKE;LPGAT1;PLA2G4D;DGKB;DGKA;PLD1;LCLAT1;AGPAT2;GPAM;CEPT1;PLPP3;LPIN1;PLPP1;DGKH |
| Circadian entrainment | 16/99 | 9.15E-09 | 1.00E-07 | 7.1284834 | 131.9479 | GRIA1;GRIA2;KCNJ5;GUCY1A1;KCNJ6;GUCY1B1;ADCY4;ADCY2;ADCY8;ADCY6;ADCY5;GRIN1;NOS1;GRIA3;GRIA4;KCNJ3 |
| Calcium signaling pathway | 22/189 | 9.24E-09 | 1.00E-07 | 4.9065498 | 90.772271 | PDE1C;PDE1B;CHRNA7;PDE1A;CACNA1B;ATP2A3;ADCY4;ATP2B4;ITPR2;ADCY2;ATP2B2;ATP2B1;ADCY8;CACNA1E;SLC8A2;GRIN1;P2RX6;P2RX5;ORAI2;PLCE1;NOS1;SLC25A5 |
| Taste transduction | 15/88 | 1.26E-08 | 1.31E-07 | 7.5878669 | 138.02516 | KCNK5;HCN4;ENTPD2;PDE1C;PDE1B;PDE1A;GABRA4;GABRA3;ADCY4;HTR3B;ADCY8;ADCY6;SCN9A;ASIC2;SCN3A |
| Steroid biosynthesis | 8/19 | 1.56E-08 | 1.55E-07 | 26.587833 | 477.94808 | EBP;SC5D;DHCR24;MSMO1;DHCR7;HSD17B7;LSS;CYP51 |
| Pancreatic secretion | 16/105 | 2.18E-08 | 2.09E-07 | 6.645853 | 117.23649 | SLC12A2;TRPC1;ATP2A3;ATP1A4;ADCY4;ATP2B4;ITPR2;ATP1A2;ADCY2;ATP2B2;ATP2B1;ADCY8;SLC4A4;ADCY6;ADCY5;BST1 |
| Oxytocin signaling pathway | 19/154 | 3.69E-08 | 3.39E-07 | 5.2203739 | 89.345101 | KCNJ5;GUCY1A1;KCNJ6;GUCY1B1;PLA2G4D;NPR2;ADCY4;CACNA2D2;ITPR2;ADCY2;ADCY8;PTGS2;ADCY6;ADCY5;TRPM2;CACNG2;CACNG3;KCNJ3;CACNG5 |
| cGMP-PKG signaling pathway | 20/172 | 4.44E-08 | 3.93E-07 | 4.8856275 | 82.712602 | GUCY1A1;GUCY1B1;NPR2;ATP2A3;ATP1A4;ADCY4;ATP2B4;ITPR2;ATP1A2;ADCY2;ATP2B2;ATP2B1;ADCY8;ADCY6;SLC8A2;ADCY5;KCNU1;KCNMB2;PDE3A;SLC25A5 |
| Bile secretion | 13/72 | 5.80E-08 | 4.95E-07 | 8.1115685 | 135.16308 | ATP1A4;ADCY4;ATP1A2;ADCY2;HMGCR;ADCY8;SLC4A4;ADCY6;AQP1;ADCY5;SLC9A3;SLC22A8;ABCB1B |
| Choline metabolism in cancer | 15/99 | 6.50E-08 | 5.36E-07 | 6.5904762 | 109.06066 | DGKG;SLC22A4;DGKE;SLC44A3;SLC22A5;PLA2G4D;DGKB;DGKA;PIK3R3;PIK3R1;PIK3CB;PLD1;PLPP3;PLPP1;DGKH |
| Glyoxylate and dicarboxylate metabolism | 9/31 | 8.47E-08 | 6.75E-07 | 14.975347 | 243.86657 | MCEE;MDH1;PCCA;GLDC;PCCB;HAO1;ACO1;GLUL;HAO2 |
| Steroid hormone biosynthesis | 14/89 | 1.10E-07 | 8.49E-07 | 6.8793409 | 110.21579 | UGT1A1;SRD5A2;AKR1D1;HSD3B1;CYP7B1;HSD17B7;CYP17A1;HSD11B1;SULT1E1;CYP11A1;CYP11B2;CYP11B1;CYP1B1;CYP2E1 |
| Phospholipase D signaling pathway | 18/149 | 1.17E-07 | 8.72E-07 | 5.0879179 | 81.22106 | DGKG;DGKE;PLA2G4D;DGKB;DGKA;ADCY4;PIK3R3;ADCY2;PIK3R1;PIK3CB;PLD1;ADCY8;AGPAT2;ADCY6;ADCY5;PLPP3;PLPP1;DGKH |
| Regulation of lipolysis in adipocytes | 11/56 | 2.49E-07 | 1.80E-06 | 8.9714346 | 136.42131 | ADCY4;PIK3R3;ADCY2;PIK3CB;ABHD5;PIK3R1;ADCY8;PTGS2;ADCY6;ADCY5;PTGS1 |
| Cortisol synthesis and secretion | 12/69 | 2.89E-07 | 2.03E-06 | 7.7364434 | 116.49608 | STAR;CYP11A1;CYP11B2;HSD3B1;ADCY4;ITPR2;KCNA4;ADCY2;ADCY8;ADCY6;ADCY5;CYP17A1 |
| Butanoate metabolism | 8/27 | 3.79E-07 | 2.59E-06 | 15.386624 | 227.5193 | ECHS1;BDH2;ACSM3;HMGCS1;GAD1;EHHADH;HMGCLL1;AACS |
| Drug metabolism | 15/114 | 4.34E-07 | 2.88E-06 | 5.5875902 | 81.858327 | GSTK1;UGT1A1;MGST1;MGST2;NME3;FMO1;FMO2;FMO3;ADH7;FMO4;CES1D;ALDH3B1;DPYD;CMPK1;CYP2E1 |
| Serotonergic synapse | 16/132 | 5.61E-07 | 3.63E-06 | 5.0918663 | 73.288249 | KCNJ5;GABRB1;DDC;KCNJ6;PLA2G4D;TRPC1;CACNA1B;ITPR2;HTR3B;PTGS2;SLC6A4;ADCY5;PTGS1;SLC18A1;SLC18A2;KCNJ3 |
| Fatty acid degradation | 10/50 | 7.37E-07 | 4.63E-06 | 9.1603774 | 129.35185 | ALDH3A2;CPT1A;ECHS1;ACOX1;EHHADH;ACSL6;ACSL4;ACSL3;ADH7;ACADSB |
| Tyrosine metabolism | 9/40 | 9.25E-07 | 5.67E-06 | 10.622745 | 147.58884 | PNMT;AOC3;GSTZ1;DDC;GOT1;ALDH3B1;DBH;ADH7;FAH |
| Proximal tubule bicarbonate reclamation | 7/22 | 1.21E-06 | 7.21E-06 | 17.025016 | 232.01622 | SLC9A3;MDH1;ATP1A4;ATP1A2;PCK1;SLC4A4;AQP1 |
| Metabolism of xenobiotics by cytochrome P450 | 11/66 | 1.41E-06 | 8.20E-06 | 7.3364839 | 98.852139 | HSD11B1;GSTK1;UGT1A1;ALDH3B1;CYP2S1;DHDH;MGST1;CYP1B1;MGST2;CYP2E1;ADH7 |
| beta-Alanine metabolism | 8/32 | 1.59E-06 | 9.06E-06 | 12.177945 | 162.57838 | AOC3;ALDH3A2;ALDH6A1;ECHS1;ALDH3B1;GAD1;EHHADH;DPYD |
| Adrenergic signaling in cardiomyocytes | 16/148 | 2.62E-06 | 1.45E-05 | 4.4709692 | 57.466794 | ATP1A4;ADCY4;ATP2B4;CACNA2D2;ATP1A2;ADCY2;ATP2B2;ATP2B1;ADCY8;ADCY6;ADCY5;SCN7A;CACNG2;SCN4B;CACNG3;CACNG5 |
| PPAR signaling pathway | 12/85 | 2.91E-06 | 1.58E-05 | 6.0358032 | 76.937823 | CYP27A1;FADS2;CPT1A;ACOX1;EHHADH;ACSL6;UCP1;ACSL4;SCD1;ACSL3;PCK1;SLC27A2 |
| Thyroid hormone synthesis | 11/73 | 3.93E-06 | 2.09E-05 | 6.5058235 | 80.975352 | IYD;GPX3;ATP1A4;ADCY4;ITPR2;ATP1A2;ADCY2;ADCY8;SLC26A4;ADCY6;ADCY5 |
| Thermogenesis | 20/231 | 5.01E-06 | 2.61E-05 | 3.5087495 | 42.817668 | COX8A;CPT1A;NDUFB8;COX16;NDUFB6;COX15;ACSL6;ADCY4;NDUFA4L2;UCP1;NDUFA1;ACSL4;ADCY2;ACSL3;UQCR10;ADCY8;ADCY6;ADCY5;NDUFS8;NDUFV3 |
| Alanine, aspartate and glutamate metabolism | 8/37 | 5.19E-06 | 2.64E-05 | 10.075707 | 122.60827 | FOLH1;GOT1;AGXT2;GAD1;GPT2;GLUL;ASPA;DDO |
| Renin secretion | 11/76 | 5.88E-06 | 2.93E-05 | 6.204595 | 74.726207 | GUCY1A1;GUCY1B1;ACE;PDE1C;PDE1B;PDE1A;PDE3A;ITPR2;ADCY6;AQP1;ADCY5 |
| Sphingolipid signaling pathway | 14/124 | 6.62E-06 | 3.23E-05 | 4.681991 | 55.835084 | CERS4;CERS6;ASAH2;SGMS1;PTEN;PIK3R3;PIK3R1;PIK3CB;PLD1;SGPP2;SGPP1;SPTLC1;SMPD1;DEGS1 |
| Chemical carcinogenesis | 12/94 | 8.48E-06 | 4.05E-05 | 5.3708426 | 62.719637 | HSD11B1;SULT1A1;GSTK1;UGT1A1;CHRNA7;ALDH3B1;MGST1;CYP1B1;MGST2;CYP2E1;PTGS2;ADH7 |
| Amphetamine addiction | 10/68 | 1.35E-05 | 6.33E-05 | 6.3116461 | 70.772745 | GRIA1;GRIA2;DDC;GRIN3A;SLC18A1;SLC18A2;GRIA3;ADCY5;GRIA4;GRIN1 |
| Biosynthesis of unsaturated fatty acids | 7/32 | 1.88E-05 | 8.66E-05 | 10.209756 | 111.08096 | FADS2;ELOVL4;ELOVL5;ACOX1;SCD1;FADS1;ACOT4 |
| Fatty acid biosynthesis | 5/14 | 2.30E-05 | 1.04E-04 | 20.198339 | 215.68305 | FASN;ACSL6;ACSL4;ACSL3;ACACA |
| Platelet activation | 13/125 | 3.41E-05 | 1.51E-04 | 4.2613852 | 43.839408 | GUCY1A1;GUCY1B1;PLA2G4D;ADCY4;ITPR2;PIK3R3;ADCY2;PIK3R1;PIK3CB;ADCY8;ADCY6;ADCY5;PTGS1 |
| Long-term depression | 9/61 | 3.55E-05 | 1.54E-04 | 6.3259452 | 64.823543 | GRIA1;GRIA2;GUCY1A1;GRID2;GUCY1B1;PLA2G4D;ITPR2;NOS1;GRIA3 |
| Tryptophan metabolism | 8/48 | 3.91E-05 | 1.67E-04 | 7.3007519 | 74.100859 | ALDH3A2;DDC;ECHS1;TDO2;EHHADH;CYP1B1;KMO;INMT |
| Inflammatory mediator regulation of TRP channels | 13/127 | 4.03E-05 | 1.69E-04 | 4.1861913 | 42.362908 | PLA2G4D;ADCY4;ITPR2;PIK3R3;TRPV3;ADCY2;PIK3R1;PIK3CB;ADCY8;ADCY6;ADCY5;ASIC4;ASIC2 |
| Nicotinate and nicotinamide metabolism | 7/36 | 4.25E-05 | 1.75E-04 | 8.7997024 | 88.569752 | BST1;NNMT;NT5C1A;QPRT;NMNAT2;NADK;NUDT12 |
| Folate biosynthesis | 6/26 | 5.47E-05 | 2.22E-04 | 10.921348 | 107.171 | MOCS2;GCH1;GGH;PCBD2;GPHN;PTS |
| Pyruvate metabolism | 7/38 | 6.14E-05 | 2.44E-04 | 8.2311324 | 79.82933 | ALDH3A2;PDHA1;MDH1;LDHD;ACYP2;PCK1;ACACA |
| Nitrogen metabolism | 5/17 | 6.66E-05 | 2.61E-04 | 15.146417 | 145.66316 | CAR3;CAR13;CAR12;GLUL;CAR5B |
| Dopaminergic synapse | 13/135 | 7.62E-05 | 2.94E-04 | 3.9100694 | 37.076705 | GRIA1;GRIA2;KCNJ5;DDC;KCNJ6;CACNA1B;ITPR2;ADCY5;SLC18A1;GRIA3;SLC18A2;GRIA4;KCNJ3 |
| Neuroactive ligand-receptor interaction | 23/348 | 7.98E-05 | 3.03E-04 | 2.6192828 | 24.716135 | GABRQ;CHRNB2;GRIA1;GRIA2;GABRB1;CHRNB4;CHRNA3;GRID2;CHRNA5;GRID1;GABRA4;CHRNA7;GABRA3;GRIK4;GRIK2;GABRG3;GABRG2;GRIN1;P2RX6;P2RX5;GRIN3A;GRIA3;GRIA4 |
| Glycosaminoglycan biosynthesis | 8/53 | 8.16E-05 | 3.05E-04 | 6.4878864 | 61.077491 | GLCE;DSE;NDST3;CSGALNACT2;CHST15;CHSY3;HS3ST5;HS3ST2 |
| Mucin type O-glycan biosynthesis | 6/28 | 8.56E-05 | 3.13E-04 | 9.927477 | 92.984435 | GALNT11;GALNT14;GALNT16;GALNT15;GALNT17;GCNT4 |
| Glycine, serine and threonine metabolism | 7/40 | 8.65E-05 | 3.13E-04 | 7.7314799 | 72.32742 | AOC3;SRR;ALAS1;GLDC;AGXT2;CHDH;SDSL |
| Insulin secretion | 10/86 | 1.07E-04 | 3.80E-04 | 4.8123138 | 44.018186 | KCNU1;KCNMB2;ATP1A4;ADCY4;ATP1A2;ADCY2;KCNN3;ADCY8;ADCY6;ADCY5 |
| Carbohydrate digestion and absorption | 7/43 | 1.39E-04 | 4.90E-04 | 7.0860955 | 62.907753 | MGAM;ATP1A4;PIK3R3;ATP1A2;AMY1;PIK3CB;PIK3R1 |
| Inositol phosphate metabolism | 9/73 | 1.49E-04 | 5.09E-04 | 5.1366525 | 45.260291 | IPPK;ALDH6A1;INPP5F;SYNJ1;PTEN;PLCE1;PIK3C2G;PIK3CB;PI4K2A |
| Taurine and hypotaurine metabolism | 4/11 | 1.49E-04 | 5.09E-04 | 20.738806 | 182.72533 | GGT7;GGT6;GAD1;CDO1 |
| Mineral absorption | 7/44 | 1.62E-04 | 5.45E-04 | 6.8942244 | 60.170374 | SLC9A3;SLC31A1;ATP1A4;ATP1A2;ATP2B1;CLCN2;SLC39A4 |
| Citrate cycle (TCA cycle) | 6/32 | 1.88E-04 | 6.23E-04 | 8.3984443 | 72.060901 | ACLY;PDHA1;MDH1;OGDH;ACO1;PCK1 |
| Cardiac muscle contraction | 9/78 | 2.49E-04 | 8.14E-04 | 4.7632031 | 39.533915 | COX8A;SLC9A6;ATP1A4;CACNA2D2;ATP1A2;UQCR10;CACNG2;CACNG3;CACNG5 |
| Estrogen signaling pathway | 12/134 | 2.82E-04 | 8.92E-04 | 3.602459 | 29.439261 | KCNJ5;KCNJ6;ADCY4;ITPR2;PIK3R3;ADCY2;PIK3R1;PIK3CB;ADCY8;ADCY6;ADCY5;KCNJ3 |
| Oxidative phosphorylation | 12/134 | 2.82E-04 | 8.92E-04 | 3.602459 | 29.439261 | ATP6V1A;COX8A;NDUFB8;NDUFS8;ATP6AP1;NDUFB6;COX15;NDUFA4L2;NDUFA1;ATP6V1H;NDUFV3;UQCR10 |
| Cocaine addiction | 7/48 | 2.84E-04 | 8.92E-04 | 6.2203359 | 50.805555 | GRIA2;DDC;GRIN3A;SLC18A1;SLC18A2;ADCY5;GRIN1 |
| Vascular smooth muscle contraction | 12/140 | 4.22E-04 | 0.0013111 | 3.4325284 | 26.669106 | GUCY1A1;KCNU1;GUCY1B1;PLA2G4D;NPR2;KCNMB2;ADCY4;ITPR2;ADCY2;ADCY8;ADCY6;ADCY5 |
| Parkinson disease | 12/144 | 5.45E-04 | 0.0016702 | 3.3278237 | 25.007114 | COX8A;NDUFB8;NDUFS8;NDUFB6;NDUFA4L2;NDUFA1;NDUFV3;UQCR10;SLC25A5;SLC18A1;SLC18A2;ADCY5 |
| AMPK signaling pathway | 11/126 | 6.22E-04 | 0.0018819 | 3.4979042 | 25.823252 | GYS2;CPT1A;PFKFB4;FASN;PIK3R3;PIK3R1;PIK3CB;SCD1;HMGCR;PCK1;ACACA |
| Dilated cardiomyopathy (DCM) | 9/90 | 7.24E-04 | 0.0021357 | 4.0550324 | 29.321781 | ADCY4;CACNA2D2;ADCY2;ADCY8;CACNG2;CACNG3;ADCY6;ADCY5;CACNG5 |
| Protein digestion and absorption | 9/90 | 7.24E-04 | 0.0021357 | 4.0550324 | 29.321781 | KCNK5;SLC36A1;SLC9A3;ACE2;SLC15A1;SLC7A8;ATP1A4;ATP1A2;SLC8A2 |
| Insulin resistance | 10/110 | 7.94E-04 | 0.0023135 | 3.6528302 | 26.07653 | GYS2;CPT1A;PTEN;PIK3R3;PYGM;PIK3R1;PIK3CB;PYGL;PCK1;SLC27A2 |
| Non-alcoholic fatty liver disease (NAFLD) | 12/151 | 8.32E-04 | 0.002395 | 3.1590909 | 22.404227 | COX8A;NDUFB8;NDUFS8;NDUFB6;NDUFA4L2;NDUFA1;PIK3R3;CYP2E1;PIK3R1;NDUFV3;PIK3CB;UQCR10 |
| Gastric acid secretion | 8/74 | 8.46E-04 | 0.0024069 | 4.4187742 | 31.263029 | ATP1A4;ADCY4;ITPR2;ATP1A2;ADCY2;ADCY8;ADCY6;ADCY5 |
| Pyrimidine metabolism | 7/58 | 9.19E-04 | 0.002583 | 4.998087 | 34.949779 | ENTPD3;ENTPD5;NT5C1A;DPYD;CMPK1;NME3;UPRT |
| Fatty acid elongation | 5/29 | 9.78E-04 | 0.0027177 | 7.5685358 | 52.450573 | ECHS1;MECR;ELOVL4;ELOVL5;ACOT4 |
| Synaptic vesicle cycle | 8/77 | 0.0011009 | 0.0030242 | 4.2259998 | 28.78609 | ATP6V1A;CACNA1B;SLC1A3;ATP6V1H;SLC6A2;SLC18A1;SLC18A2;SLC6A4 |
| Cushing syndrome | 12/159 | 0.0013035 | 0.0035401 | 2.9859307 | 19.834695 | STAR;CYP11A1;CYP11B2;HSD3B1;ADCY4;ITPR2;KCNA4;ADCY2;ADCY8;ADCY6;ADCY5;CYP17A1 |
| Arginine biosynthesis | 4/19 | 0.001475 | 0.003961 | 9.6741294 | 63.066477 | GOT1;GPT2;NOS1;GLUL |
| Ether lipid metabolism | 6/47 | 0.0015809 | 0.0041981 | 5.321732 | 34.324039 | PLA2G4D;CEPT1;PLD1;ENPP6;PLPP3;PLPP1 |
| Glucagon signaling pathway | 9/102 | 0.001767 | 0.0045903 | 3.5296155 | 22.372402 | GYS2;CPT1A;PDHA1;ITPR2;PYGM;ADCY2;PYGL;PCK1;ACACA |
| Longevity regulating pathway | 9/102 | 0.001767 | 0.0045903 | 3.5296155 | 22.372402 | ADCY4;PIK3R3;ADCY2;PIK3R1;PIK3CB;ADCY8;SOD2;ADCY6;ADCY5 |
| Starch and sucrose metabolism | 5/33 | 0.0017885 | 0.0045964 | 6.4859813 | 41.032619 | GYS2;MGAM;PYGM;AMY1;PYGL |
| Cysteine and methionine metabolism | 6/50 | 0.0021857 | 0.0054987 | 4.9581205 | 30.372565 | AHCYL2;GOT1;MDH1;AGXT2;SDSL;CDO1 |
| N-Glycan biosynthesis | 6/50 | 0.0021857 | 0.0054987 | 4.9581205 | 30.372565 | DPAGT1;MGAT4C;MAN1A;GANAB;ALG9;MGAT5 |
| Gap junction | 8/86 | 0.0022489 | 0.0055989 | 3.7366493 | 22.783487 | GUCY1A1;GUCY1B1;ADCY4;ITPR2;ADCY2;ADCY8;ADCY6;ADCY5 |
| Synthesis and degradation of ketone bodies | 3/11 | 0.002747 | 0.0067684 | 13.583799 | 80.106891 | BDH2;HMGCS1;HMGCLL1 |
| Alzheimer disease | 12/175 | 0.0029143 | 0.0071074 | 2.6906023 | 15.708049 | COX8A;NDUFB8;NDUFS8;NDUFB6;ATP2A3;NDUFA4L2;NDUFA1;ITPR2;NDUFV3;NOS1;UQCR10;GRIN1 |
| GnRH signaling pathway | 8/90 | 0.0029926 | 0.0071524 | 3.5536402 | 20.652339 | PLA2G4D;ADCY4;ITPR2;ADCY2;PLD1;ADCY8;ADCY6;ADCY5 |
| Progesterone-mediated oocyte maturation | 8/90 | 0.0029926 | 0.0071524 | 3.5536402 | 20.652339 | ADCY4;PIK3R3;ADCY2;PIK3CB;PIK3R1;ADCY8;ADCY6;ADCY5 |
| Phenylalanine metabolism | 4/23 | 0.003094 | 0.0072496 | 7.6358995 | 44.122531 | AOC3;DDC;GOT1;ALDH3B1 |
| Terpenoid backbone biosynthesis | 4/23 | 0.003094 | 0.0072496 | 7.6358995 | 44.122531 | FDPS;IDI1;HMGCS1;HMGCR |
| Aldosterone-regulated sodium reabsorption | 5/38 | 0.0033869 | 0.0078589 | 5.5018408 | 31.293641 | ATP1A4;PIK3R3;ATP1A2;PIK3CB;PIK3R1 |
| Histidine metabolism | 4/24 | 0.0036345 | 0.0083524 | 7.2537313 | 40.746219 | ALDH3A2;ALDH3B1;HNMT;ASPA |
| Fat digestion and absorption | 5/40 | 0.0042484 | 0.0096702 | 5.1869159 | 28.326808 | ABCA1;DGAT2;PLPP3;AGPAT2;PLPP1 |
| Huntington disease | 12/192 | 0.0061021 | 0.0137584 | 2.4343434 | 12.413033 | COX8A;NDUFB8;NDUFS8;NDUFB6;UCP1;NDUFA4L2;NDUFA1;NDUFV3;UQCR10;SLC25A5;SOD2;GRIN1 |
| Glutathione metabolism | 6/64 | 0.0075193 | 0.0167954 | 3.7586207 | 18.380719 | GSTK1;GGT7;GGT6;GPX3;MGST1;MGST2 |
| Primary bile acid biosynthesis | 3/16 | 0.0084332 | 0.0186625 | 8.3571122 | 39.910005 | CYP27A1;AKR1D1;CYP7B1 |
| Relaxin signaling pathway | 9/131 | 0.0092231 | 0.0201504 | 2.6865796 | 12.589431 | ADCY4;PIK3R3;ADCY2;PIK3R1;PIK3CB;NOS1;ADCY8;ADCY6;ADCY5 |
| Type II diabetes mellitus | 5/48 | 0.0092742 | 0.0201504 | 4.2201695 | 19.752565 | CACNA1B;PIK3R3;PIK3CB;PIK3R1;CACNA1E |
| Arachidonic acid metabolism | 7/89 | 0.0103248 | 0.0222308 | 3.1036013 | 14.19342 | CYP4F18;PLA2G4D;GPX3;EPHX2;CYP2E1;PTGS2;PTGS1 |
| Arginine and proline metabolism | 5/50 | 0.0109844 | 0.02344 | 4.0321911 | 18.190328 | ALDH3A2;GOT1;AZIN2;NOS1;PRODH |
| Apelin signaling pathway | 9/138 | 0.0126956 | 0.0268517 | 2.5398765 | 11.090377 | UCP1;ADCY4;ITPR2;ADCY2;NOS1;ADCY8;ADCY6;SLC8A2;ADCY5 |
| Pentose and glucuronate interconversions | 4/34 | 0.0128347 | 0.0269079 | 4.8333333 | 21.052063 | UGT1A1;DCXR;DHDH;CRYL1 |
| Insulin signaling pathway | 9/139 | 0.0132615 | 0.0275609 | 2.5202086 | 10.894579 | GYS2;FASN;PIK3R3;PYGM;PIK3R1;PIK3CB;PYGL;PCK1;ACACA |
| One carbon pool by folate | 3/19 | 0.0137448 | 0.028319 | 6.7891061 | 29.105559 | ALDH1L1;MTHFD2L;MTHFS |
| VEGF signaling pathway | 5/58 | 0.0199552 | 0.0407633 | 3.4221478 | 13.395187 | PLA2G4D;PIK3R3;PIK3CB;PIK3R1;PTGS2 |
| Ferroptosis | 4/40 | 0.0222976 | 0.0451621 | 4.026534 | 15.314023 | ACSL6;ACSL4;ACSL3;CP |
| Sulfur metabolism | 2/9 | 0.0230994 | 0.0462136 | 10.330855 | 38.926126 | BPNT1;SUOX |
| Mannose type O-glycan biosynthesis | 3/23 | 0.0232035 | 0.0462136 | 5.4301676 | 20.436185 | POMT2;FUT9;B3GAT1 |
| Porphyrin and chlorophyll metabolism | 4/41 | 0.0241965 | 0.047793 | 3.9175071 | 14.579195 | ALAS1;UGT1A1;COX15;CP |
| Parathyroid hormone synthesis, secretion and action | 7/107 | 0.0259681 | 0.0505184 | 2.5425891 | 9.2827089 | ADCY4;ITPR2;ADCY2;PLD1;ADCY8;ADCY6;ADCY5 |
| Vitamin digestion and absorption | 3/24 | 0.025999 | 0.0505184 | 5.1713222 | 18.873754 | BTD;FOLH1;MMACHC |
| Glycosylphosphatidylinositol (GPI)-anchor biosynthesis | 3/25 | 0.0289668 | 0.0553066 | 4.9360081 | 17.481398 | PIGYL;PIGK;PIGG |
| alpha-Linolenic acid metabolism | 3/25 | 0.0289668 | 0.0553066 | 4.9360081 | 17.481398 | FADS2;PLA2G4D;ACOX1 |
| Central carbon metabolism in cancer | 5/64 | 0.0291575 | 0.0553066 | 3.0731823 | 10.863836 | PDHA1;PTEN;PIK3R3;PIK3CB;PIK3R1 |
| Fc gamma R-mediated phagocytosis | 6/87 | 0.0303209 | 0.0570606 | 2.6881676 | 9.3976119 | PIK3R3;PIK3CB;PIK3R1;PLPP3;PLD1;PLPP1 |
| Glycolysis / Gluconeogenesis | 5/67 | 0.034617 | 0.0641354 | 2.9240277 | 9.8347027 | ALDH3A2;PDHA1;ALDH3B1;ADH7;PCK1 |
| Long-term potentiation | 5/67 | 0.034617 | 0.0641354 | 2.9240277 | 9.8347027 | GRIA1;GRIA2;ITPR2;ADCY8;GRIN1 |
| Thyroid hormone signaling pathway | 7/115 | 0.0364345 | 0.0669834 | 2.3532764 | 7.7946168 | SLCO1C1;ATP1A4;PLCE1;PIK3R3;ATP1A2;PIK3R1;PIK3CB |
| Adipocytokine signaling pathway | 5/71 | 0.0428201 | 0.0775942 | 2.7462475 | 8.6527335 | CPT1A;ACSL6;ACSL4;ACSL3;PCK1 |
| Cholesterol metabolism | 4/49 | 0.0428554 | 0.0775942 | 3.2197347 | 10.141919 | CYP27A1;ABCA1;STAR;LIPG |
| Human cytomegalovirus infection | 12/255 | 0.0448759 | 0.0806416 | 1.7973251 | 5.5786361 | ADCY4;TAP2;PIK3R3;TAP1;ITPR2;ADCY2;PIK3R1;PIK3CB;ADCY8;PTGS2;ADCY6;ADCY5 |
| Lysosome | 7/124 | 0.0511574 | 0.0911074 | 2.1712448 | 6.4547819 | MFSD8;GNPTG;ATP6AP1;SMPD1;SLC17A5;ATP6V1H;ABCB9 |
| Amyotrophic lateral sclerosis (ALS) | 4/52 | 0.0514624 | 0.0911074 | 3.0180348 | 8.9542218 | GRIA1;GRIA2;NOS1;GRIN1 |
| Rap1 signaling pathway | 10/209 | 0.057958 | 0.1018527 | 1.8262065 | 5.201102 | ADCY4;PLCE1;PIK3R3;ADCY2;PIK3R1;PIK3CB;ADCY8;ADCY6;ADCY5;GRIN1 |
| Thiamine metabolism | 2/15 | 0.0605845 | 0.1055445 | 5.5610523 | 15.59161 | TPK1;AK5 |
| Endocrine and other factor-regulated calcium reabsorptio | 4/55 | 0.060942 | 0.1055445 | 2.8400644 | 7.9460251 | ATP1A4;ATP1A2;ATP2B1;ADCY6 |
| Human T-cell leukemia virus 1 infection | 11/245 | 0.0694899 | 0.1194826 | 1.7084808 | 4.5557911 | FDPS;ADCY4;PTEN;PIK3R3;ADCY2;PIK3R1;PIK3CB;ADCY8;SLC25A5;ADCY6;ADCY5 |
| Endometrial cancer | 4/58 | 0.0712788 | 0.1216831 | 2.6818684 | 7.0832347 | PTEN;PIK3R3;PIK3CB;PIK3R1 |
| Lysine degradation | 4/59 | 0.0749111 | 0.1269769 | 2.6329715 | 6.8232234 | ALDH3A2;COLGALT2;ECHS1;EHHADH |
| Selenocompound metabolism | 2/17 | 0.0757676 | 0.1275244 | 4.819083 | 12.433638 | SCLY;INMT |
| Hypertrophic cardiomyopathy (HCM) | 5/86 | 0.0831709 | 0.139006 | 2.2359525 | 5.5604946 | ACE;CACNA2D2;CACNG2;CACNG3;CACNG5 |
| Pantothenate and CoA biosynthesis | 2/18 | 0.0837627 | 0.1390228 | 4.517658 | 11.202742 | PANK3;DPYD |
| Oocyte meiosis | 6/116 | 0.0943943 | 0.1555879 | 1.9765066 | 4.6650974 | ADCY4;ITPR2;ADCY2;ADCY8;ADCY6;ADCY5 |
| Retinol metabolism | 5/91 | 0.0999236 | 0.1635736 | 2.1054119 | 4.8494987 | UGT1A1;RDH11;CYP2S1;ADH7;DHRS3 |
| Small cell lung cancer | 5/92 | 0.1034631 | 0.1682155 | 2.0811043 | 4.721069 | PTEN;PIK3R3;PIK3CB;PIK3R1;PTGS2 |
| Fc epsilon RI signaling pathway | 4/68 | 0.1115834 | 0.1801921 | 2.2616604 | 4.9597827 | PLA2G4D;PIK3R3;PIK3CB;PIK3R1 |
| Other types of O-glycan biosynthesis | 2/22 | 0.1179821 | 0.1892465 | 3.6133829 | 7.7226013 | POMT2;COLGALT2 |
| Prostate cancer | 5/97 | 0.1220671 | 0.1944935 | 1.9674929 | 4.1380008 | SRD5A2;PTEN;PIK3R3;PIK3R1;PIK3CB |
| mTOR signaling pathway | 7/154 | 0.1241224 | 0.1964586 | 1.7254534 | 3.6001363 | ATP6V1A;PTEN;PIK3R3;ATP6V1H;PIK3R1;PIK3CB;LPIN1 |
| Arrhythmogenic right ventricular cardiomyopathy (ARVC) | 4/72 | 0.1300128 | 0.2017731 | 2.1281826 | 4.3417528 | CACNA2D2;CACNG2;CACNG3;CACNG5 |
| Melanoma | 4/72 | 0.1300128 | 0.2017731 | 2.1281826 | 4.3417528 | PTEN;PIK3R3;PIK3CB;PIK3R1 |
| Prolactin signaling pathway | 4/72 | 0.1300128 | 0.2017731 | 2.1281826 | 4.3417528 | PIK3R3;PIK3CB;PIK3R1;CYP17A1 |
| Melanogenesis | 5/100 | 0.1339257 | 0.2065048 | 1.9050664 | 3.8300788 | ADCY4;ADCY2;ADCY8;ADCY6;ADCY5 |
| Glioma | 4/75 | 0.1446046 | 0.2201306 | 2.0379441 | 3.9408783 | PTEN;PIK3R3;PIK3CB;PIK3R1 |
| Pancreatic cancer | 4/75 | 0.1446046 | 0.2201306 | 2.0379441 | 3.9408783 | PIK3R3;PIK3CB;PIK3R1;PLD1 |
| FoxO signaling pathway | 6/132 | 0.1476111 | 0.2232852 | 1.7240949 | 3.2984937 | PTEN;PIK3R3;PIK3R1;PIK3CB;PCK1;SOD2 |
| Phosphonate and phosphinate metabolism | 1/6 | 0.1514681 | 0.2276784 | 7.2189239 | 13.624856 | CEPT1 |
| Ascorbate and aldarate metabolism | 2/27 | 0.1645302 | 0.2449869 | 2.8899628 | 5.2154043 | ALDH3A2;UGT1A1 |
| Chemokine signaling pathway | 8/197 | 0.165033 | 0.2449869 | 1.5332776 | 2.7623681 | ADCY4;PIK3R3;ADCY2;PIK3R1;PIK3CB;ADCY8;ADCY6;ADCY5 |
| Epstein-Barr virus infection | 9/229 | 0.1683166 | 0.2483189 | 1.4822804 | 2.6412886 | ENTPD3;OAS2;OAS3;TAP2;PIK3R3;TAP1;OAS1B;PIK3R1;PIK3CB |
| Influenza A | 7/168 | 0.1701496 | 0.2494832 | 1.574272 | 2.7881571 | FDPS;OAS2;OAS3;PIK3R3;OAS1B;PIK3R1;PIK3CB |
| C-type lectin receptor signaling pathway | 5/112 | 0.186016 | 0.2710843 | 1.690366 | 2.8430649 | ITPR2;PIK3R3;PIK3R1;PIK3CB;PTGS2 |
| Measles | 6/144 | 0.1946562 | 0.2798073 | 1.5731965 | 2.574568 | OAS2;OAS3;PIK3R3;OAS1B;PIK3R1;PIK3CB |
| Phenylalanine, tyrosine and tryptophan biosynthesis | 1/8 | 0.1966846 | 0.2798073 | 5.1558442 | 8.3841947 | GOT1 |
| Riboflavin metabolism | 1/8 | 0.1966846 | 0.2798073 | 5.1558442 | 8.3841947 | RFK |
| Sulfur relay system | 1/8 | 0.1966846 | 0.2798073 | 5.1558442 | 8.3841947 | MOCS2 |
| Primary immunodeficiency | 2/36 | 0.2537571 | 0.3567527 | 2.1239886 | 2.9127906 | TAP2;TAP1 |
| Renin-angiotensin system | 2/36 | 0.2537571 | 0.3567527 | 2.1239886 | 2.9127906 | ACE2;ACE |
| Non-small cell lung cancer | 3/66 | 0.2636316 | 0.3676879 | 1.7200497 | 2.2931745 | PIK3R3;PIK3CB;PIK3R1 |
| Hepatitis C | 6/160 | 0.2646122 | 0.3676879 | 1.4085802 | 1.8726932 | OAS2;OAS3;PIK3R3;OAS1B;PIK3R1;PIK3CB |
| Renal cell carcinoma | 3/68 | 0.2783218 | 0.3845024 | 1.6669532 | 2.131995 | PIK3R3;PIK3CB;PIK3R1 |
| Acute myeloid leukemia | 3/69 | 0.2856967 | 0.3924224 | 1.6416116 | 2.0566515 | PIK3R3;PIK3CB;PIK3R1 |
| AGE-RAGE signaling pathway in diabetic complications | 4/101 | 0.2908506 | 0.3972188 | 1.4896907 | 1.839687 | PLCE1;PIK3R3;PIK3R1;PIK3CB |
| Chagas disease (American trypanosomiasis) | 4/103 | 0.3030084 | 0.4114716 | 1.4594452 | 1.7425699 | ACE;PIK3R3;PIK3R1;PIK3CB |
| B cell receptor signaling pathway | 3/72 | 0.3079018 | 0.4150393 | 1.5699943 | 1.8494132 | PIK3R3;PIK3CB;PIK3R1 |
| HIF-1 signaling pathway | 4/104 | 0.3091088 | 0.4150393 | 1.4447761 | 1.6962568 | PDHA1;PIK3R3;PIK3R1;PIK3CB |
| Hepatocellular carcinoma | 6/171 | 0.3159414 | 0.4218435 | 1.3139258 | 1.5139034 | MGST1;PTEN;MGST2;PIK3R3;PIK3R1;PIK3CB |
| Bacterial invasion of epithelial cells | 3/74 | 0.3227419 | 0.4285296 | 1.5256118 | 1.7253177 | PIK3R3;PIK3CB;PIK3R1 |
| Chronic myeloid leukemia | 3/76 | 0.3375845 | 0.4457608 | 1.4836611 | 1.6111659 | PIK3R3;PIK3CB;PIK3R1 |
| Necroptosis | 6/176 | 0.33979 | 0.4462078 | 1.2749504 | 1.3762164 | PLA2G4D;SMPD1;PYGM;PYGL;SLC25A5;GLUL |
| Fluid shear stress and atherosclerosis | 5/143 | 0.3435237 | 0.4486457 | 1.3085467 | 1.3981811 | MGST1;MGST2;PIK3R3;PIK3R1;PIK3CB |
| TNF signaling pathway | 4/110 | 0.3459032 | 0.4492981 | 1.3625739 | 1.4465036 | PIK3R3;PIK3R1;PIK3CB;PTGS2 |
| Axon guidance | 6/180 | 0.3590045 | 0.4613014 | 1.2453829 | 1.2757958 | TRPC5;DPYSL5;TRPC1;PIK3R3;PIK3R1;PIK3CB |
| Phagosome | 6/180 | 0.3590045 | 0.4613014 | 1.2453829 | 1.2757958 | ATP6V1A;ATP6AP1;TAP2;TAP1;ATP6V1H;NOS1 |
| Cellular senescence | 6/185 | 0.38311 | 0.4896433 | 1.2102818 | 1.1611845 | PTEN;ITPR2;PIK3R3;PIK3R1;PIK3CB;SLC25A5 |
| Linoleic acid metabolism | 2/50 | 0.392598 | 0.4988333 | 1.5034077 | 1.4056398 | PLA2G4D;CYP2E1 |
| ErbB signaling pathway | 3/84 | 0.3965621 | 0.4988333 | 1.3365749 | 1.2362285 | PIK3R3;PIK3CB;PIK3R1 |
| Rheumatoid arthritis | 3/84 | 0.3965621 | 0.4988333 | 1.3365749 | 1.2362285 | ATP6V1A;ATP6AP1;ATP6V1H |
| Colorectal cancer | 3/88 | 0.4255564 | 0.5325025 | 1.2734144 | 1.0879516 | PIK3R3;PIK3CB;PIK3R1 |
| Ras signaling pathway | 7/233 | 0.4412094 | 0.5492138 | 1.1177174 | 0.9145563 | PLA2G4D;PLCE1;PIK3R3;PIK3R1;PIK3CB;PLD1;GRIN1 |
| Alcoholism | 6/199 | 0.4503787 | 0.5577229 | 1.1216743 | 0.894722 | DDC;GRIN3A;SLC18A1;SLC18A2;ADCY5;GRIN1 |
| Autophagy | 4/130 | 0.4673682 | 0.5757783 | 1.1451078 | 0.8710124 | PTEN;PIK3R3;PIK3R1;PIK3CB |
| Toll-like receptor signaling pathway | 3/99 | 0.5024106 | 0.6157751 | 1.1268622 | 0.7756616 | PIK3R3;PIK3R1;PIK3CB |
| T cell receptor signaling pathway | 3/101 | 0.5158233 | 0.6289886 | 1.103751 | 0.7306733 | PIK3R3;PIK3R1;PIK3CB |
| Collecting duct acid secretion | 1/27 | 0.5226514 | 0.6308772 | 1.3867561 | 0.8997836 | ATP6V1A |
| Phototransduction | 1/27 | 0.5226514 | 0.6308772 | 1.3867561 | 0.8997836 | PDE6A |
| Apoptosis | 4/141 | 0.5307747 | 0.6374631 | 1.0525656 | 0.6667137 | ITPR2;PIK3R3;PIK3R1;PIK3CB |
| Amoebiasis | 3/106 | 0.5484821 | 0.6554361 | 1.0498997 | 0.6305704 | PIK3R3;PIK3R1;PIK3CB |
| Breast cancer | 4/147 | 0.5637562 | 0.670337 | 1.0080889 | 0.5777694 | PTEN;PIK3R3;PIK3R1;PIK3CB |
| Gastric cancer | 4/150 | 0.5797636 | 0.6839501 | 0.9872214 | 0.5381689 | PIK3R3;PIK3R1;PIK3CB;ABCB1B |
| Galactose metabolism | 1/32 | 0.58379 | 0.6839501 | 1.1627865 | 0.6258279 | MGAM |
| Pentose phosphate pathway | 1/32 | 0.58379 | 0.6839501 | 1.1627865 | 0.6258279 | RBKS |
| Leukocyte transendothelial migration | 3/115 | 0.6038674 | 0.704021 | 0.9650838 | 0.4867889 | PIK3R3;PIK3R1;PIK3CB |
| Fructose and mannose metabolism | 1/35 | 0.6166536 | 0.7154379 | 1.060024 | 0.5124663 | PFKFB4 |
| Natural killer cell mediated cytotoxicity | 3/118 | 0.6213013 | 0.7172908 | 0.939762 | 0.4472695 | PIK3R3;PIK3R1;PIK3CB |
| Human immunodeficiency virus 1 infection | 6/238 | 0.6242531 | 0.7172908 | 0.9312282 | 0.4387941 | TAP2;PIK3R3;TAP1;ITPR2;PIK3R1;PIK3CB |
| Neurotrophin signaling pathway | 3/121 | 0.6382056 | 0.7272143 | 0.9157277 | 0.4112486 | PIK3R3;PIK3R1;PIK3CB |
| Human papillomavirus infection | 9/360 | 0.639436 | 0.7272143 | 0.9227389 | 0.41262 | ATP6V1A;ATP6AP1;PTEN;PIK3R3;ATP6V1H;PIK3R1;PIK3CB;PTGS2;OASL2 |
| Proteoglycans in cancer | 5/203 | 0.6438667 | 0.7272143 | 0.9091853 | 0.4002812 | PLCE1;PIK3R3;ITPR2;PIK3R1;PIK3CB |
| Hepatitis B | 4/163 | 0.6450604 | 0.7272143 | 0.9058951 | 0.3971546 | ATP6AP1;PIK3R3;PIK3R1;PIK3CB |
| NOD-like receptor signaling pathway | 5/205 | 0.652367 | 0.7319987 | 0.9 | 0.3844332 | TRPM2;OAS2;OAS3;ITPR2;OAS1B |
| Osteoclast differentiation | 3/128 | 0.675565 | 0.7544862 | 0.8641341 | 0.3389184 | PIK3R3;PIK3R1;PIK3CB |
| MAPK signaling pathway | 7/294 | 0.684061 | 0.7604213 | 0.8773624 | 0.3331417 | PLA2G4D;CACNA1B;CACNA2D2;CACNG2;CACNA1E;CACNG3;CACNG5 |
| Vasopressin-regulated water reabsorption | 1/43 | 0.6921717 | 0.7658752 | 0.8577613 | 0.3155886 | ADCY6 |
| Kaposi sarcoma-associated herpesvirus infection | 5/216 | 0.69677 | 0.7674102 | 0.8525933 | 0.3080419 | PIK3R3;ITPR2;PIK3R1;PIK3CB;PTGS2 |
| Antigen processing and presentation | 2/90 | 0.702896 | 0.7706062 | 0.8183508 | 0.2885066 | TAP2;TAP1 |
| Signaling pathways regulating pluripotency of stem cells | 3/137 | 0.7193105 | 0.785001 | 0.80572 | 0.2654542 | PIK3R3;PIK3R1;PIK3CB |
| Amino sugar and nucleotide sugar metabolism | 1/49 | 0.7388915 | 0.8027048 | 0.7503092 | 0.2270467 | UAP1 |
| Pathways in cancer | 12/535 | 0.7828464 | 0.8466076 | 0.8229185 | 0.2014659 | ADCY4;PTEN;MGST1;MGST2;PIK3R3;ADCY2;PIK3CB;PIK3R1;ADCY8;PTGS2;ADCY6;ADCY5 |
| Focal adhesion | 4/199 | 0.788787 | 0.8491897 | 0.7372752 | 0.1749251 | PTEN;PIK3R3;PIK3R1;PIK3CB |
| JAK-STAT signaling pathway | 3/164 | 0.8231888 | 0.8822517 | 0.6696624 | 0.130296 | PIK3R3;PIK3R1;PIK3CB |
| Leishmaniasis | 1/67 | 0.8406954 | 0.896992 | 0.5451734 | 0.0946017 | PTGS2 |
| Type I diabetes mellitus | 1/69 | 0.84921 | 0.9020497 | 0.5290844 | 0.0864782 | GAD1 |
| p53 signaling pathway | 1/71 | 0.8572702 | 0.9065822 | 0.5139147 | 0.0791439 | PTEN |
| ECM-receptor interaction | 1/83 | 0.8973603 | 0.9447978 | 0.4384361 | 0.0474817 | SV2A |
| Herpes simplex virus 1 infection | 8/433 | 0.9017544 | 0.9452601 | 0.6735073 | 0.0696495 | OAS2;OAS3;TAP2;TAP1;PIK3R3;OAS1B;PIK3CB;PIK3R1 |
| IL-17 signaling pathway | 1/91 | 0.9176239 | 0.9576948 | 0.3992991 | 0.0343268 | PTGS2 |
| PI3K-Akt signaling pathway | 6/357 | 0.9226429 | 0.9587463 | 0.6117033 | 0.0492501 | GYS2;PTEN;PIK3R3;PIK3R1;PIK3CB;PCK1 |
| Regulation of actin cytoskeleton | 3/217 | 0.9349952 | 0.9633131 | 0.5024278 | 0.0337701 | PIK3R3;PIK3R1;PIK3CB |
| Protein processing in endoplasmic reticulum | 2/163 | 0.936967 | 0.9633131 | 0.4456118 | 0.0290125 | MAN1A;GANAB |
| NF-kappa B signaling pathway | 1/102 | 0.9391295 | 0.9633131 | 0.355609 | 0.0223329 | PTGS2 |
| MicroRNAs in cancer | 4/281 | 0.9475955 | 0.9650138 | 0.5168113 | 0.0278187 | PTEN;CYP1B1;PTGS2;ABCB1B |
| Viral carcinogenesis | 3/229 | 0.9488629 | 0.9650138 | 0.4754536 | 0.024957 | PIK3R3;PIK3R1;PIK3CB |
| Tuberculosis | 2/178 | 0.9551668 | 0.9673088 | 0.4073167 | 0.0186833 | ATP6AP1;ATP6V1H |
| Transcriptional misregulation in cancer | 2/183 | 0.9600278 | 0.9681293 | 0.3959621 | 0.0161525 | HPGD;GRIA3 |
| Endocytosis | 1/269 | 0.9993939 | 0.9999952 | 0.1328607 | 8.05E-05 | PLD1 |
| Olfactory transduction | 6/1133 | 0.9999952 | 0.9999952 | 0.1827762 | 8.73E-07 | SLC24A4;PDE1C;PDE1B;PDE1A;CNGA3;SLC8A2 |
