## Supplementary material for "Glutamine antagonism suppresses tumor growth in adrenocortical carcinoma through inhibition of de novo nucleotide biosynthesis": Suppl. Table 2B

**Suppl. Table 2b Down-regulated pathways Human ACC (TCGA) versus normal adrenals (GTEx)**

| Term | Overlap | P-value | Adjusted P-va | Odds Ratio | Combined Sc | Genes |
|---|---|---|---|---|---|---|
| Phosphatidylinositol signaling system | 47/99 | 1.52E-34 | 3.91E-32 | 17.755326 | 1382.6293 | DGKG;DGKE;DGKD;DGKB;INPP1;DGKA;INPPL1;PTEN;ITPR2;PIK3CD;ITPR3;[illegible] |
| Inositol phosphate metabolism | 36/74 | 3.16E-27 | 3.06E-25 | 18.413927 | 1123.5943 | INPP1;INPPL1;PTEN;PIK3CD;PIK3C2G;PIK3C2A;PIK3CG;PIK3C2B;INPP5B;[illegible] |
| Purine metabolism | 46/129 | 3.56E-27 | 3.06E-25 | 10.858056 | 661.24656 | GUCY2D;CANT1;PDE1C;GDA;PDE1B;NPR1;PDE1A;NPR2;AK2;GMPS;ADCY4;[illegible] |
| Arachidonic acid metabolism | 31/63 | 8.09E-24 | 5.22E-22 | 18.739381 | 996.38844 | CYP4A22;ALOX15;ALOX12;PLA2G6;CYP2C19;PTGS2;LTC4S;PLB1;PTGS1;[illegible] |
| Serotonergic synapse | 39/113 | 1.24E-22 | 6.40E-21 | 10.255881 | 517.32256 | GABRB3;TPH2;GABRB2;TPH1;DDC;ALOX15;CACNA1A;ITPR2;CACNA1D;[illegible] |
| Drug metabolism | 35/108 | 1.98E-19 | 7.98E-18 | 9.292331 | 400.16431 | TPMT;ADH1B;DPYS;GSTP1;MGST1;GMPS;CYP2C19;ADH7;MPO;CYP3A5;[illegible] |
| Glycerolipid metabolism | 27/61 | 2.17E-19 | 7.98E-18 | 15.297256 | 657.42507 | DGKG;PNLIPRP1;DGKE;DGKD;MOGAT2;DGKB;MOGAT1;DGKA;AKR1B1;[illegible] |
| Nicotine addiction | 22/40 | 1.15E-18 | 3.72E-17 | 23.444669 | 968.34968 | GABRB3;CHRNB2;GABRA2;GRIA2;GABRB2;GABRP;GABRA1;SLC32A1;[illegible] |
| ABC transporters | 23/45 | 1.71E-18 | 4.89E-17 | 20.069976 | 821.10666 | ABCD4;ABCA1;ABCC3;ABCG8;ABCD3;ABCB1;ABCC2;ABCA5;ABCB7;[illegible] |
| Retrograde endocannabinoid signaling | 39/148 | 5.92E-18 | 1.53E-16 | 6.9498244 | 275.68411 | GABRB3;GRIA2;GABRB2;FAAH;NDUFA10;ADCY4;CACNA1A;ITPR2;[illegible] |
| Glycerophospholipid metabolism | 31/97 | 3.63E-17 | 8.51E-16 | 9.0694651 | 343.32849 | DGKG;DGKE;DGKD;DGKB;DGKA;LCAT;PLA2G6;PLD1;PLB1;AGPAT3;PLD2;[illegible] |
| Morphine addiction | 30/91 | 4.40E-17 | 9.47E-16 | 9.4891462 | 357.37489 | GABRB3;GABRB2;PDE1C;PDE1B;PDE1A;ADCY4;CACNA1A;ADCY2;ADCY1;[illegible] |
| Glutamatergic synapse | 33/114 | 9.88E-17 | 1.96E-15 | 7.8765432 | 290.27881 | GRIA2;GRIK5;ADCY4;CACNA1A;ITPR2;CACNA1D;ITPR3;GRIK1;ADCY2;[illegible] |
| Valine, leucine and isoleucine degradation | 22/48 | 1.89E-16 | 3.49E-15 | 16.22408 | 587.37746 | ACAD8;ACAA2;AGXT2;BCKDHB;MCCC1;ACADSB;HADHB;ALDH3A2;HMGCL;[illegible] |
| Pancreatic secretion | 30/98 | 4.47E-16 | 7.69E-15 | 8.5091743 | 300.74744 | PNLIPRP1;AMY2A;AMY2B;ATP1A4;ADCY4;ITPR2;ATP2A2;ITPR3;ATP2A1;[illegible] |
| Oxytocin signaling pathway | 37/153 | 8.75E-16 | 1.41E-14 | 6.18053 | 214.29517 | NPR1;NPR2;ADCY4;ITPR2;CACNA1D;ITPR3;ADCY2;ADCY1;CACNA1F;[illegible] |
| Retinol metabolism | 23/67 | 7.57E-14 | 1.15E-12 | 10.023348 | 302.82113 | DGAT1;RETSAT;ADH1B;CYP4A22;CYP4A11;UGT2B15;ADH7;CYP3A5;[illegible] |
| Cholinergic synapse | 29/112 | 1.73E-13 | 2.48E-12 | 6.7267931 | 197.65862 | CHRNA3;ADCY4;CACNA1A;ITPR2;PIK3CD;CACNA1D;ITPR3;ADCY2;ADCY1;[illegible] |
| Taste transduction | 25/83 | 1.91E-13 | 2.59E-12 | 8.2757746 | 242.38856 | KCNK5;HCN4;PDE1C;PDE1B;PDE1A;ADCY4;CACNA1A;ITPR3;ADCY6;SCN9A;[illegible] |
| Adrenergic signaling in cardiomyocytes | 33/145 | 2.03E-13 | 2.62E-12 | 5.6870892 | 166.21317 | KCNE1;ATP1A4;ADCY4;ATP2A2;CACNA1D;ATP1A2;ADCY2;ADCY1;CACNA1F;[illegible] |
| Propanoate metabolism | 16/32 | 4.53E-13 | 5.56E-12 | 19.068342 | 541.98505 | ACSS3;BCKDHB;ACACB;ACACA;HADHA;MCEE;ALDH6A1;SUCLA2;PCCA;[illegible] |
| GABAergic synapse | 25/89 | 1.10E-12 | 1.29E-11 | 7.4975437 | 206.42406 | GABRB3;GABRB2;ADCY4;CACNA1A;ADCY2;CACNA1D;ADCY1;CACNA1F;[illegible] |
| Bile secretion | 22/72 | 3.87E-12 | 4.34E-11 | 8.4258443 | 221.40745 | ABCC3;ABCG8;ABCC2;ABCB1;SLC10A1;AQP9;SLC22A1;ATP1A4;ADCY4;[illegible] |
| Tyrosine metabolism | 16/36 | 4.55E-12 | 4.89E-11 | 15.251457 | 398.31898 | PNMT;DDC;AOC2;HGD;ADH1B;TAT;DBH;ADH7;ALDH1A3;ALDH3A1;TPO;TH;[illegible] |
| Gastric acid secretion | 22/75 | 9.65E-12 | 9.96E-11 | 7.9476506 | 201.58109 | KCNJ10;KCNK10;ATP1A4;ADCY4;ITPR2;ATP1B3;ITPR3;ATP1A2;ADCY2;ADCY1;[illegible] |
| Phospholipase D signaling pathway | 31/148 | 1.10E-11 | 1.10E-10 | 5.1023199 | 128.72536 | DGKG;DGKE;DGKD;DGKB;DGKA;ADCY4;PIK3CD;ADCY2;ADCY1;PLD1;[illegible] |
| Renin secretion | 21/69 | 1.30E-11 | 1.24E-10 | 8.3703914 | 209.80765 | GUCY1A2;ACE;PDE1C;NPR1;PDE1B;PDE1A;ITPR2;ITPR3;CACNA1D;[illegible] |
| Fructose and mannose metabolism | 15/33 | 1.43E-11 | 1.32E-10 | 15.872657 | 396.36724 | PFKFB2;PFKFB1;PFKFB3;MPI;SORD;AKR1B1;GMPPB;HK3;AKR1B10;GMDS;[illegible] |
| Fatty acid degradation | 17/44 | 1.52E-11 | 1.35E-10 | 12.011104 | 299.23054 | ACADVL;CPT1A;ACAA2;ADH1B;ACSL1;CYP4A22;CYP4A11;ADH7;CPT1B;[illegible] |
| Insulin secretion | 23/86 | 2.69E-11 | 2.31E-10 | 6.993413 | 170.21688 | KCNJ11;ABCC8;ATP1A4;ADCY4;SLC2A2;ATP1B3;ITPR3;ATP1A2;ADCY2;[illegible] |
| Glycine, serine and threonine metabolism | 16/40 | 3.23E-11 | 2.69E-10 | 12.706868 | 306.95115 | DMGDH;ALAS2;ALAS1;SDS;AOC2;GLDC;AGXT2;GLYCTK;AMT;PIPOX;GCSH;[illegible] |
| Metabolism of xenobiotics by cytochrome P450 | 21/74 | 5.74E-11 | 4.62E-10 | 7.5787307 | 178.71939 | ADH1B;GSTP1;UGT2B15;MGST1;ADH7;CYP3A5;HSD11B1;ALDH1A3;[illegible] |
| Circadian entrainment | 24/97 | 5.99E-11 | 4.68E-10 | 6.3008702 | 148.31194 | GUCY1A2;GRIA2;KCNJ5;KCNJ9;ADCY4;CACNA1D;ITPR3;ADCY2;ADCY1;[illegible] |
| beta-Alanine metabolism | 14/31 | 7.70E-11 | 5.73E-10 | 15.671013 | 364.9252 | AOC2;DPYS;UPB1;CNDP1;ALDH3A2;ALDH1A3;HADHA;ALDH3A1;ALDH6A1;[illegible] |
| Tryptophan metabolism | 16/42 | 7.77E-11 | 5.73E-10 | 11.728179 | 273.00499 | TPH2;TPH1;DDC;AADAT;HAAO;INMT;ACMSD;ALDH3A2;HADHA;AANAT;KYNU;[illegible] |
| Inflammatory mediator regulation of TRP channels | 24/100 | 1.19E-10 | 8.54E-10 | 6.0511918 | 138.27358 | PLA2G4F;PLA2G4E;CYP4A22;CYP4A11;ADCY4;ITPR2;TRPV3;PIK3CD;[illegible] |
| Glycosphingolipid biosynthesis | 16/45 | 2.62E-10 | 1.82E-09 | 10.513256 | 231.96867 | B4GALT1;ST8SIA1;SLC33A1;B3GALT4;B3GALT5;ABO;FUT9;B3GALT1;ST3GAL4;[illegible] |
| Cortisol synthesis and secretion | 19/65 | 2.68E-10 | 1.82E-09 | 7.8873817 | 173.84453 | HSD3B2;ADCY4;ITPR2;ITPR3;ADCY2;CACNA1D;ADCY1;CACNA1F;ADCY6;[illegible] |
| Calcium signaling pathway | 33/188 | 3.45E-10 | 2.29E-09 | 4.1000198 | 89.323692 | PDE1C;PDE1B;PDE1A;ADCY4;CACNA1A;ITPR2;ATP2A2;CACNA1D;ITPR3;[illegible] |
| Linoleic acid metabolism | 13/29 | 4.16E-10 | 2.68E-09 | 15.446455 | 333.66603 | PLA2G4F;PLA2G4E;PLA2G2C;ALOX15;PLA2G2A;PLA2G4A;PLA2G6;CYP2C19;[illegible] |
| Thyroid hormone synthesis | 20/74 | 4.30E-10 | 2.71E-09 | 7.0766528 | 152.61746 | GPX2;GPX3;GSR;ATP1A4;ADCY4;ITPR2;ATP1B3;ITPR3;ATP1A2;ADCY2;ADCY1;[illegible] |
| Choline metabolism in cancer | 23/99 | 5.81E-10 | 3.57E-09 | 5.7931893 | 123.20391 | DGKG;SLC22A4;DGKE;SLC44A3;PLA2G4F;DGKD;PCYT1A;DGKB;PLA2G4E;[illegible] |
| Peroxisome | 21/83 | 6.02E-10 | 3.61E-09 | 6.4755132 | 137.48059 | ABCD4;ABCD3;NOS2;ACSL1;IDH1;EPHX2;PIPOX;HSD17B4;SOD2;HMGCL;[illegible] |
| Amino sugar and nucleotide sugar metabolism | 16/48 | 7.90E-10 | 4.63E-09 | 9.5261307 | 199.65516 | GALT;MPI;UAP1;NPL;GMPPB;HK3;CYB5R1;UGDH;CYB5R4;GMDS;FPGT;[illegible] |
| cGMP-PKG signaling pathway | 30/166 | 1.06E-09 | 6.08E-09 | 4.2392966 | 87.605242 | NPR1;NPR2;ATP1A4;ADCY4;ITPR2;ATP2A2;CACNA1D;ITPR3;ATP2A1;ATP1A2;[illegible] |
| Vascular smooth muscle contraction | 26/132 | 2.04E-09 | 1.15E-08 | 4.7022124 | 94.084093 | NPR1;NPR2;CYP4A22;ADCY4;ITPR2;CACNA1D;ITPR3;ADCY2;ADCY1;[illegible] |
| cAMP signaling pathway | 34/212 | 2.19E-09 | 1.20E-08 | 3.6776994 | 73.332092 | HCN4;GRIA2;NPR1;ADCY4;ATP1A4;PIK3CD;ATP2A2;ADCY2;CACNA1D;[illegible] |
| Aldosterone synthesis and secretion | 22/98 | 2.74E-09 | 1.47E-08 | 5.5357086 | 109.14291 | KCNJ5;HSD3B2;NPR1;ATP1A4;ADCY4;ITPR2;ATP1B3;CACNA1D;ITPR3;[illegible] |
| PPAR signaling pathway | 19/74 | 3.02E-09 | 1.59E-08 | 6.593585 | 129.35371 | CPT1A;GK;ACSL1;CYP4A22;CYP4A11;UCP1;AQP7;CPT1B;CYP8B1;GK2;[illegible] |
| Chemical carcinogenesis | 20/82 | 3.09E-09 | 1.59E-08 | 6.1609323 | 120.7313 | ADH1B;GSTP1;UGT2B15;MGST1;CYP2C19;PTGS2;ADH7;CYP3A5;HSD11B1;[illegible] |
| Glycolysis / Gluconeogenesis | 18/68 | 4.62E-09 | 2.34E-08 | 6.8661027 | 131.78008 | PKLR;ADH1B;ADH7;ENO3;ALDH3A2;HK3;ALDH1A3;ALDH3A1;ALDH3B1;[illegible] |
| Cysteine and methionine metabolism | 15/47 | 5.31E-09 | 2.64E-08 | 8.9217809 | 169.98727 | AHCYL1;SDS;AGXT2;TAT;BHMT2;MTR;MTAP;BHMT;MAT2A;CTH;GCLM;BCAT1;[illegible] |
| Pyrimidine metabolism | 16/57 | 1.32E-08 | 6.43E-08 | 7.431499 | 134.8273 | CANT1;ENTPD3;ENTPD4;DPYS;ENTPD5;ENTPD6;CAD;UPB1;TYMP;UCKL1;[illegible] |
| Salivary secretion | 20/90 | 1.72E-08 | 8.24E-08 | 5.4545192 | 97.503464 | GUCY1A2;ATP1A4;ADCY4;ITPR2;ATP1B3;ITPR3;ATP1A2;ADCY2;ADCY1;BEST2;[illegible] |
| Mineral absorption | 15/51 | 1.86E-08 | 8.74E-08 | 7.9287985 | 141.11861 | SLC34A2;SLC6A19;SLC40A1;ATP1A4;CYBRD1;ATP1B3;SLC30A1;ATP1A2;[illegible] |
| Glycosaminoglycan biosynthesis | 15/53 | 3.32E-08 | 1.53E-07 | 7.5107007 | 129.33653 | HS3ST3B1;B4GALT1;CHST7;CSGALNACT2;HS6ST1;HS3ST2;NDST2;EXTL2;[illegible] |
| Platelet activation | 23/124 | 5.60E-08 | 2.54E-07 | 4.3534694 | 72.69263 | GUCY1A2;PLA2G4F;PLA2G4E;ADCY4;ITPR2;PIK3CD;PLA2G4A;ITPR3;ADCY2;[illegible] |
| Glutathione metabolism | 15/56 | 7.49E-08 | 3.33E-07 | 6.9600353 | 114.19487 | GGT5;GPX2;NAT8B;GGT7;GPX3;GSTP1;IDH1;GSR;MGST1;GSTA5;NAT8;[illegible] |
| Starch and sucrose metabolism | 12/36 | 1.06E-07 | 4.62E-07 | 9.491992 | 152.46337 | GYS2;HK3;AMY2A;AMY2B;TREH;SI;AGL;PGM2;PYGM;PYGL;PGM1;GANC |
| Pantothenate and CoA biosynthesis | 9/19 | 1.21E-07 | 5.22E-07 | 17.047006 | 271.45952 | PANK4;PANK2;PANK3;DPYS;DPYD;BCAT1;UPB1;COASY;BCAT2 |
| Carbohydrate digestion and absorption | 13/44 | 1.57E-07 | 6.62E-07 | 7.9660612 | 124.82755 | AMY2A;AMY2B;ATP1A4;SLC2A2;PIK3CD;ATP1A2;CACNA1D;ATP1B3;SLC2A5;[illegible] |
| Glucagon signaling pathway | 20/103 | 1.87E-07 | 7.77E-07 | 4.597036 | 71.222132 | PFKFB1;CPT1A;ITPR2;SLC2A2;ITPR3;PYGM;ADCY2;PYGL;CPT1B;ACACB;[illegible] |
| Long-term depression | 15/60 | 2.03E-07 | 8.30E-07 | 6.3400268 | 97.709634 | GUCY1A2;GRIA2;PLA2G4F;GRID2;PLA2G4E;PLA2G4A;ITPR2;CACNA1A;ITPR3;[illegible] |
| Cardiac muscle contraction | 17/78 | 2.72E-07 | 1.10E-06 | 5.3068575 | 80.222539 | COX8C;CACNA2D1;ATP1A4;ATP2A2;CACNA2D2;ATP1B3;ATP1A2;CACNA1D;[illegible] |
| Pyruvate metabolism | 12/39 | 2.87E-07 | 1.14E-06 | 8.4359915 | 127.07798 | ALDH3A2;PC;PKLR;ME1;ACOT12;ME3;ME2;PCK1;ACACB;ACACA;LDHAL6A;LDHAL6B |
| Sphingolipid metabolism | 13/47 | 3.68E-07 | 1.44E-06 | 7.262024 | 107.58272 | ARSA;ASAH2;CERK;SGMS1;GBA2;SGPP2;ACER2;NEU3;UGCG;ACER1;SPTLC2;ACER3;B4GALT6 |
| Dilated cardiomyopathy (DCM) | 18/91 | 5.70E-07 | 2.19E-06 | 4.6970989 | 67.535722 | CACNA2D1;ADCY4;ATP2A2;CACNA2D2;ADCY2;CACNA1D;CACNA2D4;[illegible] |
| Ovarian steroidogenesis | 13/49 | 6.26E-07 | 2.37E-06 | 6.8578546 | 97.960659 | PLA2G4F;HSD3B2;PLA2G4E;ADCY4;PLA2G4A;ADCY2;ADCY1;PTGS2;ADCY6;ADCY5;ALOX5;ACOT2;CYP1B1 |
| Phenylalanine metabolism | 8/17 | 6.70E-07 | 2.51E-06 | 16.820649 | 239.11935 | ALDH1A3;ALDH3A1;DDC;AOC2;TAT;ALDH3B1;GLYAT;HPD |
| GnRH signaling pathway | 18/93 | 7.98E-07 | 2.93E-06 | 4.5713595 | 64.186166 | PLA2G4F;PLA2G4E;ADCY4;PLA2G4A;ITPR2;ITPR3;ADCY2;CACNA1D;ADCY1;[illegible] |
| N-Glycan biosynthesis | 13/50 | 8.07E-07 | 2.93E-06 | 6.6721551 | 93.613072 | DPAGT1;B4GALT1;ST6GAL2;ALG14;ALG13;ALG11;MGAT4C;MAN2A2;MAN1A2;[illegible] |
| Histidine metabolism | 9/23 | 8.94E-07 | 3.20E-06 | 12.173867 | 169.55547 | ALDH3A2;ALDH1A3;ALDH3A1;HAL;AMDHD1;ALDH3B1;ASPA;CNDP1;FTCD |
| Glyoxylate and dicarboxylate metabolism | 10/30 | 1.24E-06 | 4.38E-06 | 9.475025 | 128.86696 | GCSH;MCEE;PCCA;GLDC;GLYCTK;CAT;AMT;ACO1;GLUL;HAO2 |
| Thyroid hormone signaling pathway | 20/116 | 1.36E-06 | 4.70E-06 | 3.9717878 | 53.636815 | PFKFB2;DIO1;DIO2;ATP1A4;DIO3;PIK3CD;ATP2A2;ATP1B3;SLC16A10;ATP1A2;[illegible] |
| Vitamin digestion and absorption | 9/24 | 1.37E-06 | 4.70E-06 | 11.361677 | 153.42437 | SLC5A6;BTD;TCN2;SLC23A1;MMACHC;PLB1;SLC19A1;PNLIP;SLC19A2 |
| Galactose metabolism | 10/31 | 1.75E-06 | 5.93E-06 | 9.0233576 | 119.6268 | HK3;GALT;B4GALT1;AKR1B10;SI;AKR1B1;PGM2;GALM;PGM1;GANC |
| Glycosylphosphatidylinositol (GPI)-anchor biosynt | 9/25 | 2.04E-06 | 6.83E-06 | 10.65101 | 139.5625 | PIGS;PGAP1;PIGO;PIGN;PIGZ;PIGM;PIGL;PIGV;GPLD1 |
| Dopaminergic synapse | 21/131 | 2.54E-06 | 8.41E-06 | 3.6405785 | 46.899857 | GRIA2;KCNJ5;DDC;KCNJ9;CACNA1A;ITPR2;CACNA1D;ITPR3;GRIN2B;[illegible] |
| Regulation of lipolysis in adipocytes | 13/55 | 2.61E-06 | 8.52E-06 | 5.8763002 | 75.55248 | NPR1;ADCY4;AQP7;PIK3CD;ADCY2;ABHD5;ADCY1;PTGS2;ADCY6;ADCY5;PTGS1;PIK3CA;PNPLA2 |
| Parathyroid hormone synthesis, secretion and acti | 18/106 | 5.69E-06 | 1.83E-05 | 3.8933672 | 47.021157 | SLC34A2;PDE4C;ADCY4;ITPR2;ITPR3;ADCY2;ADCY1;PLD1;ADCY6;PLD2;[illegible] |
| Steroid hormone biosynthesis | 13/60 | 7.34E-06 | 2.34E-05 | 5.2497761 | 62.063311 | DHRS11;HSD3B2;UGT2B15;AKR1C4;CYP7B1;CYP3A5;HSD11B1;SULT1E1;[illegible] |
| Primary bile acid biosynthesis | 7/17 | 1.03E-05 | 3.23E-05 | 13.232371 | 151.98408 | CH25H;ACOX2;HSD17B4;AKR1C4;CYP7B1;CYP46A1;CYP8B1 |
| Pentose phosphate pathway | 9/30 | 1.14E-05 | 3.53E-05 | 8.112917 | 92.371039 | TKTL2;RPIA;H6PD;GLYCTK;PGM2;FBP1;PGM1;RBKS;FBP2 |
| Apelin signaling pathway | 20/137 | 1.84E-05 | 5.66E-05 | 3.2552804 | 35.487728 | NOS2;UCP1;ADCY4;ITPR2;ITPR3;ADCY2;ADCY1;PIK3R6;RYR3;PIK3CG;[illegible] |
| alpha-Linolenic acid metabolism | 8/25 | 2.08E-05 | 6.23E-05 | 8.9012961 | 95.981062 | PLA2G4F;PLA2G4E;PLA2G2C;ACOX1;PLA2G2A;PLA2G4A;PLA2G6;PLB1 |
| Fatty acid biosynthesis | 6/13 | 2.08E-05 | 6.23E-05 | 16.189339 | 174.5631 | OLAH;ACSL1;ACSBG1;ACACB;ACSBG2;ACACA |
| Synaptic vesicle cycle | 14/78 | 3.22E-05 | 9.56E-05 | 4.1523007 | 42.945425 | ATP6V1G2;SLC32A1;SLC6A12;CACNA1A;TCIRG1;SLC1A7;SLC6A4;SLC6A5;[illegible] |
| Neuroactive ligand-receptor interaction | 35/338 | 4.87E-05 | 1.43E-04 | 2.2115255 | 21.960659 | GABRB3;GABRB2;GRIA2;CHRNA3;GRIK5;GRIK1;GRIK2;GABRR2;CHRND;[illegible] |
| Arrhythmogenic right ventricular cardiomyopathy ( | 13/72 | 5.74E-05 | 1.66E-04 | 4.1793757 | 40.81218 | CACNA2D1;ATP2A2;CACNA2D2;CACNA1D;CACNA2D4;CACNA1F;CACNB1;[illegible] |
| Type II diabetes mellitus | 10/46 | 8.05E-05 | 2.31E-04 | 5.2594628 | 49.58433 | HK3;KCNJ11;PKLR;PIK3CA;ABCC8;CACNA1A;SLC2A2;PIK3CD;CACNA1D;CACNA1E |
| Hypertrophic cardiomyopathy (HCM) | 14/85 | 8.61E-05 | 2.44E-04 | 3.7415345 | 35.02029 | ACE;CACNA2D1;ATP2A2;CACNA2D2;CACNA1D;CACNA2D4;CACNA1F;[illegible] |
| Nicotinate and nicotinamide metabolism | 8/30 | 8.96E-05 | 2.51E-04 | 6.8764615 | 64.08731 | BST1;NNMT;NMNAT1;NT5C1A;NT5C1B;AOX1;NADSYN1;NADK |
| Ether lipid metabolism | 10/47 | 9.76E-05 | 2.69E-04 | 5.1170451 | 47.252374 | PLA2G4F;PLA2G4E;PLA2G2C;PLA2G2A;GDPD1;PLA2G4A;PLA2G6;PLD1;PLB1;PLD2 |
| Mannose type O-glycan biosynthesis | 7/23 | 9.92E-05 | 2.69E-04 | 8.267617 | 76.216912 | B4GALT1;FUT9;B3GAT2;POMT1;B3GAT1;POMGNT1;ST3GAL3 |
| Proximal tubule bicarbonate reclamation | 7/23 | 9.92E-05 | 2.69E-04 | 8.267617 | 76.216912 | SLC9A3;ATP1A4;ATP1A2;ATP1B3;PCK1;GLS;AQP1 |
| Selenocompound metabolism | 6/17 | 1.26E-04 | 3.38E-04 | 10.300136 | 92.511287 | TXNRD1;CTH;MTR;SCLY;INMT;PAPSS2 |
| Amphetamine addiction | 12/68 | 1.38E-04 | 3.67E-04 | 4.0611326 | 36.094582 | GRIA2;DDC;GRIN3B;TH;CACNA1D;SLC18A1;GRIN2B;SLC18A2;GRIA3;[illegible] |
| Fat digestion and absorption | 9/41 | 1.68E-04 | 4.44E-04 | 5.3210142 | 46.23314 | PNLIPRP1;LIPF;ABCA1;ABCG8;DGAT1;MOGAT2;PLA2G2C;PLA2G2A;PNLIP |
| Folate biosynthesis | 7/26 | 2.33E-04 | 6.07E-04 | 6.961103 | 58.228253 | TPH2;TPH1;AKR1B10;TH;FPGS;MOCOS;AKR1B1 |
| Glycosaminoglycan degradation | 6/19 | 2.53E-04 | 6.52E-04 | 8.7145809 | 72.189054 | GALNS;HGSNAT;HPSE2;IDUA;HYAL4;HPSE |
| Cushing syndrome | 19/155 | 3.16E-04 | 8.08E-04 | 2.6551159 | 21.398372 | HSD3B2;ADCY4;ITPR2;CACNA1D;ITPR3;ADCY2;ADCY1;CACNA1F;ADCY6;[illegible] |
| One carbon pool by folate | 6/20 | 3.45E-04 | 8.74E-04 | 8.0916844 | 64.49493 | ALDH1L1;AMT;MTHFR;MTR;ALDH1L2;FTCD |
| Butanoate metabolism | 7/28 | 3.84E-04 | 9.61E-04 | 6.2974768 | 49.53313 | HMGCL;HADHA;ACSM2A;ACSM5;ACSM4;HMGCLL1;ACSM2B |
| Arginine biosynthesis | 6/21 | 4.63E-04 | 0.0011489 | 7.5518408 | 57.979167 | NOS2;ARG1;NOS1;GLUL;GLS;OTC |
| Endocrine and other factor-regulated calcium reab | 9/48 | 5.87E-04 | 0.0014427 | 4.3643482 | 32.471789 | PLCB3;PLCB4;TRPV5;ATP1A4;ATP1A2;ATP1B3;PLCB1;PLCB2;ADCY6 |
| Other types of O-glycan biosynthesis | 6/22 | 6.10E-04 | 0.0014842 | 7.0794776 | 52.405333 | POFUT2;B4GALT1;ST6GAL2;MFNG;POMT1;ST3GAL3 |
| Arginine and proline metabolism | 9/49 | 6.87E-04 | 0.0016422 | 4.255015 | 30.98726 | ALDH3A2;DAO;CKM;NOS2;ARG1;SAT2;NOS1;PRODH;CNDP1 |
| Cocaine addiction | 9/49 | 6.87E-04 | 0.0016422 | 4.255015 | 30.98726 | GRIA2;DDC;GRIN3B;TH;SLC18A1;GRIN2B;SLC18A2;ADCY5;GRIN1 |
| Mucin type O-glycan biosynthesis | 7/31 | 7.46E-04 | 0.0017635 | 5.5094207 | 39.668854 | GALNT6;GALNT13;GALNT3;B3GNT6;GCNT4;GALNTL6;GALNT8 |
| Ferroptosis | 8/40 | 7.52E-04 | 0.0017635 | 4.7250748 | 33.987234 | TF;ACSL1;ALOX15;SLC40A1;CYBB;SAT2;GCLM;CP |
| Sphingolipid signaling pathway | 15/119 | 9.77E-04 | 0.0022704 | 2.734737 | 18.955146 | ASAH2;SGMS1;PTEN;PIK3CD;PLD1;SGPP2;PLD2;ACER2;ACER1;PLCB3;[illegible] |
| Porphyrin and chlorophyll metabolism | 8/42 | 0.0010553 | 0.0024309 | 4.44666 | 30.477194 | ALAD;ALAS2;ALAS1;FECH;HEPH;UGT2B15;PPOX;CP |
| AMPK signaling pathway | 15/120 | 0.0010653 | 0.0024323 | 2.7085485 | 18.538616 | PFKFB2;PFKFB1;CPT1A;PFKFB3;PIK3CD;CPT1B;ACACB;ACACA;GYS2;PIK3CA;[illegible] |
| Nitrogen metabolism | 5/17 | 0.0012145 | 0.0027487 | 7.8599238 | 52.766857 | CA1;CA3;CA5A;GLUL;CA13 |
| Pentose and glucuronate interconversions | 7/34 | 0.001338 | 0.0030018 | 4.8964881 | 32.397967 | UGDH;AKR1B10;UGT2B15;SORD;AKR1B1;CRYL1;XYLB |
| Estrogen signaling pathway | 16/137 | 0.0015319 | 0.0034045 | 2.5074795 | 16.251559 | KCNJ5;KCNJ9;ADCY4;ITPR2;PIK3CD;ITPR3;ADCY2;ADCY1;ADCY6;ADCY5;[illegible] |
| Gap junction | 12/88 | 0.0015439 | 0.0034045 | 2.9892524 | 19.350761 | GUCY1A2;PLCB3;PLCB4;ADCY4;ITPR2;ITPR3;ADCY2;ADCY1;PLCB1;PLCB2;[illegible] |
| Alanine, aspartate and glutamate metabolism | 7/35 | 0.0016005 | 0.0034887 | 4.7213645 | 30.393363 | ADSL;AGXT2;CAD;GLUL;ASPA;NIT2;GLS |
| Taurine and hypotaurine metabolism | 4/11 | 0.0016091 | 0.0034887 | 10.771457 | 69.28272 | GGT5;GGT7;GGT1;CSAD |
| Long-term potentiation | 10/67 | 0.0018764 | 0.0040042 | 3.3180854 | 20.832258 | GRIA2;PLCB3;PLCB4;ITPR2;ITPR3;ADCY1;PLCB1;PLCB2;GRIN2B;GRIN1 |
| Protein digestion and absorption | 12/90 | 0.0018779 | 0.0040042 | 2.9122969 | 18.282213 | KCNK5;SLC9A3;SLC15A1;SLC6A19;SLC7A7;KCNQ1;KCNJ13;ATP1A4;ATP1B3;ATP1A2;SLC16A10;SLC38A2 |
| Fatty acid elongation | 6/27 | 0.0019486 | 0.0041209 | 5.3924662 | 33.652398 | HADHB;HADHA;ACAA2;THEM5;ACOT2;THEM4 |
| Fc gamma R-mediated phagocytosis | 12/91 | 0.0020659 | 0.0043333 | 2.8752803 | 17.775529 | PLA2G4F;PIK3CA;PLA2G4E;INPP5D;INPPL1;PIK3CD;PLA2G4A;PIP5K1A;PLCG1;PLD1;PLA2G6;PLD2 |
| Aldosterone-regulated sodium reabsorption | 7/37 | 0.0022444 | 0.0046698 | 4.4061421 | 26.87444 | PIK3CA;SCNN1A;ATP1A4;PIK3CD;ATP1A2;ATP1B3;KCNJ1 |
| Alzheimer disease | 18/171 | 0.0026163 | 0.0054 | 2.2316213 | 13.269229 | COX8C;NDUFA10;ITPR2;ATP2A2;CACNA1D;ITPR3;ATP2A1;CACNA1F;GRIN2B;[illegible] |
| Insulin resistance | 13/108 | 0.0031303 | 0.0064097 | 2.5906761 | 14.939434 | CPT1A;PTEN;SLC2A2;PIK3CD;PYGM;PYGL;CPT1B;ACACB;GYS2;PIK3CA;[illegible] |
| Citrate cycle (TCA cycle) | 6/30 | 0.0034351 | 0.0069783 | 4.7176617 | 26.766704 | PC;SUCLA2;IDH1;SUCLG2;ACO1;PCK1 |
| Fc epsilon RI signaling pathway | 9/68 | 0.007021 | 0.0141517 | 2.8818634 | 14.290727 | PLA2G4F;PIK3CA;PLA2G4E;ALOX5;INPP5D;ALOX5AP;PIK3CD;PLA2G4A;PLCG1 |
| Insulin signaling pathway | 14/137 | 0.0095534 | 0.0191069 | 2.153811 | 10.017062 | PKLR;INPPL1;PIK3CD;PYGM;PYGL;ACACB;ACACA;GYS2;HK3;PIK3CA;INPP5K;[illegible] |
| Other glycan degradation | 4/18 | 0.0112477 | 0.0223223 | 5.3837424 | 24.160052 | NEU3;MANBA;MAN2C1;GBA2 |
| AGE-RAGE signaling pathway in diabetic complicat | 11/100 | 0.0121599 | 0.0239485 | 2.3359551 | 10.300657 | PLCB3;PLCB4;PIK3CA;CYBB;PIK3CD;PLCG1;PLCB1;PLCD3;PLCB2;NOX1;[illegible] |
| Cholesterol metabolism | 7/50 | 0.0123517 | 0.024142 | 3.0719448 | 13.498003 | ABCA1;ABCG8;NPC1;SOAT1;SOAT2;LCAT;ABCB11 |
| Steroid biosynthesis | 4/19 | 0.0136876 | 0.026552 | 5.0245614 | 21.561709 | CYP27B1;SOAT1;SOAT2;CYP2R1 |
| Relaxin signaling pathway | 13/130 | 0.0144788 | 0.027877 | 2.1010911 | 8.8982728 | NOS2;ADCY4;PIK3CD;ADCY2;ADCY1;ADCY6;ADCY5;PLCB3;PLCB4;PIK3CA;[illegible] |
| Lysosome | 12/123 | 0.0218617 | 0.0417802 | 2.0429078 | 7.8100721 | MFSD8;GALNS;ARSA;HGSNAT;MANBA;NPC1;ENTPD4;IDUA;SLC11A1;CTNS;[illegible] |
| Thermogenesis | 19/231 | 0.025636 | 0.048633 | 1.6964157 | 6.2152565 | KLB;CPT1A;COX8C;ACSL1;NPR1;NDUFA10;ADCY4;UCP1;ADCY2;ADCY1;[illegible] |
| VEGF signaling pathway | 7/59 | 0.0286164 | 0.0538908 | 2.5390553 | 9.0232305 | PLA2G4F;PIK3CA;PLA2G4E;PIK3CD;PLA2G4A;PLCG1;PTGS2 |
| Melanogenesis | 10/101 | 0.031553 | 0.0589903 | 2.0746287 | 7.1700986 | PLCB3;PLCB4;DCT;ADCY4;ADCY2;ADCY1;PLCB1;PLCB2;ADCY6;ADCY5 |
| Vibrio cholerae infection | 6/50 | 0.0393139 | 0.0729711 | 2.5705563 | 8.3187777 | ATP6V1G2;KCNQ1;TCIRG1;PLCG1;ATP6V1E2;CFTR |
| Collecting duct acid secretion | 4/27 | 0.0451176 | 0.0826724 | 3.2755062 | 10.149097 | ATP4B;ATP6V1G2;TCIRG1;ATP6V1E2 |
| Central carbon metabolism in cancer | 7/65 | 0.0451814 | 0.0826724 | 2.2756732 | 7.0479163 | HK3;PIK3CA;IDH1;PTEN;SLC2A2;PIK3CD;GLS |
| Phototransduction | 4/28 | 0.0506087 | 0.091951 | 3.1388613 | 9.3652071 | GUCY2D;CNGA1;PDE6B;CNGB1 |
| Amoebiasis | 9/96 | 0.0536423 | 0.0967811 | 1.9514764 | 5.7088844 | PLCB3;PLCB4;PIK3CA;NOS2;ARG1;PIK3CD;ADCY1;PLCB1;PLCB2 |
| Adipocytokine signaling pathway | 7/69 | 0.0590686 | 0.1058312 | 2.1284057 | 6.0213802 | CPT1A;ACSL1;ACSBG1;PCK1;CPT1B;ACSBG2;ACACB |
| HIF-1 signaling pathway | 9/100 | 0.0661173 | 0.1176433 | 1.8653024 | 5.066766 | HK3;TF;PFKFB3;PIK3CA;NOS2;CYBB;PIK3CD;PLCG1;ENO3 |
| Longevity regulating pathway | 9/102 | 0.0729859 | 0.1289751 | 1.8249952 | 4.7769049 | PIK3CA;CAT;ADCY4;PIK3CD;ADCY2;ADCY1;SOD2;ADCY6;ADCY5 |
| Chagas disease (American trypanosomiasis) | 9/103 | 0.0765794 | 0.1335729 | 1.8054848 | 4.6390619 | PLCB3;ACE;PLCB4;PIK3CA;NOS2;PIK3CD;ADCY1;PLCB1;PLCB2 |
| Lysine degradation | 6/59 | 0.0766232 | 0.1335729 | 2.1330329 | 5.4794521 | ALDH3A2;HADHA;AADAT;PIPOX;BBOX1;AASS |
| Synthesis and degradation of ketone bodies | 2/10 | 0.0877442 | 0.1519329 | 4.7029237 | 11.443763 | HMGCL;HMGCLL1 |
| Chemokine signaling pathway | 14/190 | 0.1017436 | 0.1749989 | 1.5009916 | 3.4302156 | ADCY4;PIK3CD;ADCY2;ADCY1;PIK3R6;PIK3CG;ADCY6;PIK3R5;ADCY5;[illegible] |
| Ubiquinone and other terpenoid-quinone biosynthe | 2/11 | 0.1037631 | 0.1772905 | 4.1801564 | 9.4707517 | TAT;HPD |
| Amyotrophic lateral sclerosis (ALS) | 5/51 | 0.1140019 | 0.1935032 | 2.0467413 | 4.4445807 | GRIA2;CAT;NOS1;GRIN2B;GRIN1 |
| African trypanosomiasis | 4/37 | 0.1150738 | 0.1940461 | 2.2817249 | 4.933503 | PLCB3;PLCB4;PLCB1;PLCB2 |
| Ascorbate and aldarate metabolism | 3/27 | 0.1539202 | 0.2562027 | 2.3518105 | 4.4009921 | ALDH3A2;UGDH;UGT2B15 |
| Biosynthesis of unsaturated fatty acids | 3/27 | 0.1539202 | 0.2562027 | 2.3518105 | 4.4009921 | ACOX1;ACOT2;HSD17B4 |
| Rap1 signaling pathway | 14/206 | 0.160846 | 0.2660146 | 1.3747388 | 2.5120708 | ADCY4;PIK3CD;ADCY2;ADCY1;GRIN2B;ADCY6;ADCY5;GRIN1;PLCB3;PLCB4;[illegible] |
| Thiamine metabolism | 2/15 | 0.1738821 | 0.2857425 | 2.8933445 | 5.0615528 | AK2;AK7 |
| Caffeine metabolism | 1/5 | 0.2284765 | 0.36613 | 4.6992574 | 6.9376172 | CYP1A2 |
| D-Glutamine and D-glutamate metabolism | 1/5 | 0.2284765 | 0.36613 | 4.6992574 | 6.9376172 | GLS |
| Neomycin, kanamycin and gentamicin biosynthesis | 1/5 | 0.2284765 | 0.36613 | 4.6992574 | 6.9376172 | HK3 |
| Phenylalanine, tyrosine and tryptophan biosynthes | 1/5 | 0.2284765 | 0.36613 | 4.6992574 | 6.9376172 | TAT |
| Progesterone-mediated oocyte maturation | 7/99 | 0.234219 | 0.3730155 | 1.4320869 | 2.0786721 | PIK3CA;ADCY4;PIK3CD;ADCY2;ADCY1;ADCY6;ADCY5 |
| MAPK signaling pathway | 18/295 | 0.2377945 | 0.3763864 | 1.2245138 | 1.7588286 | PLA2G4F;PLA2G4E;CACNA2D1;CACNA2D2;CACNA1A;PLA2G4A;CACNA1D;[illegible] |
| Human cytomegalovirus infection | 14/225 | 0.2492583 | 0.3921258 | 1.2496827 | 1.7361413 | ADCY4;ITPR2;PIK3CD;ADCY2;ITPR3;ADCY1;PTGS2;ADCY6;ADCY5;PLCB3;[illegible] |
| Phosphonate and phosphinate metabolism | 1/6 | 0.2674867 | 0.4157323 | 3.7592079 | 4.9572128 | PCYT1A |
| Vitamin B6 metabolism | 1/6 | 0.2674867 | 0.4157323 | 3.7592079 | 4.9572128 | AOX1 |
| Fluid shear stress and atherosclerosis | 9/139 | 0.2705081 | 0.4169555 | 1.303017 | 1.703634 | GSTA5;PIK3CA;GSTA4;TRPV4;GSTP1;MGST1;PIK3CD;NOX1;GSTM5 |
| Huntington disease | 12/193 | 0.2715059 | 0.4169555 | 1.2481874 | 1.6273509 | COX8C;PLCB3;PLCB4;NDUFA10;UCP1;NDUFS2;NDUFS1;PLCB1;SOD2;GRIN2B;PLCB2;GRIN1 |
| Prolactin signaling pathway | 5/70 | 0.2793191 | 0.4264162 | 1.4470102 | 1.8455173 | GALT;PIK3CA;TH;SLC2A2;PIK3CD |
| Oocyte meiosis | 8/125 | 0.2979975 | 0.452255 | 1.2865336 | 1.5575678 | PLCZ1;ADCY4;ITPR2;ITPR3;ADCY2;ADCY1;ADCY6;ADCY5 |
| Necroptosis | 10/162 | 0.3039124 | 0.4585344 | 1.2380383 | 1.4745233 | PLA2G4F;PLA2G4E;ALOX15;CYBB;PLA2G4A;TRPM7;PYGM;PGAM5;PYGL;GLUL |
| Renin-angiotensin system | 2/23 | 0.325339 | 0.4880085 | 1.7903629 | 2.0103762 | ACE;LNPEP |
| Small cell lung cancer | 6/93 | 0.3302082 | 0.4924493 | 1.2971007 | 1.4372288 | PIK3CA;NOS2;PTEN;PIK3CD;PTGS2;FHIT |
| Hepatocellular carcinoma | 10/168 | 0.343962 | 0.5100126 | 1.1906448 | 1.2706849 | GSTA5;PIK3CA;GSTA4;GSTP1;TXNRD1;MGST1;PTEN;PIK3CD;PLCG1;GSTM5 |
| Oxidative phosphorylation | 8/133 | 0.358831 | 0.5290194 | 1.2036849 | 1.2336613 | ATP4B;COX8C;ATP6V1G2;NDUFA10;NDUFS2;NDUFS1;TCIRG1;ATP6V1E2 |
| Sulfur metabolism | 1/9 | 0.3730865 | 0.5469109 | 2.3491337 | 2.3161167 | PAPSS2 |
| Maturity onset diabetes of the young | 2/26 | 0.3811467 | 0.5555698 | 1.5663198 | 1.5108264 | PKLR;SLC2A2 |
| Ras signaling pathway | 13/232 | 0.3918474 | 0.5679586 | 1.1164338 | 1.0459675 | PLA2G4F;PLA2G4E;PLA2G2C;PLA2G2A;PIK3CD;PLA2G4A;PLA2G6;PLD1;[illegible] |
| Parkinson disease | 8/142 | 0.4285344 | 0.6158635 | 1.1223047 | 0.9510234 | COX8C;TH;NDUFA10;NDUFS2;NDUFS1;SLC18A1;SLC18A2;ADCY5 |
| Non-small cell lung cancer | 4/66 | 0.4296722 | 0.6158635 | 1.2126085 | 1.0243301 | PIK3CA;PIK3CD;PLCG1;FHIT |
| Epithelial cell signaling in Helicobacter pylori infec | 4/68 | 0.4524024 | 0.6448608 | 1.1745904 | 0.9316655 | ATP6V1G2;TCIRG1;PLCG1;ATP6V1E2 |
| Kaposi sarcoma-associated herpesvirus infection | 10/186 | 0.4670125 | 0.6620287 | 1.0678526 | 0.8130621 | PIK3CA;ITPR2;PIK3CD;ITPR3;PIK3C3;PLCG1;PTGS2;PIK3R6;PIK3CG;PIK3R5 |
| Non-alcoholic fatty liver disease (NAFLD) | 8/149 | 0.482316 | 0.6799865 | 1.0661915 | 0.7774197 | COX8C;PIK3CA;PKLR;NDUFA10;PIK3CD;NDUFS2;NDUFS1;CYP2E1 |
| B cell receptor signaling pathway | 4/71 | 0.4859118 | 0.6813328 | 1.1218189 | 0.8096484 | PIK3CA;INPP5D;INPPL1;PIK3CD |
| Glioma | 4/75 | 0.5292271 | 0.7380573 | 1.0583941 | 0.6734959 | PIK3CA;PTEN;PIK3CD;PLCG1 |
| Alcoholism | 9/180 | 0.5616767 | 0.7777953 | 0.9884442 | 0.5701631 | GRIN3B;DDC;TH;SLC29A1;GRIN2B;SLC18A1;SLC18A2;ADCY5;GRIN1 |
| Cellular senescence | 8/160 | 0.5637508 | 0.7777953 | 0.9884557 | 0.5665263 | PIK3CA;TRPV4;PTEN;ITPR2;PIK3CD;CACNA1D;ITPR3;TRPM7 |
| Endometrial cancer | 3/58 | 0.5672254 | 0.7784263 | 1.0245671 | 0.5809281 | PIK3CA;PTEN;PIK3CD |
| C-type lectin receptor signaling pathway | 5/104 | 0.6093764 | 0.8318471 | 0.9483503 | 0.4697361 | PIK3CA;ITPR2;PIK3CD;ITPR3;PTGS2 |
| Phagosome | 7/152 | 0.6530763 | 0.8849045 | 0.906086 | 0.3860481 | ATP6V1G2;CYBB;PIK3C3;TCIRG1;NOS1;MPO;ATP6V1E2 |
| Acute myeloid leukemia | 3/66 | 0.6551037 | 0.8849045 | 0.8940854 | 0.3781639 | PIK3CA;PIK3CD;MPO |
| Vasopressin-regulated water reabsorption | 2/44 | 0.6592344 | 0.8854498 | 0.8941904 | 0.3725877 | AQP3;ADCY6 |
| FoxO signaling pathway | 6/132 | 0.6623713 | 0.8854498 | 0.8937692 | 0.3681694 | PIK3CA;CAT;PTEN;PIK3CD;PCK1;SOD2 |
| Toxoplasmosis | 5/113 | 0.6817937 | 0.9036417 | 0.8689069 | 0.3328158 | NOS2;ALOX5;PIK3R6;PIK3CG;PIK3R5 |
| NOD-like receptor signaling pathway | 8/178 | 0.682985 | 0.9036417 | 0.8829511 | 0.3366537 | PLCB3;PLCB4;CYBB;ITPR2;ITPR3;TRPM7;PLCB1;PLCB2 |
| Melanoma | 3/72 | 0.7120949 | 0.9373495 | 0.8160801 | 0.2770951 | PIK3CA;PTEN;PIK3CD |
| Prostate cancer | 4/97 | 0.7290003 | 0.9504284 | 0.8070816 | 0.2551032 | PIK3CA;GSTP1;PTEN;PIK3CD |
| Leishmaniasis | 3/74 | 0.7293986 | 0.9504284 | 0.793008 | 0.2502218 | NOS2;CYBB;PTGS2 |
| Pancreatic cancer | 3/75 | 0.737739 | 0.9564657 | 0.7819527 | 0.2378427 | PIK3CA;PIK3CD;PLD1 |
| Asthma | 1/31 | 0.7999681 | 0.9999955 | 0.6257096 | 0.139648 | EPX |
| ErbB signaling pathway | 3/85 | 0.8102727 | 0.9999955 | 0.6862297 | 0.144372 | PIK3CA;PIK3CD;PLCG1 |
| TNF signaling pathway | 4/110 | 0.8131031 | 0.9999955 | 0.7076127 | 0.1464032 | PIK3CA;PIK3CD;PGAM5;PTGS2 |
| Axon guidance | 7/181 | 0.8147695 | 0.9999955 | 0.7539096 | 0.1544384 | TRPC5;PIK3CA;DPYSL5;TRPC4;DPYSL2;PIK3CD;PLCG1 |
| Leukocyte transendothelial migration | 4/112 | 0.8239301 | 0.9999955 | 0.6944352 | 0.134491 | PIK3CA;CYBB;PIK3CD;PLCG1 |
| Rheumatoid arthritis | 3/91 | 0.8450826 | 0.9999955 | 0.6392384 | 0.1075972 | ATP6V1G2;TCIRG1;ATP6V1E2 |
| Pathways in cancer | 22/530 | 0.857404 | 0.9999955 | 0.8092601 | 0.1245015 | NOS2;GSTP1;TXNRD1;ADCY4;PTEN;MGST1;PIK3CD;ADCY2;ADCY1;PTGS2;[illegible] |
| Human T-cell leukemia virus 1 infection | 8/219 | 0.8692623 | 0.9999955 | 0.7098326 | 0.0994549 | PIK3CA;ADCY4;PTEN;PIK3CD;ADCY2;ADCY1;ADCY6;ADCY5 |
| Renal cell carcinoma | 2/69 | 0.8700598 | 0.9999955 | 0.5597976 | 0.0779201 | PIK3CA;PIK3CD |
| Bladder cancer | 1/41 | 0.8810371 | 0.9999955 | 0.4690347 | 0.0594058 | TYMP |
| Proteoglycans in cancer | 7/201 | 0.8871699 | 0.9999955 | 0.6754682 | 0.0808663 | HPSE2;PIK3CA;ITPR2;PIK3CD;ITPR3;PLCG1;HPSE |
| mTOR signaling pathway | 5/152 | 0.8879792 | 0.9999955 | 0.637062 | 0.0756874 | ATP6V1G2;PIK3CA;PTEN;PIK3CD;ATP6V1E2 |
| T cell receptor signaling pathway | 3/101 | 0.8908315 | 0.9999955 | 0.5737063 | 0.0663205 | PIK3CA;PIK3CD;PLCG1 |
| Autophagy | 4/128 | 0.8930544 | 0.9999955 | 0.6043182 | 0.0683531 | PIK3CA;PTEN;PIK3CD;PIK3C3 |
| Bacterial invasion of epithelial cells | 2/74 | 0.8941084 | 0.9999955 | 0.5207852 | 0.0582906 | PIK3CA;PIK3CD |
| Chronic myeloid leukemia | 2/76 | 0.9025146 | 0.9999955 | 0.5066563 | 0.0519679 | PIK3CA;PIK3CD |
| RNA degradation | 2/79 | 0.9139624 | 0.9999955 | 0.4868392 | 0.0437989 | NUDT16;ENO3 |
| Notch signaling pathway | 1/48 | 0.9173273 | 0.9999955 | 0.399031 | 0.0344328 | MFNG |
| Apoptosis | 4/143 | 0.9349627 | 0.9999955 | 0.5386753 | 0.0362252 | PIK3CA;ITPR2;PIK3CD;ITPR3 |
| Colorectal cancer | 2/86 | 0.9359509 | 0.9999955 | 0.4461041 | 0.0295287 | PIK3CA;PIK3CD |

| | | | | | | |
|---|---|---|---|---|---|---|
| Autoimmune thyroid disease | 1/53 | 0.9362565 | 0.9999955 | 0.3605674 | 0.0237491 | TPO |
| Neurotrophin signaling pathway | 3/119 | 0.9437043 | 0.9999955 | 0.4842211 | 0.0280569 | PIK3CA;PIK3CD;PLCG1 |
| NF-kappa B signaling pathway | 2/95 | 0.9564672 | 0.9999955 | 0.4027409 | 0.0179255 | PLCG1;PTGS2 |
| Osteoclast differentiation | 3/127 | 0.9585261 | 0.9999955 | 0.452789 | 0.0191795 | PIK3CA;PIK3CD;NOX1 |
| Wnt signaling pathway | 4/158 | 0.9613999 | 0.9999955 | 0.48582 | 0.0191242 | PLCB3;PLCB4;PLCB1;PLCB2 |
| Natural killer cell mediated cytotoxicity | 3/131 | 0.9644824 | 0.9999955 | 0.4385463 | 0.0158595 | PIK3CA;PIK3CD;PLCG1 |
| Mitophagy | 1/65 | 0.9658571 | 0.9999955 | 0.2927754 | 0.0101708 | PGAM5 |
| Protein processing in endoplasmic reticulum | 4/165 | 0.9699611 | 0.9999955 | 0.4645247 | 0.0141677 | EDEM3;MAN1A2;EDEM1;MAN1A1 |
| Toll-like receptor signaling pathway | 2/104 | 0.9706029 | 0.9999955 | 0.3670301 | 0.0109514 | PIK3CA;PIK3CD |
| p53 signaling pathway | 1/72 | 0.9762827 | 0.9999955 | 0.2638126 | 0.0063323 | PTEN |
| Pertussis | 1/76 | 0.9807415 | 0.9999955 | 0.2496898 | 0.0048556 | NOS2 |
| Breast cancer | 3/147 | 0.9811508 | 0.9999955 | 0.3894883 | 0.0074116 | PIK3CA;PTEN;PIK3CD |
| Tuberculosis | 4/179 | 0.9820315 | 0.9999955 | 0.427045 | 0.0077431 | CYP27B1;NOS2;PIK3C3;TCIRG1 |
| Gastric cancer | 3/149 | 0.98261 | 0.9999955 | 0.384112 | 0.0067385 | ABCB1;PIK3CA;PIK3CD |
| Human immunodeficiency virus 1 infection | 5/212 | 0.9841591 | 0.9999955 | 0.4509657 | 0.0072009 | PIK3CA;ITPR2;PIK3CD;ITPR3;PLCG1 |
| Regulation of actin cytoskeleton | 5/214 | 0.985238 | 0.9999955 | 0.4466027 | 0.0066419 | PIK3CA;PIP4K2A;PIP5K1A;PIK3CD;PIP4K2B |
| MicroRNAs in cancer | 8/299 | 0.9856468 | 0.9999955 | 0.5124969 | 0.0074093 | SLC45A3;ABCB1;PIK3CA;PTEN;CYP1B1;PLCG1;PTGS2;GLS |
| ECM-receptor interaction | 1/82 | 0.9859094 | 0.9999955 | 0.2311209 | 0.0032798 | SV2C |
| Salmonella infection | 1/86 | 0.9885594 | 0.9999955 | 0.220198 | 0.0025337 | NOS2 |
| JAK-STAT signaling pathway | 3/162 | 0.9897683 | 0.9999955 | 0.3524633 | 0.0036249 | PIK3CA;PIK3CD;AOX1 |
| Hepatitis B | 3/163 | 0.9901818 | 0.9999955 | 0.3502418 | 0.0034557 | PIK3CA;SLC10A1;PIK3CD |
| Th1 and Th2 cell differentiation | 1/92 | 0.9916305 | 0.9999955 | 0.2056142 | 0.0017281 | PLCG1 |
| IL-17 signaling pathway | 1/93 | 0.9920554 | 0.9999955 | 0.2033685 | 0.0016221 | PTGS2 |
| Measles | 2/138 | 0.9936222 | 0.9999955 | 0.274777 | 0.0017581 | PIK3CA;PIK3CD |
| PI3K-Akt signaling pathway | 9/354 | 0.9938963 | 0.9999955 | 0.4853944 | 0.0029718 | GYS2;PIK3CA;PTEN;THEM4;PIK3CD;PCK1;PIK3R6;PIK3CG;PIK3R5 |
| Signaling pathways regulating pluripotency of stem | 2/139 | 0.9939077 | 0.9999955 | 0.2727569 | 0.0016668 | PIK3CA;PIK3CD |
| Human papillomavirus infection | 8/330 | 0.9946265 | 0.9999955 | 0.4623892 | 0.0024913 | ATP6V1G2;PIK3CA;MFNG;PTEN;PIK3CD;TCIRG1;PTGS2;ATP6V1E2 |
| Endocytosis | 5/244 | 0.9950376 | 0.9999955 | 0.3899199 | 0.0019397 | PIP5K1A;FOLR3;PLD1;FOLR1;PLD2 |
| Th17 cell differentiation | 1/107 | 0.9961698 | 0.9999955 | 0.1763777 | 6.77E-04 | PLCG1 |
| Transcriptional misregulation in cancer | 3/186 | 0.996257 | 0.9999955 | 0.3058483 | 0.0011469 | SLC45A3;MPO;GRIA3 |
| Hepatitis C | 2/155 | 0.9970883 | 0.9999955 | 0.244026 | 7.12E-04 | PIK3CA;PIK3CD |
| Focal adhesion | 3/199 | 0.9978548 | 0.9999955 | 0.285365 | 6.13E-04 | PIK3CA;PTEN;PIK3CD |
| Epstein-Barr virus infection | 3/201 | 0.9980321 | 0.9999955 | 0.2824525 | 5.56E-04 | ENTPD3;PIK3CA;PIK3CD |
| Tight junction | 2/170 | 0.9985539 | 0.9999955 | 0.222061 | 3.21E-04 | CACNA1D;CFTR |
| Influenza A | 2/171 | 0.9986202 | 0.9999955 | 0.2207353 | 3.05E-04 | PIK3CA;PIK3CD |
| Systemic lupus erythematosus | 1/133 | 0.9990118 | 0.9999955 | 0.1414416 | 1.40E-04 | GRIN2B |
| Viral carcinogenesis | 2/201 | 0.9996649 | 0.9999955 | 0.1871598 | 6.27E-05 | PIK3CA;PIK3CD |
| Olfactory transduction | 7/444 | 0.999969 | 0.9999955 | 0.2959877 | 9.17E-06 | SLC24A4;GUCY2D;PDE1C;PDE1B;PDE1A;CNGA4;CNGB1 |
| Herpes simplex virus 1 infection | 2/492 | 0.9999955 | 0.9999955 | 0.0748326 | 3.36E-07 | PIK3CA;PIK3CD |
