## Supplementary material for "Glutamine antagonism suppresses tumor growth in adrenocortical carcinoma through inhibition of de novo nucleotide biosynthesis": Suppl. Table 3

**Suppl. Table 3 Significantly dysregulated polar metabolites BPCre-ACC tumors vs Control adrenals (Fig. 2c)**

| Metabolite | FC (ACC vs Ctrl) | log2(FC) | p.ajusted |
|---|---|---|---|
| Creatine Phosphate | 0.37611 | -1.4108 | 0.00017579 |
| XMP | 0.072634 | -3.7832 | 0.00020499 |
| Glutathione | 2.4828 | 1.312 | 0.00020499 |
| Saccharate/Galactarate | 0.46576 | -1.1023 | 0.00065315 |
| Ornithine | 4.1594 | 2.0564 | 0.00085236 |
| CDP-choline | 2.6074 | 1.3826 | 0.0008971 |
| Citramalate | 0.41202 | -1.2792 | 0.0012074 |
| Cytidine | 3.2955 | 1.7205 | 0.0012543 |
| Ophthalmic acid | 2.5297 | 1.339 | 0.0012702 |
| NADH | 2.2367 | 1.1613 | 0.0012702 |
| S-Adenosyl-L-homocysteine | 0.38479 | -1.3778 | 0.0013756 |
| L-Aspartate | 2.2556 | 1.1735 | 0.0013756 |
| O-Phosphorylethanolamine | 1.8681 | 0.90158 | 0.0018438 |
| GABA/2-aminoisobutyric acid | 2.0282 | 1.0202 | 0.0046204 |
| Glycerol | 0.545 | -0.87567 | 0.0068063 |
| Lysine | 2.1176 | 1.0825 | 0.0091693 |
| UDP-glucose/Uridine 5'-diphos | 1.9347 | 0.95214 | 0.0091693 |
| Creatine | 0.58907 | -0.7635 | 0.0091693 |
| Cyclic AMP | 1.6757 | 0.7448 | 0.0091693 |
| HMG | 2.6579 | 1.4103 | 0.0096952 |
| D-Fructose 1,6-bisphosphate | 0.42884 | -1.2215 | 0.0096952 |
| D-Galacturonate | 0.54387 | -0.87866 | 0.0096952 |
| NAD | 12.843 | 3.683 | 0.0099879 |
| Phosphorylcholine | 0.50424 | -0.98781 | 0.010224 |
| 3-Ureidopropionic acid | 1.4036 | 0.48912 | 0.010224 |
| Xanthosine | 0.41379 | -1.273 | 0.011258 |
| Xanthine | 0.48576 | -1.0417 | 0.011258 |
| UDP-N-acetylglucosamine/UDF | 1.8232 | 0.86649 | 0.011258 |
| Pantothenic acid | 0.55376 | -0.85266 | 0.011258 |
| 2PG/3PG | 0.38083 | -1.3928 | 0.011893 |
| cis-4-Hydroxy-D-proline | 0.55556 | -0.84798 | 0.012966 |
| Uracil | 0.5904 | -0.76023 | 0.012966 |
| L-Glutamate | 1.8533 | 0.89012 | 0.013881 |
| Guanidinoacetate | 0.54707 | -0.8702 | 0.013881 |
| 1,2-Dihydroxy-5-(methylthio)pe | 0.64881 | -0.62414 | 0.015315 |
| L-Proline | 1.936 | 0.95306 | 0.015332 |
| Phosphoenol pyruvate | 0.36577 | -1.451 | 0.01568 |
| D-Glucose | 0.19146 | -2.3849 | 0.015844 |
| L-Cysteine | 3.7023 | 1.8884 | 0.015844 |
| Uric Acid | 0.54065 | -0.88724 | 0.015844 |
| Cystine | 0.55291 | -0.85489 | 0.015844 |
| Nicotinamide | 0.67297 | -0.57139 | 0.015844 |
| Glucose 6-phosphate/Galactose | 0.28385 | -1.8168 | 0.017731 |
| CDP-ethanolamine | 2.4215 | 1.2759 | 0.01807 |
| Thymidine | 1.8484 | 0.88624 | 0.020101 |
| Carnosine | 0.65692 | -0.6062 | 0.024881 |
| dTTP | 0.7232 | -0.46752 | 0.024881 |
| N-Formimino-L-glutamate | 1.6922 | 0.75887 | 0.026278 |
| D-Ribose 1-phosphate/D-Ribos | 0.36771 | -1.4433 | 0.027981 |
| L-carnitine | 1.7511 | 0.80824 | 0.028496 |
| N,N-Dimethylarginine | 3.1176 | 1.6405 | 0.048143 |
