## Supplementary material for "Glutamine antagonism suppresses tumor growth in adrenocortical carcinoma through inhibition of de novo nucleotide biosynthesis": Suppl. Table 4

| LC-MS Assay | Peak Identifier | *m/z* | RT(s) | p-value (BH FDR corrected) | Fold change (ACC/ACA) | Metabolite/lipid annotation | Data applied for metabolite annotation |
|---|---|---|---|---|---|---|---|
| HILIC NEG | M449T72 | 449.1318 | 72.5 | 6.5592E-20 | 15.74 | ST 19:1;O2;S2 | MS1 |
| HILIC NEG | M379T84 | 379.1592 | 83.9 | 9.3538E-16 | 13.89 | Octanoylglucuronide | MS1 |
| HILIC POS | M273T303 | 273.2211 | 303.3 | 6.82E-23 | 10.38 | ST 19:2;O | MS1 |
| HILIC POS | M484T303 | 484.2908 | 303.3 | 1.16E-16 | 9.78 | ST 19:1;O2;GlcA | MS1 |
| HILIC NEG | M512T352 | 512.2952 | 351.7 | 4.6886E-21 | 8.52 | ST 21:1;O4; GlcA | MS1 |
| HILIC NEG | M451T76 | 451.1479 | 76.1 | 8.1911E-13 | 6.34 | ST 19:1;O3;S2 | MS1 |
| HILIC NEG | M495T323 | 495.2973 | 322.7 | 1.5024E-16 | 6.30 | ST 21:0;O2;GlcA | MS1 |
| HILIC NEG | M481T337 | 481.2453 | 336.6 | 0.0071 | 4.58 | ST 19:1;O3;GlcA | MS1 |
| HILIC NEG | M525T352 | 525.2712 | 352.0 | 4.3186E-07 | 4.46 | ST 19:1;O2;GlcA | MS1 |
| LIPID NEG | M431T91 | 431.3167 | 90.7 | 2.31E-22 | 4.09 | ST 27:2;O4 / ST 28:1;O4 | MS1 |
| HILIC POS | M264T481 | 264.1189 | 481.3 | 0.00025441 | 3.04 | Creatine riboside; 5,6-Dihydrouridine | MS1 |
| HILIC POS | M432T323 | 432.3103 | 322.9 | 0.0034109 | 2.56 | 11'-Carboxy-alpha-tocotrienol | MS1 |
| HILIC NEG | M143T70 | 143.0465 | 70.2 | 0.031023 | 2.43 | Imidazolone; N-Nitrosoproline | MS1 |
| HILIC NEG | M227T357 | 227.1041 | 357.5 | 0.00038861 | 2.27 | Pyridoxamine | MS1 |
| HILIC NEG | M275T305 | 275.1042 | 305.3 | 0.00014447 | 2.23 | N-lactoyl-Tryptophan; L-1,2,3,4-Tetrahydro-beta-carboline-3-carboxylic a | MS1 |
| HILIC POS | M287T502 | 287.2439 | 502.0 | 3.15E-05 | 2.17 | N1,N12-Diacetylspermine | MS1 |
| HILIC POS | M279T453 | 279.1062 | 453.0 | 0.048497 | 2.15 | Isovalerylglucuronide; 2-(3-Carboxy-3-aminopropyl)-L-histidine | MS1 |
| HILIC POS | M247T429 | 247.1398 | 429.5 | 0.0069823 | 2.13 | Octopine | MS1 |
| HILIC POS | M414T323 | 414.2997 | 322.5 | 0.00028184 | 2.13 | N-Docosahexaenoyl GABA | MS1 |
| HILIC NEG | M496T325 | 496.3050 | 325.5 | 1.6664E-06 | 2.07 | LPE P-16:0; Galactosylsphingosine; Glucosylsphingosine | MS1 |
| HILIC NEG | M256T497 | 256.1307 | 496.6 | 0.0014846 | 1.97 | L-Histidine trimethylbetaine; | MS1 |
| HILIC POS | M273T147 | 273.0841 | 147.2 | 0.006478 | 1.95 | Hypoxanthine | MS/MS and RT |
| HILIC POS | M323T367 | 323.0384 | 366.9 | 0.037445 | 1.90 | Xanthosine; L-arginino-succinate | MS1 |
| HILIC POS | M267T470 | 267.1086 | 469.8 | 3.01E-05 | 1.87 | 6-Nitrotryptophan; Nitrotryptophan | MS1 |
| HILIC NEG | M185T429_2 | 185.0569 | 429.0 | 0.0071 | 1.85 | Thymine; Imidazoleacetic acid; Pyroglutamylglycine; 1H-Imidazole-1-ace | MS1 |
| HILIC POS | M282T500_2 | 282.0965 | 500.4 | 0.0022446 | 1.85 | 4-Hydroxyphenylacetylglutamic acid; 4-hydroxyphenylacetylglutamine | MS1 |
| HILIC POS | M296T500 | 296.0658 | 500.4 | 0.00016483 | 1.84 | Glycerophosphocholine | MS1 |
| HILIC NEG | M257T496 | 257.0782 | 496.5 | 3.3508E-06 | 1.79 | 2'-O-Methyluridine; 3-Methyluridine; 5-Hydroxymethyl-2'-deoxyuridine; A | MS1 |
| HILIC POS | M264T467 | 264.1188 | 466.8 | 0.0034109 | 1.76 | Creatine riboside; 5,6-Dihydrouridine | MS1 |
| HILIC NEG | M289T309_2 | 289.1215 | 309.0 | 0.029499 | 1.72 | Tetradecadienedioic acid; N-Palmitoyl Cysteine | MS1 |
| HILIC POS | M149T479_1 | 149.0574 | 478.8 | 0.025495 | 1.72 | heptadienoic acid; 1-Cyclohexene-1-carboxylic acid | MS1 |
| HILIC POS | M245T471 | 245.0533 | 470.8 | 0.019285 | 1.69 | L-Glutamic acid 5-phosphate; (2S,3R)-2-Amino-3-[(2S)-2-amino-3-hydro | MS1 |
| HILIC NEG | M242T451 | 242.0784 | 450.9 | 0.0021918 | 1.67 | Cytidine; Arabinofuranosylcytosine | MS1 |
| HILIC POS | M448T209 | 448.3416 | 209.0 | 0.025495 | 1.67 | Arachidonoylcarnitine | MS1 and MS/MS |
| HILIC POS | M229T471 | 229.0793 | 470.8 | 0.022337 | 1.67 | (2S,3R)-2-Amino-3-[(2S)-2-amino-3-hydroxypropanoyl]oxybutanoic acid | MS1 |
| HILIC POS | M568T497 | 568.1842 | 497.3 | 0.0074244 | 1.66 | N-Acetylgalactosaminyl lactose | MS1 |
| LIPID NEG | M466T326 | 466.3074 | 325.7 | 4.32E-07 | 1.59 | ST 27:1;O;S | MS1 |
| HILIC POS | M241T79 | 241.0929 | 79.1 | 0.0047467 | 1.59 | D-glycero-L-galacto-Octulose | MS1 |
| HILIC NEG | M529T347 | 529.2676 | 346.7 | 0.0003952 | 1.57 | PA 21:0 | MS1 |
| HILIC POS | M238T480 | 238.0447 | 479.7 | 0.0034109 | 1.56 | Galactosamine; Glycerylphosphorylethanolamine | MS1 |
| LIPID NEG | M549T326 | 549.2659 | 325.8 | 3.75E-07 | 1.56 | ST 27:1;O4;S | MS1 |
| HILIC POS | M131T479_1 | 131.0338 | 478.8 | 0.012773 | 1.55 | Citraconic acid; Gamma-delta-Dioxovaleric acid; Glutaconic acid; Itaconi | MS1 |
| HILIC NEG | M189T552 | 189.0520 | 552.0 | 0.0091317 | 1.54 | L-beta-aspartyl-L-glycine; N-carbamoylglutamic Acid; 5-Hydroxy-1-methy | MS1 |
| HILIC POS | M247T443 | 247.0922 | 442.5 | 0.015156 | 1.53 | 5,6-Dihydrouridine; 4,5-seco-dopa; 2,6-Diamino-9-(2-hydroxyethoxymeth | MS1 |
| LIPID NEG | M601T326 | 601.2793 | 325.9 | 2.03E-06 | 1.52 | LPG 22:6 | MS1 |
| HILIC POS | M430T285 | 430.3157 | 285.1 | 0.019285 | 1.52 | Hexadecanedioic acid mono-L-carnitine ester | MS1 |
| HILIC POS | M148T479 | 148.0603 | 478.8 | 0.011436 | 1.51 | L-4-Hydroxyglutamate semialdehyde; Mesaconic acid; N-Acetylserine; N- | MS1 |
| HILIC NEG | M257T82 | 257.1763 | 82.2 | 0.04475 | 1.51 | Dodecenoic acid; Tetradecanedioic acid | MS1 |
| HILIC POS | M102T479 | 102.0549 | 478.8 | 0.012626 | 1.50 | 4-Hydroxy-2-butenoic acid gamma-lactone; N-methyl-beta-alanine; 1-Am | MS1 |
| HILIC NEG | M219T552 | 219.0626 | 551.8 | 0.02173 | 1.50 | Thymine glycol; (Z)-2-methylureidoacrylate peracid | MS1 |
| HILIC POS | M258T435 | 258.1083 | 434.6 | 0.0061823 | 1.49 | 2'-C-Methylcytidine; 2'-O-Methylcytidine; 3-Methylcytidine; 5-Methylcyti | MS1 |
| HILIC NEG | M528T347 | 528.2644 | 346.7 | 0.0018039 | 1.48 | Glycochenodeoxycholate-3-sulfate; glycodeoxycholate sulfate | MS1 |
| LIPID NEG | M311T96 | 311.2226 | 96.1 | 0.0077279 | 1.48 | FA 18:2;O2 / FA 17:2 / SFE 17:2 | MS1 |
| HILIC NEG | M256T443 | 256.0943 | 443.4 | 0.0003952 | 1.48 | 2'-C-Methylcytidine; 2'-O-Methylcytidine; 3-Methylcytidine; 5-Methylcyti | MS1 |
| LIPIDS POS | M337T503 | 337.2731 | 503.0 | 0.027757 | 1.47 | FA 21:3;O | MS1 |
| LIPID NEG | M617T326 | 617.2525 | 325.9 | 3.67E-06 | 1.46 | LPI 17:2 | MS1 |
| HILIC NEG | M243T499 | 243.0625 | 499.1 | 0.000016197 | 1.46 | Uridine; Pseudouridine | MS1 |
| LIPID NEG | M860T499 | 859.5493 | 498.8 | 0.044913 | 1.43 | PI O-34:0 | MS1 |
| LIPID NEG | M612T519 | 611.5441 | 518.9 | 0.021694 | 1.43 | Cer 36:1;O2 | MS1 |
| HILIC POS | M300T164 | 300.2895 | 163.6 | 0.022337 | 1.41 | Sphingosine | MS1 and MS/MS |
| HILIC POS | M246T370 | 246.1082 | 369.7 | 0.048412 | 1.40 | Deoxyuridine | MS1 |
| HILIC POS | M260T500_1 | 260.0737 | 500.4 | 0.024261 | 1.38 | Fructoseglycine; N-(1-Deoxy-1-fructosyl)glycine; N-Glycolylglucosamine | MS1 |
| HILIC NEG | M202T231 | 202.1088 | 230.5 | 0.02069 | 1.37 | Acetylcarnitine; N-Lactoylleucine; 1-Aminocyclohexanecarboxylic acid; L- | MS1 |
| LIPID NEG | M1538T509 | 1538.1298 | 509.1 | 0.020862 | 1.37 | CL 74:0 | MS1 |
| LIPIDS POS | M781T501 | 780.5507 | 500.7 | 0.043103 | 1.36 | PC 34:2 / PC 36:5 | MS1 |
| LIPID NEG | M584T496 | 583.5133 | 496.3 | 0.00012567 | 1.36 | Cer 34:1;O2 | MS1 |
| HILIC NEG | M308T483 | 308.0992 | 482.5 | 0.000027072 | 1.36 | N-Acetylneuraminic acid; 9-O-Acetylneuraminic acid; Glucose-6-glutami | MS1 |
| HILIC POS | M450T213 | 450.3575 | 213.3 | 0.037445 | 1.35 | Stearoylcarnitine; 1,24,25-trihydroxyvitamin D3; 1alpha,23(S),25-trihydro | MS1 |
| LIPID NEG | M526T129_1 | 526.2940 | 129.1 | 0.0013627 | 1.35 | LPE 22:5 | MS1 |
| LIPID NEG | M766T496_1 | 765.5470 | 496.4 | 0.023517 | 1.35 | PC 31:0 / PE 34:0 | MS1 |
| HILIC POS | M125T500 | 124.9999 | 500.4 | 0.029089 | 1.34 | But-2-enoic acid; Cyclopropanecarboxylic acid; gamma-Butyrolactone; Is | MS1 |
| HILIC POS | M246T539 | 246.1811 | 538.5 | 0.0066083 | 1.34 | N,N,N-Trimethyl-L-alanyl-L-proline betaine | MS1 |
| HILIC POS | M336T500 | 336.0872 | 500.4 | 0.018557 | 1.33 | S-Formylglutathione | MS1 |
| HILIC POS | M260T500_2 | 260.1141 | 500.4 | 0.0034109 | 1.33 | Elenaic acid; Fumarycarnitine; Lumichrome | MS1 |
| LIPID NEG | M715T483 | 714.5071 | 483.4 | 0.010231 | 1.33 | PE(16:0/18:2) | MS1 |
| LIPIDS POS | M755T505 | 754.5927 | 505.4 | 0.011054 | 1.32 | PG O-34:0 | MS1 |
| HILIC POS | M133T247 | 133.0607 | 246.8 | 0.04877 | 1.32 | Asparagine; N-Carbamoylsarcosine; Ureidopropionic acid | MS1 |
| HILIC POS | M243T425 | 243.0951 | 425.0 | 0.00068439 | 1.32 | N-Acetyl-b-glucosaminylamine | MS1 |
| HILIC POS | M151T121 | 151.0614 | 121.4 | 0.011474 | 1.30 | 1-Methylhypoxanthine; 7-Methylhypoxanthine; 1,6-Hexanedithiol | MS1 |
| LIPID NEG | M785T503 | 784.5098 | 503.2 | 0.029233 | 1.30 | PS 36:3 / PC 33:4 / PE 36:4 / PE O-36:5;O | MS1 |
| HILIC NEG | M214T487 | 214.0489 | 486.8 | 0.00057902 | 1.29 | Glucosamine; N-Methylethanolaminium phosphate; Galactosamine; Gly | MS1 |
| HILIC POS | M538T313 | 538.4210 | 312.8 | 0.008237 | 1.27 | ST 24:1;O4;Lys | MS1 |
| HILIC POS | M282T86 | 282.1196 | 86.0 | 0.0032386 | 1.27 | 1-Methyladenosine; 2'-O-Methyladenosine; 3'-O-Methyladenosine; N-(1- | MS1 |
| HILIC POS | M207T311 | 207.1125 | 310.8 | 0.010955 | 1.26 | Indole-3-propionic acid; methyl indole-3-acetic acid; Phenylethylmalonan | MS1 |
| LIPID NEG | M694T529 | 693.5471 | 528.7 | 0.044913 | 1.26 | DG 42:7 | MS1 |
| LIPIDS POS | M631T541 | 630.6178 | 540.5 | 0.00061727 | 1.26 | Cer 40:0;O | MS1 |
| HILIC NEG | M151T233 | 151.0264 | 232.8 | 0.027866 | 1.26 | Xanthine; Oxypurinol; 6,8-Dihydroxypurine | MS1 |
| LIPIDS POS | M645T541 | 644.5947 | 540.6 | 0.015546 | 1.25 | Cer 40:1;O2 | MS1 |
| HILIC POS | M280T537_2 | 280.1879 | 537.0 | 0.014384 | 1.25 | N-Lauroylglycine | MS1 |
| LIPIDS POS | M637T503 | 636.5555 | 503.4 | 0.032841 | 1.24 | DG(18:1/18:2) | MS1 and MS/MS |
| HILIC POS | M208T479 | 207.9981 | 479.2 | 0.027558 | 1.24 | Phosphoserine; 3-phospho-L-serine; DL-O-Phosphoserine | MS1 and RT |
| LIPID NEG | M795T510 | 794.5435 | 509.9 | 0.00065339 | 1.24 | PC 34:1 [PC 16:0/18:1] | MS1 and MS/MS |
| LIPIDS POS | M754T505 | 753.5883 | 505.4 | 0.0079896 | 1.24 | CerPE 39:1;O2 / SM 36:1;O2 | MS1 |
| HILIC POS | M665T271 | 664.5263 | 270.9 | 0.02314 | 1.24 | LPC 28:0 | MS1 |
| LIPID NEG | M476T126 | 476.2779 | 125.9 | 0.042573 | 1.23 | LPE 18:2 | MS1 |
| LIPID NEG | M452T161 | 452.2781 | 161.4 | 0.0048875 | 1.23 | LPE 16:0 | MS1 |
| LIPIDS POS | M775T451 | 774.5630 | 451.2 | 0.025528 | 1.23 | PS O-36:2 / PG O-36:4 | MS1 |
| LIPIDS POS | M723T465 | 722.5528 | 465.1 | 0.0176 | 1.23 | HexCer 34:1;O2 | MS1 |
| LIPID NEG | M790T476 | 789.5473 | 475.6 | 0.01898 | 1.22 | PA 43:6 | MS1 |
| HILIC POS | M121T351_1 | 121.0645 | 350.9 | 0.022375 | 1.22 | 3-Methylbenzaldehyde; 4-Methylbenzaldehyde | MS1 |
| LIPID NEG | M500T112 | 500.2785 | 111.6 | 0.0052198 | 1.21 | ST 24:1;O4;Gly / LPE 20:4 | MS1 |
| LIPID NEG | M501T120 | 501.2818 | 120.5 | 0.0071828 | 1.21 | ST 24:1;O4;Gly / LPE 20:4 | MS1 |
| LIPID NEG | M453T161 | 453.2816 | 161.3 | 0.0072565 | 1.21 | LPC 13:0 / LPE 16:0 / LPE O-16:1;O | MS1 |
| LIPID NEG | M520T161 | 520.2659 | 160.9 | 0.011824 | 1.21 | LPS 18:2 / LPE 18:3 | MS1 |
| HILIC POS | M149T351 | 149.0596 | 350.9 | 0.00020887 | 1.21 | Cinnamic acid; p-Coumaraldehyde; trans-Cinnamic acid | MS1 |
| LIPIDS POS | M547T437 | 546.5239 | 436.6 | 0.04088 | 1.21 | FA 36:4 / Cer 34:0;O | MS1 |
| LIPID NEG | M568T120 | 568.2654 | 120.5 | 0.012985 | 1.21 | LPS 22:6 | MS1 |
| HILIC POS | M138T119 | 138.0549 | 118.6 | 0.026202 | 1.20 | 2-Aminobenzoic acid; 2-Pyridylacetic acid; 3-Pyridylacetic acid; 6-Methyl- | MS1 |
| LIPID NEG | M647T556 | 646.6126 | 556.4 | 0.047018 | 1.20 | Cer 42:2;O2 | MS1 |
| LIPID NEG | M526T129_2 | 526.3149 | 129.4 | 0.030112 | 1.20 | LPC 15:0 / LPE 18:0 / LPE O-18:1;O | MS1 |
| LIPIDS POS | M745T475 | 744.5526 | 474.9 | 0.021928 | 1.20 | PC 33:2 | MS1 |

| | | | | | | |
|---|---|---|---|---|---|---|
| HILIC POS | M120T351_2 | 120.0807 | 350.9 | 0.00031984 | 1.19 Indoline; Isoindoline | MS1 |
| HILIC POS | M167T351_2 | 167.0892 | 350.9 | 0.00046419 | 1.19 Phenylalanine; Methyl N-methylanthranilate; N,N-Dimethylanthranilic ac | MS1 |
| LIPID NEG | M483T139 | 483.2729 | 139.0 | 0.044913 | 1.18 LPG 16:0 / LPA 18:0 | MS1 |
| HILIC POS | M274T452 | 274.0919 | 452.3 | 0.048412 | 1.18 5'-Deoxyadenosine; Deoxyadenosine; 2',3'-Dideoxyguanosine | MS1 |
| LIPID NEG | M1567T509 | 1567.1598 | 509.4 | 0.026811 | 1.18 CL 76:0 | MS1 |
| HILIC NEG | M205T319 | 205.0985 | 318.6 | 0.016241 | 1.18 Indole-3-methanamine | MS1 |
| HILIC POS | M148T351 | 148.0754 | 351.0 | 0.0023632 | 1.17 3-Methyloxindole; 5-Methoxyindole; Cinnamamide; Indole-3-carbinol; Me | MS1 |
| LIPID NEG | M542T149 | 542.3368 | 149.3 | 0.010103 | 1.17 LPC O-18:1 | MS1 |
| LIPID NEG | M608T149 | 608.3181 | 149.3 | 0.0055978 | 1.17 PE 23:2;O3 / PE 22:2;O | MS1 |
| LIPIDS POS | M834T533 | 833.6492 | 532.7 | 0.038042 | 1.17 SM 40:0;O2 / SM 42:3;O2 | MS1 |
| LIPID NEG | M540T149 | 540.3307 | 149.3 | 0.010222 | 1.17 LPS O-19:0;O / LPC 16:0 / LPE 19:0 / LPE O-19:1;O | MS1 |
| HILIC POS | M190T351 | 190.0860 | 351.0 | 0.0048852 | 1.16 2-Methyl-2-phenylsuccinimide; 2-Naphthoic acid; Indole-3-methyl acetat | MS1 |
| LIPID NEG | M847T509 | 846.5465 | 508.6 | 0.034171 | 1.16 PS 37:2 | MS1 |
| LIPID NEG | M480T164 | 480.3093 | 164.1 | 0.0063865 | 1.16 LPC 15:0 / LPE 18:0 / LPE O-18:1;O | MS1 |
| LIPID NEG | M766T531 | 765.5832 | 531.3 | 0.025696 | 1.16 TG 43:3 / PA O-42:4 | MS1 |
| LIPIDS POS | M882T606 | 881.6987 | 606.0 | 0.024698 | 1.16 TG 51:3 | MS1 |
| LIPID NEG | M591T66 | 591.3539 | 66.4 | 0.022509 | 1.15 ST 27:3;O4;Hex | MS1 |
| LIPIDS POS | M765T510 | 764.5965 | 509.9 | 0.019555 | 1.15 PC 33:1 / PE 36:1 | MS1 |
| LIPID NEG | M541T164 | 541.3338 | 164.1 | 0.016882 | 1.15 LPS O-19:0;O / LPC 16:0 / LPE 19:0 / LPE O-19:1;O | MS1 |
| LIPID NEG | M776T507 | 775.5958 | 507.5 | 0.0063797 | 1.14 SM 36:1;O2 | MS1 |
| LIPIDS POS | M721T536 | 720.5895 | 535.6 | 0.040056 | 1.14 PC O-32:0 | MS1 |
| LIPID NEG | M530T164 | 530.3020 | 164.3 | 0.019382 | 1.12 LPC 16:0 / LPE 19:0 / LPE O-19:1;O | MS1 |
| LIPID NEG | M746T509 | 745.5566 | 509.4 | 0.022026 | 1.12 DG 43:6 | MS1 |
| LIPID NEG | M608T164 | 608.3179 | 164.1 | 0.026771 | 1.12 PE 23:2;O3 / PE 22:2;O | MS1 |
| LIPID NEG | M806T509 | 805.5777 | 509.4 | 0.011603 | 1.12 PA 44:5 | MS1 |
| LIPID NEG | M793T488 | 792.5302 | 487.8 | 0.0089454 | 1.11 PC 34:2 [PC 16:0/18:2] | MS1 and MS/MS |
| LIPID NEG | M872T488 | 871.5489 | 487.8 | 0.015334 | 1.10 PI O-35:1 | MS1 |
| HILIC NEG | M147T356 | 147.0454 | 355.7 | 0.034938 | 1.10 Cinnamic acid; p-Coumaraldehyde; trans-Cinnamic acid | MS1 |
| LIPIDS POS | M735T509 | 734.5687 | 509.0 | 0.032485 | 1.10 PC 32:0 | MS1 |
| LIPIDS POS | M759T487 | 758.5685 | 487.2 | 0.001002 | 1.09 PC 34:2 | MS1 |
| LIPID NEG | M779T508 | 778.5592 | 508.4 | 0.019323 | 1.09 PC 32:0 [PC 16:0/16:0] | MS1 and MS/MS |
| LIPID NEG | M804T488 | 803.5620 | 487.6 | 0.047752 | 1.08 PA 44:6 | MS1 |
| HILIC NEG | M175T490 | 175.0726 | 490.3 | 0.038753 | 1.02 N5-formyl-N5-hydroxy-L-ornithine | MS1 |
| HILIC NEG | M307T417 | 307.0278 | 417.1 | 0.03629 | 0.94 2-Hydroxy-4-methoxybenzophenone sulfate | MS1 |
| HILIC POS | M162T446_2 | 162.1122 | 445.9 | 0.047562 | 0.93 Carnitine | MS1 and MS/MS |
| HILIC POS | M103T446 | 103.0389 | 445.9 | 0.037445 | 0.92 Methylmalonic acid semialdehyde; 2-Ketobutyric acid; 2-Methyl-3-oxopro | MS1 |
| HILIC NEG | M174T289_2 | 174.0755 | 288.7 | 0.040964 | 0.92 Indole-3-acetamide | MS1 |
| HILIC NEG | M471T417 | 471.0344 | 417.3 | 0.0032038 | 0.92 dIDP | MS1 |
| HILIC NEG | M239T539_2 | 238.8917 | 539.3 | 0.013906 | 0.92 Trimetaphosphoric acid | MS1 |
| LIPIDS POS | M958T616 | 957.7871 | 616.3 | 0.047232 | 0.92 TG 58:6 / TG 56:3 / TG 60:9 | MS1 |
| HILIC NEG | M116T413 | 116.0718 | 412.6 | 0.044481 | 0.91 Valine; N-Methyl-4-aminobutyric acid; N-Methyl-a-aminoisobutyric acid; | MS1 |
| HILIC NEG | M118T461 | 118.0512 | 461.3 | 0.020501 | 0.91 Threonine; N-Methyl-L-serine; O-Methyl-L-serine; 4-Amino-3-hydroxybut | MS1 |
| LIPID NEG | M853T483 | 852.5736 | 483.0 | 0.048842 | 0.91 PC 38:5 [PC 18:0/20:5] | MS1 and MS/MS |
| LIPIDS POS | M605T611 | 604.5379 | 610.7 | 0.015449 | 0.91 DG O-34:1 | MS1 |
| HILIC NEG | M257T112 | 257.0784 | 112.4 | 0.023985 | 0.91 2'-O-Methyluridine; 3-Methyluridine; 5-Hydroxymethyl-2'-deoxyuridine; A | MS1 |
| HILIC POS | M140T406_2 | 140.0680 | 405.6 | 0.029904 | 0.90 Valine; N-Methyl-4-aminobutyric acid; N-Methyl-a-aminoisobutyric acid; | MS1 |
| HILIC NEG | M183T339 | 183.0414 | 339.4 | 0.028255 | 0.90 Urocanic acid | MS1 |
| HILIC POS | M787T263 | 786.5981 | 263.1 | 0.022337 | 0.90 PC 36:2 | MS1 |
| HILIC POS | M790T263 | 789.6156 | 263.1 | 0.028468 | 0.90 PC 35:2 ; PE 38:2 | MS1 |
| LIPID NEG | M845T527 | 844.6053 | 527.3 | 0.048132 | 0.90 PC 37:2 [PC 19:0/18:2] | MS1 and MS/MS |
| LIPID NEG | M750T491 | 749.5797 | 491.4 | 0.028408 | 0.89 SM 34:0;O2 | MS1 |
| HILIC NEG | M177T539_2 | 176.9338 | 539.4 | 0.0304 | 0.89 diphosphate | MS1 |
| LIPIDS POS | M948T617 | 947.7455 | 616.9 | 0.0057878 | 0.89 TG 56:5 | MS1 |
| HILIC POS | M801T262_1 | 800.6127 | 262.5 | 0.047664 | 0.89 PC 37:2 ; PA 42:3 ; PE 40:2 | MS1 |
| HILIC NEG | M858T306 | 857.5174 | 306.4 | 0.033619 | 0.89 PI 36:4 [PI 16:0_20:4] | MS1 and MS/MS |
| HILIC NEG | M686T303 | 685.5261 | 303.0 | 0.0023507 | 0.89 SM d33:2 | MS1 |
| HILIC NEG | M101T415 | 101.0245 | 415.4 | 0.023669 | 0.88 Methylmalonic acid semialdehyde; 2-Ketobutyric acid; 2-Methyl-3-oxopro | MS1 |
| HILIC NEG | M85T415 | 85.0296 | 415.4 | 0.042203 | 0.88 But-2-enoic acid; gamma-Butyrolactone; Isocrotonic acid; Methylmalond | MS1 |
| HILIC NEG | M218T405 | 218.0227 | 405.3 | 0.019972 | 0.88 3,5-Dihydroxyphenylglycine | MS1 |
| HILIC POS | M160T477_1 | 160.0736 | 477.1 | 0.020357 | 0.88 Tyramine; 2-Anilinoethanol; 2-Hydroxyphenethylamine; 2-Methoxy-5-me | MS1 |
| LIPIDS POS | M835T523 | 834.5972 | 522.7 | 0.026415 | 0.88 PC 38:3 | MS1 |
| HILIC NEG | M163T405 | 163.0403 | 405.4 | 0.0033131 | 0.88 2-Hydroxycinnamic acid; 4-Hydroxycinnamic acid; Coumarinic acid; Enol-p | MS1 |
| HILIC NEG | M161T415 | 161.0458 | 415.4 | 0.032214 | 0.88 2-Hydroxyadipic acid; 3-Hydroxyadipic acid; 4-Hydroxycrotonic acid; Aceto | MS1 |
| HILIC NEG | M283T339 | 283.0828 | 339.4 | 0.034072 | 0.88 2-O-Benzoyl-D-glucose; 6-O-Benzoyl-alpha-D-glucose; p-Cresol glucuroni | MS1 |
| HILIC NEG | M583T314 | 583.3819 | 313.5 | 0.014332 | 0.87 LPC 18:0 | MS1 |
| HILIC POS | M829T255 | 828.5263 | 254.8 | 0.022337 | 0.87 PC P-38:6 ; 3-O-Sulfogalactosylceramide d36:2 | MS1 |
| HILIC NEG | M179T415 | 179.0562 | 415.1 | 0.028979 | 0.87 Glucose; Galactose; Fructose; Mannose; myo-Inositol; 3,4-Dihydroxybutyr | MS1 |
| HILIC NEG | M146T473_1 | 146.0592 | 473.2 | 0.0019742 | 0.87 (R)-pantoate; 3-Methyloxindole; 5-Methoxyindole; Cinnamamide; Indole- | MS1 |
| HILIC NEG | M295T112 | 295.0518 | 112.5 | 0.038401 | 0.87 Biotin sulfoxide | MS1 |
| LIPID NEG | M925T506 | 924.5840 | 506.2 | 0.010222 | 0.87 PS 43:6 | MS1 |
| HILIC NEG | M862T309 | 861.5485 | 308.7 | 0.029499 | 0.87 PI 36:2 | MS1 |
| HILIC NEG | M146T486 | 146.0824 | 486.4 | 0.010702 | 0.87 (2R,3R,4R)-2-Amino-4-hydroxy-3-methylpentanoic acid; 4-Aminobutyralc | MS1 |
| HILIC NEG | M180T405 | 180.0668 | 405.4 | 0.0022387 | 0.87 Tyrosine; L-Threo-3-Phenylserine; 4-Hydroxy-4-(3-pyridyl)-butanoic acid; I | MS1 |
| LIPIDS POS | M816T522 | 815.6249 | 521.8 | 0.023157 | 0.87 PC 37:3 / PE 40:3 | MS1 |
| HILIC NEG | M127T473 | 127.0515 | 473.2 | 0.0011788 | 0.87 5,6-dihydrothymine; Dihydrothymine; L-Cyclo(alanylglycyl); Pyroglutamine | MS1 |
| HILIC NEG | M164T198 | 164.0581 | 197.8 | 0.014394 | 0.87 1-Methylguanine; 2-Amino-6-methoxypurine; 3-Methylguanine; 7-Methyl | MS1 |
| HILIC POS | M540T323 | 540.3655 | 323.1 | 0.012626 | 0.87 Cholylasparagine; PA 23:0 | MS1 |
| HILIC NEG | M847T271 | 846.6161 | 270.5 | 0.032054 | 0.87 PE P-42:2 | MS1 |
| LIPIDS POS | M801T529 | 800.6153 | 529.2 | 0.046259 | 0.87 PC 37:2 | MS1 |
| HILIC NEG | M220T486 | 220.1193 | 486.4 | 0.0091331 | 0.87 L-Carnitine | MS1 |
| HILIC NEG | M131T415 | 131.0351 | 415.4 | 0.013906 | 0.87 Ethylmalonic acid; Glutaric acid; Methylsuccinic acid; 2-Acetolactate; 2-C | MS1 |
| HILIC NEG | M508T313 | 508.3419 | 313.5 | 0.010199 | 0.87 LPE 20:0; LPC 17:0 | MS1 |
| HILIC POS | M634T305_2 | 634.4784 | 304.7 | 0.037445 | 0.86 LPC 26:1 | MS1 |
| HILIC NEG | M845T271 | 844.6056 | 270.5 | 0.015108 | 0.86 PC 36:2 ; PS 40:1 | MS1 |
| HILIC NEG | M143T415 | 143.0349 | 415.2 | 0.022027 | 0.86 2-Methylglutaconic acid; 3-Hexenedioic acid; 3-Hydroxyadipic acid 3,6-lac | MS1 |
| LIPIDS POS | M907T613 | 906.7586 | 612.7 | 0.010814 | 0.86 TG 52:1 / TG 54:4 / TG 56:7 | MS1 |
| LIPIDS POS | M812T506 | 811.6031 | 506.1 | 0.0018544 | 0.86 PC 38:4 / PE 41:4 / PC 36:1 / PE 39:1 | MS1 |
| HILIC NEG | M771T271 | 770.5688 | 270.6 | 0.0003952 | 0.86 PC 35:2 ; PE 38:2 | MS1 |
| HILIC POS | M243T80 | 242.9941 | 79.8 | 0.0082767 | 0.86 2-Methoxyhydroquinone sulfate; 2,5 Dihydroxybenzyl alcohol sulfate; 5-M | MS1 |
| HILIC NEG | M737T303 | 736.5253 | 302.8 | 0.042352 | 0.86 PC P-34:4 | MS1 |
| HILIC NEG | M314T473_2 | 314.1171 | 473.2 | 0.0045777 | 0.86 Feruloylcholine; phenylacetylcarnitine | MS1 |
| HILIC NEG | M870T267 | 869.6093 | 266.6 | 0.011314 | 0.86 PC 38:4 ; PE 42:4 ; PS 42:3 | MS1 |
| HILIC NEG | M145T473_1 | 145.0621 | 473.2 | 0.00051028 | 0.86 Glutamine; Glycyl-D-Alanine; Glycylsarcosine; Ureidoisobutyric acid; 2-Im | MS1 |
| HILIC NEG | M128T473_1 | 128.0355 | 473.2 | 0.00014398 | 0.86 1-Pyrroline-4-hydroxy-2-carboxylate; 4-Oxoproline; Pyroglutamic acid; Pyr | MS1 |
| HILIC POS | M484T301 | 484.2820 | 301.1 | 0.04064 | 0.86 N-Oleoyl tyrosine | MS1 |
| HILIC NEG | M319T473 | 319.0275 | 473.2 | 0.0012366 | 0.86 6-Thioinosine; 6-Thioinosinic acid | MS1 |
| LIPID NEG | M911T551 | 910.6586 | 551.3 | 0.012985 | 0.86 PC 42:4 | MS1 |
| HILIC NEG | M736T303 | 735.5219 | 302.8 | 0.044228 | 0.86 SM d34:2 | MS1 |
| LIPID NEG | M859T538 | 858.6210 | 537.5 | 0.0027837 | 0.85 PC 38:2 / PE 41:2 / PS 41:1 | MS1 |
| HILIC POS | M812T259 | 811.6022 | 258.6 | 0.0029146 | 0.85 PC 36:1 ; PC 38:4 ; PE 39:1 | MS1 |
| HILIC POS | M816T262 | 815.6284 | 261.8 | 0.0026344 | 0.85 PC 37:3 ; PE 40:3 | MS1 |
| LIPIDS POS | M831T450 | 830.5679 | 450.3 | 0.018135 | 0.85 PC 40:8 | MS1 |
| HILIC NEG | M891T265 | 890.5908 | 265.1 | 0.043485 | 0.85 PC 40:7 ; PS 44:6 | MS1 |
| LIPID NEG | M926T506 | 925.5877 | 506.2 | 0.0092483 | 0.85 TG 53:14;O2 | MS1 |
| LIPIDS POS | M814T522 | 813.6187 | 521.7 | 0.003172 | 0.85 PC 38:3 / PE 41:3 / PA 43:4 / PC 36:0 / PE 39:0 | MS1 |
| LIPID NEG | M855T507 | 854.5891 | 506.7 | 0.00023041 | 0.85 PC 38:4 [PC 18:0/20:4] | MS1 and MS/MS |
| HILIC POS | M836T257 | 835.6022 | 257.4 | 0.037445 | 0.85 Ubiquinol-9 | MS1 |
| HILIC NEG | M146T411 | 146.0825 | 411.3 | 0.0028015 | 0.85 4-Aminobutyraldehyde; Butyramide; Isobutyramide; (2R,3R,4R)-2-Amino | MS1 |
| HILIC POS | M102T464_1 | 102.0549 | 464.3 | 0.034617 | 0.85 (S)-2-Azetidinecarboxylic acid; 1-Aminocyclopropanecarboxylic acid; 4-Hy | MS1 |
| HILIC NEG | M886T304 | 885.5485 | 304.5 | 0.00029806 | 0.85 PI 38:4 [PI 18:0_20:4] | MS1 and MS/MS |
| HILIC NEG | M129T65_2 | 129.0559 | 65.1 | 0.043295 | 0.85 2-Methyl-3-ketovaleric acid; 3-Methyl-2-oxovaleric acid | MS1 and RT |
| LIPIDS POS | M852T588 | 851.7007 | 588.4 | 0.0018544 | 0.85 SM 43:1;O2 | MS1 |
| LIPID NEG | M757T553 | 756.5898 | 552.7 | 0.003251 | 0.84 PE O-38:2 | MS1 |

| | | | | | | | |
|---|---|---|---|---|---|---|---|
| HILIC POS | M636T304 | 636.4951 | 303.9 | 0.013825 | 0.84 | LPC 26:0 | MS1 |
| HILIC POS | M220T476 | 220.0666 | 476.4 | 0.019313 | 0.84 | 2-Oxo-4-hydroxy-4-carboxy-5-ureidoimidazole; 2-Oxo-4-hydroxy-4-carbox | MS1 |
| LIPID NEG | M876T452 | 875.5619 | 452.4 | 0.021849 | 0.84 | PI 37:2 | MS1 |
| LIPID NEG | M927T538 | 926.6098 | 538.1 | 0.012985 | 0.84 | PS 43:4 | MS1 |
| LIPID NEG | M851T461 | 850.5581 | 460.9 | 0.020011 | 0.84 | PC 38:6 | MS1 |
| LIPID NEG | M833T534 | 832.6064 | 534.3 | 0.012985 | 0.84 | PC 36:1 [PC 18:0/18:1] | MS1 and MS/MS |
| LIPIDS POS | M699T431 | 698.5556 | 431.4 | 0.02584 | 0.84 | HexCer 34:2;O2 | MS1 |
| HILIC NEG | M819T265 | 818.5693 | 265.5 | 0.0092013 | 0.84 | PC 39:6 ; PE 42:6 | MS1 |
| LIPID NEG | M858T520 | 857.6083 | 520.3 | 0.0029633 | 0.84 | PA 48:7 | MS1 |
| LIPIDS POS | M703T504 | 702.5416 | 504.2 | 0.043372 | 0.84 | PC O-31:2 | MS1 |
| HILIC POS | M858T256 | 857.5871 | 256.1 | 0.0031513 | 0.84 | PC 40:6 | MS1 |
| LIPID NEG | M729T518 | 728.5507 | 518.3 | 0.010176 | 0.83 | PE O-36:3 | MS1 |
| HILIC NEG | M690T303 | 689.5488 | 302.9 | 0.0013621 | 0.83 | SM d33:1 | MS1 |
| LIPIDS POS | M751T482 | 750.5422 | 482.2 | 0.037524 | 0.83 | PE O-38:6 / PA O-40:7 / PC O-33:3 / PE O-36:3 | MS1 |
| LIPID NEG | M729T535 | 728.5593 | 535.1 | 0.0052568 | 0.83 | PE O-36:2 [PE-O 18:1_18:1] | MS1 and MS/MS |
| LIPID NEG | M889T530 | 888.6764 | 530.2 | 0.019544 | 0.83 | HexCer 42:1;O4 | MS1 |
| LIPID NEG | M1179T532 | 1178.7763 | 531.7 | 0.03514 | 0.83 | Hex3Cer d42:2 | MS1 |
| LIPIDS POS | M930T623 | 929.8339 | 622.9 | 0.0046086 | 0.83 | TG 56:4 | MS1 |
| HILIC NEG | M740T299 | 739.5536 | 298.9 | 0.012933 | 0.83 | DG 40:7 ; SM d34:0 | MS1 |
| LIPIDS POS | M833T474 | 832.5834 | 473.8 | 0.011054 | 0.83 | PC 40:7 | MS1 |
| HILIC NEG | M725T232 | 724.5287 | 232.1 | 0.020501 | 0.83 | PE 36:4 [PE 18:2_18:2] | MS1 and MS/MS |
| HILIC NEG | M313T473 | 313.1134 | 473.2 | 0.00059882 | 0.83 | Galactosylglycerol | MS1 |
| HILIC POS | M833T257 | 832.5835 | 257.1 | 0.012047 | 0.83 | PC 38:4 ; PC 40:7 | MS1 |
| LIPIDS POS | M524T242 | 524.3708 | 242.1 | 0.014243 | 0.83 | LPC(18:0) | MS1 |
| HILIC POS | M145T418 | 145.0494 | 417.8 | 0.037993 | 0.83 | Hexenedioic acid; 3-Hydroxyadipic acid 3,6-lactone; 3-Methylglutaconic a | MS1 |
| HILIC POS | M834T257 | 833.5870 | 257.1 | 0.011431 | 0.83 | Coenzyme Q9 | MS1 |
| LIPID NEG | M847T521 | 846.5767 | 520.7 | 0.014603 | 0.83 | PC 38:3 [PC 18:0/20:3] | MS1 and MS/MS |
| LIPID NEG | M907T529 | 906.6320 | 529.0 | 9.06E-05 | 0.83 | SHexCer 42:1;O3 | MS1 |
| HILIC NEG | M176T403 | 176.0930 | 403.2 | 0.0097489 | 0.82 | Valine; Norvaline; N-Methyl-4-aminobutyric acid; N-Methyl-a-aminoisob | MS1 |
| LIPIDS POS | M1600T521 | 1600.2157 | 520.6 | 0.019555 | 0.82 | CL 80:0 | MS1 |
| LIPIDS POS | M825T526 | 824.5559 | 525.8 | 0.030916 | 0.82 | PC 36:2 / PE 39:2 / SHexCer 36:1;O3 / PE O-40:5 | MS1 |
| HILIC POS | M276T525 | 276.1189 | 525.4 | 0.023209 | 0.82 | 2'-O-Methyluridine; 3-Methyluridine; 5-Hydroxymethyl-2'-deoxyuridine; A | MS1 |
| HILIC POS | M229T496 | 229.1182 | 495.8 | 0.028572 | 0.82 | 3-Methoxytyrosine; 3-O-Methyl-a-methyldopa; Melevodopa; Methyldopa | MS1 |
| LIPID NEG | M881T499 | 880.5911 | 499.1 | 0.012985 | 0.82 | PC O-42:7 / Hex2Cer 32:0;O2 | MS1 |
| LIPIDS POS | M835T483 | 834.5987 | 483.4 | 0.011054 | 0.82 | PC 38:3 / PE 41:3 / PA 43:4 / PC 36:0 / PE 39:0 / PE 43:6 | MS1 |
| HILIC POS | M865T256 | 864.6442 | 256.2 | 0.028472 | 0.82 | PC 40:2 ; PC 42:5 | MS1 |
| LIPID NEG | M819T513 | 818.5304 | 513.3 | 0.039093 | 0.82 | PS O-40:7 / PE O-40:8 | MS1 |
| LIPID NEG | M864T493 | 863.5976 | 492.8 | 0.019323 | 0.82 | PC O-40:7 | MS1 |
| HILIC POS | M254T477_2 | 254.0799 | 476.6 | 0.039463 | 0.82 | Hydroxyoctenoylglycine; Propenoylcarnitine | MS1 |
| HILIC POS | M816T255 | 815.5752 | 254.7 | 0.028653 | 0.81 | PG 37:0 | MS1 |
| HILIC POS | M860T256 | 859.6026 | 255.7 | 0.018557 | 0.81 | PC 40:5 ; PC 42:8 ; PC 40:5 | MS1 |
| HILIC POS | M108T97_2 | 108.0477 | 96.6 | 0.034617 | 0.81 | (Methylthio)acetaldehyde | MS1 |
| HILIC NEG | M259T471 | 259.0130 | 470.8 | 0.0053956 | 0.81 | Glucose 6-sulfate; Galactose 6-sulfate | MS1 |
| HILIC POS | M607T113 | 607.2520 | 113.3 | 0.032527 | 0.81 | Bilirubin | MS1 and MS/MS |
| HILIC NEG | M266T406 | 266.0479 | 405.7 | 0.014536 | 0.81 | indole-3-acetyl-glycine | MS1 |
| HILIC NEG | M707T305 | 707.4906 | 304.7 | 0.012979 | 0.81 | SM d32:2 | MS1 |
| HILIC NEG | M750T225 | 749.5318 | 224.7 | 0.00081768 | 0.81 | PA 35:0 ; PG 34:0 | MS1 |
| HILIC NEG | M164T84 | 164.0357 | 84.4 | 0.041192 | 0.81 | 4-Pyridoxolactone; 5-Pyridoxolactone; Formylanthranilic acid; Noradrenoc | MS1 |
| HILIC POS | M178T464 | 178.0860 | 464.3 | 0.007741 | 0.81 | 1,2-Dehydrosalsolinol; 1,2,3,4-Tetrahydroisoquinoline-1-carboxylic acid; 1 | MS1 |
| HILIC POS | M408T305 | 408.3233 | 304.9 | 0.034617 | 0.81 | Stearoylcholine | MS1 |
| HILIC NEG | M348T485 | 348.0542 | 485.4 | 0.018736 | 0.81 | 2-Methylthioadenosine; (S)-5'-Deoxy-5'-(methylsulfinyl)adenosine | MS1 |
| HILIC NEG | M255T477 | 254.9819 | 477.0 | 0.0045777 | 0.81 | Ascorbic acid 2-sulfate; Ascorbic acid 3-sulfate | MS1 |
| LIPIDS POS | M818T541 | 817.6426 | 541.0 | 0.032298 | 0.81 | PC 37:2 / PE 40:2 | MS1 |
| LIPIDS POS | M973T643 | 972.8952 | 642.6 | 0.047037 | 0.81 | TG 59:3 [TG 18:1_18:1_23:1] | MS1 and MS/MS |
| HILIC NEG | M131T485_1 | 131.0828 | 485.3 | 0.0012366 | 0.81 | Ornithine | MS1 |
| HILIC NEG | M727T233 | 726.5445 | 232.6 | 0.00066167 | 0.81 | PE P-36:2 | MS1 |
| HILIC POS | M820T256 | 819.6080 | 256.2 | 0.012794 | 0.81 | PC P-38:3 ; PC P-40:6 ; PC 40:2 ; PC O-38:4 | MS1 |
| LIPID NEG | M730T518 | 729.5553 | 518.4 | 0.013644 | 0.81 | CerPE 38:2;O3 | MS1 |
| HILIC NEG | M174T485 | 174.0885 | 485.3 | 0.0015189 | 0.80 | Citrulline; Argininic acid; Carbamoyl (2R)-2,5-diaminopentanoate | MS1 |
| HILIC NEG | M146T107 | 146.0614 | 106.8 | 0.029552 | 0.80 | 3-Methyloxindole; 5-Methoxyindole; Cinnamamide; Indole-3-carbinol; Me | MS1 |
| LIPID NEG | M831T631 | 830.5911 | 631.3 | 0.015779 | 0.80 | PE 39:2 / PC 36:2 / PS 39:1 | MS1 |
| LIPIDS POS | M943T627 | 942.8477 | 626.7 | 0.02998 | 0.80 | TG 57:4 | MS1 |
| HILIC NEG | M283T153 | 283.0518 | 152.6 | 0.0045189 | 0.80 | 6-Thioinosine; 6-Thioinosinic acid | MS1 |
| LIPID NEG | M695T587 | 694.6715 | 587.5 | 0.011181 | 0.80 | Cer 44:0;O3 | MS1 |
| LIPID NEG | M723T491 | 722.5122 | 490.5 | 0.026726 | 0.80 | PE O-36:5 [PE O-16:1_20:4] | MS1 and MS/MS |
| HILIC NEG | M749T225 | 748.5284 | 224.8 | 0.034072 | 0.80 | PE 38:6 [PE 18:2_20:4] | MS1 and MS/MS |
| LIPID NEG | M790T491 | 790.5000 | 490.7 | 0.030921 | 0.80 | PS O-38:7 | MS1 |
| LIPID NEG | M837T535 | 836.6180 | 534.9 | 0.037401 | 0.80 | PE 43:4 | MS1 |
| LIPID NEG | M836T485 | 835.5678 | 485.0 | 0.032068 | 0.80 | PI O-35:1 / PG 37:1 | MS1 |
| HILIC POS | M536T302 | 536.4069 | 301.6 | 0.020357 | 0.80 | N-Nervonoyl Phenylalanine | MS1 |
| LIPID NEG | M873T555 | 872.6821 | 555.1 | 2.06E-05 | 0.80 | HexCer 42:1;O3 | MS1 |
| HILIC NEG | M860T308 | 859.5313 | 308.1 | 0.00075053 | 0.80 | PI 36:3 | MS1 |
| HILIC NEG | M264T328_2 | 264.1074 | 328.2 | 0.018946 | 0.80 | Tryptophan; Acetyl-N-formyl-5-methoxykynurenamine; di-Hydroxymelato | MS1 |
| HILIC NEG | M777T224 | 776.5581 | 223.7 | 0.00034954 | 0.80 | PE P-40:5 | MS1 |
| HILIC POS | M226T417 | 226.1046 | 417.4 | 0.038524 | 0.80 | Acetylcarnitine | MS1 and MS/MS |
| HILIC NEG | M215T463 | 215.1152 | 463.4 | 0.015108 | 0.79 | N-a-Acetyl-L-arginine | MS1 |
| HILIC POS | M725T195 | 724.5263 | 194.7 | 0.048412 | 0.79 | PE P-36:4 | MS1 |
| HILIC NEG | M796T267 | 795.5723 | 266.7 | 0.0012366 | 0.79 | PC 37:4 ; PE 40:4 | MS1 |
| LIPID NEG | M878T474 | 877.5774 | 474.5 | 0.0039774 | 0.79 | PC 40:7 / PS 43:6 | MS1 |
| HILIC POS | M244T334_2 | 244.1541 | 334.2 | 0.022337 | 0.79 | Dodecadienedioic acid; Tiglylcarnitine | MS1 |
| HILIC POS | M324T146 | 324.0590 | 145.9 | 0.018032 | 0.79 | Cytidine 2'-phosphate; Cytidine 3'-monophosphate; Cytidine monophosph | MS1 |
| HILIC NEG | M821T275 | 820.5638 | 275.3 | 0.0064638 | 0.79 | PC 36:2 [PC 18:0/18:2] | MS1 and MS/MS |
| HILIC NEG | M246T152 | 246.0814 | 151.5 | 0.00029022 | 0.79 | 5,6-Dihydrouridine | MS1 |
| HILIC NEG | M699T236 | 698.5135 | 235.7 | 0.0042656 | 0.79 | PE 34:3 [PE 16:1_18:2] | MS1 and MS/MS |
| LIPIDS POS | M753T502 | 752.5581 | 502.3 | 0.0046086 | 0.79 | PE O-38:5 / PE O-40:8 / PC O-33:2 / PE O-36:2 | MS1 |
| LIPIDS POS | M970T604 | 969.7288 | 603.6 | 0.03117 | 0.79 | TG 58:8 | MS1 |
| HILIC POS | M791T255 | 790.5737 | 255.5 | 0.020616 | 0.79 | PC O-38:7 | MS1 |
| LIPID NEG | M748T568 | 747.6313 | 567.9 | 0.037265 | 0.78 | CE 22:3 | MS1 |
| LIPID NEG | M778T512 | 777.5625 | 512.2 | 0.0063865 | 0.78 | PG 36:0 / PA 38:0 | MS1 |
| HILIC POS | M168T417 | 168.0346 | 417.4 | 0.045385 | 0.78 | Thioguanine | MS1 |
| HILIC NEG | M747T226 | 746.5135 | 226.4 | 0.0068333 | 0.78 | PE 38:7 [PE 18:3_20:4] | MS1 and MS/MS |
| HILIC POS | M147T525 | 147.0764 | 525.4 | 0.010129 | 0.78 | Glutamine | MS1 and RT |
| HILIC POS | M864T256 | 863.6329 | 256.4 | 0.0068233 | 0.78 | PC 40:3 ; PC 42:6 ; Galabiosylceramide d34:1 ; LacCer d34:1 | MS1 |
| HILIC POS | M150T469 | 150.0760 | 469.5 | 0.020357 | 0.78 | 2-Acetolactate; 2-C-Methyl-1,4-erythrono-D-lactone; 2-Deoxy-L-ribono-1, | MS1 |
| HILIC NEG | M752T225 | 751.5475 | 224.9 | 0.00020876 | 0.78 | PE P-38:4 | MS1 |
| HILIC NEG | M884T305 | 883.5327 | 304.6 | 0.000094016 | 0.78 | PI 36:5 [PI 16:0_20:5] | MS1 and MS/MS |
| LIPIDS POS | M731T523 | 730.5736 | 523.1 | 0.020255 | 0.78 | PC O-33:2 | MS1 |
| LIPID NEG | M750T492 | 749.5305 | 492.2 | 0.0063227 | 0.78 | PG 34:0 / PA 36:0 | MS1 |
| HILIC NEG | M723T228 | 722.5134 | 228.2 | 0.0021267 | 0.78 | PE 36:5 [PE 16:1_20:4] | MS1 and MS/MS |
| HILIC NEG | M284T107 | 284.2236 | 107.4 | 0.0000271 | 0.78 | Myristoylglycine | MS1 |
| HILIC NEG | M159T421 | 159.0299 | 421.4 | 0.0010031 | 0.77 | 3-Oxoadipic acid; Oxoadipic acid; Succinic anhydride; 2-Methyl-4-oxopent | MS1 |
| HILIC POS | M862T256 | 861.6181 | 255.8 | 0.017762 | 0.77 | PC 40:4 ; PC 42:7 | MS1 |
| LIPIDS POS | M775T474 | 774.5419 | 473.5 | 0.0049024 | 0.77 | PE O-38:5 / PE O-40:8 / PC O-33:2 / PE O-36:2 | MS1 |
| LIPIDS POS | M973T660 | 972.9317 | 659.9 | 0.044863 | 0.77 | TG 60:3e | MS1 |
| HILIC POS | M825T258 | 824.6131 | 258.0 | 0.0082767 | 0.77 | PA 44:5 ; PC 37:1 ; PE 40:1 ; PE 42:4 | MS1 |
| HILIC NEG | M748T226 | 747.5168 | 226.4 | 0.013906 | 0.77 | DG 42:10; PA 36:1; PG 34:1 | MS1 |
| HILIC NEG | M223T114 | 223.1344 | 114.0 | 0.01654 | 0.77 | 13-Oxo-9,11-tridecadienoic acid | MS1 |
| HILIC POS | M225T417 | 225.1208 | 417.2 | 0.032733 | 0.77 | N-Acetylisoputreanine; N(6)-Acetonyllysine; n2-Acetyl,n6-methyllysine | MS1 |
| LIPID NEG | M795T506 | 794.5703 | 506.2 | 0.029774 | 0.77 | PE 40:4 | MS1 |
| LIPID NEG | M751T513 | 750.5426 | 513.1 | 0.00053592 | 0.76 | PE O-38:5 | MS1 |
| HILIC POS | M762T195 | 762.4817 | 195.0 | 0.022216 | 0.76 | PE P-36:4 | MS1 |
| LIPIDS POS | M950T652 | 949.8557 | 651.7 | 0.011083 | 0.76 | FA 59:1;O3 | MS1 |
| LIPIDS POS | M838T500 | 837.6103 | 500.0 | 0.019555 | 0.76 | PC 39:6 / PE 42:6 | MS1 |

| | | | | | | | |
|---|---|---|---|---|---|---|---|
| LIPIDS POS | M777T495 | 776.5578 | 494.8 | 0.03101 | 0.76 | PE O-40:7 / PC O-35:4 / PE O-38:4 / PC O-33:1 / PE O-36:1 | MS1 |
| LIPIDS POS | M1006T652 | 1005.8815 | 651.9 | 0.037524 | 0.75 | TG 61:3 / TG 59:0 / TG 63:6 | MS1 |
| HILIC NEG | M245T328 | 245.0934 | 328.1 | 0.031575 | 0.75 | N-Acetyltryptophan; Indolepropionylglycine | MS1 |
| LIPID NEG | M773T483 | 772.5271 | 483.2 | 0.0021007 | 0.75 | PE O-40:8 | MS1 |
| LIPID NEG | M709T549 | 708.6142 | 549.1 | 0.0022052 | 0.75 | Cer t42:2 | MS1 |
| LIPIDS POS | M727T489 | 726.5417 | 488.5 | 0.020061 | 0.75 | PC O-31:1 / PE O-34:1 / PE O-36:4 / PA O-38:5 | MS1 |
| HILIC NEG | M775T223 | 774.5438 | 222.9 | 9.448E-07 | 0.75 | PE 40:7 [PE 18:1_22:6] | MS1 and MS/MS |
| HILIC NEG | M487T283 | 487.3284 | 283.4 | 0.042544 | 0.74 | Glyceryl lactooleate | MS1 |
| HILIC NEG | M101T65_1 | 101.0245 | 64.8 | 0.045696 | 0.74 | Methylmalonic acid semialdehyde; 2-Ketobutyric acid; 2-Methyl-3-oxopro | MS1 |
| HILIC NEG | M279T513 | 279.0725 | 512.9 | 0.0023147 | 0.74 | 3-Hydroxy-3-carboxymethyl-adipic acid | MS1 |
| LIPIDS POS | M1012T664 | 1011.8716 | 664.3 | 0.030916 | 0.74 | TG 60:1 | MS1 |
| HILIC NEG | M160T68 | 160.1063 | 67.6 | 0.019196 | 0.74 | 2-Hydroxycaprylic acid; 3-Hydroxyoctanoic acid; 4-hydroxyoctanoic acid; 5 | MS1 |
| LIPID NEG | M891T532 | 890.6358 | 532.0 | 0.019932 | 0.74 | SHexCer 42:1;O2 | MS1 |
| HILIC NEG | M256T535 | 256.0943 | 534.6 | 0.019766 | 0.74 | 2'-C-Methylcytidine; 2'-O-Methylcytidine; 3-Methylcytidine; 5-Methylcytid | MS1 |
| LIPIDS POS | M956T647 | 955.8084 | 647.0 | 0.041523 | 0.74 | TG 56:1 | MS1 |
| HILIC NEG | M148T479 | 148.0617 | 479.3 | 1.623E-09 | 0.74 | Alanine; beta-Alanine; 4-Amino-4-deoxyarabinose; (2S)-2-Amino-3-hydro | MS1 |
| HILIC POS | M818T254 | 817.5921 | 253.8 | 0.0013327 | 0.73 | PC P-38:4 ; PC P-40:7 ; PC O-38:5 ; PC P-38:4 | MS1 |
| HILIC NEG | M211T65 | 211.0590 | 64.8 | 0.0071261 | 0.73 | Cystamine; 8-Azaguanine | MS1 |
| HILIC NEG | M774T223 | 773.5318 | 222.5 | 0.014684 | 0.73 | PA 37:2 ; PG 36:2 | MS1 |
| LIPID NEG | M776T505 | 775.5467 | 504.6 | 6.45E-05 | 0.73 | PE O-40:7 | MS1 |
| HILIC NEG | M262T344 | 262.0394 | 344.3 | 0.00060389 | 0.73 | Epinephrine 3-sulfate; Epinephrine sulfate | MS1 |
| HILIC POS | M175T67 | 175.0477 | 67.0 | 0.038772 | 0.73 | N-Methyl-4-oxo-1,4-dihydropyridine-3-carboxamide; n-Methyl-6-oxo-1,6- | MS1 |
| LIPIDS POS | M951T614 | 950.8165 | 613.9 | 0.0239 | 0.73 | TG 58:7 [TG 18:1_18:1_22:5] | MS1 and MS/MS |
| HILIC POS | M855T254 | 854.5695 | 253.8 | 0.0020771 | 0.73 | PC 42:10 | MS1 |
| LIPIDS POS | M996T656 | 995.8893 | 656.0 | 0.011083 | 0.73 | TG 60:2 / TG 62:5 | MS1 |
| HILIC NEG | M144T339 | 144.1033 | 338.8 | 0.016518 | 0.73 | [(3-Methylbutyl)amino]acetic acid; 1-Methylpyrrolidine; 2-Aminoheptano | MS1 |
| HILIC NEG | M120T84 | 120.0457 | 84.5 | 0.0091331 | 0.73 | p-Aminobenzaldehyde; 2-Hydroxybenzyl alcohol; 3-Hydroxybenzyl alcohol | MS1 |
| HILIC POS | M542T323 | 542.3715 | 323.2 | 0.0029146 | 0.73 | N-Nervonoyl Histidine | MS1 |
| HILIC NEG | M128T384 | 128.0719 | 384.0 | 0.03924 | 0.73 | 2-Methylpyrrolidine-1-carboxylic acid; 1-Methylpyrrolidine-2-carboxylic a | MS1 |
| HILIC POS | M856T254 | 855.5729 | 253.8 | 0.0063877 | 0.73 | PC 40:7 ; PC 42:10 | MS1 |
| HILIC POS | M219T163 | 219.0086 | 163.2 | 4.34E-05 | 0.72 | 5-Methylthioribose | MS1 |
| LIPIDS POS | M992T648 | 991.8661 | 647.6 | 0.041143 | 0.72 | TG 60:3 / TG 58:0 / TG 62:6 | MS1 |
| HILIC NEG | M189T566 | 189.1247 | 565.8 | 0.0069166 | 0.72 | N-Acetylputrescine; (S)-2-Amino-4-methylpentanamide; L-Isoleucinamid | MS1 |
| HILIC POS | M137T342_2 | 137.0709 | 341.8 | 0.010955 | 0.72 | 1-Methylnicotinamide; 2-Acetyl-3-methylpyrazine; 2-Methylnicotinamide | MS1 |
| HILIC POS | M883T254 | 882.5992 | 254.1 | 8.74E-05 | 0.71 | PC 42:7 ; PC 44:10 | MS1 |
| HILIC POS | M893T253 | 892.5237 | 253.4 | 0.048412 | 0.71 | PC 42:10 | MS1 |
| HILIC NEG | M162T486 | 162.0774 | 485.7 | 0.00092603 | 0.71 | 2-Aminoisobutyric acid; 3-Aminobutanoic acid; 3-Aminoisobutanoic acid; | MS1 |
| LIPID NEG | M391T84 | 391.2854 | 83.6 | 0.011876 | 0.71 | ST 24:1;O4 / ST 23:1;O2 / FA 23:4 | MS1 |
| LIPIDS POS | M996T664 | 995.8981 | 664.2 | 0.006129 | 0.71 | TG 60:1 / TG 62:4 | MS1 |
| HILIC NEG | M773T223 | 772.5285 | 222.5 | 5.7892E-06 | 0.71 | PE 40:8 [PE 18:2_22:6] | MS1 and MS/MS |
| HILIC NEG | M155T151 | 155.0465 | 151.4 | 0.026866 | 0.71 | 4-Imidazolone-5-propionic acid; 5-Hydroxymethyl-4-methyluracil; 5-Hydro | MS1 |
| LIPIDS POS | M776T473 | 775.5457 | 473.4 | 0.00022563 | 0.71 | PG 36:2 | MS1 |
| HILIC NEG | M130T453 | 130.0512 | 452.6 | 0.000035223 | 0.70 | 5-Aminolevulinic acid; Glutaramic acid; L-Glutamic gamma-semialdehyd | MS1 |
| HILIC NEG | M246T497 | 246.1098 | 497.4 | 0.016652 | 0.70 | 6-Guanidino-2-oxocaproic acid | MS1 |
| LIPID NEG | M607T366 | 607.3852 | 366.5 | 0.00039435 | 0.70 | ST 27:1;O;GlcA | MS1 |
| HILIC NEG | M158T317 | 158.1189 | 316.9 | 0.014684 | 0.70 | 8-Aminooctanoic acid; DL-2-Aminooctanoic acid; Pregabalin; Propionylch | MS1 |
| LIPID NEG | M879T484 | 878.5896 | 484.2 | 0.00061335 | 0.70 | PC 40:6 [PC 18:0/22:6] | MS1 and MS/MS |
| HILIC NEG | M201T464 | 201.1248 | 464.3 | 0.00028125 | 0.70 | N-Acetylisoputreanine; n2-Acetyl,n6-methyllysine | MS1 |
| HILIC NEG | M124T119 | 124.0406 | 119.0 | 0.00029022 | 0.70 | 3-Aminobenzene-1,2-diol | MS1 |
| LIPID NEG | M797T535 | 796.5472 | 535.3 | 0.037265 | 0.70 | PS O-38:4 / PE O-38:5 | MS1 |
| LIPID NEG | M591T367 | 591.3902 | 367.2 | 0.020463 | 0.69 | ST 27:2;O;Hex | MS1 |
| HILIC NEG | M777T233 | 776.5232 | 233.3 | 0.010168 | 0.69 | PC 36:6 | MS1 |
| LIPIDS POS | M1003T660 | 1002.9413 | 659.8 | 0.028392 | 0.69 | TG 61:2 [TG 25:1_18:0_18:1] | MS1 and MS/MS |
| HILIC NEG | M776T223 | 775.5475 | 222.9 | 3.6092E-10 | 0.69 | PA 38:1 ; PG 36:1 | MS1 |
| HILIC POS | M132T443 | 132.0659 | 442.7 | 0.019844 | 0.69 | 4-Hydroxyproline | MS/MS and RT |
| HILIC NEG | M702T235 | 701.5321 | 235.4 | 0.0022092 | 0.69 | PE P-34:1 | MS1 |
| LIPIDS POS | M1000T643 | 999.9122 | 643.4 | 0.017518 | 0.69 | FA 63:1;O3 | MS1 |
| LIPID NEG | M741T383 | 741.3804 | 382.9 | 0.011513 | 0.69 | PI 25:1;O2 | MS1 |
| LIPID NEG | M818T492_1 | 817.5186 | 492.4 | 0.00065646 | 0.68 | PG O-38:5 | MS1 |
| HILIC POS | M130T548 | 130.0498 | 548.1 | 0.008809 | 0.68 | 1-pyrroline-3-hydroxy-5-carboxylic Acid; 1-Pyrroline-4-hydroxy-2-carboxyl | MS1 |
| HILIC NEG | M299T71 | 299.1140 | 71.1 | 0.00013713 | 0.67 | 3-(3,4,5-Trimethoxyphenyl)propanoic acid | MS1 |
| HILIC NEG | M187T67 | 187.1342 | 67.0 | 0.01431 | 0.67 | 3-Hydroxydecanoic acid; 10-Hydroxydecanoic acid; 2-Hydroxydecanoic aci | MS1 |
| LIPID NEG | M900T443 | 899.5622 | 443.3 | 0.024669 | 0.67 | PI 39:4 | MS1 |
| LIPIDS POS | M1022T664 | 1021.9132 | 664.5 | 0.026524 | 0.67 | TG 62:2 / TG 64:5 | MS1 |
| HILIC POS | M431T107 | 431.3515 | 107.3 | 0.0031592 | 0.67 | ST 28:2;O3 | MS1 |
| HILIC POS | M881T254 | 880.5831 | 254.3 | 0.017671 | 0.67 | PC 42:8 ; PC 44:11 | MS1 |
| HILIC NEG | M720T229 | 720.4977 | 229.3 | 0.010423 | 0.66 | PE P-36:5 | MS1 |
| HILIC NEG | M183T328 | 183.1393 | 328.2 | 0.001136 | 0.66 | Undecenoic acid; 3-Methyl-3-decenoic acid; 3-Methyl-4-decenoic acid | MS1 |
| LIPID NEG | M459T84 | 459.2732 | 83.6 | 0.0096696 | 0.66 | ST 27:4;O6 | MS1 |
| HILIC NEG | M276T294 | 276.0549 | 293.6 | 0.0022041 | 0.66 | Tyramine-O-sulfate; Metanephrine sulfate | MS1 |
| HILIC NEG | M139T94 | 139.0073 | 93.5 | 0.036255 | 0.66 | Methanesulfinic acid | MS1 |
| HILIC POS | M308T279 | 308.1853 | 278.8 | 0.0097645 | 0.65 | 3-Polyprenyl-4,5-dihydroxybenzoate | MS1 |
| LIPID NEG | M826T538 | 825.6632 | 538.3 | 0.0015585 | 0.65 | HexCer 36:1;O / HexCer 42:2;O3 | MS1 |
| LIPIDS POS | M674T659 | 673.6496 | 659.4 | 0.012579 | 0.64 | FA 43:0;O | MS1 |
| LIPID NEG | M712T565 | 711.6332 | 565.4 | 7.04E-05 | 0.64 | CE 19:0 | MS1 |
| LIPIDS POS | M590T634 | 589.5552 | 633.6 | 0.00064051 | 0.64 | FA 37:0;O | MS1 |
| HILIC POS | M134T423 | 134.0811 | 422.9 | 0.025029 | 0.63 | 2-Oxovaleric acid; 3-Hydroxynorvaline; 3-Oxopentanoic acid; 1,4-Dideoxy- | MS1 |
| HILIC POS | M280T311 | 280.1541 | 311.4 | 0.015478 | 0.63 | phenylacetylcarnitine; Feruloylcholine | MS1 |
| LIPIDS POS | M1039T665 | 1038.8916 | 664.5 | 0.015546 | 0.62 | TG 62:2 | MS1 |
| LIPIDS POS | M1036T656 | 1035.8713 | 656.0 | 0.0097393 | 0.62 | TG 62:3 | MS1 |
| HILIC POS | M305T523 | 305.1454 | 523.0 | 0.017546 | 0.61 | N-Ribosylhistidine | MS1 |
| HILIC NEG | M211T296 | 211.1706 | 295.7 | 0.033619 | 0.61 | Tridecenoic acid | MS1 |
| HILIC POS | M158T386 | 158.1175 | 385.5 | 0.00339 | 0.61 | Octadienoic acid | MS1 |
| HILIC POS | M307T304 | 307.2629 | 303.5 | 0.010677 | 0.61 | Eicosatrienoic acid | MS1 |
| HILIC POS | M105T98 | 105.0368 | 97.9 | 0.018557 | 0.60 | (Methylthio)acetone | MS1 |
| LIPIDS POS | M844T445 | 843.5118 | 444.9 | 0.0061021 | 0.60 | PG 38:4 / PG 40:7 | MS1 |
| HILIC POS | M144T463 | 144.1019 | 463.0 | 0.00031984 | 0.59 | (2S)-1,2-Dimethylpyrrolidine-2-carboxylic acid; (R)-1-Methylpiperidine-2- | MS1 |
| LIPID NEG | M684T549 | 683.6012 | 549.1 | 9.39E-05 | 0.59 | CE 17:0 | MS1 |
| HILIC POS | M197T452 | 197.0895 | 451.9 | 0.021316 | 0.59 | N2-Acetylornithine; N5-Acetylornithine | MS1 and MS/MS |
| HILIC NEG | M222T393 | 222.0985 | 392.9 | 0.015108 | 0.57 | 2-Amino-4-ethoxy-3-hydroxybutanoic acid | MS1 |
| HILIC POS | M222T180 | 222.0196 | 180.2 | 0.021316 | 0.57 | Choline sulfate | MS1 |
| LIPID NEG | M637T549 | 636.5928 | 549.0 | 3.92E-05 | 0.56 | Cer 40:1;O3 | MS1 |
| HILIC POS | M778T185 | 777.5598 | 185.0 | 0.0037004 | 0.56 | PE P-38:3 ; PE P-40:6 ; PE O-38:4 | MS1 |
| LIPID NEG | M766T558 | 765.6045 | 557.6 | 6.83E-05 | 0.56 | TG 46:6 | MS1 |
| LIPIDS POS | M1021T656 | 1020.9003 | 656.2 | 0.037524 | 0.56 | TG 63:7 | MS1 |
| HILIC POS | M188T424 | 188.1279 | 424.4 | 0.021782 | 0.53 | 3-hydroxynonadienoic acid; 8-Amino-7-oxononanoic acid; N-Heptanoylgly | MS1 |
| LIPIDS POS | M835T625 | 834.7909 | 625.1 | 0.0086928 | 0.48 | TG 50:2e | MS1 |
| LIPIDS POS | M1018T648 | 1017.8820 | 647.6 | 0.0092915 | 0.48 | TG 62:4 / TG 60:1 / TG 64:7 | MS1 |
| HILIC NEG | M217T84 | 216.9815 | 84.4 | 0.031023 | 0.46 | 3-hydroxybenzoic acid-3-O-sulphate; 4-hydroxybenzoic acid-4-O-sulphate | MS1 |
| HILIC POS | M208T274 | 208.1330 | 274.5 | 0.0026344 | 0.43 | Cinnamyl propionate; Isopropyl cinnamate; Phenylalanine betaine; Prenyl | MS1 |
