## Supplementary material for "Glutamine antagonism suppresses tumor growth in adrenocortical carcinoma through inhibition of de novo nucleotide biosynthesis": Suppl. Table 5

| **Pathway Name** | **Match Status** | **FDR** | Metabolites in Pathway (matches in red) |
|---|---|---|---|
| Arginine and proline metabolis | 11/36 | 1.06E-04 | L-Arginine; N(omega)-Hydroxyarginine; Guanidinoacetate; Creatine; 4-Guanidinobutanoate; **4-Aminobutanoate**; Putrescine; **4-Aminobutyraldehyde**; S-Adenosylmethioninamine; S-Adenosyl-L-methionine; Spermidine; **N-Acetylputrescine**; **N4-Acetylaminobutanal**; Spermine; trans-3-Hydroxy-L-proline; D-Proline; (4R)-4-Hydroxy-2-oxoglutarate; L-erythro-4-Hydroxyglutamate; **L-4-Hydroxyglutamate semialdehyde**; **L-1-Pyrroline-3-hydroxy-5-carboxylate**; **Hydroxyproline**; L-Proline; **L-Glutamyl 5-phosphate**; **L-Glutamate**; **L-Glutamate 5-semialdehyde**; **L-Ornithine**; Peptide; Nitric oxide; Phosphocreatine; Agmatine; 4-Acetamidobutanoate; Homocarnosine; 1-Pyrroline-2-carboxylate; Glyoxylate; Pyruvate; (S)-1-Pyrroline-5-carboxylate |
| Valine, leucine and isoleucine b | 5/8 | 8.52E-04 | **L-Threonine**; **(S)-3-Methyl-2-oxopentanoic acid**; L-Leucine; **3-Methyl-2-oxobutanoic acid**; **2-Oxobutanoate**; L-Isoleucine; 4-Methyl-2-oxopentanoate; **L-Valine** |
| Phenylalanine metabolism | 4/8 | 0.0096787 | Phenylacetaldehyde; **L-Phenylalanine**; Phenethylamine; **Phenylpyruvate**; Phenylacetic acid; 2-Hydroxyphenylacetate; **2-Hydroxy-3-phenylpropenoate**; **L-Tyrosine** |
| Arginine biosynthesis | 5/14 | 0.0096787 | **L-Glutamate**; L-Arginine; **N-Acetylornithine**; N-(L-Arginino)succinate; L-Aspartate; **L-Citrulline**; Carbamoyl phosphate; **L-Ornithine**; Ammonia; **L-Glutamine**; 2-Oxoglutarate; N-Acetyl-L-glutamate; Urea; Fumarate |
| Phenylalanine, tyrosine and tryp | 3/4 | 0.0096787 | **Phenylpyruvate**; **L-Phenylalanine**; **L-Tyrosine**; 3-(4-Hydroxyphenyl)pyruvate |
| Pyrimidine metabolism | 8/39 | 0.011617 | UDP; **L-Glutamine**; Carbamoyl phosphate; (S)-Dihydroorotate; Orotidine 5'-phosphate; UMP; UTP; **Uridine**; 5,6-Dihydrouracil; **3-Ureidopropionate**; CTP; CDP; **CMP**; **Cytidine**; dCDP; dCMP; dCTP; Deoxycytidine; dUTP; dUDP; dUMP; **Deoxyuridine**; dTDP; dTTP; dTMP; Thymidine; **Thymine**; (R)-5,6-Dihydrothymine; (R)-3-Ureidoisobutyrate; P1,P4-Bis(5'-uridyl) tetraphosphate; 2'-Deoxy-5-hydroxymethylcytidine-5'-diphosphate; N-Carbamoyl-L-aspartate; Orotate; 5-Phospho-alpha-D-ribose 1-diphosphate; Uracil; **beta-Alanine**; 2-Deoxy-D-ribose 1-phosphate; (R)-3-Amino-2-methylpropanoate; 2'-Deoxy-5-hydroxymethylcytidine-5'-triphosphate |
| Glycine, serine and threonine m | 7/33 | 0.01732 | L-Serine; Choline; Betaine aldehyde; **Betaine**; Guanidinoacetate; 3-Phospho-D-glycerate; **N,N-Dimethylglycine**; L-Cystathionine; Glycine; **O-Phospho-L-serine**; **Sarcosine**; 5,10-Methylenetetrahydrofolate; **L-Threonine**; Lipoylprotein; Aminoacetone; D-Glycerate; S-Aminomethyldihydrolipoylprotein; Tetrahydrofolate; Dihydrolipoylprotein; 2-Phospho-D-glycerate; D-Serine; Hydroxypyruvate; Creatine; 3-Phosphonooxypyruvate; L-Cysteine; **2-Oxobutanoate**; Glyoxylate; L-2-Amino-3-oxobutanoic acid; Pyruvate; CO2; **5-Aminolevulinate**; Methylglyoxal; Ammonia |
| Alanine, aspartate and glutam | 6/28 | 0.028191 | N-Acetyl-L-aspartate; 2-Oxosuccinamate; L-Aspartate; **L-Asparagine**; D-Aspartate; N-(L-Arginino)succinate; N6-(1,2-Dicarboxyethyl)-AMP; **L-Alanine**; **Succinate semialdehyde**; **L-Glutamate**; **4-Aminobutanoate**; **L-Glutamine**; Ammonia; 2-Oxoglutaramate; (S)-1-Pyrroline-5-carboxylate; N-Acetylaspartylglutamate; N-Acetylaspartylglutamylglutamate; Citrate; Oxaloacetate; Fumarate; Pyruvate; N-Carbamoyl-L-aspartate; Succinate; 2-Oxoglutarate; Carbamoyl phosphate; D-Glucosamine 6-phosphate; 5-Phosphoribosylamine; beta-Citryl-L-glutamate |
