## Supplementary material for "Glutamine antagonism suppresses tumor growth in adrenocortical carcinoma through inhibition of de novo nucleotide biosynthesis": Suppl. Methods, Figures and Table Legends

### Supplementary Data

#### Supplementary Methods

##### Untargeted serum metabolome analysis

The samples were aliquoted on the day of collection and stored locally at -80°C. The samples were transported on dry ice to the University of Birmingham, United Kingdom, and stored at -80°C prior to untargeted serum metabolome profiling.

##### Sample Extraction

Serum samples: Samples were randomised using the RAND() function in Microsoft Excel to determine the sample preparation order. This order was assessed to ensure that all the metadata (age, sex, body mass index, and disease classification) was randomised and there were no patterns/correlations with the sample preparation order. Serum samples were thawed on ice on the day of preparation. For the HILIC assay, plasma (50 µL) and 1/1 acetonitrile/methanol (v/v) (150 µL) were mixed by vortex (120 s) and centrifuged (17,000 g, 20 min, 4°C). For lipid assays, plasma (50 µL) and isopropanol (150 µL) were mixed by vortex (120 s) and centrifuged (17,000 g, 20 min, 4°C).

Pooled QC samples: 120 µL serum aliquots from all biological samples were transferred to a single tube and vortex mixed. This single tube sample was called the pooled quality control (QC) sample. Multiple 50 µL aliquots were prepared as described for serum samples.

Process blank samples: Blank extraction solutions were prepared by performing the extraction protocol as described above for serum samples with no serum present.

Ultra High Performance Liquid Chromatography-Mass Spectrometry: All of the three types of samples (biological, QC and blank) were analyzed by applying two UHPLC–MS methods in positive and negative ion modes separately using a Dionex UltiMate 3000 UHPLC system coupled with a heated electrospray Q Exactive Focus mass spectrometer (Thermo Fisher Scientific). Analysis order for the samples was randomized, and all of the samples were analyzed in two analytical batches. Ten QC samples were injected at the start of each assay/ion mode to condition the analytical system and were then injected after every 6th biological sample, with two QC samples being analyzed at the end of each batch. The blank sample was analyzed as the 5th and as the final injection of each batch.

HILIC assays: These assays used an Accucore150-Amide-HILIC column (100 x 2.1mm, 2.6 µm, ThermoFisher Scientific). For positive ion analysis, mobile phase A was 10 mM ammonium formate dissolved in acetonitrile/water/formic acid (95:4.9:0.1 (v/v)) and mobile phase B was 10 mM ammonium formate dissolved in acetonitrile/water/formic acid (50/49.9/0.1 (v/v)). For negative ion analysis, mobile phase A was 10 mM ammonium acetate dissolved in acetonitrile/water/acetic acid (95:4.9:0.1 (v/v)) and mobile phase B was 10 mM ammonium acetate dissolved in acetonitrile/water/acetic acid (50/49.9/0.1 (v/v)). The gradient elution applied for positive and negative ion mode was t=0.0, 1% B; t=1.0, 1% B; t=3.0, 15% B; t=6.0, 50% B; t=9.0, 95% B; t=10.0, 95% B; t=10.5, 1% B; t=14.0, 1% B. All changes were linear (curve = 5) and the flow rate was 0.50 $mL.min^{-1}$. Column temperature was 35°C and injection volume was 2 µL. Data were acquired in positive and negative ionization modes separately (70 – 1050 m/z) with a mass resolution 70,000 (FWHM, m/z 200). Ion source parameters applied were Sheath gas = 55 arbitrary units, Aux gas = 35 arbitrary units, Sweep gas = 4 arbitrary units, Spray Voltage = 3.2kV (positive ion) and 2.7kV (negative ion), Capillary temp. = 380°C, Aux gas heater temp. = 440°C. Thermo Exactive Tune (2.8 SP1, build 2806) software controlled the instruments and data acquisition. All data were collected as MS1 data in the Profile mode setting with the exception of five QC sample injections (injections 6-10) where MS/MS data were collected in the "Discovery mode" setting over different precursor m/z ranges (70−210 m/z; 200−310 m/z; 300−410 m/z; 400-510 m/z; 500-1050 m/z) using stepped normalized collision energies (positive ion mode: 20, 40, 100%; negative ion mode: 40, 60, 130%). All samples were maintained at a temperature of 4°C in the autosampler. Instrument maintenance (source and column cleaning) was performed between the end of batch 1 and start of batch 2.

Lipid assays: These assays used a reversed-phase Hypersil GOLD $C_{18}$ column (100 x 2.1 mm, 1.9μm; Thermo Fisher Scientific). Mobile phase A was 10 mM ammonium formate dissolved in acetonitrile/water/formic acid (60:39.9:0.1 (v/v)) and mobile phase B was 10 mM ammonium formate dissolved in isopropanol/acetonitrile/water/formic acid (85.5/9.5/4.9/0.1 (v/v)). The gradient elution applied was t=0.0, 20% B; t=0.5, 20% B, t=8.5, 100% B; t=9.5, 100% B; t=11.5, 20% B; t=14.0, 20% B. All changes were linear (curve = 5) and the flow rate was 0.40 mL/min. Column temperature was 55 °C and injection volume was 2 μL. Data were acquired in positive and negative ionization mode separately (150 – 2000 *m/z*) with a mass resolution 70,000 (FWHM, m/z 200). Ion source parameters applied were Sheath gas = 48 arbitrary units, Aux gas = 15 arbitrary units, Sweep gas = 0 arbitrary units, Spray Voltage = 3.2kV (positive ion) / 2.7kV (negative ion), Capillary temp. = 380°C, Aux gas heater temp. = 450°C. Thermo Exactive Tune (2.8 SP1, build 2806) software controlled the instruments and data acquisition. All data were acquired in Profile mode. All data were collected as MS1 data in profile mode with the exception of five QC sample injections (injections 6-10) where MS/MS data were collected in Discovery mode over different precursor m/z ranges (150−510 m/z; 500−710 m/z; 700−860 m/z; 850−1010 m/z; 1000−2000 m/z) using stepped normalized collision energies (positive ion mode: 20, 40, negative ion mode: 40, 60, 130%). All samples were maintained at a temperature of 4°C in the autosampler. Instrument maintenance (source and column cleaning) was performed between the end of batch 1 and start of batch 2.

Raw data processing: Vendor format raw data files (.RAW) were converted to the mzML file format using the open-source ProteoWizard software[1]. Deconvolution was performed by the R package XCMS (version 3.6.1 running in the Galaxy environment)[2]. The R package IPO[3] was used to optimise and obtain XCMS peak picking parameters for each of the different assays:

(a) HILIC negative ion mode assay: min peak width = 6.23; max peak width = 40; ppm = 9.98; mzdiff = -0.02025; bw = 0.25; mzwid = 0.001; minfrac = 0.5; snr_thr = 10.

(b) HILIC positive ion mode assay: min peak width = 5.70; max peak width = 40; ppm = 7.0; mzdiff = 0.00021; bw = 0.25; mzwid = 0.0059; minfrac = 0.5; snr_thr = 10.

(c) Lipids negative and positive ion mode assays: min peak width = 5.44; max peak width = 40; ppm = 6.5; mzdiff = -0.0065; bw = 0.25; mzwid = 0.04265; minfrac = 0.5; snr_thr = 10.

A data matrix of metabolite features (i.e., m/z-retention time pairs) versus samples was constructed for each of four assay types (HILIC-negative, HILIC-positive, Lipids-negative, Lipids-positive).

Data quality assessment and filtering: Each of the four data matrices constructed by raw data processing was filtered based on the data collected for QC and blank samples[4]. Metabolite features were retained in the data matrix if they were: present in >90% of all the QC samples, had a peak area relative standard deviation (RSD) <30% across all the QC samples (QC11 onwards) and had an extract blank/mean QC area ratio of <5%.

Metabolite annotation: Metabolite annotation was performed applying the Python package BEAMSpy https://github.com/computational-metabolomics/beamspy (RT diff = 2 s, Pearson correlation >0.50; m/z values of all experimentally observed peaks were searched against LIPIDMAPS[5] and HMDB[6], all matches < 5ppm mass error tolerance were reported). To generate more robust compound annotations using MS/MS data, QC sample UHPLC-MS/MS data were matched to MS/MS databases using the LipidSearch software (lipid assay metabolite annotation; version 4.2.18, Thermo Fisher Scientific) and were searched against the entire in-silico HCD MS/MS database with 5ppm mass error. HILIC MS/MS data were searched in mzCloud using Compound Discoverer (v3.1, Thermo Fisher Scientific) using a >70% match as the criteria for a successful annotation. RT data were matched to an in-house RT library constructed with authentic chemical standards, matches within ± 5 seconds were applied as the criteria for a successful match. LipidSearch and mzCloud annotations were aligned to the XCMS outputs using the R programming language (https://www.R-project.org), applying a 5ppm mass error for the MS1 data and a +/- 5 second retention time tolerance window. All annotations described are reported to either level 2 or 3 as defined by the Metabolomics Standards Initiative[7].

Pathway enrichment analysis: Pathway enrichment analysis was performed in the software MetaboAnalyst[8] for HILIC based results. Pathway enrichment analysis applied pathway analysis, scatter plot for visualization method, hypergeometric test for enrichment method, relative-betweenness centrality for topology analysis and Homo sapiens (KEGG) as the pathway library (accessed on 13/05/2024). Only pathways with a p-value (after FDR correction) < 0.05 and with a minimum of five metabolites in the pathway listed as biologically important were reported. For lipids based results, lipids were classified according to LipidMaps ontologies[5] manually.

#### REFERENCES


1. Chambers MC, Maclean B, Burke R, et al. A cross-platform toolkit for mass spectrometry and proteomics. Nat Biotechnol 2012;30(10):918-20. DOI: 10.1038/nbt.2377.

2. Smith CA, Want EJ, O'Maille G, Abagyan R, Siuzdak G. XCMS: processing mass spectrometry data for metabolite profiling using nonlinear peak alignment, matching, and identification. Anal Chem 2006;78(3):779-87. DOI: 10.1021/ac051437y.

3. Libiseller G, Dvorzak M, Kleb U, et al. IPO: a tool for automated optimization of XCMS parameters. BMC Bioinformatics 2015;16:118. DOI: 10.1186/s12859-015-0562-8.

4. Lloyd GR, Jankevics A, Weber RJM. struct: an R/Bioconductor-based framework for standardized metabolomics data analysis and beyond. Bioinformatics 2021;36(22-23):5551-5552. DOI: 10.1093/bioinformatics/btaa1031.

5. Conroy MJ, Andrews RM, Andrews S, et al. LIPID MAPS: update to databases and tools for the lipidomics community. Nucleic Acids Res 2024;52(D1):D1677-D1682. DOI: 10.1093/nar/gkad896.

6. Wishart DS, Guo A, Oler E, et al. HMDB 5.0: the Human Metabolome Database for 2022. Nucleic Acids Res 2022;50(D1):D622-D631. DOI: 10.1093/nar/gkab1062.

7. Sumner LW, Amberg A, Barrett D, et al. Proposed minimum reporting standards for chemical analysis Chemical Analysis Working Group (CAWG) Metabolomics Standards Initiative (MSI). Metabolomics 2007;3(3):211-221. DOI: 10.1007/s11306-007-0082-2.

8. Pang Z, Lu Y, Zhou G, et al. MetaboAnalyst 6.0: towards a unified platform for metabolomics data processing, analysis and interpretation. Nucleic Acids Res 2024;52(W1):W398-W406. DOI: 10.1093/nar/gkae253.

Suppl. Fig. 1

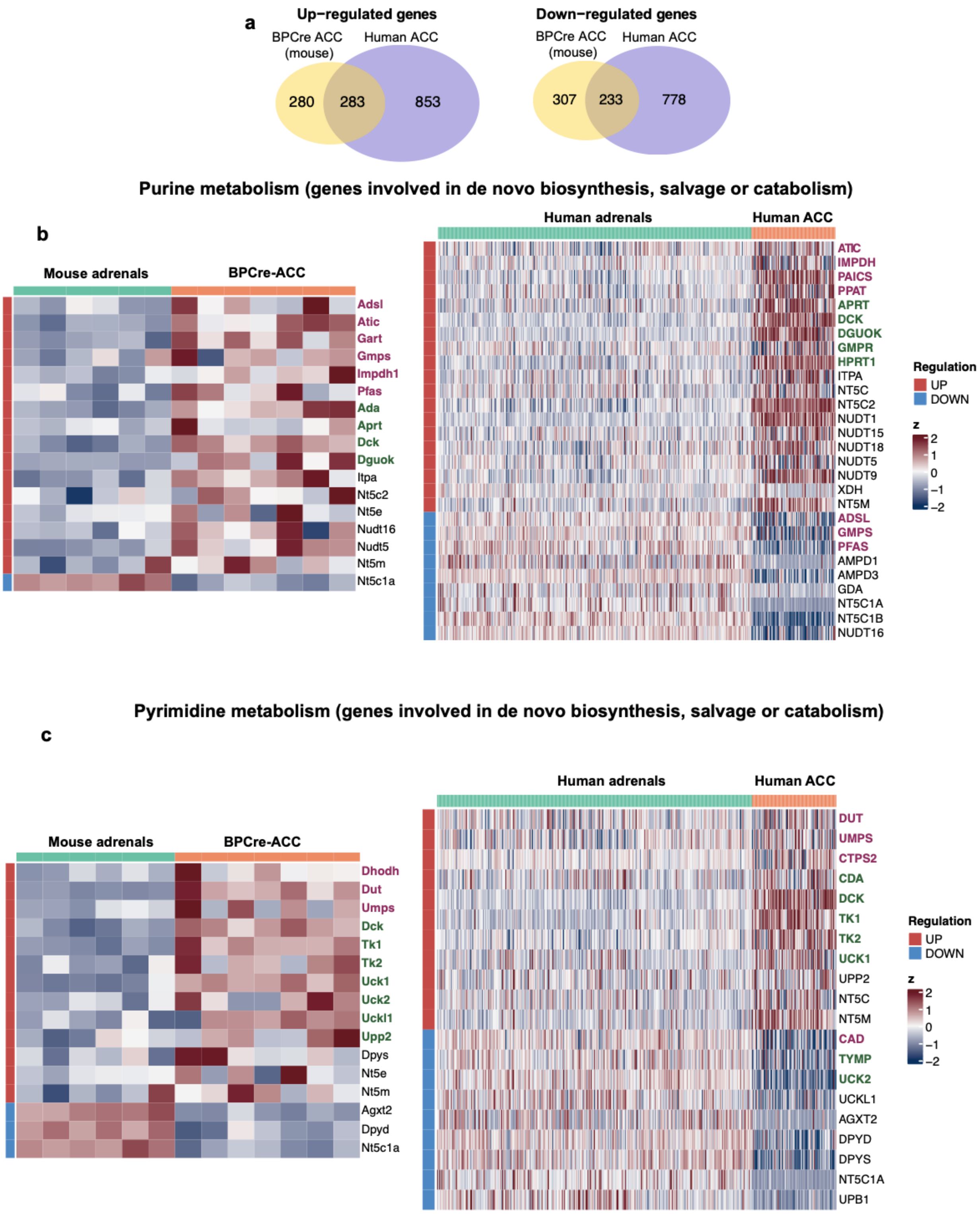


**Suppl. Fig. 1 (Related to Fig. 1): Cross-species comparison of transcriptional metabolic dysregulation in ACC (purine and pyrimidine metabolism). a** Venn diagrams showing overlap in significantly up-regulated and down-regulated metabolic genes between human and mouse (BPCre) ACC tumors, as assessed by RNA-seq. Mouse tissue: 6 control adrenals (no mutations) and 7 BPCre ACC tumors. Human tissue: 79 ACC tumors (TCGA) and 295 healthy adrenals (GTEx). **b-c** Heat-maps illustrating significant expression changes in genes involved in *de novo* and salvage purine (b) and pyrimidine (c) biosynthesis and catabolism, as assessed by RNA-seq. Genes involved in *de novo* nucleotide biosynthesis are highlighted in bold purple font. Genes involved in salvage biosynthesis are highlighted in bold green font.

#### Suppl. Fig. 2

**a** Amino-sugar and nucleotide sugar metabolism

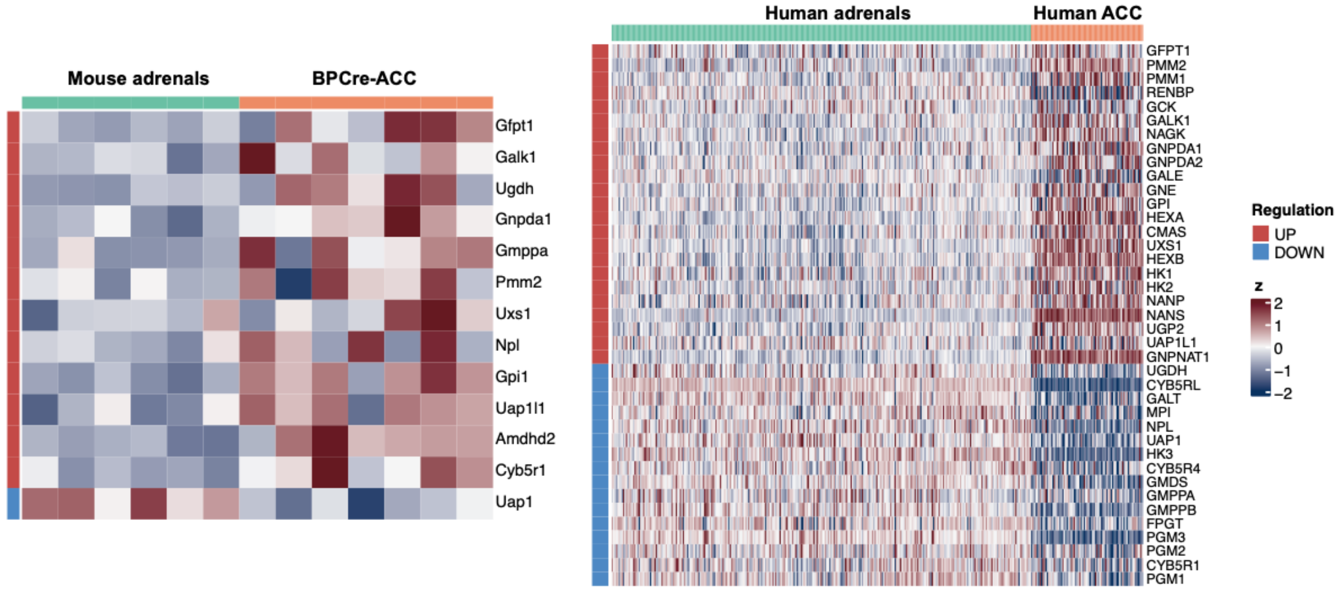


**b** Glutathione metabolism

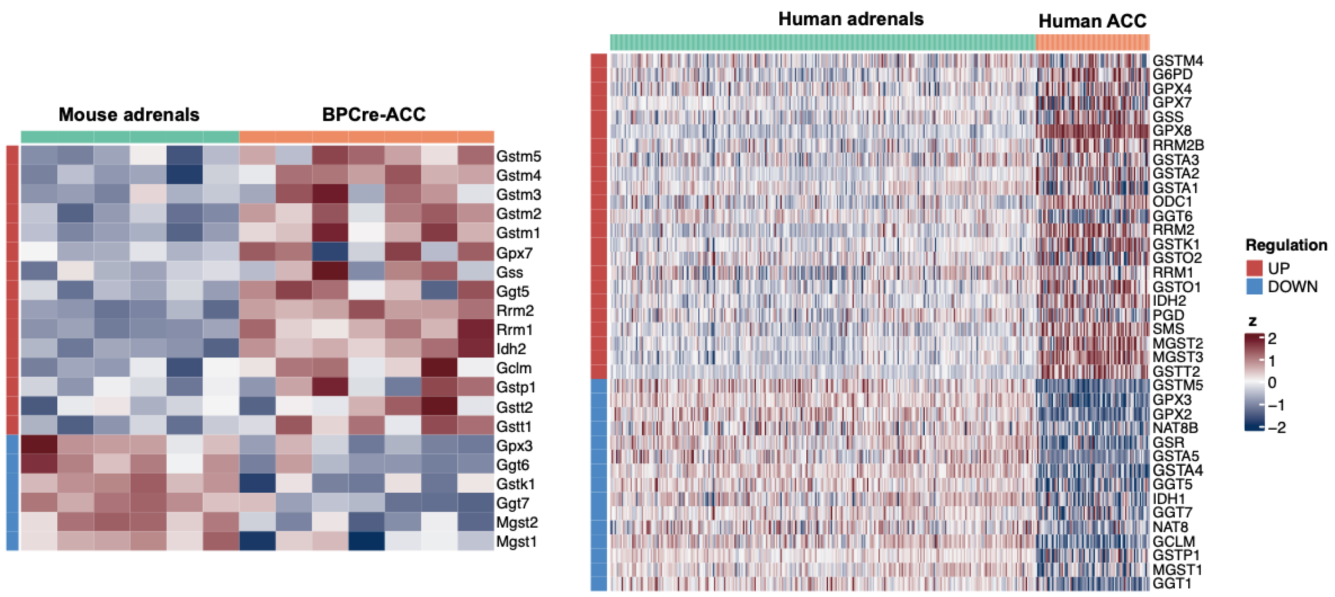

**Suppl. Fig. 2 (Related to Fig. 1): Cross-species comparison of transcriptional metabolic dysregulation in ACC (amino sugar and nucleotide sugar metabolism, glutathione metabolism).** Heat-maps illustrating significant expression changes in genes involved in amino sugar and nucleotide sugar metabolism (**a**) and glutathione metabolism (**b**) in mouse and human ACC, as assessed by RNA-seq. Mouse tissue: 6 control adrenals (no mutations) and 7 BPCre ACC tumors. Human tissue: 79 ACC tumors (TCGA) and 295 healthy adrenals (GTEx).

Suppl. Fig. 3

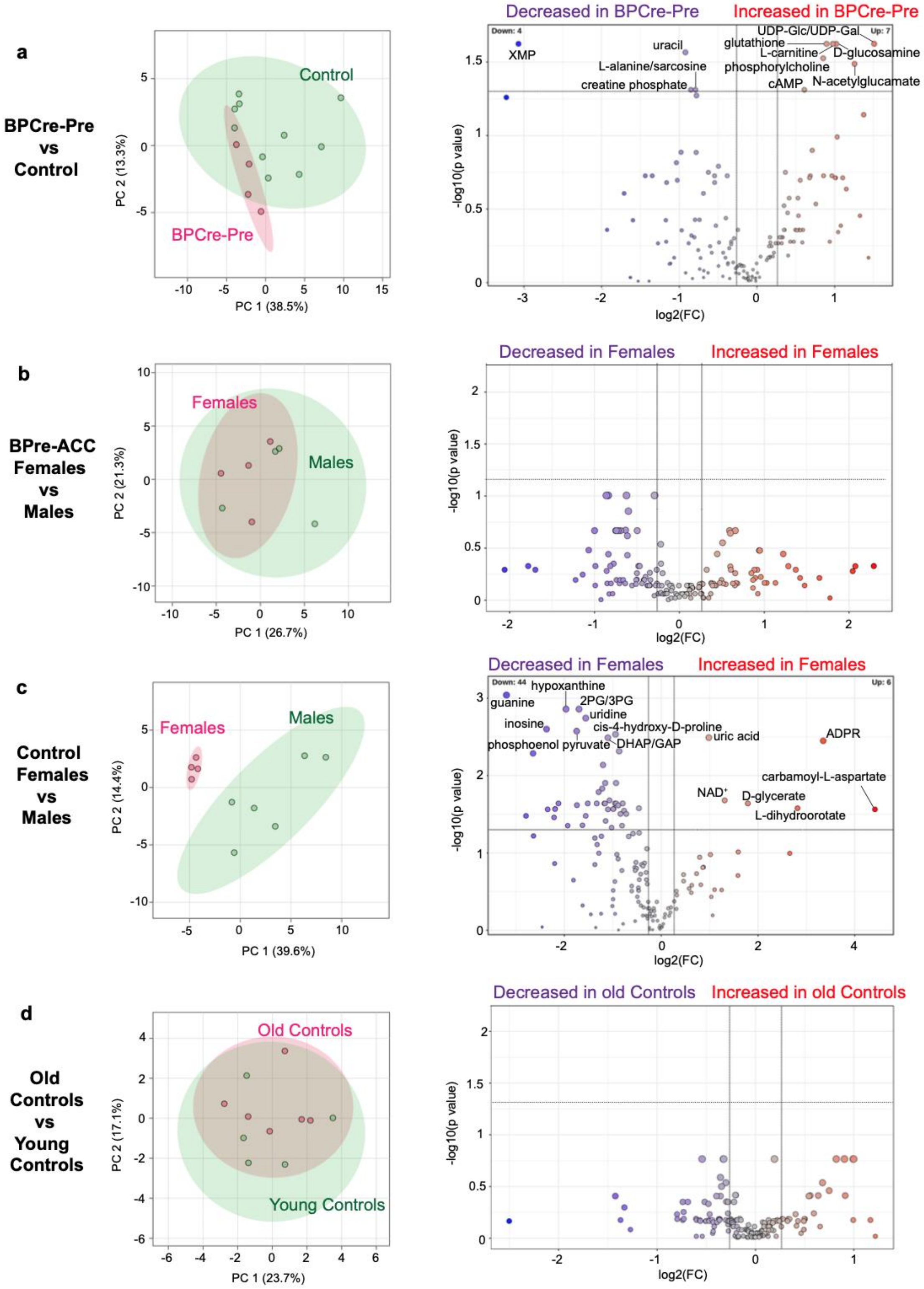


**Suppl. Fig. 3 (related to Fig. 2): Tissue metabolome analysis from Control and BPCre mice, including comparisons by sex and age. a-d** PCA and volcano plots highlighting changes in the polar tissue metabolome of selected groups. For mice without tumors (a, c and d), each sample represents tissue from two adrenals taken from either the same mouse or from two mice of the same genotype. For the volcano plots, metabolites with fold change threshold (x) 1.2 and t-test threshold (y) 0.05 (FDR-corrected p values) are annotated (colored red for increase and purple for decrease). Dashed lines indicate statistical thresholds. **a** Macroscopically normal adrenals from BPCre mice prior to tumor formation (BPCre-Pre) (n=4 samples from male and female mice, 4 months) vs. control adrenals from C57BL/6J mice (n=6 males and n=4 females, 7-11 months). Samples also shown in Fig. 2b. **b** ACC tumors from female and male BPCre mice (n=4 tumors from each sex, 8-13 months) **c** Control adrenals from female and male C57BL/6J mice (n=4 female and n=6 male samples, 7-11 months). **d** Adrenals from control younger (n=5 samples, age 3–5 months) and older (n=6 samples, age 6-10 months) male C57BL/6J mice. Metabolites that could not be unambiguously distinguished are reported using dual annotations. 2PG: 2-phosphoglycerate; 3PG: 3-phosphoglycerate; ADPR: adenosine 5′-diphosphoribose; cAMP, cyclic adenosine monophosphate; DHAP, dihydroxyacetone phosphate; GAP: glyceraldehyde 3-phosphate; NAD, oxidized nicotinamide adenine dinucleotide; UDP: uridine 5’-diphosphate; UDP-Gal: UDP-galactose; UDP-Glc: UDP-glucose XMP, xanthosine monophosphate.

**Suppl. Fig. 4**

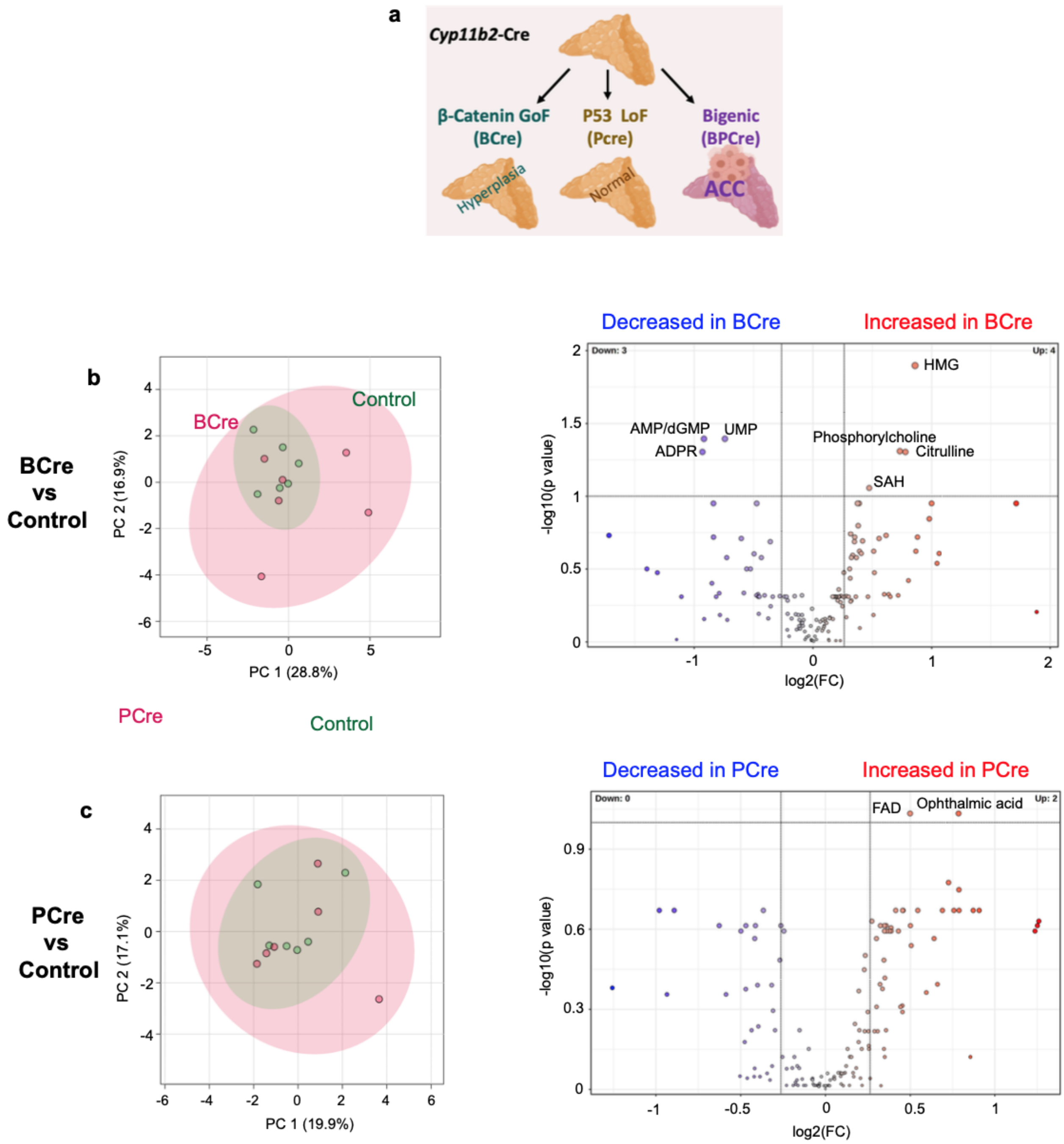
a
Cyp11b2-Cre
β-Catenin GoF
(BCre)
P53 LoF
(Pcre)
Bigenic
(BPCre)
Hyperplasia
Normal
ACC
b
BCre
vs
Control
BCre
Control
PC 2 (16.9%)
PC 1 (28.8%)
Decreased in BCre
Increased in BCre
Down: 3
Up: 4
HMG
AMP/dGMP
UMP
ADPR
Phosphorylcholine
Citrulline
SAH
-log10(p value)
log2(FC)
PCre
Control
c
PCre
vs
Control
PC 2 (17.1%)
PC 1 (19.9%)
Decreased in PCre
Increased in PCre
Down: 0
Up: 2
FAD
Ophthalmic acid
-log10(p value)
log2(FC)

**Suppl. Fig. 4 (related to Fig. 2): Metabolic dysregulation in mice bearing β-Catenin gain-of-function (GOF) mutations or P53 loss-of-function (LOF) mutations. a** Schematic of phenotypes in *Cyp11b2 (AS)*-Cre mouse models. **b,c:** PCA and volcano plots highlighting changes in the polar metabolome of BCre mice (bearing β-GOF mutations) and PCre mice (bearing p53 LOF mutations), as compared to AS-Cre mice (no mutations). Each sample represents tissue from two adrenals taken from either the same mouse or from two mice of the same genotype. For the volcano plots, metabolites with fold change threshold (x) 1.2 and t-test threshold (y) 0.1 (FDR-corrected p values) are annotated (colored red for increase and purple for decrease). Dashed lines indicate statistical thresholds. **b** BCre mouse adrenals (n=6 samples from male mice, 7-11 months) vs. control adrenals from C57BL/6J mice (n=6 samples from male mice, 7-10 months). **c** Adrenals from PCre mice (n=6 samples from male mice, 7-12 months) vs. control adrenals from C57BL/6J mice (n=6 samples from male mice, 7-10 months). Metabolites that could not be unambiguously distinguished are reported using dual annotations. ADPR, adenosine diphosphate ribose; AMP, adenosine monophosphate; dGMP, deoxyguanosine monophosphate; FAD, flavin adenine dinucleotide; HMG, 3-hydroxy-3-methylglutarate; SAH, S-adenosylhomocysteine; UMP, uridine monophosphate.

**Suppl. Fig. 5**

**a**

| Cell line | Species | Mutations | Doubling time (h) |
|---|---|---|---|
| BCH-ACC3 | Mouse | *Ctnnb1* GOF; *p53* LOF | 19.5±1.1 |
| NCI-H295R | Human | *CTNNB1* GOF; *P53* LOF | 61.0±11.0 |
| CU-ACC1 | Human | *CTNNB1* GOF | 76.6±13.7 |
| CU-ACC2 | Human | *P53* LOF; *MSH2* LOF | 52.3±14.9 |

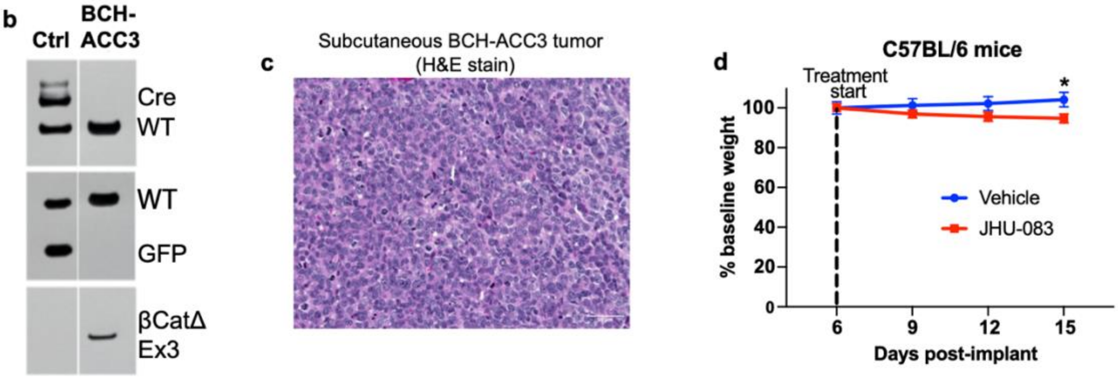


**Suppl. Fig. 5 (related to Fig. 3). ACC cell line description and subcutaneous implant model in C57BL/6 mice. a** Table summarising ACC cell line characteristics. Doubling time (mean, S.E.M.) for each cell line was calculated from 4-10 independent cell proliferation experiments, using the formula Doubling Time (hours) = (72 * ln(2)) / ln(final cell number / initial cell number). **b** PCR analysis showing amplification of AS-Cre and $ROSA^{mT/mG}$ alleles in control samples from an $AS^{Cre/+}::ROSA^{mTmG/+}$ mouse, and loss of $ROSA^{mT/mG}$ and Cre alleles along with deletion of β-catenin exon 3 in BCH-ACC3 cells. **c** Hematoxylin-Eosin staining from BCH-ACC3 subcutaneous implants. **d** Body weight (as % of weight at the beginning of treatment) in C57BL/6 mice treated with JHU-083 or vehicle. JHU-083 was administered at 1 mg/kg from day 6-10 post-implantation and 0.33 mg/kg from days 11-14. N=11,7; *p<0.05 (t-test).

#### Suppl. Fig. 6

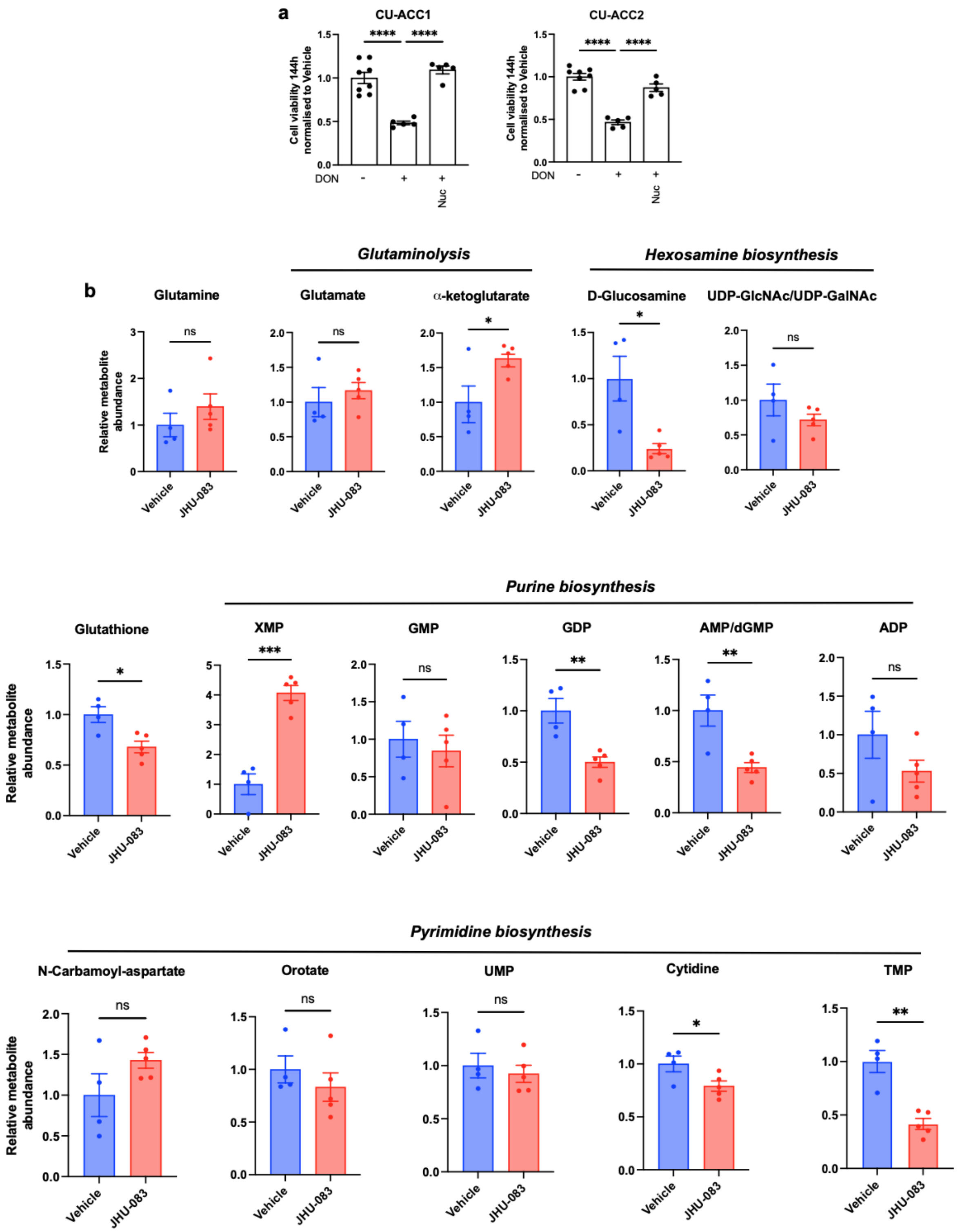


**Suppl. Fig. 6 (related to Fig. 4): Gln antagonism drives nucleotide depletion in ACC. a** Viable CU-ACC1 and CU-ACC2 cell number normalized to vehicle (DON-) after treatment with DON (DON +), alone or supplemented with the indicated metabolites. DON was administered at 5 μM for 144h. Nuc: Embryomax nucleoside cocktail 1x. Data representative of three independent experiments (5-8 technical replicates per experiment). ****p<0.0001, one-way ANOVA followed by Tukey's test for multiple comparisons between DON monotherapy and each other group. **b** Relative abundance of selected metabolites in BCH-ACC3 SQ mouse implants after 48h treatment with JHU-083 or vehicle. N=5,4; ns: not significant, *p<0.05, **p<0.01, ***p<0.001, t-test. Metabolites that could not be unambiguously distinguished are reported using dual annotations. ADP, adenosine diphosphate; AMP, adenosine monophosphate; dGMP, deoxyguanosine monophosphate; GDP, guanosine diphosphate; GMP, guanosine monophosphate; TMP, thymidine monophosphate; UDP, uridine diphosphate; UDP-GalnNAc: UDP-N-Acetylgalctosamine; UDP-GlcNAc: UDP-N-acetylglucosamine; UMP, uridine monophosphate; XMP, xanthosine monophosphate.

**Suppl. Fig. 7**

**a**

***Purine biosynthesis***

***PFAS***

| | Logrank p | HR |
|---|---|---|
| ACC | 0.046 | 2.2 |
| LIHC | 0.038 | 1.4 |
| MESO | 0.006 | 2 |
| SARC | 0.023 | 1.6 |
| SKCM | 0.007 | 1.4 |
| UVM | 0.068 | 2.3 |

***GMPS***

| | Logrank p | HR |
|---|---|---|
| ACC | <0.001 | 4.3 |
| KIRP | 0.010 | 2.3 |
| LGG | 0.002 | 1.8 |
| LIHC | 0.004 | 1.7 |
| LUAD | 0.004 | 1.6 |
| MESO | 0.006 | 2 |
| PAAD | <0.001 | 2.1 |
| PCPG | 0.016 | 5.8 e08 |

***PPAT***

| | Logrank p | HR |
|---|---|---|
| KICH | 0.012 | 9 |
| LGG | <0.001 | 2 |
| LIHC | 0.008 | 1.6 |
| MESO | 0.007 | 2 |
| SARC | <0.001 | 2.1 |
| THCA | 0.002 | 7.3 |

***Pyrimidine biosynthesis***

***CAD***

| | Logrank p | HR |
|---|---|---|
| ACC | 0.002 | 3.7 |
| BLCA | 0.043 | 1.4 |
| KIRC | <0.001 | 1.7 |
| KIRP | 0.016 | 2.2 |
| LAML | 0.046 | 1.8 |
| LGG | 0.043 | 1.4 |
| LIHC | <0.001 | 1.9 |
| MESO | 0.003 | 2.1 |

***CTPS1***

| | Logrank p | HR |
|---|---|---|
| ACC | 0.0024 | 2.5 |
| LGG | 0.00065 | 1.9 |
| MESO | 0.0038 | 2 |
| SKCM | 0.00021 | 1.7 |
| UVM | 0.01 | 3.4 |

**b**

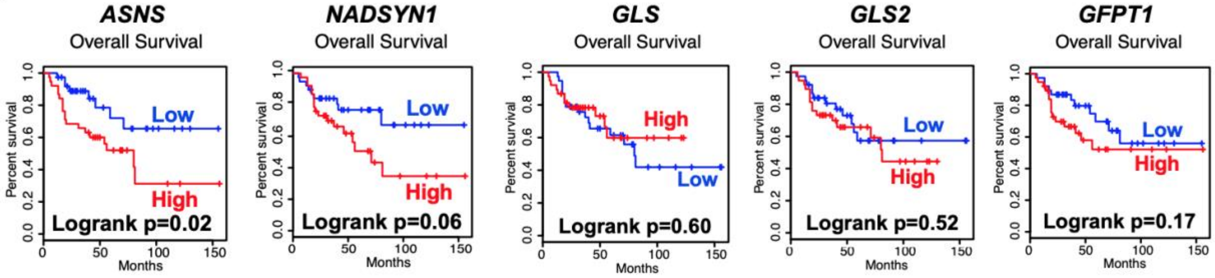


**Suppl. Fig. 7 (related to Fig. 4): Associations between Gln-metabolising enzyme expression and survival (TCGA database). a** Cancer types with significant (logrank p<0.05) association between high enzyme gene expression levels and poor prognosis, based on The Cancer Genome Atlas (TCGA). HR: Hazard ratios. BLCA: Bladder urothelial carcinoma, KICH: kidney chromophobe cell carcinoma, KIRC: kidney renal clear cell carcinoma, KIRP: kidney renal papillary cell carcinoma, LAML: acute myeloid leukemia, LGG: low-grade glioma, LIHC: hepatocellular carcinoma, LUAD: lung adenocarcinoma, MESO: mesothelioma, PAAD: pancreatic adenocarcinoma, PCPG: pheochromocytoma and paraganglioma, SARC: sarcoma, SKCM: skin cutaneous melanoma, THCA: thyroid carcinoma, UVM: uveal melanoma. **b** Survival curves comparing overall survival in ACC patients with high (top 50%) versus low (bottom 50%) gene expression levels of the indicated Gln-metabolizing enzymes, based on

The Cancer Genome Atlas (TCGA). Significance and hazards ratio were assessed by Log-rank (Mantel-Cox) tests.

**Suppl. Fig. 8**

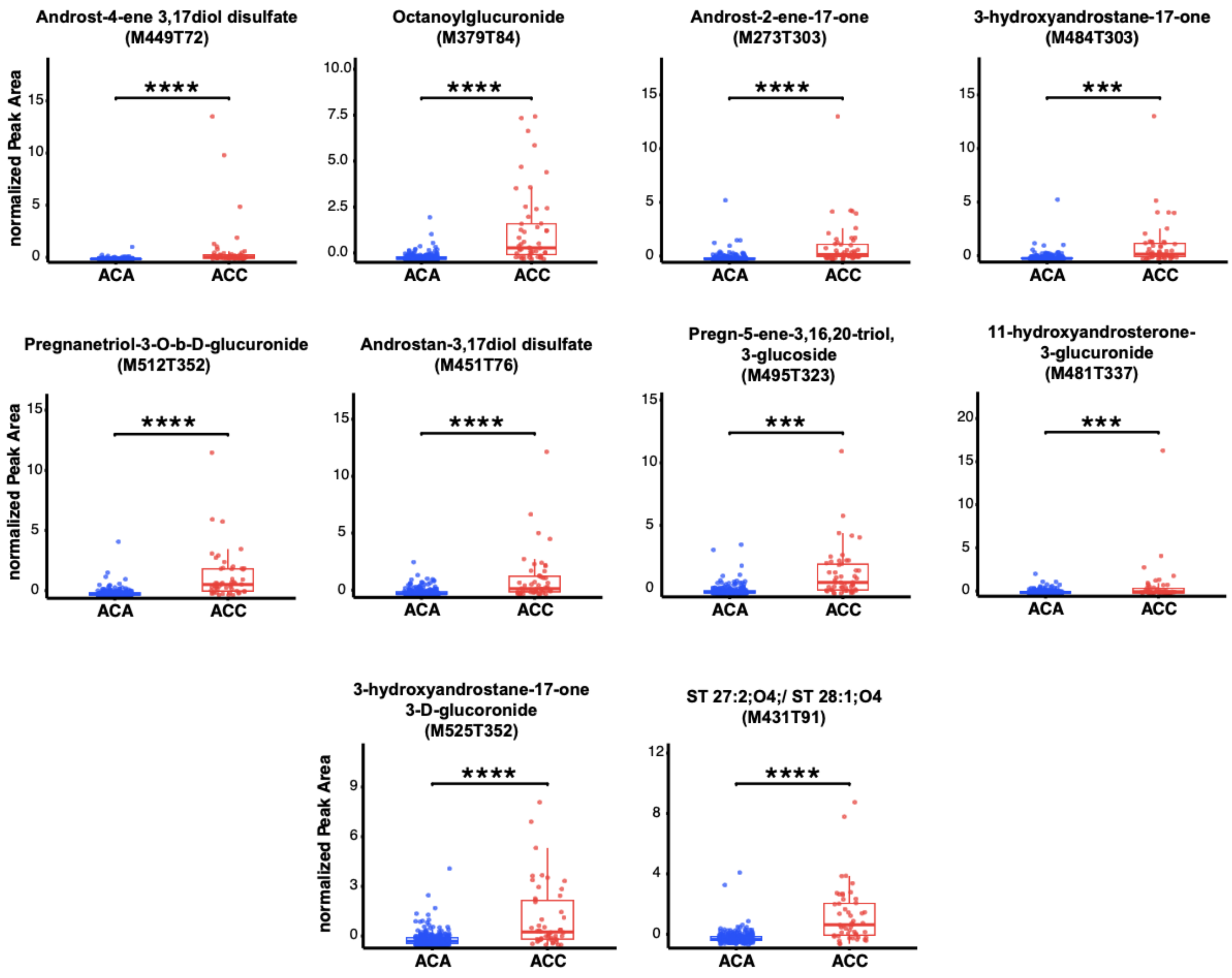


**Suppl. Fig. 8 (related to Fig. 6): Top 10 most up-regulated metabolites in the serum of patients with ACC.** Box and whiskers plots comparing relative serum abundance of the 10 most increased metabolites in patients with ACC vs ACA controls (n=54, 291). The box edges represent the 1st and 3rd quartile and the upper and lower whiskers represent Q3 + 1.5x Interquartile range (IQR) and Q1 - 1.5x IQR, respectively. The median is depicted as the horizontal line within the box. ***p<0.001, ****p<0.0001 (t-test).

#### Supplementary Table Legends

**Suppl. Table 1.** Significantly up-regulated pathways (Benjamini-Hochberg-corrected FDR<0.05) in murine (BPCre) ACC (A) and human ACC (B). Significantly up-regulated genes are provided for each pathway.

**Suppl. Table 2.** Significantly down-regulated pathways (Benjamini-Hochberg-corrected FDR<0.05) in murine (BPCre) ACC (A) and human ACC (B). Significantly down-regulated genes are provided for each pathway.

**Suppl. Table 3.** Significantly dysregulated (Benjamini-Hochberg-corrected FDR<0.05 and >1.2 fold-change) polar metabolites in BPCre-ACC tissue, as compared to Control (normal) adrenals.

**Suppl. Table 4.** Significantly dysregulated (Benjamini-Hochberg-corrected FDR<0.05) serum metabolic features in patients with ACC, as compared to ACA. CE: cholesterol ester; Cer: Ceramide; CL: cardiolipin; DG: diacylglyceride; FA: fatty acid; HexCer: hexose-ceramide; LPA: lysophosphatidic acid; LPC: lysoglycerophosphocholine; LPE: lysoglycerophosphoethanolamine; LPG: lysoglycerophosphoglycerol; LPI: lysoglycerophosphoinositol; LPS: lysoglycerophosphoserine; PC:glycerophosphocholine; PE:glycerophosphoethanolamine; PG: glycerophosphoglycerol; PI: glycerophosphoinositol; PS: glycerophosphoserine; SHexCer: Sulfoglycosphingolipids; SM: sphingomyelin; ST: sterol lipid; TG: triacylglyceride

**Suppl. Table 5.** Significantly dysregulated (FDR<0.05) metabolic pathways based on analysis of significantly dysregulated polar serum metabolites in patients with ACC vs ACA (with reference to the KEGG database). Significant metabolites in each pathway are highlighted in red font.
